# Revisiting the burden borne by fumarase: enzymatic hydration of an olefin

**DOI:** 10.1101/2022.09.03.506457

**Authors:** Asutosh Bellur, Soumik Das, Vijay Jayaraman, Sudarshan Behera, Arpitha Suryavanshi, Sundaram Balasubramanian, Padmanabhan Balaram, Garima Jindal, Hemalatha Balaram

## Abstract

Fumarate hydratase (FH) is a remarkable catalyst that decreases the free energy of the catalyzed reaction by 30 kcal mol^−1^, much larger than most exceptional enzymes with extraordinary catalytic rates. Two classes of FH are observed in nature: class-I and class-II, that have different folds, yet catalyze the same reversible hydration/dehydration reaction of the dicarboxylic acids fumarate/malate, with equal efficiencies. Using class-I FH from the hyperthermophilic archaeon *Methanocaldococcus jannaschii* (Mj) as a model along with comparative analysis with the only other available class-I FH structure from *Leishmania major (*Lm), we provide insights into the molecular mechanism of catalysis in this class of enzymes. The structure of MjFH apo-protein has been determined, revealing that large inter-subunit rearrangements occur across apo- and the holo-protein forms, with a largely preorganized active site for substrate binding. Site-directed mutagenesis of active site residues, kinetic analysis and computational studies including DFT and natural population analysis, together show that residues interacting with the carboxylate group of the substrate play a pivotal role in catalysis. Our study establishes that an electrostatic network at the active site of class-I FH, polarizes the substrate fumarate through interactions with its carboxylate groups, thereby permitting an easier addition of a water molecule across the olefinic bond. We propose a mechanism of catalysis in FH that occurs through transition state stabilization involving the distortion of the electronic structure of the substrate olefinic bond mediated by the charge polarization of the bound substrate at the enzyme active site.

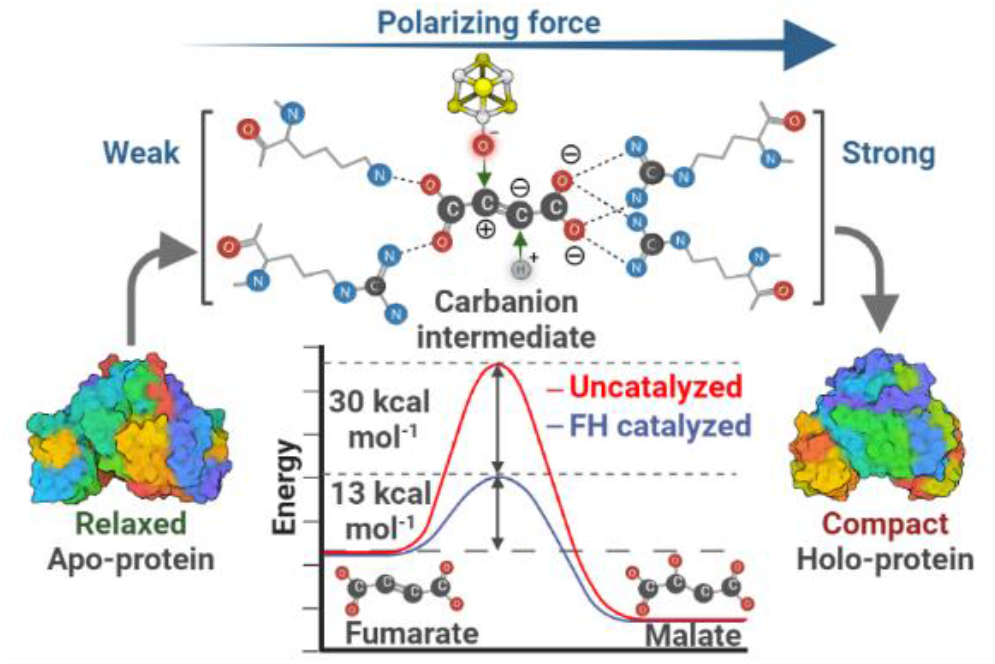

## INTRODUCTION

Fumarate, a symmetric olefinic dicarboxylic acid, is an important intermediate in the tricarboxylic acid (TCA, Krebs’) cycle. In living cells, fumarate is efficiently and stereospecifically converted to L-malate in a hydration reaction, catalyzed by the enzyme fumarate hydratase (fumarase, FH, E.C. 4.2,1.2). Yet, in organic chemistry, olefin hydration presents many challenges,^1–3^ leading to the provocative conclusion that “*enzymes are the best chemists*”.^1^ On the scale of ‘enzyme proficiency’, which is the ratio of the rates of the enzyme catalyzed and uncatalyzed reaction in aqueous solution, fumarase (FH) ranks high with a 10^15^-fold rate enhancement in the enzymatic reaction. The 36 kcal mol^−1^ activation barrier, that must be surmounted for catalysis, has been described, by Bearne and Wolfenden, as the ‘*burden borne by fumarase*’.^4^ The biochemical equilibrium in the reversible hydration reaction in the TCA cycle lies in the direction of malate (K_eq_=4, first measured by Krebs),^5,6^ suggesting that the rates of hydration are 4-fold higher than the rates of dehydration.^7^ How does the enzyme lower this formidable activation barrier for this deceptively simple reaction? While kinetic and thermodynamic analyses have provided valuable insights, the molecular mechanisms remain to be clarified.^7,8^

Fumarate hydratases (FH) are distinctly divided into two classes, class-I and class-II, based on the presence or absence of a 4Fe-4S cluster, respectively.^9^ Both classes of FH are ubiquitous across all kingdoms of life. While, class-II enzymes occur predominantly in eukaryotes, class-I enzymes are widely distributed across archaea and prokaryotes, with sparse representation in eukaryotes.^10^ These two enzymes share no sequence or structural similarity, yet catalyze the same reaction with close to equal efficiencies.^4,11^ There is support for the chemical mechanism in class-II fumarases taking place through all three possible pathways: carbonium (E1),^12^ concerted (E2)^13^ and carbanion (E1cB)^14^ pathway (**Scheme 1**), with the weight of evidence tilting in favor of the carbanion pathway.^15^ Class-II FH has been placed under the aspartase/fumarase family of enzymes, which share a common fold. Structural and mechanistic studies have focused on the malate dehydration reaction, with discussions centered on general acid-base catalysis. A conserved serine residue is believed to be the catalytic base while the catalytic acid has remained elusive in these class of enzymes.^16^ Extensive kinetic isotope experiments in class-II FH have revealed that the rate limiting step of the reaction does not involve a carbon-hydrogen bond breakage or formation. The rate limiting step is thought to be either the product release or proton exchange alongside a slow conformational isomerization.^17^

Unlike class-II fumarases, a comprehensive understanding of enzyme chemistry and catalysis is lacking for class-I FHs due to their thermolabile and oxygen-sensitive nature. Class-I FHs are further divided into single-subunit and two-subunit types, depending on the number of genes coding for the functional enzyme. Two-subunit proteins are found only in prokaryotes and archaea, but not in eukaryotes. The alpha (α) and beta (β) subunits of the two-subunit type class-I FH correspond to the N-terminal and C-terminal domains (NTD and CTD) of single-subunit class-I FH.^18^ Both these enzymes have high sequence similarity and a small polypeptide insertion between the two subunits is the only distinguishing feature between them. Single-subunit class-I FH have been biochemically characterized in a handful of organisms.^10,19–25^ Thus far, only two-subunit Class-I FH from *Pelotomaculum thermoprpionicum*^18^ *and Pyrococcus furiosus*^26^ have been biochemically characterized. The recent report of the structure of a substrate bound class-I FH from *Leishmania major* is a major step forward in probing the molecular mechanism of the fumarase reaction.^25^

Transition state stabilization in the fumarase reaction is driven by both enthalpic and entropic factors (ΔH=−24 kcal mol^−1^ and ΔS= 19 cal mol^−1^ K^−1^).^4^ The large enthalpic stabilization must arise from the electrostatic interactions of the tri-negative aci-carboxylate anion at the substrate binding site, while the positive entropic gain can arise from the liberation of the water solvent shell around the fumarate dianion upon binding to the enzyme active site. Bearns and Wolfenden concluded over 25 years ago that the “*nature of enzyme - ligand interactions will be clearer when structural information about this remarkable catalyst becomes available*”.^4^ In this report, we compare the structure of the apo form of the thermostable, two-subunit FH from *Methanocaldococcus jannaschi* with *Leishmania major* FH (MjFH and LmFH), probe mutational effects on catalysis using several mutant enzymes and report DFT calculations to gain insights into the mechanism of the fumarase reaction. Our results provide strong support for electrostatic stabilization at the active site of a charge polarized, *aci*-carboxylate form of fumarate as the transition state in the hydration reaction.

**Scheme 1.**
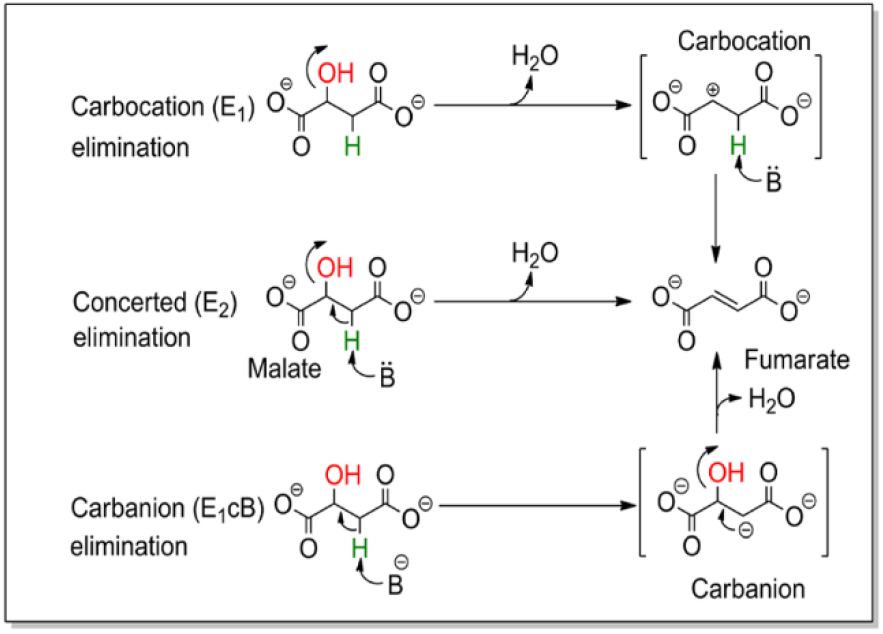
Mechanisms for dehydration of malate to fumarate. E_1_ and E_1_cb mechanisms proceed through a two-step reaction involving a carbocation and a carbanion intermediate, respectively, while the E_2_ mechanism is a one-step reaction that proceeds without an intermediate with simultaneous removal of proton and elimination of water molecule.

## EXPERIMENTAL PROCEDURES

### Generation of plasmid constructs

The genes for the subunits of MjFHα and MjFHβ were amplified by PCR from *M. jannaschii* genomic DNA (procured from ATCC) using primers listed in **Table S1** and cloned into two tandem multiple cloning sites of pET-Duet vector (Novagen, Merck), using the restriction enzymes BamHI and SalI for MjFHα subunit, and NdeI and XhoI for MjFHβ subunit such that they can be co-expressed from the same plasmid (pETduet_MjFHαβ). For expressing the individual subunits separately, the same primers and restriction sites were used. Site-directed mutants were generated using the PCR-driven overlap extension method using pETduet_MjFHαβ as a template. DNA fragments with the required mutations, generated using appropriate oligonucleotides were then assembled into the plasmid by ligation-dependent cloning.^27^ All the clones were confirmed by DNA sequencing.

### Protein expression, purification, and reconstitution of iron-sulfur cluster

Recombinant MjFHαβ was expressed in the *E. coli* strain BL21(DE3)-RIL carrying the plasmid pETduet_MjFHαβ. Cells were induced with 0.3 mM IPTG and grown for 18 hours at 16 °C followed by lysis using French press (Thermo IEC Inc., USA) at 1000 psi. The lysate was incubated at 70 °C for 30 minutes to precipitate the bacterial proteins followed by addition of 0.01% PEI to precipitate nucleic acids. Recombinant protein was purified using Q-sepharose anion-exchange column, concentration determined by Bradford method^28^ and stored at −80 °C as aliquots. Similar protocol was adopted for purification of all the mutants. Selenomethinone incorporated MjFHαβ was purified in a manner similar to that of the WT. Growth conditions for cells expressing WT and selenomethionine incorporated proteins and details of protein purification protocol are provided in supporting infromation.

Purified protein aliquots stored at −80 °C were reconstituted with Fe-S cluster following standard protocols.^29,30^ The protein solution was incubated in anaerobic chamber (Coy, USA) for 1 hour to remove all traces of oxygen. All the subsequent steps were performed under anaerobic conditions in the chamber. Briefly, around 10-50 μM protein was used for reconstitution. The reconstitution procedure was initiated by the addition of 50-fold excess DTT to the protein solution followed by stirring for 30 min. Subsequently, 10-fold excess ferrous ammonium sulfate and sodium sulfide were added and stirred for another 3.5 hours.

### Interaction of MjFHα and MjFHβ subunits

Interaction of MjFHα subunit with the β-subunit was checked by incubating α-subunit bound nickel-Sepharose beads with an excess of β-subunit at 26 °C for 30 min in buffer containing 20 mM Tris-HCl, pH 7.4. Following this, beads were washed with the same buffer and α-subunit was eluted in same buffer containing 500 mM imidazole. An aliquot of the eluate was loaded onto SDS-PAGE for separation and stained with Coomassie Brilliant Blue (CBB) for visualization. Oligomeric state of co-purified protein was probed by size-exclusion chromatography on an analytical Superdex 200 10/300 GL column. Details of the buffer and standards used are provided in supporting information.

The thermodynamics of the interaction of MjFHα and β-subunits and the dissociation constant (K_d_) were estimated by isothermal calorimetry using a VP-isothermal titration calorimeter (MicroCal VP-ITC machine). 681 μM MjFHβ was taken in the syringe with 75 μM MjFHα kept in the cell. 30 injections of 10 μl each of MjFHβ were added to the sample cell containing MjFHα and the cell contents were stirred at 300 rpm throughout the titration. The titration was performed at a constant temperature of 25 °C. Data were processed using Origin software and the best fit model was used for K_d_ estimation.

### Characterization of Fe-S cluster

Metal-thiolate charge transfer band was detected in the visible region using a 20 μM solution of 4Fe-4S cluster reconstituted protein taken anaerobically in a 1 cm path length cuvette, sealed, and used for acquiring CD spectra in the region 250 to 600 nm using a spectropolarimeter (Jasco J-810).

### Enzyme activity

All initial velocity measurements were conducted at 70 °C using a spectrophotometer (Hitachi U2010 spectrophotometer) and initiated by the addition of enzyme. The activity of MjFH was found to be maximal at pH 7.2 and hence all assays were carried out at this pH in 50 mM Tris-HCl. Conversion of fumarate to malate was measured spectrophotometrically as a decrease in absorbance caused by the depletion of fumarate. Details of the wavelengths and extinction coefficient values used are provided in supporting information. Kinetics of inhibition by *RS*-2-thiomalate and details of analysis of initial velocity plots of WT and mutants are provided in supporting information.

### Crystallization and data collection

MjFHα, MjFHβ, MjFHαβ apo-protein and holo-protein were set up for crystallization using the Microbatch method.^31^ Details of the methods adopted for setting up crystal trays are provided in supporting information. Crystals of MjFHβ were obtained in 0.1 M Bis-Tris, pH 5.5, and 25 % polyethylene glycol-3350 (PEG-3350). Crystals of MjFHαβ apo-protein were obtained in 0.2 M magnesium chloride hexahydrate, 0.1 M BIS-TRIS, pH 5.5, 25% w/v polyethylene glycol 3,350. Selenomethionine incorporated MjFHαβ (Se-MjFHαβ) also crystallized in the same condition, but crystals were not well formed. Well-formed crystals were obtained in the presence of an additive, non-detergent sulfobetaines (0.3 M NDSB-195) obtained from Hampton additive screen™. Crystals were cryo-protected by placing them for a brief period, in the initial condition with 20% (v/v) glycerol, before collecting diffraction data.

X-ray diffraction data on MjFHβ protein crystal was collected on a Rigaku RU200 X-ray diffractometer equipped with a rotating anode type light source with an osmic mirror that gives a monochromatic light source of wavelength 1.54179 Å. Selenium-SAD data was collected on Se-MjFHαβ crystals at a wavelength of 0.9787 at the BM-14 beamline of European Synchrotron Radiation facility (ESRF).

### Structure determination and refinement

Crystal structure of MjFHβ was solved by molecular replacement method by making use of *Archaeoglobus fulgidus* FH β-subunit (PDB ID: 2ISB) as the template for MR. Crystal structure of Se-MjFHαβ was solved by selenium-SAD, making use of the eleven anomalously scattering selenium atoms per functional dimer. Various packages available in CCP4 suite^32^ and Phenix modules^33^ were used. Briefly, iMOSFLM^34^ was used for data processing, SCALA^35^ for data reduction and scaling, PHASER^36^ for phasing in MjFHβ structure solution and PHENIX AutoSol^33^ for determining the position of selenium atom sites in Se-MjFHαβ. Automatic model building was performed for Se-MjFHαβ using PHENIX AutoBuild.^37^ REFMAC 5.0^38^ and Phenix.refine^39^ were used for refinement. Manual refinements were carried out using COOT^40^. Refinement statistics are summarized in **Table S2**. All the structure-related figures were created using PyMol software.^41^ Electrostatic surface potential was calculated using Adaptive Poisson-Boltzmann Solver plugin in PyMol.^42^

### Molecular dynamics simulations

Atomistic molecular dynamics simulations have been performed on five enzyme forms: (i) holo-LmFH (employing the crystal structure with PDB ID-5l2r, refer to supporting information and **Figure S5** for more details), (ii) apo-LmFH (after removing the 4Fe-4S cluster and substrate from the crystal structure of holo-LmFH), (iii) apo-MjFH (using the crystal structure obtained in this study), (iv) a model of holo-MjFH-1 (after docking the cluster and substrate into the active site of apo-LmFH), and (v) a model of holo-MjFH-2 (employing the contracted holo-MjFH structure obtained from a metadyanamics simulation^43^ (refer to supporting information, and **Figure S8 and S9** for more details). Simulations were carried out on the dimer forms of these enzymes at T=300K and P=1bar. The details on the simulation setup and simulation parameters are presented in the supporting information.

CHARMM36m force field^44^ was employed for the enzyme, whereas force field parameters for malate were generated using the CGenFF server.^45^ Force-field parameters for the 4Fe-4S cluster, and the cysteine residues bound to it were taken from a previous report.^46^ GROMACS-2018.3^47^ was used for all the MD simulations employing three-dimensional periodic boundary conditions. Refer to **Table S6** for number of atoms, cubic box length, and simulation lengths for the various simulated systems.

### Quantum Chemical Calculations

All the quantum chemical calculations were carried out using Gaussian 16 software.^48^ Geometry optimizations were performed using B3LYP functional along with the split valence Pople’s type 6-31G(d,p) basis set ^49–52^ for all atoms except Fe. For metal such as Fe, we chose LANL2DZ basis set with an effective core potential (ECP).^53^ Hessian calculations were carried out to characterize the stationary points as first order saddle points and the presence of a unique imaginary frequency was used to verify the transition states (TSs) for the small model systems. To further ascertain the correctness of the TS, intrinsic reaction coordinate (IRC) calculations were performed to identify the connecting reactant and product.^54–57^ For all stationary points, single point calculations were carried out using a higher basis set, 6-311+G** along with the D3 version of Grimme’s dispersion correction.^58^ To include the effect of reaction medium, single point calculations also included the solvent effect using Cramer and Truhlar’s solvation model density (SMD) model^59^ with water (ε=80.4) as the solvent for small systems and diethyl ether (ε=4.3) for QM cluster approach.^60–65^ In the QM cluster approach, quantum chemical methods, usually density functional theory (DFT), are used to treat a well-chosen region around the enzymatic active site while the rest of the enzyme is approximated as a homogeneous polarizable medium. To avoid any conformational changes, the C_α_ atoms are frozen at their crystallographic positions. This might result in a few imaginary frequencies below 60 cm^−1^, which do not affect the zero point energies (ZPE) much. The residues are typically truncated at C_α_, and the truncated bonds are saturated by hydrogens. The ZPE, thermal, and entropic corrections calculated at 298.15 K and 1 atm pressure obtained in the gas phase at the B3LYP/6-31G(d,p)//6-31G(d,p),LANL2DZ (Fe) level of theory are added to the “bottom-of-the-well” energies obtained from the single-point energy calculations at the SMD(water)/B3LYP/6-311+G**//SMD(diethylether)/B3LYP/6-311+G**,LANL2DZ(Fe) level of theory. It should be noted that for the QM cluster approach we only add the ZPE and thermal corrections since entropy calculations with frozen atoms are unreliable. All the energies discussed here are those obtained at the SMD(water)/B3LYP/6-311+G**//SMD(diethylether)/B3LYP/6-311+G**,LANL2DZ(Fe) level of theory unless mentioned otherwise. For the QM cluster, NPA (Natural Population Analysis) calculations were further carried out at SMD_(diethylether)_/B3LYP/6-311+G**,LANL2DZ(Fe) level of theory using the gas-phase optimized geometries at B3LYP/6-31G(d,p),LANL2DZ(Fe) level of theory. (See **SI** for full computational details).

## RESULTS

### Structure of MjFH

MjFHα and MjFHβ subunits when co-expressed in *E. coli*, coelute upon purification by ion-exchange chromatography on a Q Sepharose column. Pull down of MjFHβ using MjFHα as bait on Ni-NTA Sepharose beads confirmed formation of protein complex (**Figure 1A)**. Analytical size-exclusion chromatography of MjFHα, MjFHβ and MjFH (αβ complex) showed a molecular mass of 60, 22 and 93 kDa corresponding to a homodimer, monomer and a heterotetramer of 2α and 2β subunits, respectively (theoretical mass of α = 33 kDa, β = 22 kDa) (**Figure 1B)**. The 2α – 2β subunit stoichiometry of the complex was further verified through isothermal calorimetry, where the resultant thermogram (**Figure 1C**) fitted best to one-site binding model and the K_d_ value of 595 ± 19 nM indicated tight interaction between the α and β subunits. Visible CD, sensitive to changes in Fe-S cluster environment in proteins,^66^ did not show any circular dichroic effect in the visible region for the unreconstituted, apo-protein while, the 4Fe-4S reconstituted MjFHαβ showed a characteristic spectrum in the visible region **(Figure 1D)**.

**Figure 1.**
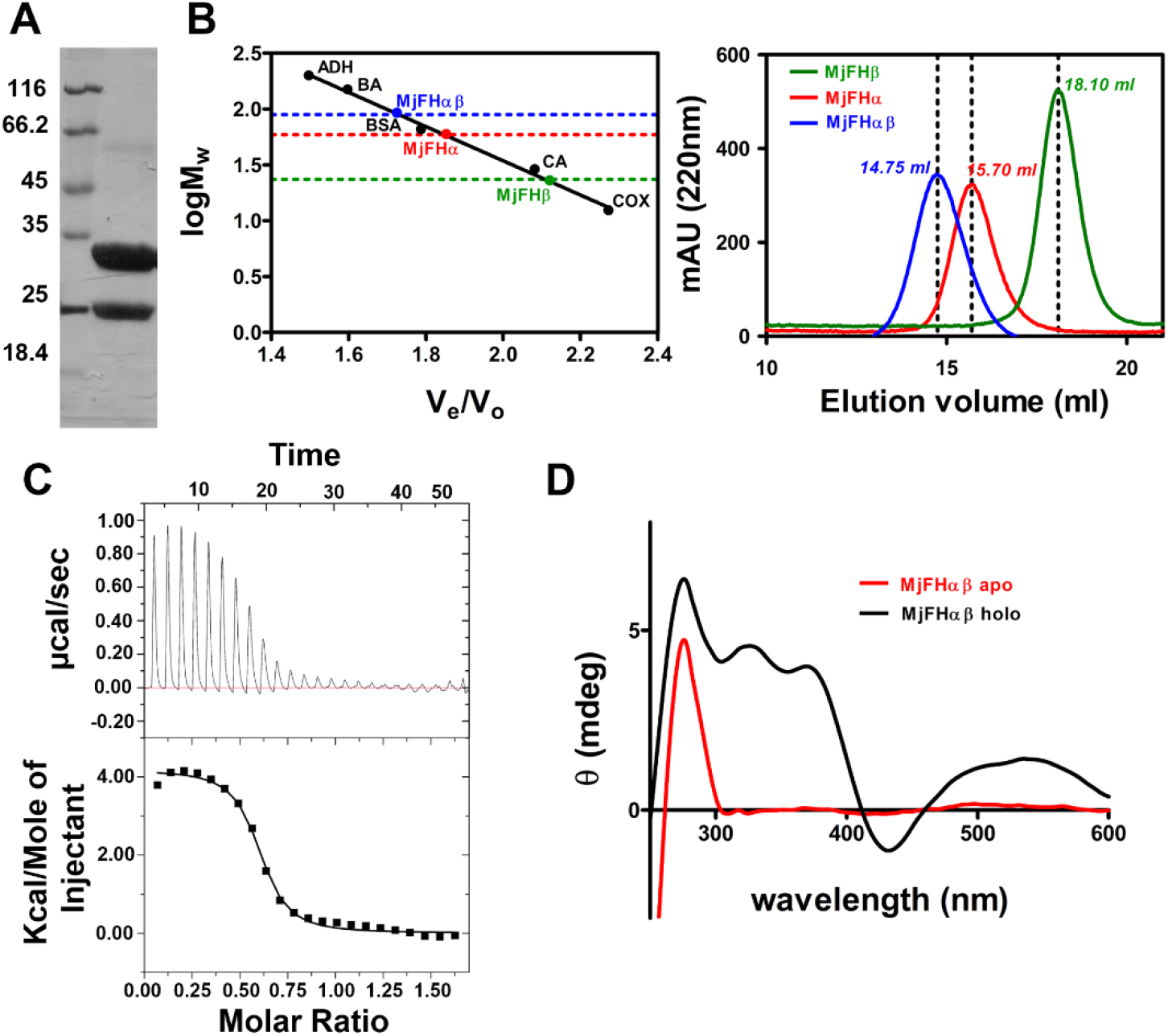
Purification and characterization of *M. jannaschii* fumarate hydratase. ***A***. α and β-subunits associate *in vitro*. SDS-PAGE of 500 mM imidazole eluate from Ni-NTA Sepharose matrix (lane2). The (His)_6_-tagged α-subunit of MjFH was bound to Ni-NTA Sepharose beads followed by incubation with β-subunit and washing to remove unbound protein. Thereafter, protein bound to Ni-NTA beads was eluted with imidazole. Lane 1, molecular mass marker with molecular mass in kDa indicated to the left of the panel. **B**. Size-exclusion chromatograms (right panel) of α-, β- and αβ-subunits of MjFH show that MjFH α is a constitutive dimer; MjFHβ a monomer and MjFHαβ a dimer of a heterodimer (2α+2β). Left panel shows calibration plot of protein standards along with elution volumes of the peaks marked in the right panel. **C**. Thermogram obtained by titration of MjFHα with MjFHβ in an isothermal calorimeter. **D**. Visible circular dichroism spectra of apo-MjFH and Fe-S reconstituted holo-MjFH. Panel shows the characteristic dichroic pattern of Fe-S cluster in holo-MjFH.

Attempts were made to crystallize individual subunits of MjFH and the complex in both apo- and holo-protein state. In all conditions attempted, only MjFHβ and MjFHαβ apo-protein complex crystallized. MjFHβ subunit structure was solved by molecular replacement using the β-subunit structure of *Archaeoglobus fulgidus* (PDB ID: 2ISB) and refined to a resolution of 2.3 Å (**Table S2)**. Structure of MjFHαβ apo-protein complex was solved by single-wavelength anomalous dispersion (SAD) using crystals of the selenium incorporated protein and refined to 2.45 Å resolution (**Table S2)**. MjFHβ forms a crystallographic dimer that does not have any physiological relevance as analytical size exclusion chromatography shows only monomers in solution. The asymmetric unit in MjFHαβ complex contains one copy each of α- and β-subunits and the functional form of the enzyme which is a dimer of a heterodimer (**Figure 2A, B**) was generated using a symmetry related mate from the neighboring unit cell (**Figure 2D**). MjFHβ structure overlays well on the β-subunit of MjFHαβ apo-protein complex with an RMSD of 0.528 Å. A 14-residue stretch at the C-terminal end of the protein that was disordered in the MjFHβ structure ordered into an α-helix in the MjFHαβ complex (**Figure 2C**). Ordering of the 14-residue α-helical stretch is aided by a helix-helix interaction with the α-subunit **(Figure S1)**. This 14-residue helical stretch is absent in the class-I single subunit *Leishmania major* FH (LmFH) and may possibly enable tight binding of the two subunits. LmFH structure is shown to have a previously unidentified α + β fold and searching MjFHα in DALI server to identify the fold, picked LmFH structure as the strongest match with an RMSD of 2.1 Å. The SCOP database^67^ classifies the fold of MjFHβ as a “swiveling β/β/α” similar to the CTD in LmFH, which is considered to be a highly mobile segment in multi-domain proteins. Surface electrostatic potential of MjFH reveals a large positively charged active site cavity between the α and β-subunits of MjFH akin to that between NTD and CTD of LmFH (**Figure 2E**). A tunnel originating from a cavity at the top of the protein, observed in LmFH as well, passes through the entire breadth of the protein without connecting to the active site cavity (**Figure 2F**).

**Figure 2.**
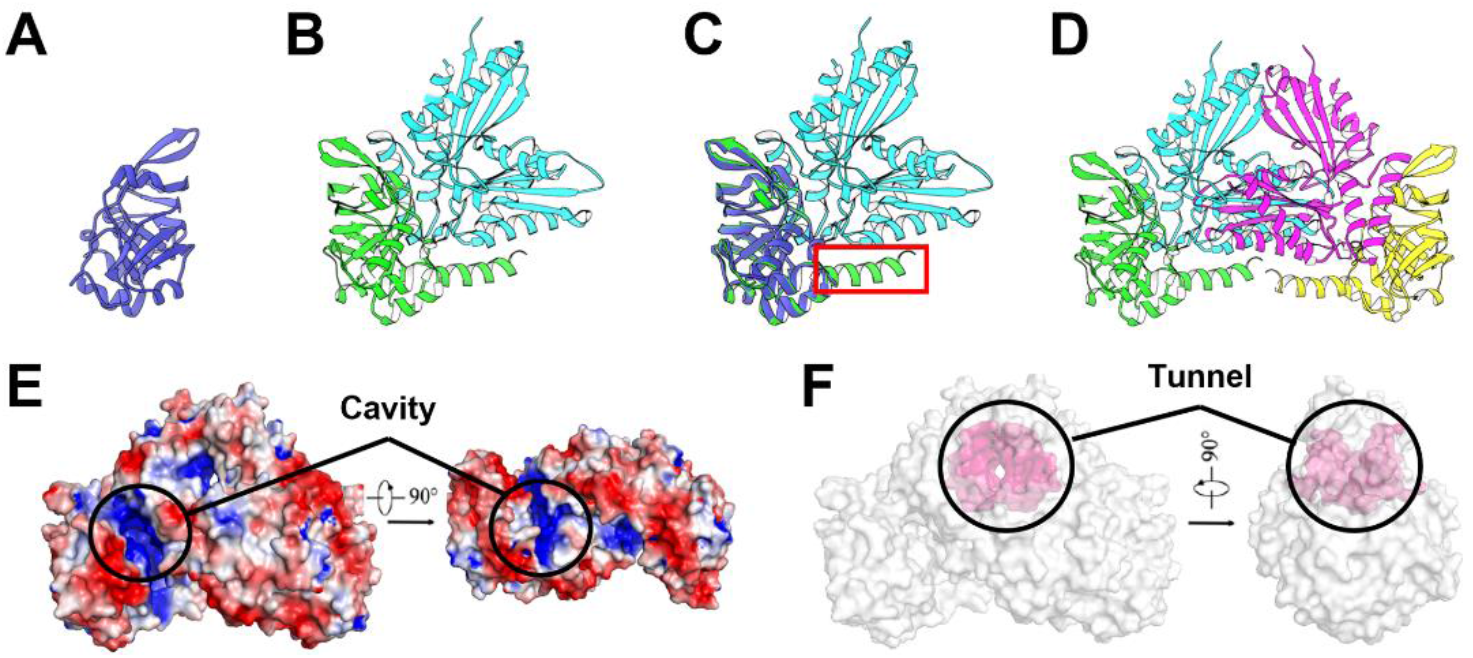
Structure of MjFH. A, B and D. Structure of MjFHβ monomer,MjFHαβ dimer and MjFHαβ biological assembly of tetramer. α-subunits are in cyan and magenta color, while β-subunits are in green and yellow color for the complex and dark blue for MjFHβ structure. **C**. Overlay of MjFHβ monomer on MjFHαβ dimer reveals ordering of the C-terminal α-helix (enclosed in red box) in β-subunit of MjFHαβ complex, which is disordered in MjFHβ structure. **E**. Surface electrostatic potential of MjFHαβ tetramer. A positively charged cavity is located between the interface of α and β-subunits creating the active site pocket. **F**. Protein tunnel. Highlighted in pink is a tunnel that is formed between the α-subunits of MjFH that passes through the entire breadth of the protein.

### Difference between class-I apo- and holo-FH

To understand structural similarities and differences between MjFH and LmFH, the structures of the two proteins were superposed. Although the MjFHα- and β-subunits individually superpose well with the N-terminal and C-terminal domains of LmFH with an RMSD of 1.8 Å and 1.4 Å, respectively (**Figure 3A, B**), structural superposition of the biological assemblies of the two proteins showed gross differences with an RMSD of 4.95 Å (**Figure 3C**). A visual inspection revealed that the LmFH structure is more compact compared to MjFH and measuring the distance between conserved residues located at the extreme termini of both proteins showed that the α-subunits and N-terminal domains of MjFH and LmFH are equidistant, while the β-subunits in MjFH are spaced farther apart compared to C-terminal domains of LmFH (**Figure 3D**). In LmFH, the CTD has been shown to be mobile and its mobility changes with substrate/inhibitor binding. The B factor values for the swiveling β-subunit, as expected, is higher than that of the α-subunit of the protein (**Figure 3E**). A major proportion of residues at the NTD/NTD or α/α dimer interface of class-I FH are found to be conserved,^25^ and an interface analysis reveals that LmFH has a larger number of interactions between the domains of the protein, possibly accounting for its compact structure. While the NTD/NTD interface in LmFH is stabilized by 42 hydrogen bonds with an overall interface interaction area of 3712 Å^2^, the α/α subunit interface in MjFH is stabilized by 34 hydrogen bonds with an interface interaction area of 2521 Å^2^. Similarly, 19 hydrogen bonds stabilize NTD-CTD domains of LmFH with an interface area of 1503 Å^2^, while only 5 hydrogen bonds are observed between the α- and β-subunits of MjFH with an interface area of 1275 Å^2^. 5 residues are involved in van der Waals interactions between the CTDs of LmFH but the β subunits of MjFH do not form any contacts (**Table S3**).

**Figure 3.**
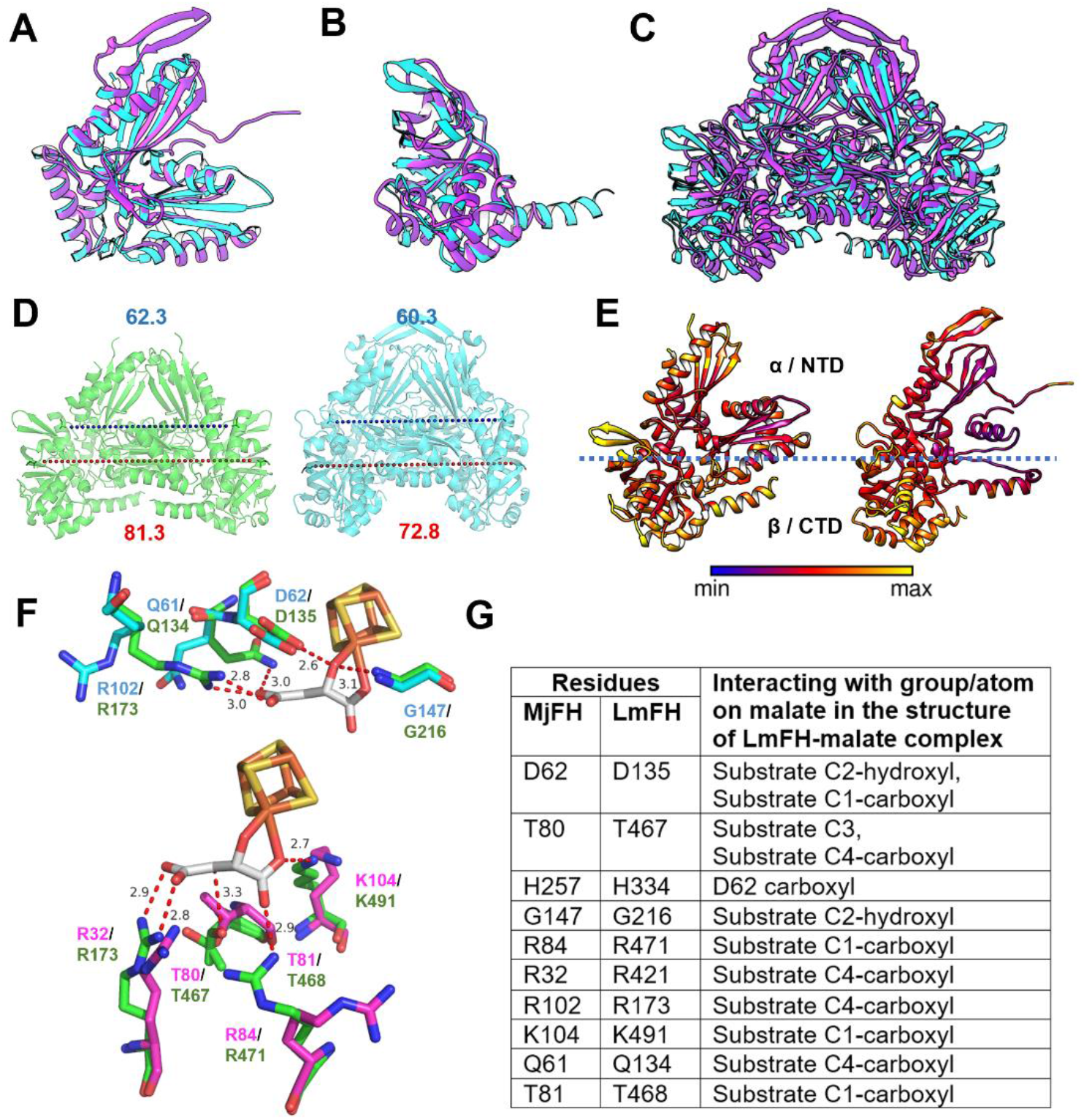
Differences between apo-MjFH and holo-LmFH (5l2r) structures. Superposition of LmFH NTD on MjFHα (**A)** and LmFH CTD on MjFHβ (**B**) show good alignment with RMSD values of 1.8 Å and 1.4 Å, respectively. **C**. Overlay of LmFH functional complex on MjFH shows gross structural differences and a lack of clear alignment, with an RMSD of 4.95 Å. **D**. Distance between **c**onserved residues A52 and A125 located at the extreme ends of MjFHα (green) and LmFH NTD (blue) dimers show similar spacing, while distance between conserved residues V160 and V546 of MjFHβ and LmFH CTD show that β-subunits of MjFH are more distantly spaced compared to CTD of LmFH. **E**. B-factor analysis reveals that MjFHβ and LmFH CTD have higher B-factor values compared to the overall average of the protein. **F**. Active site residues in MjFH compared to LmFH. Upper panel shows the superposition of active site residues from MjFHα (blue) on LmFH NTD (green) and lower panel shows the superposition of active site residues from MjFHβ (magenta) on LmFH CTD (green). Malate bound to the cluster is highlighted in grey. Residues Gln61 and Arg102 from the α-subunit and Arg84 from β-subunit of MjFH show different side chain rotameric conformation. **G**. Numbering of the active site residues of MjFH and LmFH. Note that the contacts with the atoms on the substrate are from the LmFH structure.

Transposing the Fe-S cluster from LmFH onto MjFH reveals that three cysteine residues, Cys60, Cys182 and Cys269 in MjFH are well positioned to bind to the cluster (**Figure S2**). LmFH active site has 9 residues present in loops, hydrogen-bonding with the substrate at the active site (Residue numberings of all the 9 residues for both MjFH and LmFH are provided in **Figure 3G**). 6 of these residues are positioned similarly in MjFH apo-protein active site, while three other residues (Gln61 and Arg102 from the α-subunit and Arg84 from β-subunit) adopt a different side chain rotameric conformation going to show that the active site is *largely pre-organized* even in the absence of cluster and substrate (**Figure 3G**).

### Binding of cluster and substrate compacts quaternary structure

The inter-CTD distance (the distance between the centers of mass (COM) of the CTDs (LmFH)/β-subunits (MjFH)) and radius of gyration (Rg) in the crystal structure of apo-MjFH are greater than those of holo-LmFH (**Table S7**) as can also be visually inferred from the overlay of the two structures (**Figure 3C**). To identify the molecular determinants for this difference, atomistic molecular dynamics simulations were carried out on both apo- and holo-forms of both MjFH and LmFH (see supporting information). Force field parameters employed herein were validated by examining the stability of holo-LmFH and the cluster and substrate during the course of simulation (**Figure S6 A)**. A visual inspection into the structure of the 4Fe-4S cluster and the bound malate at 200 ns simulation time frame of holo-LmFH (**Figure S6 B**) also illustrates that the malate and cluster are intact and stable. Next, we created a model for holo-MjFH by docking the cluster and substrate into the experimentally determined apo-MjFH structure (see supporting information, and **Tables S4** and **S5**, and **Figure S10** for docking details). This simulation, hereinafter referred to as holo-MjFH-1, however, did not result in the contraction of the holo-MjFH enzyme (**Figure S7**). Thus, we resorted to metadynamics based MD simulations to arrive at a model structure for holo-MjFH, and these simulations, hereafter referred to as holo-MjFH-2, (see supporting information for details) indeed resulted in a stable structure, contracted with respect to the apo-MjFH enzyme. Both the inter-CTD distance and Rg values of the holo-MjFH-2 stay stable during the entire course of simulation (**Figure 4A and 4B**). An increase in the inter-CTD distance and Rg values in the simulation of apo-LmFH as well, strongly suggests that the class-I FH structure contracts or expands based on the presence or absence of the bound Fe-S cluster and substrate. Both these values for apo-LmFH increase initially and remain saturated over the last 300 ns of simulation (**Figure 4A and 4B**). Structures of all the enzyme forms obtained as that representing the major cluster (obtained using GROMACS *cluster* module) over the last 100 ns of the corresponding trajectories are overlaid and presented in **Figure 4C and 4D**. The cartoon representations show that the CTDs of both the apo- (expanded) and holo- (contracted) forms of MjFH superpose better with the corresponding forms of LmFH than the alignment of apo-FH (expanded) on holo-FH (contracted) (**Figure 3C** and **Figure S11 A & B**). The superposition of apo-LmFH on holo-LmFH (**Figure S11. C**) and apo-MjFH on holo-MjFH (**Figure S11. D**) clearly demonstrate that the movement of CTD/β-subunit is responsible for the expansion of apo-form of class-I FH.

**Figure 4.**
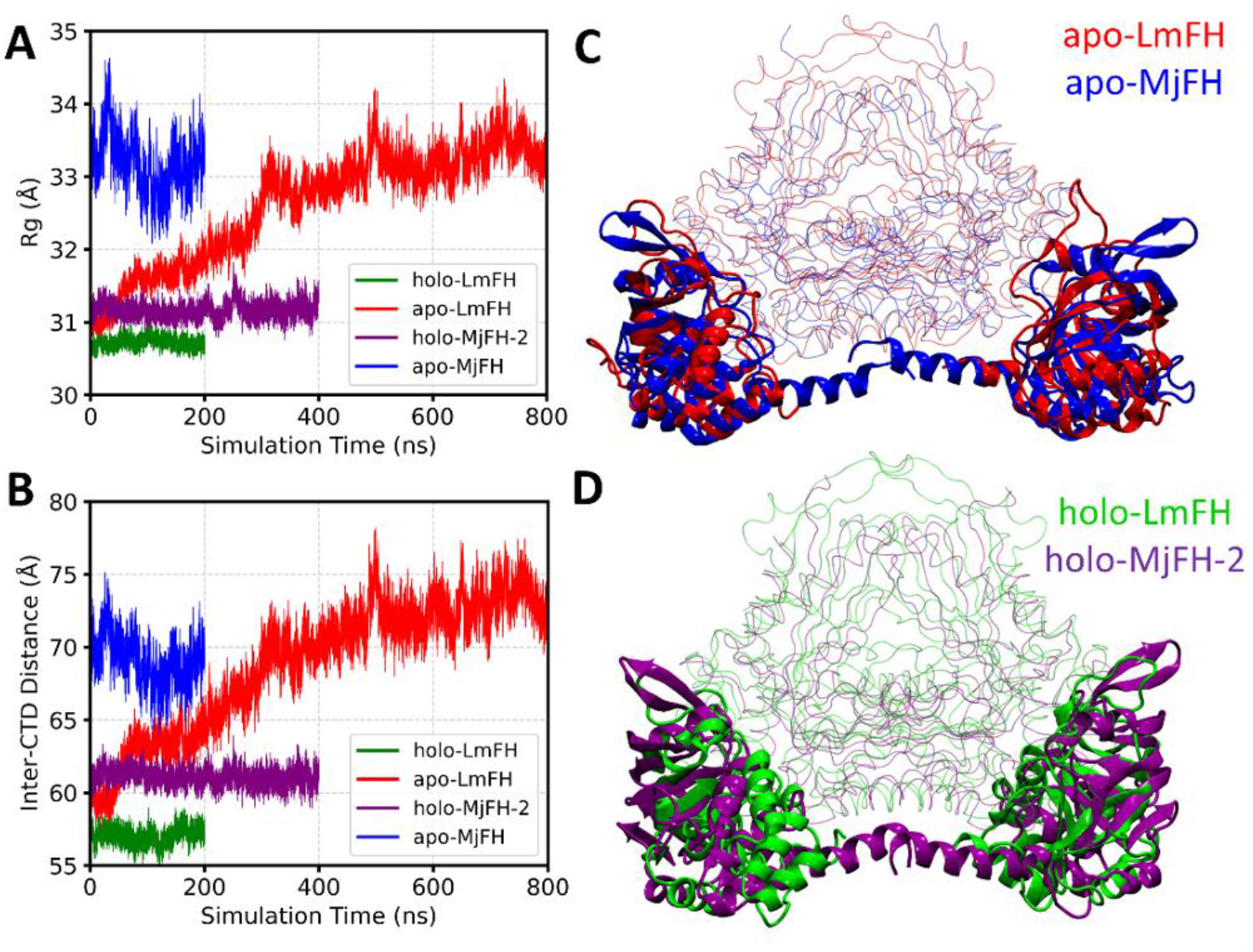
MD simulations reveal class-I FH adopts two different states based on either the presence or absence of a bound 4Fe-4S cluster and substrate. **A**. Rg and **B**. inter-CTD distance as a function of simulation time for holo-LmFH (green), apo-LmFH (red), holo-MjFH-2 (purple), and apo-MjFH (blue). Refer to **Figure S7** for the same of holo-MjFH-1 model. The apo-LmFH expands relative to the holo-LmFH, and both the Rg and inter-CTD distance values increase initially and remain saturated for the last 300 ns of simulation. **C**. Overlay of the representative structure of apo-LmFH with that of apo-MjFH and **D**. Overlay of the representative structure of holo-MjFH-2 with that of holo-LmFH. The representative structures are obtained using cluster analysis on the last 100 ns of the simulations. The CTDs (LmFH)/β-subunits (MjFH) are shown in cartoon representation, whereas the remaining parts of the enzymes are shown in transparent lines. The Rg for LmFH is calculated excluding the highly flexible 63-residues long modelled fragment, and this peptide fragment is not shown in the overlay panels (panels C and D).

### Kinetics and site-directed mutagenesis

The activity of the purified recombinant enzyme was measured at 240 nm as a time-dependent increase or decrease in absorbance of fumarate in assay mixture. The MjFH apo-protein, β-subunit and reconstituted α-subunit were found to be inactive while addition of MjFHβ to Fe-S reconstituted MjFHα yielded high levels of activity. Kinetic parameters of MjFH for activity on malate and fumarate were obtained by fitting *v vs* [S] plots to substrate inhibition equation (eq. 1, supporting information) (**Figure 5A**). *K*_M_, *K*_I_ and *V*_max_ values derived from the fits are summarized in **Table 1**. The catalytic efficiency (*k*_cat_/*K*_M_) of MjFH for both malate and fumarate is comparable to other reported class-I FH. *RS*-2-thiomalate was found to inhibit MjFH with an IC_50_ of 50.8 ± 1.2 μM and 20.6 ± 0.8 μM with malate and fumarate as substrates, respectively (**Figure 5B**). Values of the apparent inhibitory constant *K_i_* (app) for *RS*-2-thiomalate derived from the IC_50_ values with malate and fumarate as substrates were 30.8 μM and 17.4 μM, respectively, and similar to those previously reported for Class-I FH.^10,23,68^

**Figure 5.**
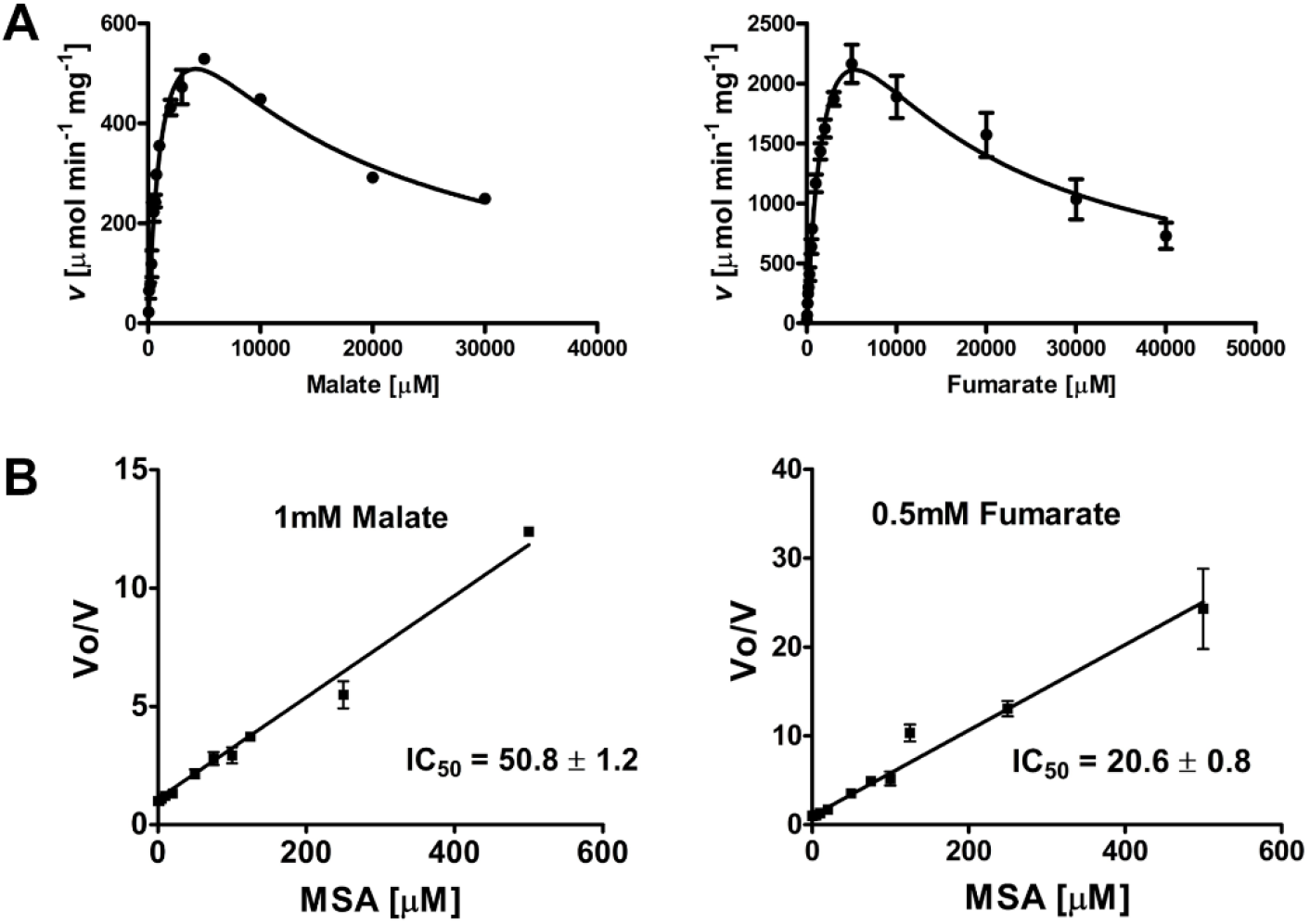
MjFH kinetics and IC_50_ plot for inhibition by *RS*-2-thiomalate. **A**. Initial velocity as a function of substrate concentration for malate (left panel) and fumarate (right panel). Data were fit to substrate inhibition equation (eq. 1, supporting information). **B**. IC_50_ plot for RS-2-thiomalate with malate (left panel) and fumarate (right panel) as substrates. The unit for IC_50_ values is μM.

**Table 1.**
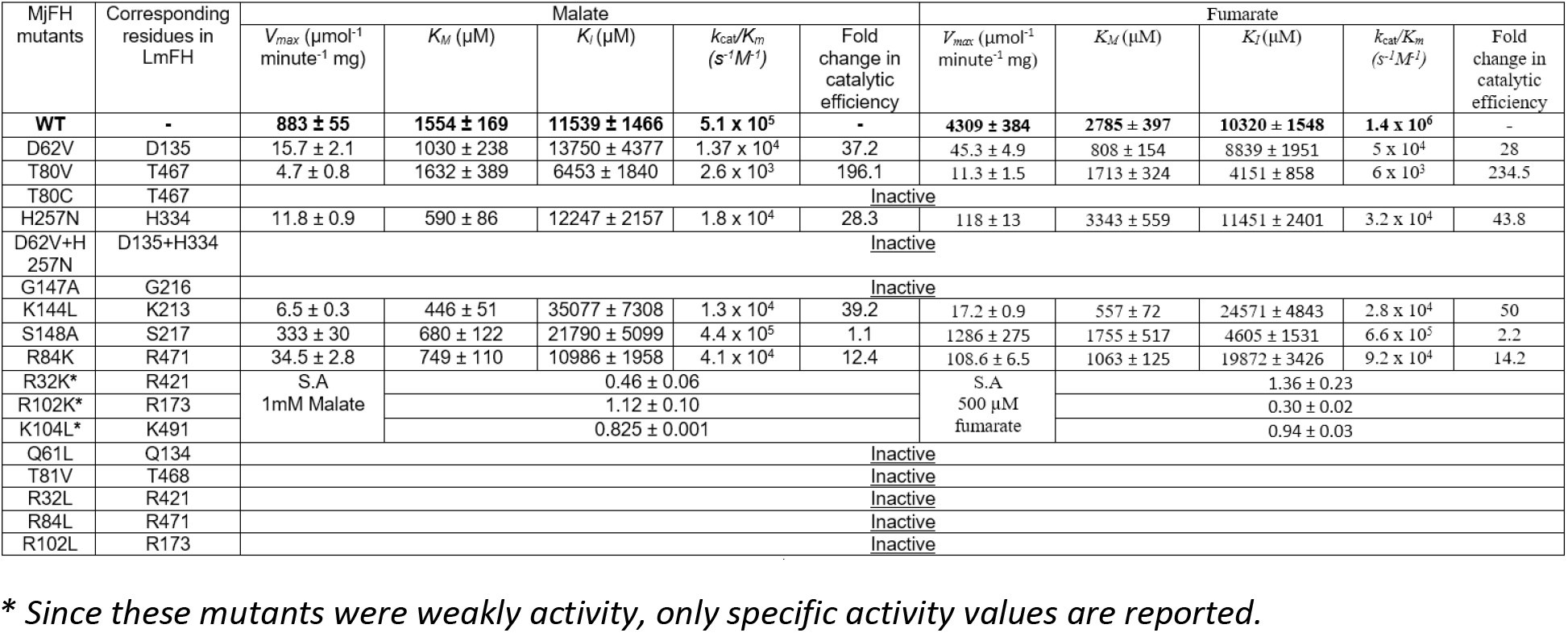
Kinetic parameters of MjFH WT and mutant enzymes

To understand the catalytic mechanism of class-I FH, residues were selected for site-directed mutagenesis in MjFH, which according to the *S*-malate bound *Leishmania major* FH (5l2r) structure and more recently solved inhibitor – thiomalate bound structure (6mso, 6msn), can interact with the substrate. A set of 10 residues are located at the active site of which 9 residues are in hydrogen bonding distance with *S*-malate (Gln61, Asp62, Arg102, Gly147 from α subunit and Arg32, Thr80, Thr81, Arg84, Lys104 from β-subunit) (**Figure 3G**). The residue/s closest to C3 carbon of *S*-malate is Thr80 at 3.34 Å and to C2 hydroxyl group are Asp62 and Gly147 at 2.64 Å and 3.1 Å distance, respectively. Another residue His257 from the α-subunit, although is not interacting with the substrate, is hydrogen-bonded to Asp62 at a distance of 2.81 Å and could stabilize the charge state of this residue. All other residues form H-bonds with the carboxyl groups of *S*-malate. Sequence alignment also shows that Gly147 is part of a motif ‘KGXGS’ that is highly conserved in class-I FH (**Figure S3**) and considering their proximity to the substrate, Lys144 and Ser148 from the motif were also chosen for mutation.

To examine the function of the 10 active site residues and 2 conserved residues from the KGXGS motif, all these 12 residues were replaced by site-directed mutagenesis. Mutant enzymes were generated and purified using protocols similar to WT protein. Steady state kinetics were conducted under conditions similar to that used for WT and the kinetic parameters along with fold change in catalytic efficiency relative to WT are summarized in **Table 1**. Mutants with measurable activities showed a small drop in *K*_M_ value, between 1.5 to 3 folds for both substrates, which is close to that observed in LmFH mutants. The K144L mutant where K144 is part of the KGXGS motif, had the highest drop in *K*_M_ values by 3.5 and 5 folds, respectively for malate and fumarate suggesting a possible role in substrate binding. Unlike in LmFH where mutation of residues corresponding to threonine80 and aspartate62, the proposed catalytic base and acid, respectively led to a large drop (2500-5500-fold) in catalytic efficiency, MjFH_T80V and MjFH_D62V retained significant activity. The mutant T80V of MjFH showed 196- and 235-fold drop in catalytic efficiency for malate and fumarate, respectively while MjFH_D62V showed a 37- and 28-fold reduction for the two substrates. Mutation of threonine to cysteine however, led to MjFH_T80C being inactive. A similar trend has been observed in adenylosuccinate lyase, where mutating the serine residue believed to be the catalytic base to alanine led to 1000-fold drop in specific activity while mutating it to cysteine abolished activity. It is suspected that the highly reactive thiol group of cysteine disrupts the active site by geometric and electrostatic perturbations.^69^ Mutation of the histidine residue in contact with the predicted catalytic acid to asparagine in MjFH_H257N also led to a 28- and 44-fold reduction in catalytic efficiency for both the substrates. A double mutant of D62V and H257N of MjFH was inactive suggesting the requirement of at least one of these residues at the active site. While mutant K144L from the KGXGS motif showed a drop in catalytic efficiency of 39- and 50-fold for malate and fumarate as substrates, S148A showed no drop in catalytic efficiency for malate and a 2-fold drop for fumarate as substrate. Mutation of Gly147 of the KGXGS motif to alanine led to an inactive enzyme. The backbone amide NH of G147 is at 3.1 Å from the C2 hydroxyl group of malate in the malate-bound structure of LmFH and mutating it to alanine could lead to the side chain occupying the active site space, resulting in short contacts that occlude substrate binding.

Mutation of all other residues (Gln61, Arg102, Gly147, Arg32, Thr81, and Arg84) at the active site that are in close contact with the carboxyl groups of the substrate to a residue with non-polar side chain led to inactive mutants except for the mutant MjFH_K104L that exhibited very feeble activity precluding estimation of *K*_M_ and *V*_max_ values. It must be noted that the corresponding mutants (R173A, R421A and R471A) in LmFH exhibited weak activity.^70^ Our inability to detect this weak activity could be due to the differences in oxygen levels across the two experimental setups: 0.1 ppm versus 6 ppm.

A cluster of positively charged residues are arrayed around the dicarboxylate substrate, thereby polarizing the reactive carbon-carbon double bond. Two arginine residues and one glutamine (R32, R102 and Q61) clamp the substrate from one side (Side A, **Figure 6**) and one lysine, arginine, and threonine residue (K104, R84 and T81V) clamp from the other side (Side B, **Figure 6**). The positively charged residues are asymmetrically distributed about the symmetric substrate facilitating charge polarization. The hydrogen bonding network on side A is mediated by 4 strong bonds formed by two arginine residues (R32 and R102), in contrast to side B which has only two hydrogen bonds from one lysine and one arginine residue (K104 and R84).

**Figure 6.**
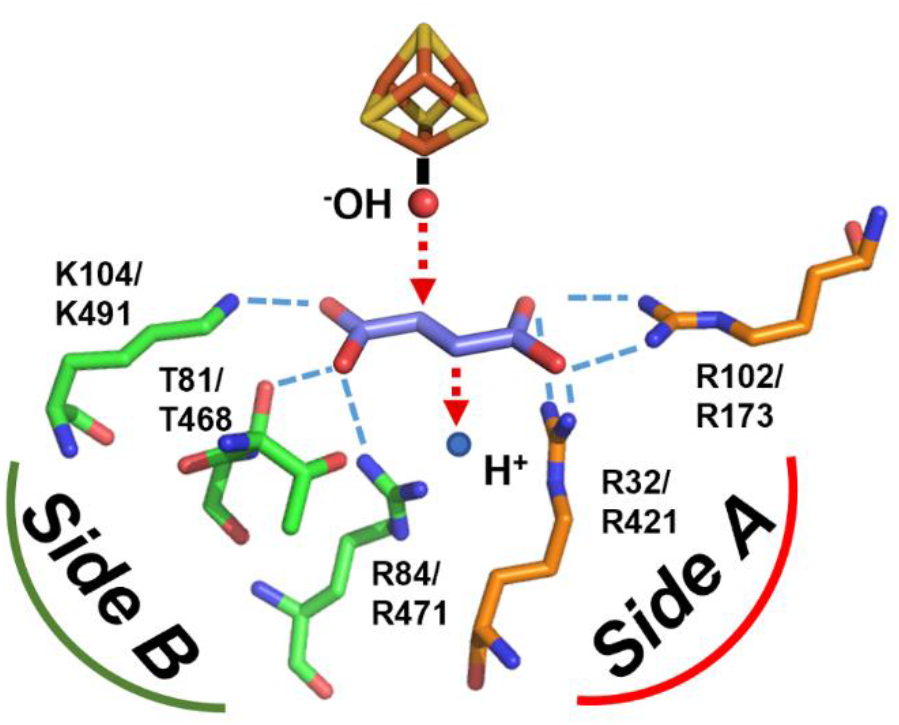
Substrate is charge-polarized at the active site with more charge interactions on one side compared to the other. Residues Arg102 and Arg32 on side A form 4 strong hydrogen bonds with carboxyl group of bound substrate, compared to one bond each formed by Lys104 and Arg84 on side B.

As the charged side chains of these residues could essentially have a role in catalysis by polarizing the substrate and priming it for reaction, the three arginine residues were mutated to lysine and the mutants probed for activity. Both the arginine residues on side A which form stronger hydrogen bonds with substrate on mutating to lysine (R32K and R102K) retained feeble activity precluding estimation of kinetic parameters, while mutating the arginine on side B to lysine (R84K) led to a 10-fold drop in catalytic efficiency for both substrates (**Table 1**). Mutating the charged residues on side A was more detrimental compared to side B. All inactive mutants showed a characteristic CD spectrum in the near-UV/visible region confirming that Fe-S cluster assembly is intact in these mutants **(Figure S4)**.

### Fumarate hydration in aqueous solution proceeds through the carbanion intermediate

Although in aqueous medium, the thermodynamics for malate (**A**) to fumarate (**C**) conversion has been worked out,^4^ the reaction mechanism has not been fully explored. A general mechanism for interconversion between fumarate and malate involving a carbanion intermediate (**Scheme 2**) has been proposed and this merits a fresh examination to enable comparison with that of the enzyme catalyzed reaction. Hence the first calculations were to understand/elucidate the mechanism in aqueous medium. Three different mechanisms; pathway w1 involving the carbanion, pathway w2 involving a carbocation, and pathway w3 which involves a concerted TS (transition state) have been studied. In pathway w1, the first step of malate to fumarate conversion involves the proton abstraction by a water molecule via **TS(A-B)_w1_** as shown in **Figure 7**. The second step involves the removal of a water molecule via **TS(B-C)_w1_**. In pathway w2, the first step involves a water removal as shown in **TS(A-B)_w2_** whereas, the second step involves proton abstraction via **TS(B-C)_w2_ (Scheme S1, Figure S12)** as opposed to pathway w1. For pathway w3, the conversion happens through a single TS of water removal (refer to supporting information for more detail). The optimized geometries for TSs involved in the carbocation pathway (w2) and concerted pathway (w3) can be seen in **Figure S12**. As seen from the relative free energies provided in parentheses in **Scheme 2**, the carbanion pathway, *i.e*., pathway w1 is found to be of the lowest energy. It presents an activation free energy barrier of 41.7 kcal mol^−1^ which is in accordance with the experimental value of 36 kcal mol^−1^. The optimized geometries for TSs involved in the carbanion pathway are shown in **Figure 7**. In the first **TS(A-B)_w1_**, the water molecule abstracts the proton from the C3 carbon and relays it to the carboxylate group via a six-membered TS. In the second **TS(B-C)_w1_**, the C-O bond cleaves where a water molecule donates a proton to the departing OH^−^ and simultaneously abstracts the proton from the carboxylate ion thereby generating the fumarate.

**Scheme 2.**
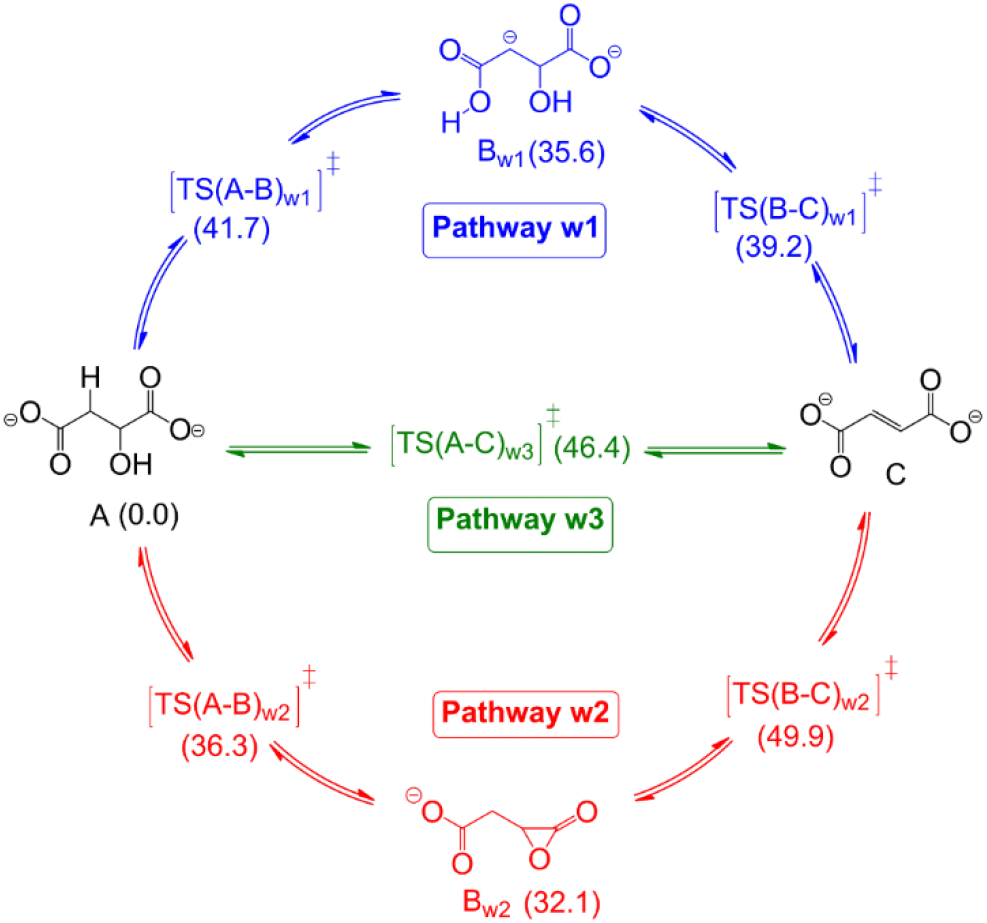
Three pathways of interconversion between malate and fumarate in aqueous medium. Values in parentheses are the relative electronic energies in kcal mol^−1^. (The energy shown for TS(A-C)_w3_ involves an external water molecule, concerted TS with no external water molecule was found to be even higher in energy (details are provided in supporting information, **Figure S13**).

**Figure 7.**
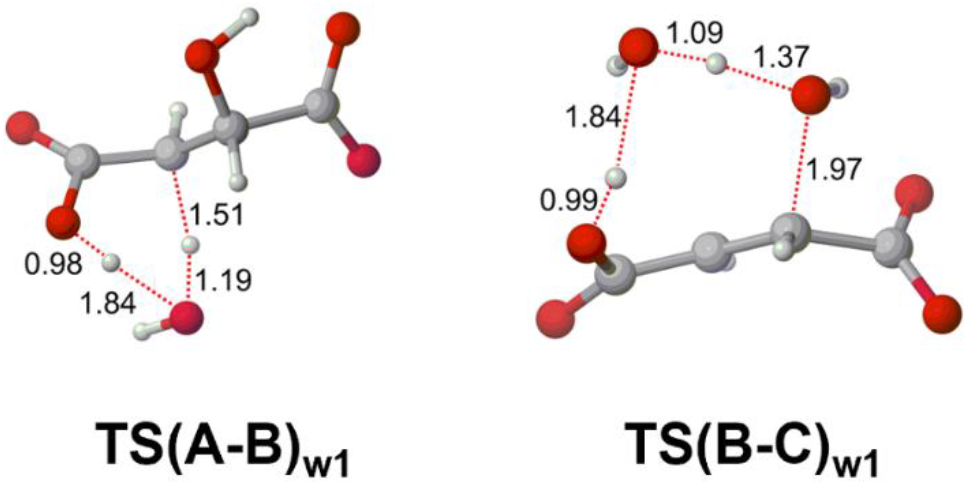
Optimized geometry of the two transition states of nucleophilic pathway (w1) in aqueous medium. Distances are in Å.

### Reaction mechanism at the enzyme active site

To understand the transition state stabilization brought out by the enzyme which corresponds to enormous catalytic rate enhancement (~ 3.5 × 10^15^ fold at pH 6.8 and 37 °C) compared to the uncatalyzed reaction, QM cluster method was utilized to probe into the reaction mechanism. Two different QM cluster models (**Figure 8**) were built for this study from the available X-ray crystal structure of FH from *Leishmania Major* (PDB ID: 5l2r). In the smaller QM cluster (QM1) consisting of 86 atoms, only five essential amino acid residues (Asp135, His334, Thr467, Thr468 & Arg471) which are believed to play an active role in the catalytic cycle were included. On the other hand, in the larger QM cluster (QM2), consisting of 207 atoms, 13 amino acid residues along with the 4Fe-4S cluster were considered. Out of these 13 residues, three cysteine residues (Cys133, Cys252, Cys346) are bound to three Fe atoms in 4Fe-4S cluster. Leaving the catalytic residues aside, five additional amino-acid residues which interact non-covalently with the bound substrate via hydrogen bonding (Gln134, Arg173, Gly216, Arg421 and Lys491) were included in QM2. Apart from the residues, a crystal water molecule was also included in QM2. Akin to the aqueous medium, here also we investigated three possible mechanistic pathways of interconversion using the QM1 model (refer to supporting information, **Scheme S2 and S3**, and **Figure S14**). The conventional pathway of olefinic hydration did not seem to be operational even inside the enzymatic active site with a high activation barrier of 42.8 kcal mol^−1^ (**Figure S15**). Although the carbanion pathway turned out to be the most favorable in this case as well, the energy barrier obtained was 21.2 kcal mol^−1^ with QM1 model which is higher than the experimentally known value of 13.8 kcal mol^−1^. However, the energy barrier reduced significantly to 16.7 kcal mol^−1^ in the larger model QM2 signifying the importance of non-covalent interactions present at the active site. In the optimized geometry of QM2 model as shown in **Figure 8**, the catalytic base T467 and catalytic acid D135 are suitably placed maintaining the H-bonding contacts as seen in the crystal structure. In **TS(A-B)_wt_**, T467 abstracts a proton from the substrate and simultaneously donates its proton to the neighboring R471 resulting in the formation of the carbanion (**Figure 9**). In the second **TS(B-C)_wt_**, OH-dissociates from the carbanion and at the same time accepts a proton from D135, thereby generating fumarate with the release of a water molecule.

**Figure 8.**
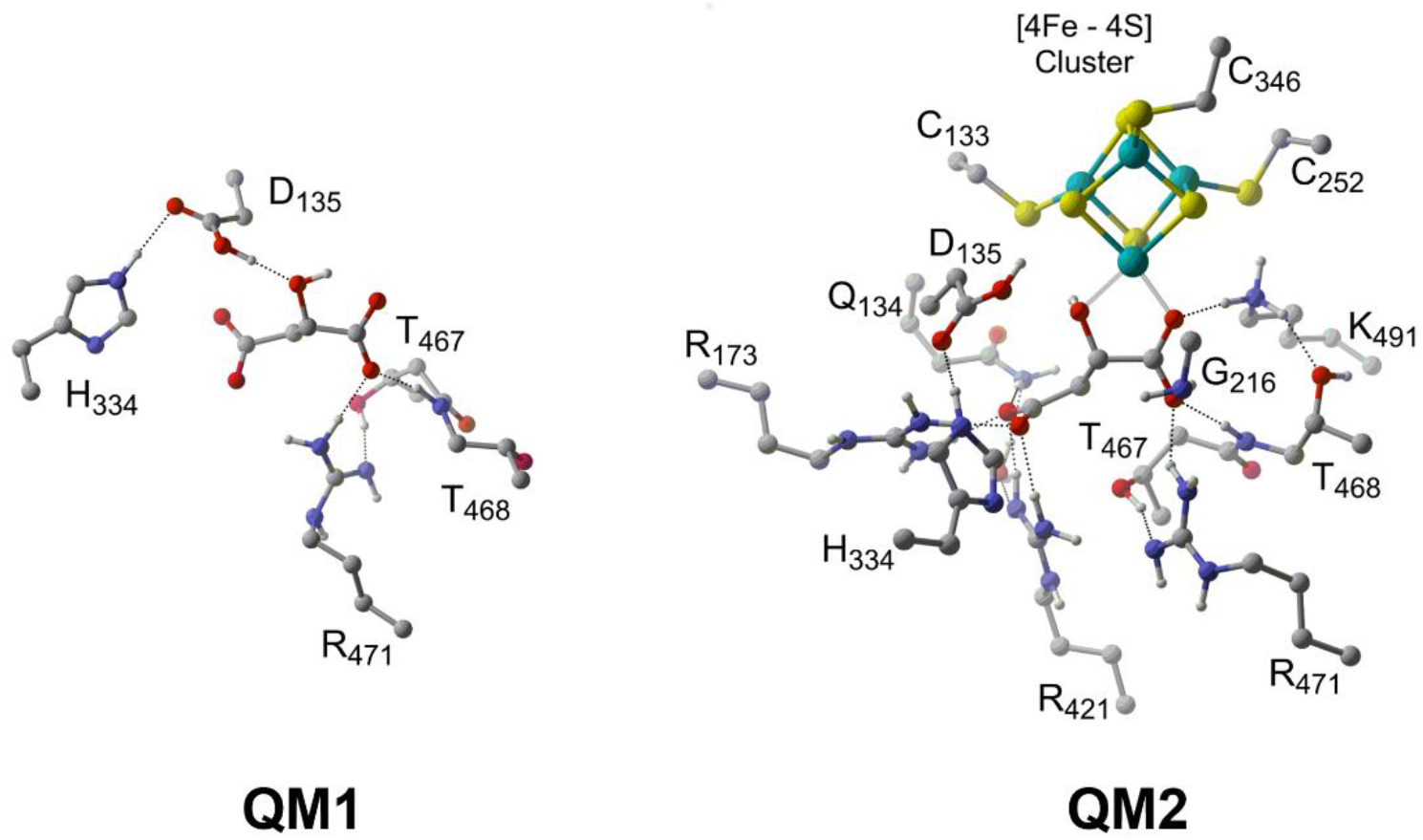
Optimized geometries of malate bound smaller (QM1) and larger QM cluster models (QM2).

**Figure 9.**
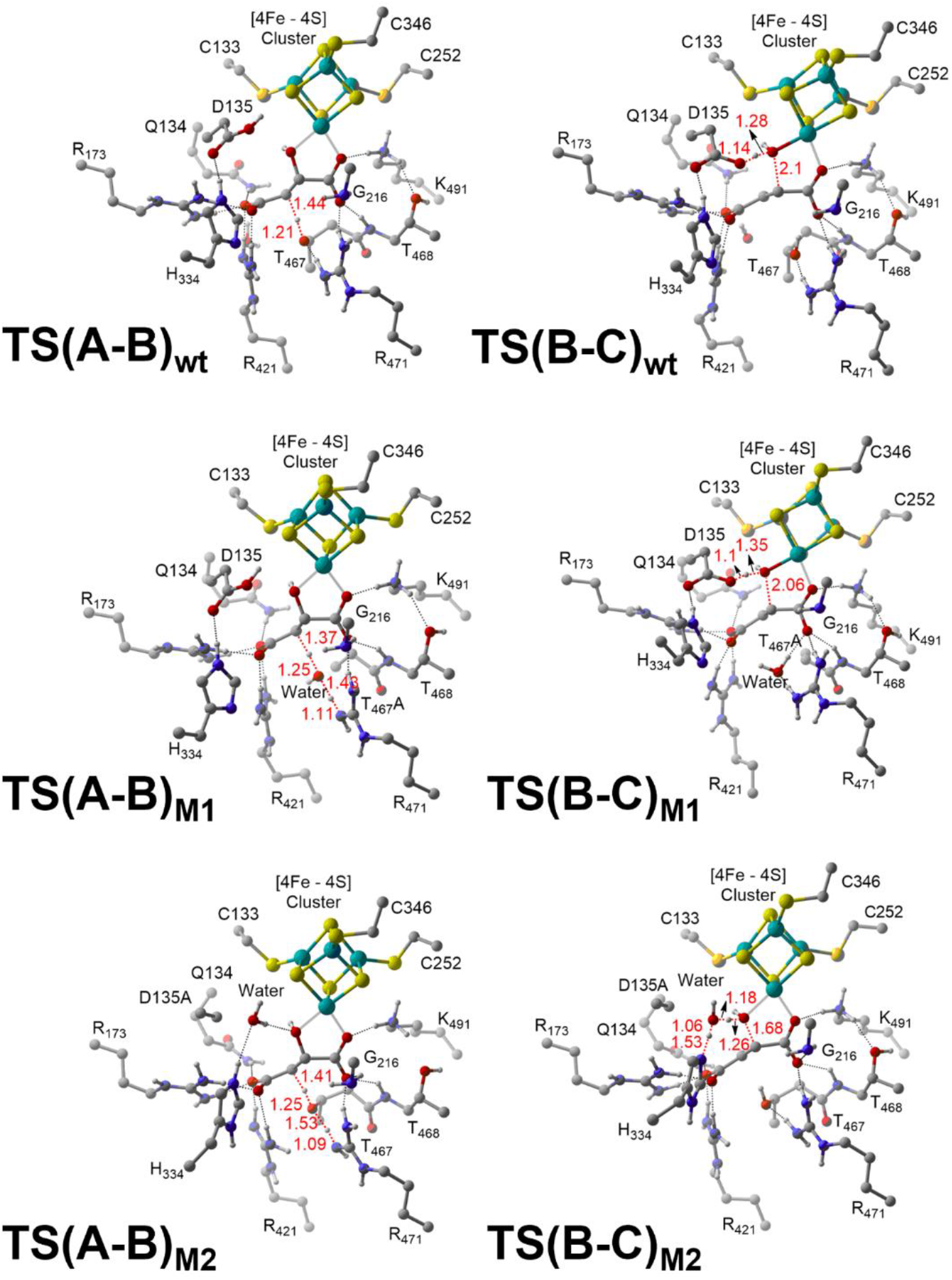
Optimized geometries of the two intermittent transition states obtained from QM2 in wild type enzyme (wt), T467A mutant (M1) and D135A mutant (M2). Distances shown are in Å.

### Alternative mechanism in the absence of catalytic residues

To evaluate the effect of mutation of catalytic residues on the reaction mechanism, we used the QM2 model and generated two mutants T467A (M1) and D135A (M2). In M1, residue Thr467 is mutated to alanine, which cannot relay the proton transfer. The obvious choice for an alternate base is the crystal water which can assist the proton transfer from C3 of malate to R471 as shown in **TS(A-B)_M1_**. In the second **TS(B-C)_M1_**, the D135 residue plays a similar role as that in the wt. In the second mutant D135A (M2), where the catalytic acid Asp135 was mutated to alanine, the second **TS(B-C)** could not be optimized with crystal water as free OH^−^ is not as stable as the negatively charged aspartate residue. We, therefore considered an alternative possibility where His334 residue exists in its protonated form (H334-H^+^). It can then transfer a proton through the water molecule to the departing OH^−^ eventually eliminating a water molecule from the carbanion via **TS(B-C)_M2_** (**Scheme S4**). The activation free energy barriers for M1 and M2 were found out to be 26.4 kcal mol^−1^ and 23.2 kcal mol^−1^, respectively (**Figure 11**). While these barriers are higher than the experimentally observed values, our model qualitatively captures the increase in activation free energies in these mutants.

## DISCUSSION

In this study, through mutational analysis and DFT calculations, we propose a mechanism for the rate enhancement elicited by class-I FH. We also see features distinct to MjFH.

### MjFH structure and catalysis

The two subunits of MjFH remain tightly associated even in the absence of ligand or Fe-S cluster. α-subunit is inactive and requires the association of the β-subunit for activity. Structure of MjFH solved is the first structure of a two-subunit class-I FH. At the time of solving the MjFH apo-protein structure, LmFH structure was not available and SAD phasing was carried out to solve the structure of the αβ-complex. The two FH structures are indicative of two different states of the protein, a slightly relaxed MjFH apo-protein state and a more compact LmFH holo-protein state. The primary difference is in the β-subunit and the CTD of the proteins which are spaced differently. MD simulations carried out with the apo- and holo-structures of LmFH and MjFH show that the enzyme expands upon removal of the Fe-S cluster and substrate, and confirms that the difference in domain assembly across the two structures is a consequence of Fe-S cluster and substrate binding. Fe-S cluster bound but substrate and inhibitor free structure of LmFH showed increased mobility of the C-terminus domain as compared to the substrate/inhibitor bound structure.^70^ Since the pocket for entry of substrate to the active site lies between the α- and β-subunit interfaces, it is possible that the β-subunit mobility may regulate substrate entry/product exit into the active site by opening/closing during the catalytic cycles.

The kinetic parameters obtained for the enzyme are in concordance with previous reports on class-I FH. However, we observed substrate inhibition in this enzyme which suggests an alternative binding site for the substrate. This is the first enzyme of this class to exhibit such a regulation with the only other known regulation reported in class-I FH being cooperativity observed in *Trypanosoma cruzi*.^23^ Interestingly, malonate, a known inhibitor of class-II FH was bound to two different regions of the tunnel in LmFH, one at the entrance and other at the center.^25^ Two of the residues in the tunnel which bind to malonate in LmFH, Gln195 and Glu267 (LmFH numbering) are conserved, while Asp197 is present either as a glutamate or histidine in other class-I FH. Residues Lys213 and Ser217 (part of KGXGS motif) are close to the tunnel with Lys213 at around 10 Å from Gln195 and Glu267. An exhaustive search for side chain rotamers of Lys213 followed by energy minimization in LmFH structure reveals that the lysine residue comes in close contact of 4.5 Å from Glu267 **(Figure 10)**. Mutating the lysine and serine residues results in significant changes in *K_i_* (**Table 1**) suggesting the alternative binding site in MjFH too might be located within the tunnel.

**Figure 10.**
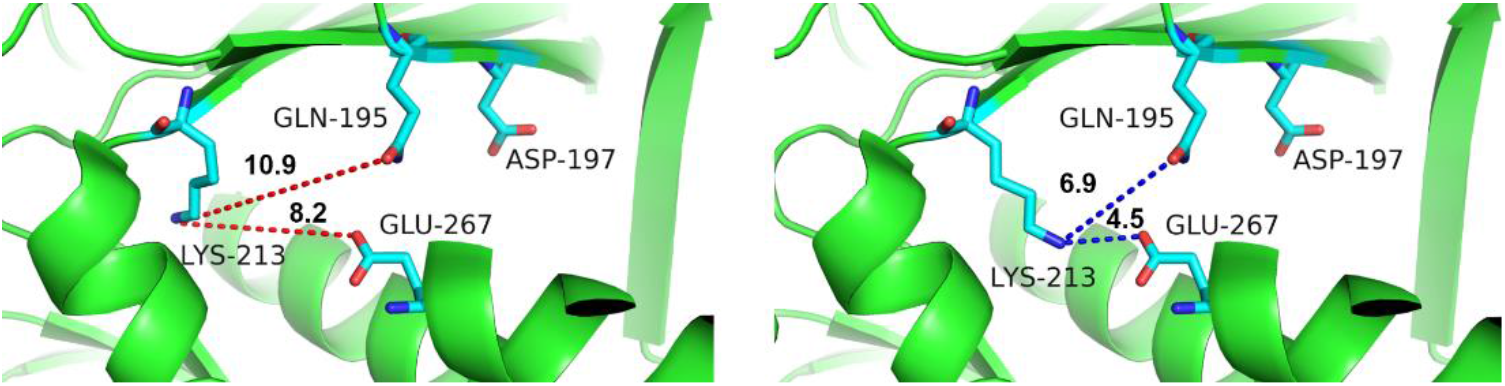
Contacts of Lys213 of the conserved KGXGS motif with Gln195 and Glu267 that interact with malonate bound at the tunnel in LmFH (5l2r). Panel on left shows interatomic distance between Lys^213^ residue, and Gln^195^ and Glu^267^ present in the tunnel. Search for possible side chain rotamers of Lys^213^ in the same structure followed by energy minimization reveals that Lys^213^ comes close to Gln^195^ and Glu^267^ as seen in the right panel. The tunnel in LmFH structure is independent of the active site. The numbers above the dashed lines are distances in Å.

**Figure 11.**
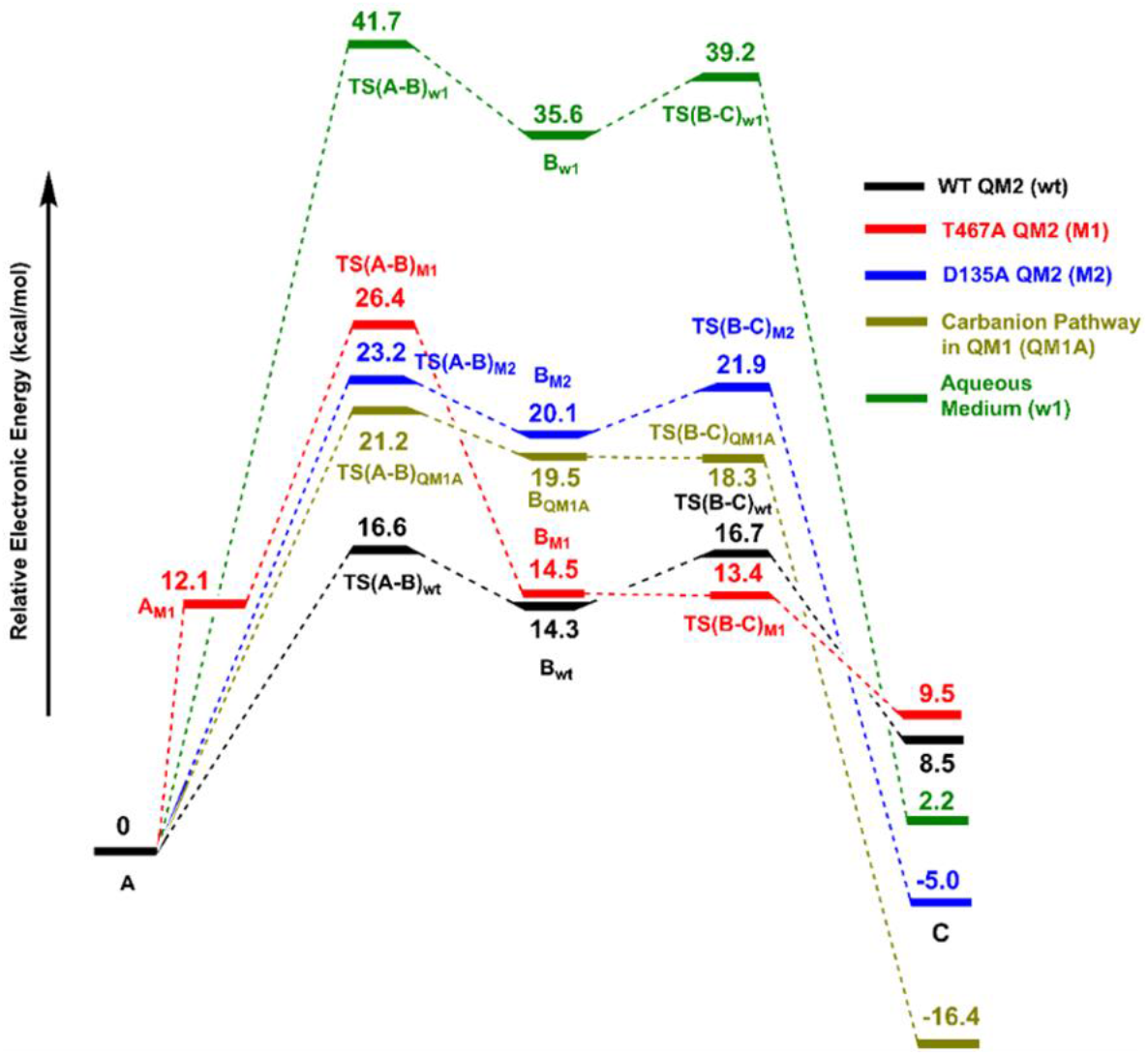
Electronic energy profiles obtained from different model systems.

Recently, the first attempt at understanding the mechanism of catalysis in class-I FH has been carried out in LmFH through mutational studies of active site residues. The residue, Thr467 (LmFH numbering) closest to the C3 carbon of bound substrate along with its hydrogen-bonded partners consisting of two arginyl residues, Arg421 and Arg471, has been predicted to act as catalytic base while Asp135 along with its hydrogen-bonded partner His334 performs the role of catalytic acid.^70^ Mutation of the predicted catalytic acid and base in LmFH led to a 2500-5500 fold drop in activity (k_*cat*_/K_M_), while in MjFH, the corresponding mutations T80V and D62V leads to a lower fall in activity (30-240 folds). It must be noted that the catalytic rate enhancement offered by the enzyme is of the order of 3*10^15^ and a 5000 fold drop in activity of the enzyme would still make it a phenomenal catalyst. This suggests that the rate determining step in the fumarase reaction in the case of class-I FH is not proton abstraction. We must therefore look elsewhere for the molecular origin on the large transition state stabilization first noted by Bearne and Wolfenden.^4^ Mutation of all the residues (Gln61, Arg102, Gly147, Arg32, Thr81, and Arg84) that form a hydrogen-bonded network with the carboxylate groups of the substrate, abolished activity in MjFH. These observations on the preferential loss of activity upon mutation of the substrate-binding residues suggest the role of charge polarization in mediating catalysis in class-I FH.

### QM calculation supports the dominant role of substrate charge polarization in catalysis

With the present studies on the catalytic mechanism in class-I FH still in its nascent state, our understanding of this aspect is largely derived from studies carried out in class-II FH. Though the biochemical equilibrium of the reaction catalyzed by FH is in the direction of fumarate to malate (K_eq_=4), the problem has been viewed in the light of dehydration rather than hydration. A serine residue has been identified as the catalytic base in class-II FH, where it is believed to form an oxyanion hole, stabilized by interaction with positively charged neighbors. Previous studies however, have revealed that the rate limiting step does not involve a carbon-hydrogen bond cleavage, clearly suggesting that the reaction may not be driven largely through catalytic base mediated proton abstraction.^17^ Both class-I and class-II FH catalyze the unidirectional hydration of acetylenedicarboxylate to oxaloacetate with good efficiencies.^71,72^ This reaction, similar to the reaction catalyzed by acetylene hydratase does not require a catalytic base and the reaction is catalyzed by a water molecule held in position bound to a tungsten cofactor. The bound water molecule is rendered nucleophilic or electrophilic by an aspartic acid residue thus allowing it to attack the triple bond.^73^ In recent years, quantum chemical studies have proven successful to understand mechanism of enzymatic reactions. DFT calculations on both electrophilic and nucleophilic pathways for water attack on triple bond of acetylene have shown very high energy barriers suggesting an alternative mechanism in acetylene hydratases. We have carried out quantum chemical studies on class-I FH to understand the mechanism that this enzyme adopts to catalyze the hydration/dehydration reaction of fumarate.

From our DFT calculations we find that the carbanion pathway is the most favorable route of interconversion between malate and fumarate in aqueous medium as well as in the enzymatic active site. The uncatalyzed reaction in aqueous medium has a significant energy barrier (41.7 kcal mol^−1^), which drastically goes down by 20.5 kcal mol^−1^ in the presence of the catalytic acid-base pair inside the protein environment (**Figure 11**). Incorporation of non-covalently interacting residues in the larger cluster QM2 results in further reduction of energy barrier suggesting that they might have an important role in modification of the electronic structure of the substrate.

A closer look at the active site of class-I FH reveals that the environment is polarized with more positively charged interaction at one end of the substrate compared to the other. We hypothesize that the active site environment polarizes the double bond of fumarate so that the nucleophilic attack of the hydroxyl group becomes favorable. To understand if the double bond of fumarate is really primed for attack of the water molecule, we performed Natural Population Analysis (NPA) calculation on fumarate bound enzyme-substrate complex in the larger model, QM2 for WT and various mutants (**Table 2**). From the NPA calculation it is observed that the charge separation is maximum for WT with K491A and Q134A mutants being the only exceptions. As the active site residues are mutated, the relative charge separation reduces making the fumarate less nucleophilic, which can in turn cause an increase in activation energies as already observed in the case of T467A and D135A mutants. Importantly, active site arginine residues play a significant role in the charge separation. As the three arginine residues are removed sequentially from the active site, the charge separation not only reduces but the polarity completely gets reversed. Thus, the nucleophilicity of the substrate totally diminishes as the arginine residues are mutated, which in turn demolishes the enzyme activity. From the good correlation obtained here, it can be said that non-covalent interactions present at the active site of FH also have an essential part in making the enzyme more efficient by polarizing the substrate.

**Table 2.** NPA charges on bound fumarate for different mutants of LmFH

| Mutant | NPA Charge on C3 | NPA Charge on C2 | Relative NPA Charge on C2 w.r.t. C3 |
| --- | --- | --- | --- |
| WT | - 0.306 | - 0.212 | + 0.094 |
| R471A | -0.286 | -0.222 | +0.064 |
| R421A | - 0.265 | - 0.231 | + 0.034 |
| R173A | -0.273 | -0.212 | +0.061 |
| R421A+R173A | - 0.237 | - 0.272 | - 0.035 |
| R421A+R173A+R471A | - 0.240 | - 0.294 | - 0.054 |
| K491A | -0.314 | -0.198 | +0.116 |
| D135A | -0.302 | -0.234 | +0.068 |
| D135A+H334A | -0.275 | -0.234 | +0.041 |
| Q134A | -0.318 | -0.214 | +0.104 |
| T467A | -0.264 | -0.228 | +0.036 |
| T468A | -0.282 | -0.201 | +0.081 |
| G216A | -0.284 | -0.223 | +0.061 |
| Fumarate | -0.285 | -0.285 | 0.000 |
| Fumarate + [4Fe-4S] cluster | -0.245 | -0.286 | -0.041 |

### Proposed mechanism of catalysis in class-I FH

With the understanding that the base mediated proton abstraction is not the rate limiting step of the reaction and reduction in free energy barriers is not achieved solely by the catalytic acid and base pair, the transition state analogues have been used to explain the series of covalent changes that take place during the fumarase reaction of an enzyme bound substrate.^15^ Rate enhancements in enzymes take place through reducing the activation barrier of the reaction by binding strongly to the altered substrate in its transition state compared to the substrate in its ground state.^4^ FH stabilizes this altered substrate in its transition state by a reduction in free energy by at least 30 kcal mol^−1^ with a release in enthalpy of −24 kcal mol^−1^ and a gain in entropy of 19 kcal mol^−1^.^4^ It is observed that the C3-carbanion intermediate binds FH very strongly, mimicking the transition state.^15^ The large favorable enthalpy gain in the fumarase reaction clearly arises from the strong network of electrostatic hydrogen bonds formed by the substrate at the preorganized active site. In the absence of substrate, the charged active site residues are likely to be heavily hydrated. Binding of the dianionic substrate will be accompanied by a release of the hydration shells around both the positive side chain of enzymes and negative charge of carboxylate groups of substrate, contributing to large gain in entropy.

The active site of class-II FH as well, reveals a pattern similar to class-I FH, where the number of hydrogen-bonds with the substrate is much larger on one end compared to the other **(Figure 12)**. We hypothesize that polarization of the substrate at the active site mediated by non-covalent interactions could potentially form the basis of catalysis where binding of the substrate and catalysis become inseparable events as predicted by the transition state theory. Upon substrate binding to the active site, the surrounding residues prime the substrate for nucleophilic attack and provide electrostatic stabilization to the negative charge on the carbanionic intermediate form of the substrate approaching the transition state **(Figure 13)**. Bearne and Wolfenden had long predicted that the electrostatic stabilization could explain the large gain in entropy (19 kcal/mol) observed for binding the altered substrate in its transition state relative to the ground state.^4^ We believe that the affinity of FH active site for the binding of the transition state far exceeds the affinity for the substrate, thus catalyzing the reaction.

**Figure 12.**
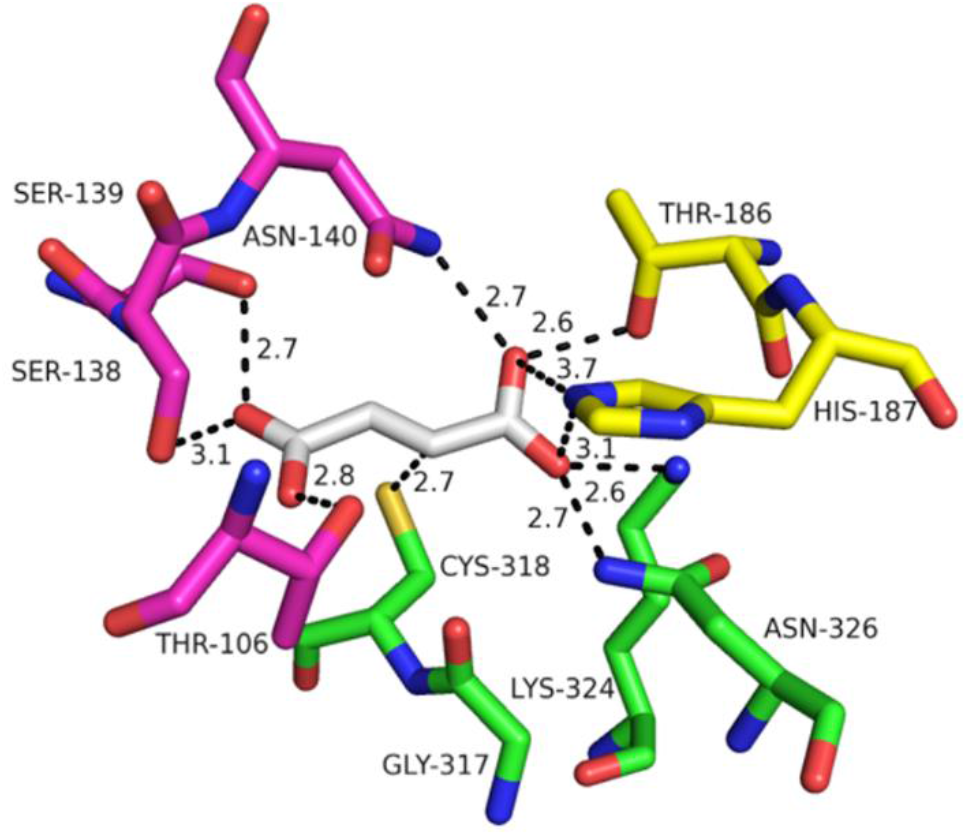
Active site architecture of class-II FH taken from *M. tuberculosis* FH reveals a larger number of H-bonds at one end of the substrate compared to other. Residues from three subunits contribute to catalysis in class-II FH, labelled as magenta, green and yellow, respectively. We see that while a serine and threonine residue are in H-bonding distance from the carboxylate group of the substrate (highlighted in grey) from one end, stronger interactions with lysine, histidine and asparagine are observed on the other end.

**Figure 13.**
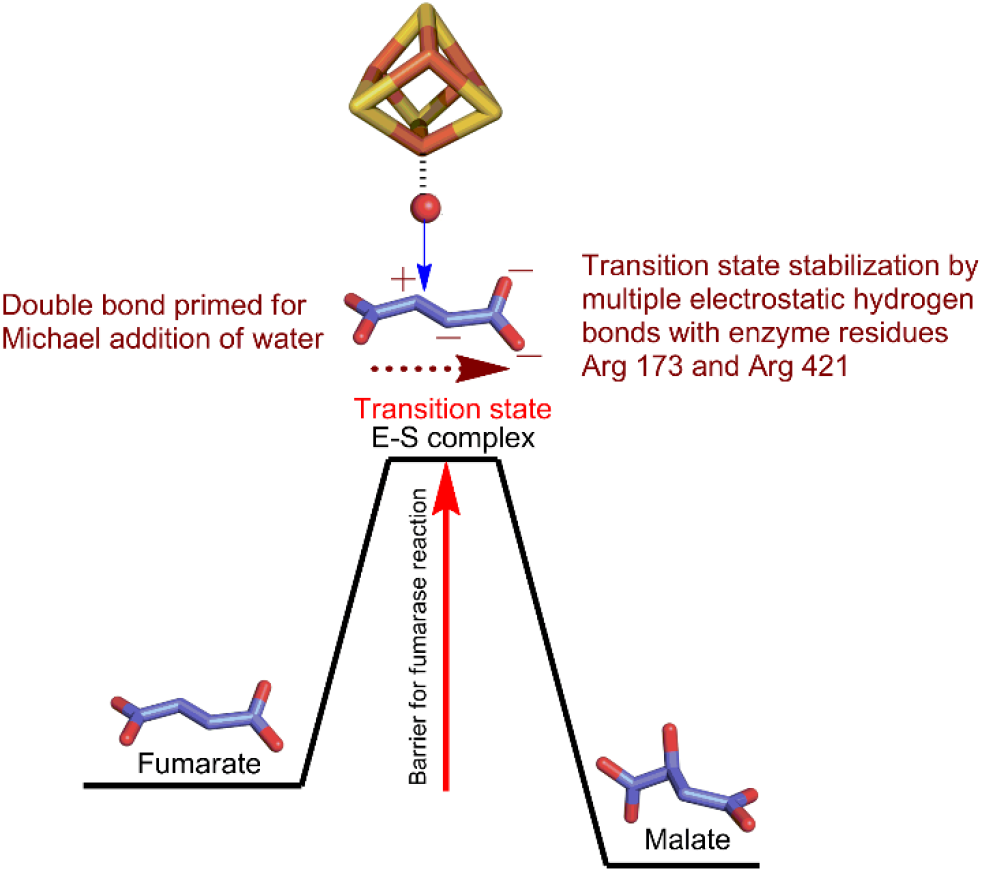
Mechanism of enzyme catalysis in FH. Upon binding to the active site of the enzyme, the substrate is raised to its transition state mediated by the positively charged residues on both sides of the substrate strongly polarizing it. The transition state complex is well primed for an attack of water molecule held in place by the Fe-S cluster, following which the product is released.

Fumarate hydratase is an example of a true “Pauling enzyme”^74^. The Fe-S cluster may have been retained as a part of enzyme evolution, and only plays a non-redox role in holding the water molecule in place for catalysis, similar to that observed in the tungsten containing enzyme, acetylene hydratase.^73^ Pre-organized polar environments of active sites have been invoked to drive enzymatic catalysis through electrostatic stabilization of the substrate.^75^ Recent studies on ketosteroid isomerase, which is an exceptional enzyme that stabilizes the substrate in the transition state by 21 kcal mol^−1^ have shown the role of electrostatic field at the active site on enzyme catalysis. Using vibrational stark effect spectroscopy, it was observed that the electrostatic field at the active site contributes significantly to reduction in free energy barrier and subsequently, its catalytic rate. Of the 11.5 kcal mol^−1^ activation barrier, reduction of 7.3 kcal mol^−1^, which corresponds to 70% of the enzyme catalytic rate compared to uncatalyzed reaction, was attributed to the electrostatic field alone. Positioning of the catalytic base, an aspartate residue contributed to the rest of the catalytic rate enhancement.^76^ Probing enzymes for electric field effects as a criterion for rate enhancements can provide useful insights on some of the exceptional enzymes in nature that carry out reactions at enormous rates, which otherwise would not have been possible. Although, catalysis in the view of electrostatic transition state stabilization has received good support over the years, theoretical studies backed by reproducible experimental observations are needed to a large extent to address the larger problem of the origin of enzyme catalysis.

## CONCLUSION

The extraordinary proficiency of fumarate hydratases in catalyzing the stereospecific addition of water to the planar, symmetrical, dianionic substrate, fumarate is in large measure due to the constellation of positively charged residues that line the pre-organized active site, polarizing the substrate electron distribution such that in the transition state, the olefinic double bond distorts to the aci-carboxylate form. Dramatic electrostatic stabilization of transition states in enzyme catalysis is provided by the example of ketosteroid isomerase, where the activation barrier is reduced by an estimated 21 kcal mol^−1^.^76^ Fumarate hydratases may provide an even more extreme example of transition state stabilization, envisaged decades ago by Polanyi, Haldane and Pauling and elaborated in later years by the seminal work of Jencks.^74,77–79^ The studies described above also provide firm experimental and theoretical support for the molecular mechanisms which enable fumarate hydratase to bear the energetic burden of catalysis clearly enunciated in the work of Bearne and Wolfenden.^4^

## ABBREVIATIONS

FH: fumarate hydratase
Mj: *Methanocaldococcus jannaschii*
Lm: *Leishmania major*
DFT: density function theory
NPA: natural population analysis
MD: molecular dynamics
NTD: N-terminal domain
CTD: C-terminal domain

## ASSOCIATED CONTENT

### Supporting information(attached)

All data described in the article are contained within the article/Supporting Information. MjFHαβ apo-protein complex (7xky) and MjFHβ protein (5dni) structures are deposited and available in the RCSB database. Additional experimental details, materials, and methods, modelling of missing residues/fragments from MjFH and LmFH for MD simulations, docking 4Fe-4S cluster into apo-MjFH, metadynamics simulation of holo-MjFH, variations between apo- and holo-forms of MjFH and LmFH, quantum chemical calculations on reaction mechanism of fumarate hydration in solution and at enzyme active site, cartesian coordinates.

## AUTHOR INFORMATION

**Asutosh Bellur** - structure solution of MjFH full-length apo-protein, construction of MjFH site directed mutants, kinetics of MjFH wild type and mutants, structure analysis, interpretation of data, manuscript writing

**Vijay Jayaraman** - cloning of MjFH and heterologous expression, purification, activity measurements, structure solution of MjFH β-subunit and full-length apo-protein, biochemical characterization.

**Arpitha Suryavanshi** - generation of site-directed mutants.

**Soumik Das** - computation: DFT and natural population analysis, manuscript writing

**Garima Jindal** - computation: DFT and natural population analysis, manuscript writing

**Sudarshan Behera** - computation: MD simulation and Metadynamics, manuscript writing

**S. Balasubramanian** - computation: MD simulation and Metadynamics, manuscript writing

**P. Balaram** - conceptualization, structure analysis, manuscript writing

**Hemalatha Balaram** - Funding, conceptualization, interpretation and analysis of all data, manuscript writing.

### Notes

The authors declare no competing financial interest.

## ACKNOWLEDGEMENTS

The diffraction data for MjFHβ was collected at the X-ray facility at Molecular Biophysics Unit, Indian Institute of Science, Bengaluru, India, and data for Apo-MjFHαβ was collected at the BM-14 beamline of European Synchrotron Radiation facility (ESRF), Grenoble, France. We acknowledge the assistance of Jyothi Kunala with analytical size exclusion chromatography. HB acknowledges financial support by the Department of Biotechnology, Ministry of Science and Technology, Government of India Grants BT/PR11294/BRB/10/1291/2014, BT/PR13760/COE/34/42/2015, and BT/INF/22/SP27679/2018; Science and Engineering Research Board (SERB), Department of Science and Technology, Government of India Grant CRG/2019/004150/IBS, EMR/2014/001276; JC Bose Fellowship, SERB and institutional funding from JNCASR. GJ acknowledges the funding (SRG/2019/001646) from the Science and Engineering Research Board (SERB). Computing time from Supercomputer Education and Research Centre (SERC), IISc, Bangalore is also gratefully acknowledged. SB thanks CSIR for fellowship and acknowledges gratefully *the support and the resources provided by ‘PARAM Yukti Facility’ under the National Supercomputing Mission, Government of India at the Jawaharlal Nehru Centre For Advanced Scientific Research*. AB was supported by CSIR for junior and senior research fellowships, and from JC Bose fellowship awarded to HB.

