## Supporting information for "Revisiting the burden borne by fumarase: enzymatic hydration of an olefin"

**Table of contents:**

1. **MATERIALS**

2. **EXPERIMENTAL PROCEDURES**
   Protein expression and purification
   Interaction of MjFHα and MjFHβ subunits
   Enzyme activity
   Crystallization

3. **COMPUTATIONAL DETAILS**
   Modelling of missing residues/fragments
   Simulation of holo-LmFH
   Simulation of apo-LmFH
   Simulation of apo-MjFH
   Docking of 4Fe-4S cluster and substrate into apo-MjFH
   Metadynamics simulation of holo-MjFH to obtain contracted structure
   Results from docking of the cluster and malate into apo-MjFH

4. **QUANTUM CHEMICAL CALCULATIONS**

5. **SCHEMES**
   **Scheme S1.** Three plausible pathways of interconversion between malate and fumarate in aqueous medium.
   **Scheme S2.** Stepwise mechanism of carbanion pathway (QM1A) and carbocation pathway (QM1B) inside the protein environment.
   **Scheme S3.** Concerted pathway (QM1C) of malate-fumarate conversion in QM1 cluster model and optimized geometry of the corresponding transition state. Distances are in Å.
   **Scheme S4.** Altered mechanism of malate-fumarate interconversion in T467A (M1) and D135A (M2) mutant via carbanion pathway.

6. **TABLES**

**Table S1.** Oligonucleotide sequences used for cloning *M. jannaschii* WT FH and generating site-directed mutants.

**Table S2.** Refinement statistics of MjFH β and MjFHαβ protein structures.

**Table S3.** Comparison of subunit/domain interfaces between MjFH and LmFH.

**Table S4.** Covalent catalytic contacts in holo-LmFH crystal structure (CS) and holo-MjFH docked structure (DS).

**Table S5.** Non-covalent catalytic contacts in holo-LmFH crystal structure (CS) and holo-MjFH docked structure (DS). M stands for malate, for example MO1 means O1 atom of malate.

**Table S6.** Details of systems studied using MD simulations.

**Table S7.** Size comparison between LmFH and MjFH.

7. **FIGURES**

**Figure S1.** Helix-helix interaction between α and β subunits stabilizes the C-terminal helix in β subunit of the protein complex.

**Figure S2.** Overlay of 4Fe-4S cluster from the crystal structure of holo-LmFH onto that from apo-MjFH shows that the active site pocket with three cysteine residues are well poised to accommodate the cluster.

**Figure S3.** Motif KGXGS is identical and conserved across all class-I FH.

**Figure S4.** Circular dichroism spectra of 4Fe-4S reconstituted wild type and mutants of MjFH.

**Figure S5.** Modelling and WTMetaD simulation of missing 63 residue peptide stretch from LmFH.

**Figure S6.** Validation of force field parameters for 4Fe-4S cluster and malate.

**Figure S7.** The Rg and inter-CTD distance values plotted against simulation time for holo-MjFH-1.

**Figure S8.** Contacts of LmFH active site with the bound substrate, malate.

**Figure S9.** CV, Rg and inter-CTD distance evolution from metadynamics run of holo-MjFH.

**Figure S10.** Ligand docked in the cavity of MjFH.

**Figure S11.** Superposition of apo- and holo- Mj and LmFH structures to observe difference in structures post MD.

**Figure S12.** Optimized geometries of the transition states involved in carbocation pathway (w2) and concerted pathway (w3). Distances shown are in Å.

**Figure S13.** Electronic energy profiles for different plausible pathways in aqueous medium.

**Figure S14.** Optimized geometries of the transition states in carbanion pathway (QM1A) and carbocation pathway (QM1B) for QM1 cluster model. Distances are in Å.

**Figure S15.** Electronic energy profiles of various plausible pathways obtained from QM1 model.

8. **SUPPORTING REFERENCES**

9. **CARTESIAN COORDINATES**
1. **MATERIALS**

Restriction enzymes, Phusion polymerase and T4 DNA ligase were obtained from New England Biolabs, USA. Primers were custom synthesized from Sigma Aldrich, Bengaluru. Media components for growing *E. coli* cultures were from Himedia, Mumbai, India. Akta HPLC and Q-sepharose resins were from GE Healthcare Life Sciences, USA. Selenomethionine was obtained from Cayman chemicals Co., Michigan. Crystallization cocktails were from Hampton Research, Aliso Viejo, CA. *RS*-2-thiomalate was obtained from Sigma Aldrich, USA. All enzyme assays were monitored using Hitachi U2010 (Hitachi High Technologies, Japan) spectrophotometer fitted with water-circulated cell holder.

2. **EXPERIMENTAL PROCEDURES**

**Protein expression and purification**

Recombinant expression of MjFHαβ was carried out by transforming the *E. coli* strain BL21 (DE3)-RIL with the plasmid pETduet_MjFHαβ, followed by selection on Luria-Bertani (LB) agar plate containing ampicillin (100 µg $ml^{-1}$) and chloramphenicol (100 µg $ml^{-1}$). Multiple colonies on the plate were picked and transferred to 10 ml LB broth. The culture was grown overnight at 37 $^{0}$C and 1% of inoculum was added to 800 ml of Terrific broth (TB). The cells were grown at 37 $^{0}$C to an $OD_{600}$ of 0.45, induced with 0.3 mM IPTG and grown further for 18 hours at 16 $^{0}$C. The cells were harvested by centrifugation and resuspended in lysis buffer containing 50mM Tris HCl, pH 8.0, 5% glycerol, 1mM DTT and 1 mM PMSF. Cell lysis was achieved by five cycles of French press (Thermo IEC Inc., USA) at 1000 psi, and the lysate was clarified by centrifugation at 30,000 x g for 30 minutes. Supernatant was incubated at 70 $^{0}$C for 30 minutes to precipitate the bacterial proteins and again clarified by centrifugation at 30,000 x g for 30 minutes. Supernatant was then treated with 0.01% PEI to precipitate nucleic acids and clarified by centrifugation at 30,000 x g for 30 minutes. The clarified lysate was filtered through a 0.44 micron filter and loaded onto Q-sepharose anion-exchange column. The protein was eluted using a linear gradient of NaCl in buffer containing 50 mM Tris HCl, pH 8.0, 5% glycerol and 1mM DTT. Fractions containing co-purified α and β-subunits of the protein were pooled, dialyzed, and stored at -80 $^{0}$C. The purified protein was analyzed by SDS-PAGE [1] and protein concentration was determined by the Bradford method [2] with bovine serum albumin (BSA) as standard. The same protocol was adopted for purification of all the mutants.

Expression and purification of MjFHαβ with selenomethionine incorporated for single-wavelength anomalous dispersion (SAD) phasing, was carried out following established protocols.[3] Overnight grown culture of *E. coli* strain BL21 (DE3)-RIL containing pETduet_MjFHαβ was centrifuged, and the cells washed with M9 minimal medium. 0.5% culture was inoculated in pre-warmed M9 minimal medium containing 5 $gL^{-1}$ glucose as carbon source. Cultures were grown till mid-log phase at 37 $^{0}$C followed by addition of inhibitory amino acid mix containing phenylalanine, lysine, and threonine at a concentration of 100 $mgL^{-1}$ and leucine, isoleucine, and valine at a concentration of 50 $mgL^{-1}$. Cultures were grown for 15 minutes followed by addition of selenomethionine to a final concentration of 100 $mgL^{-1}$ and grown for further 15 minutes. The cultures were then induced with 0.3 mM IPTG and grown at 20 $^{0}$C for 18 hours. The cells were harvested, and protein purified following the same protocol as that for the wild type (WT) protein using degassed buffer containing 50mM Tris HCl, pH 8.0, 5% glycerol and 10 mM DTT. The purified protein was checked on SDS-PAGE and complete incorporation of selenomethionine into the protein was validated by LC-ESI-MS.

**Interaction of MjFHα and MjFHβ subunits**

The oligomeric state of co-purified protein was probed by size-exclusion chromatography on an analytical Superdex 200 10/300 GL column (10 mm X 300 mm) (GE Health Care Life Sciences) attached to an AKTA Basic HPLC system. The column was equilibrated with buffer A containing 50 mM Tris-HCl, pH 7.4 and 100 mM KCl and calibrated using the molecular weight standards; *β*-amylase (200 kDa), alcohol dehydrogenase (150 kDa), bovine serum albumin (66 kDa), carbonic anhydrase (29 kDa) and cytochrome c (12.4 kDa). A standard curve was plotted using ratio of elution volume of standard proteins to void volume of column against logarithm of molecular mass. The flow rate was maintained at 0.5 ml $min^{-1}$. 100 µl of purified protein (10µM) (MjFHα, MjFHβ, and the co-expressed protein) was loaded onto the column equilibrated with buffer A containing 2mM β-mercaptoethanol or 1M KCl and the elution was monitored simultaneously at both 280 nm and 220 nm. The molecular mass of the MjFH subunits and that of the complex was calculated by interpolation of the elution volume on the standard curve.

**Enzyme activity**

The enzymatic conversion was monitored at different wavelengths as follows: 240 nm ($\varepsilon_{240}$ = 2440 $M^{-1}cm^{-1}$) [4] for fumarate concentrations up to 750 µM; 250 nm ($\varepsilon_{250}$ = 1345 $M^{-1}cm^{-1}$) for 1 mM; 260 nm ($\varepsilon_{260}$ = 805 $M^{-1}cm^{-1}$) for 1.5 mM; 270 nm ($\varepsilon_{270}$ = 475 $M^{-1}cm^{-1}$) for 2 mM; 280 nm ($\varepsilon_{280}$ = 240 $M^{-1}cm^{-1}$) for 3 mM; 290 nm ($\varepsilon_{290}$ = 110 $M^{-1}cm^{-1}$) for 5 mM; 300 nm ($\varepsilon_{300}$ = 45 $M^{-1}cm^{-1}$) for 10 and 20 mM; 305 nm ($\varepsilon_{305}$ = 18 $M^{-1}cm^{-1}$) for concentrations of 30 and 40 mM in a quartz cuvette of 1 cm path length.

Inhibition kinetics of the enzyme with *RS*-2-thiomalate was carried out using similar assay conditions with fixed concentrations of fumarate (500 µM) and malate (1000 µM) in 50 mM Tris-HCl, pH 7.2 at varying concentrations of *RS*-2-thiomalate. Apparent inhibitory constant $K_i$ (app) was calculated from the $IC_{50}$-to-Ki web-server [5] for *RS*-2-thiomalate for MjFH with malate and fumarate as substrates.

All initial velocity measurements were conducted in duplicate with data points in the plot and values derived being mean ± S.E. The initial rate *vs* substrate concentration plots were generated using GraphPad Prism version 5.0 (Graphpad software Inc) and non-linear regression method was used to fit the data points for WT and mutants using the equation, $v=V_{\max}[S]/[K_M + [S](1+[S]/K_I)]$ **(eq. 1)** that incorporates substrate inhibition. $IC_{50}$ was determined by fitting the data to the equation $v=V_0/[1 + (I/IC_{50})]$ **(eq. 2)**, where $v$ is the initial velocity, $V_{\max}$ is maximum initial velocity, [S] is substrate concentration, $K_I$ is the dissociation constant for equation 1 and $v$ is the observed velocity, $V_0$ is the uninhibited velocity, [I] is the inhibitor concentration and $IC_{50}$ in equation 2 is the 50% inhibitory concentration of the inhibitor.

**Crystallization**

A 72 well multi-well plate from Grenier-Bio was used. Conditions for crystallization were obtained by using all the conditions in the commercially available crystallization kit from Hampton (Hampton Research, USA). In case of MjFHαβ holo-protein, reconstituted protein was passed through a Zeba™ Spin Desalting Column, 7K MWCO, to remove unbound iron and sulphur and set up for crystallization within the anaerobic chamber. For MjFHαβ apo-protein, to ensure a homogenous population, the protein was subjected to apo-protein preparation. Apo-protein was prepared by addition of 50x excess EDTA and 20x excess $K_3[Fe(CN)_6]$ followed by incubation for 30 minutes. The protein was concentrated by precipitation with 100% ammonium sulfate and dissolved in a minimum volume of 20 mM potassium phosphate, pH 6.4 and dialyzed. Various concentrations ranging from 3 mg/ml to 24 mg/ml of the protein were used for setting up crystal screens. The crystallization droplet contained 3 μl of protein and 3 μl of buffer from different conditions placed under a 50% mix of paraffin and silicone oil at room temperature.

#### 3. COMPUTATIONAL DETAILS

##### Modelling of the missing residues/fragments

The crystal structure of *Leishmania major* fumarate hydratase (LmFH) was obtained from Protein Data Bank (PDB ID - 5l2r). Each chain of the crystal structure has 63 missing residues from N-terminus, and a missing ten residues linker, which connects the N-terminal and C-terminal domains. Other than these missing residues, there are 112 missing atoms (non-hydrogen atoms) from 42 different residues. These missing atoms are built using the Molefacture plugin in VMD.[6] The ten residues linker is modelled using the SWISS-MODEL server [7] whereas the modelling of the 63 missing residues from the N-terminus is elaborated below. The SWISS-MODEL server could not find a suitable model for the 63 missing residues structure. Hence the missing structure from N-terminus is modelled in two parts: one fragment containing 34 residues and the other 31 residues. These two fragments have two residues in common. After modelling these two parts individually from the SWISS-MODEL server, both were connected (by removing the two common residues) to produce a complete 63 residues structure (**Figure S5 A**). Hydrogens were added to the residues of the 63 residues peptide based on their protonation state at pH 7, and then the system was soaked in water followed by the addition of counter ions to achieve charge neutrality. The peptide, with a position restrained force constant of 1000 $kJmol^{-1}nm^{-2}$ on non-hydrogen atoms, was subjected to energy minimization using conjugate gradient algorithm.[8] The temperature was slowly increased to 300 K in five steps, with each step reducing the force constant value of position restrain, and the value becomes 0 $kJ.mol^{-1}$ $nm^{-2}$ at the final step, in the NVT ensemble. The temperature was maintained using Bussi-Donadio-Parrinello velocity rescaling thermostat [9] with a coupling constant of 0.1 ps. NPT equilibration was then performed for 10 ns at 1 bar and 300 K using Berendsen barostat [10] with a coupling constant of 0.5 ps with the same thermostat parameters. Following equilibration, well-tempered metadynamics (WTMetaD) simulation [11] was performed using the radius of gyration (Rg) of the peptide as the collective variable to obtain a globular structure of the peptide. The WTMetaD simulation was performed with Gaussian hill (σ) of 1 Å, initial hill height = 1.2 kJ/mol, deposited every 500 time steps, using a bias factor of 6. The structure of the peptide corresponding to the Rg value of 12 Å is considered as the final structure (**Figure S5 B**). A wide range of Rg values were sampled from the WTMetaD simulation of the 63 residues peptide, which can be discerned from the time evolution of Rg (**Figure S5 C**).

##### Simulation of holo-LmFH

The final structure of the 63-residue peptide is then linked with one chain of holo-LmFH, to which ten missing residues and missing atoms have already been added. It was aligned with the second chain, and then the coordinates of the missing residues and atoms from the first chain were added to the second chain. Further simulations were performed on the dimer of holo-LmFH. The hydrogen addition, energy minimization, and equilibration protocol for the final holo-LmFH dimer structure is the same as that of the 63 residues modelled peptide. Following equilibration, a production simulation run for 200 ns was carried out at ambient conditions, in the NPT ensemble, using Bussi-Donadio-Parrinello velocity rescaling thermostat [9] and Parrinello-Rahman barostat [12] with a coupling constant of 0.1 ps and 0.5 ps, respectively. CHARMM36m force field [13] was employed for the enzyme, whereas force field parameters for malate were generated using the CGenFF server.[14] Force-field parameters for the 4Fe-4S cluster, and the cysteine residues bound to it were taken from the article by Chang et al.[15] The equilibrium bond lengths and angle values for the bonds and angles formed between the 4Fe-4S cluster and malate are set to be the same as in the crystal structure of holo-LmFH. The other parameters are set as the corresponding values of 4Fe-4S and cysteine link. For example, the force constant for Fe-O is set to be the same as that for Fe-S. All the new dihedrals formed because of 4Fe-4S cluster and malate link are left free to rotate. GROMACS-2018.3 [16] was used for all the simulations employing three-dimensional periodic boundary conditions. All the bonds were constrained using the LINCS algorithm,[17] and PME [18] was used for electrostatic interaction energy calculations. A cutoff value of 12 Å was used for both electrostatic and van der Waals interactions. The treatment of solvation was carried out using the TIP3P water model.[19] Leap-frog integrator [20] with 2 fs timestep was employed for all the molecular dynamics simulations, and both the coordinates and velocities were dumped every 10 ps.

**Simulation of apo-LmFH**

From the holo-LmFH structure, after the addition of missing atoms and residues, the 4Fe-4S cluster and substrate were removed. Hydrogen atom additions (H-atoms were also added to the three cysteine residues that were bound to the iron-sulfur cluster in the holo-LmFH), energy minimization and equilibration were performed using the same protocol as mentioned for the MD simulation of 63 residues modelled peptide. A production trajectory for 800 ns was generated using the same parameters as used for the production run of holo-LmFH.

**Simulation of apo-MjFH**

In the crystal structure of apo-MjFH, one and two residues from the N-terminus and C-terminus of each chain are missing, respectively. There are also some missing atoms in the apo-MjFH structure. The missing atoms are modelled similarly as in the LmFH structure, whereas the missing residues are modelled using Pymol.[18] Applying the same simulation protocol as used for apo-LmFH, apo-MjFH was simulated for 200 ns of production run.

**Docking of 4Fe-4S cluster and substrate into apo-MjFH**

The 3D structures of the 4Fe-4S cluster and malate were obtained from the crystal structure of LmFH (PDB ID-5l2r). The ligand (4Fe-4S cluster + malate) and the receptor (one chain of MjFH (α and β subunits) with missing residues and atoms added) were initialized by adding hydrogens, adding Gasteiger charges, merging non-polar hydrogens using AutoDock Tools.[21] Rotatable bonds were set for the ligand, and both ligand and receptor were written in PDBQT format to be read by AutoDock software.[21] AutoGrid was employed to prepare a grid with grid size 40 x 40 x 40 points with a grid spacing of 0.375 Å cantered at dimensions (x, y and z): 26.72, 7.3 and 17.0, which covers the entire active site regions. Docking was performed using the Lamarckian Genetic Algorithm with a population size of 150 and a maximum number of energy evaluation to be 2500000. Default values were used for other parameters. The structural model with the highest ligand binding energy was chosen for further analysis and MD simulations. After the best docked structure was obtained for one chain of MjFH, it was overlayed with the second chain using VMD [6] and coordinates of the ligand were written. The coordinates of the ligand were then added to the second chain.

The structure of MjFH, after the addition of missing atoms and residues and the docking of the ligand, was subjected to the same simulation protocol as in the simulation of holo-LmFH, and a 200 ns of trajectory was generated for production analysis. This form is referred to as *holo-MjFH-1* hereafter.

**Metadynamics Simulation of holo-MjFH to obtain a contracted structure**

The metadynamics simulation [22] for obtaining a compact holo-MjFH structure was carried out in multiple steps. First, we used the linear combination of the 14 catalytic contacts mentioned in **Figure S8 B** as CV in the metadynamics simulation and it did not result in a stable contracted form of holo-MjFH (results not presented here). To find out more contacts which can be biased to obtain a stable contracted form of holo-MjFH, the following protocol is followed: (i) all the residues which are within 10 Å distance from the heavy atoms of the cluster and malate in holo-LmFH were considered; these are 89 in total, (ii) 51 out of these 89 residues are found to be conserved in MjFH (iii) 28 close contacts other than the 14 catalytic contacts (**Figure S8 B**) were recognized in holo-LmFH by visual inspection of the interactions of the 51 conserved residues with the cluster and substrate.

A linear combination of these 28 additional pair distances and 14 catalytic contacts per chain (42 per chain and 84 in total), with coefficients of 1, was considered a collective variable (CV) to bias in the *first* metadynamics simulation. A Gaussian hill height of 3 kJ/mol was chosen to observe the contraction of the global structure in a short simulation time. Sigma value of 4 Å was chosen as the half of the fluctuation of CV from the MD simulation of holo-MjFH-1. Gaussian hills were deposited every 500 steps, and a wall with a force constant of 10000 $kJmol^{-1}nm^{-2}$ was applied at 650 Å to avoid further expansion of the enzyme. The metadynamics simulation was run for 30 ns. A contracted form of the structure was extracted from the frame at 26.1 ns (**Figure S9 A**) with Rg and inter-CTD distance values of 30.80 Å and 58.7 Å, respectively (**Figure S9 B** and **C**).

In the next step, the 56 pair distances (other than the 28 catalytic contacts in both the chains) were restrained with force constant of 1000 $kJmol^{-1}nm^{-2}$ and the *second* metadynamics simulation was performed considering the linear combination of only the catalytic contacts (28 for both the chains) as the CV, with the same parameters as above. A wall with a force constant of 10000 $kJmol^{-1}nm^{-2}$ is applied at 85 Å to avoid further increase in the pair distances of catalytic contacts. A structure obtained from the frame at 1.25 ns (**Figure S9 D**) has all the catalytic contacts in hydrogen bonding distances. This structure was used for further equilibrations. The equilibration was done in four steps by decreasing the force constant of the distance restraints on all the 84 pair distances from 1000 to 0 $kJmol^{-1}nm^{-2}$ at the NVT ensemble. A final production run of 400 ns was performed in the NPT ensemble after 10 ns of NPT equilibration. The system is referred to as *holo-MjFH-2* hereafter. Refer to **Table S6** for the number of atoms, cubic box length and simulation length for the different enzyme forms simulated here.

##### Results from the docking of the cluster and malate into apo-MjFH

The holo-MjFH structure after docking and with the highest ligand binding energy, has the ligand very well fitted in the cavity of MjFH (**Figure S10 A and B**). The three cysteine residues (Cys60, Cys182, Cys269), which should covalently bind to three Fe of 4Fe-4S cluster, are close to covalent bonding distance from three Fe (**Figure S10 C and Table S4**). Like that of the crystal structure of LmFH, few catalytic residues (such as Lys104, Asp62, Thr81 and Gln61) are in close contact with the malate in the docked holo-MjFH structure (**Table S5**). However, since the active site region of apo-MjFH is structurally relaxed and some catalytic residues have moved away, the docking simulation could not capture the interactions of malate with them (**Table S5**). Visual inspections of other docked structures (total of 50) with lesser ligand binding energies were also carried out. No other structures represent as good a binding pose as the first one. A greater number of noncovalent catalytic contacts were seen in some structures, but at the expense of contact between cysteine residues of the enzyme and irons of the 4Fe-4S cluster. Since the close contacts between cysteine residues and irons are the most important, the first structure with the highest ligand binding energy was considered for further MD simulations.

#### 4. QUANTUM CHEMICAL CALCULATIONS

Apart from the nucleophilic mechanism involving a carbanion intermediate, we also performed a detailed analysis of two other plausible pathways i.e., carbocation pathway and concerted pathway in aqueous medium as well as in the active site of LmFH using smaller cluster model QM1. In aqueous medium, analogous to the carbanion pathway (Pathway **w1**), carbocation mechanism (Pathway **w2**) for malate (**A**) – fumarate (**C**) interconversion proceeds with two transition states (**Scheme S1**). In the concerted pathway (Pathway **w3**), we again considered the possibility of assistance via solvent water molecules and optimized two different concerted transition states with one ($\mathbf{w3^{1w}}$) and zero ($\mathbf{w3^{0w}}$) external water molecule for reversible (de)hydration. Although energetically, these pathways turned out to be unfavorable due to relatively higher activation energies (**Figure S13**).

In the smaller cluster model QM1 as well, we investigated two other possible pathways of olefinic (de)hydration apart from the favorable carbanion pathway (Pathway **QM1A**) although the carbocation mechanism (Pathway **QM1B**) and the concerted one (Pathway **QM1C**) was found to be even more energy demanding inside the protein environment (**Figure S15**). Stepwise mechanism of QM1A and QM1B are with all possible transition state intermediates can be observed in **Scheme S2 and S3**. The optimized geometries for TSs involved in the QM1A and QM1B pathway are shown in **Figure S14.**

### 5. SCHEMES

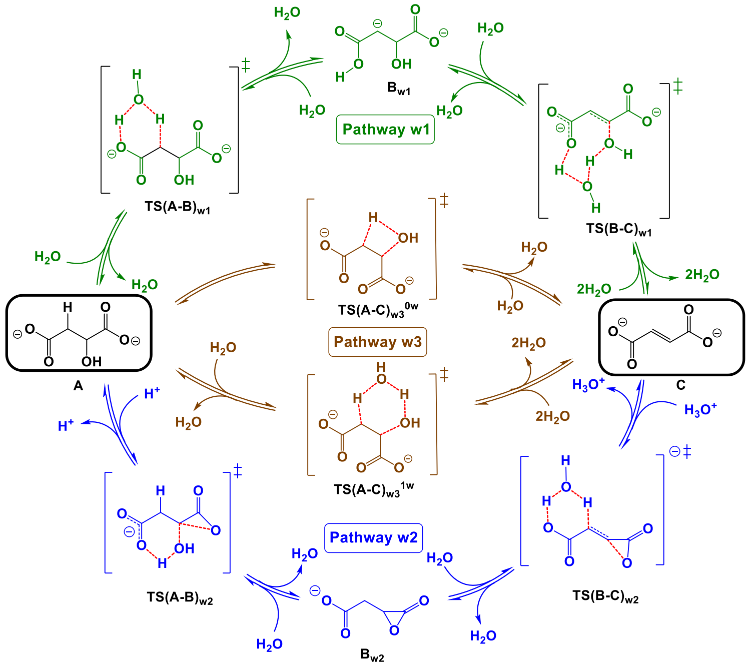


**Scheme S1.** Three plausible pathways of interconversion between malate and fumarate in aqueous medium.

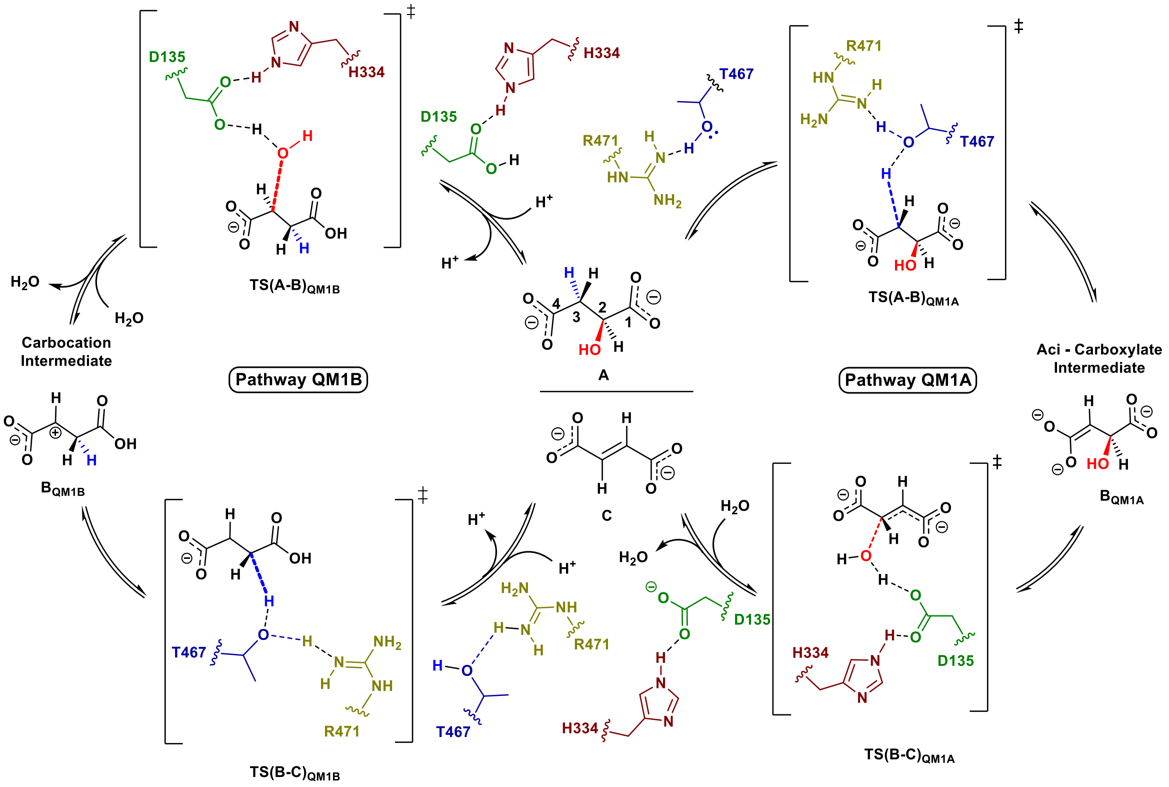


**Scheme S2.** Stepwise mechanism of carbanion pathway (**QM1A**) and carbocation pathway (**QM1B**) inside the protein environment.

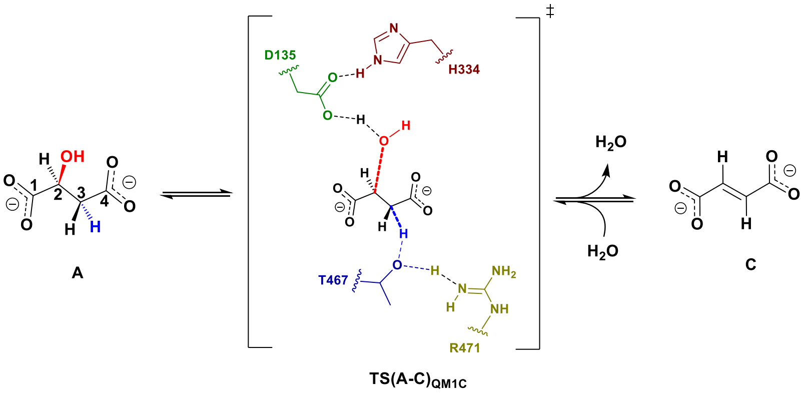


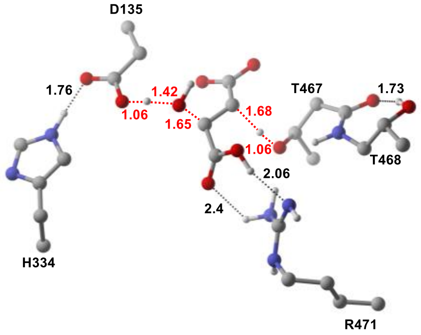


**TS(A-C)$_{QM1C}$**

**Scheme S3.** Concerted pathway (QM1C) of malate-fumarate conversion in QM1 cluster model and optimized geometry of the corresponding transition state. Distances are in Å.

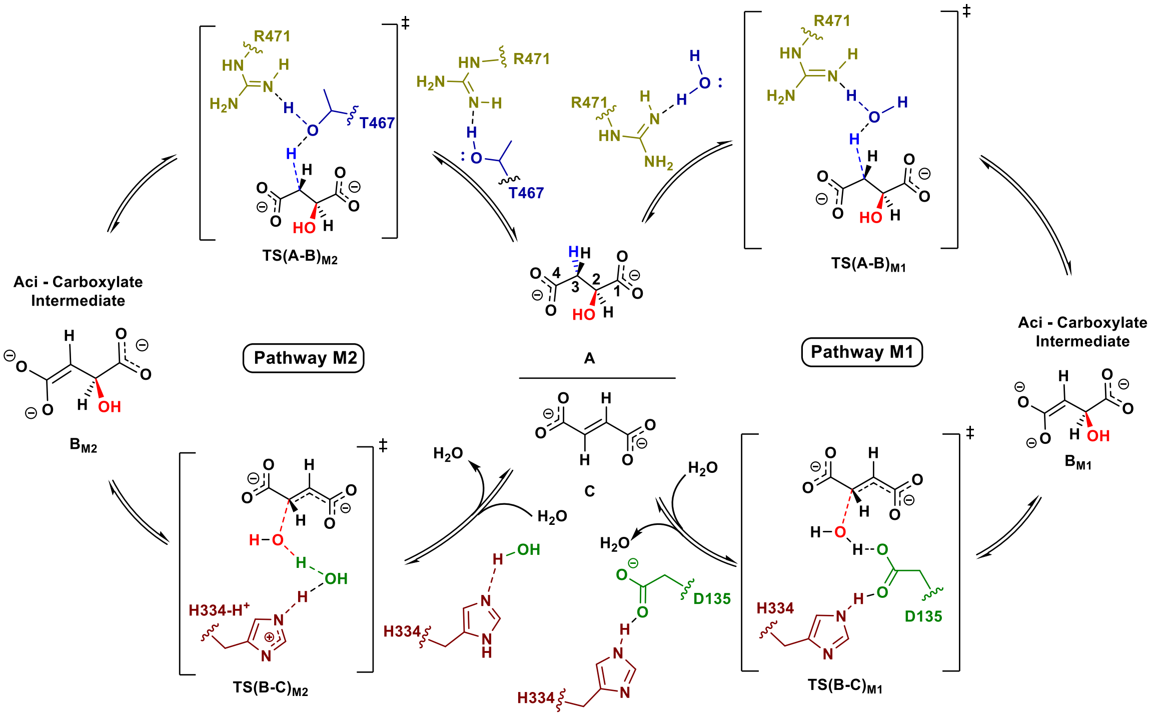


**Scheme S4.** Altered mechanism of malate-fumarate interconversion in T467A (M1) and D135A (M2) mutant via carbanion pathway.

#### 6. TABLES

**Table S1.** Oligonucleotide sequences used for cloning *M. jannaschii* WT FH and generating site-directed mutants.

| **Primer name** | **Primer sequence (5' to 3')** |
|---|---|
| MjFH_α_BamHI_up | CGCGGATCCGAAAATCTCCGATGTTGTTGTTGAATTATTTAG |
| MjFH α_SalI_down | ACGCGTCGACTTATAATTTAGCATCCAATTTTATTCTTTTAATTGCC |
| MjFH β_NdeI_up | CTAATTCCATATGGAATATACATTTAACAAATTAACAAAAAAGATG |
| MjFH β_XhoI_down | CCGCTCGAGTTATAATCCTATCAATTCATTAAGCTTTTCATAAAC |
| MjFH_D62V_FP | GTTCCTCTATGTCAAGTTACTGGTGTCCCAATAG |
| MjFH_D62V_RP | CTATTGGGACACCAGTAACTTGACATAGAGGAAC |
| MjFH_H257N_FP | GAGATTGCTGGATGCAACACAGCTTCTTTACCTGTAGG |
| MjFH_H257N_RP | CCTACAGGTAAAGAAGCTGTGTTGCATCCAGCAATCTC |
| MjFH_T80V_FP | GGTTTGTGTTTCTATAGGCCCAGTAACATCTGCAAGGATGAATG |
| MjFH_T80V_RP | CATTCATCCTTGCAGATGTTACTGGGCCTATAGAAACACAAACC |
| MjFH_T80C_FP | GGTTTGTGTTTCTATAGGCCCATGTACATCTGCAAGGATGAATG |
| MjFH_T80C_RP | CATTCATCCTTGCAGATGTACATGGGCCTATAGAAACACAAACC |
| MjFH_G147A_FP | GCATTTCCAAAAGGGGCAGCGAGCGAAAACATGAGTGC |
| MjFH_G147A_RP | GCACTCATGTTTTCGCTCGCTGCCCCTTTTGGAAATGC |
| MjFH_T81V_FP | GTGTTTCTATAGGCCCAACAGTTTCTGCAAGGATGAATGATG |
| MjFH_T81V_RP | CATCATTCATCCTTGCAGAAACTGTTGGGCCTATAGAAACAC |
| MjFH_R84L_FP | GCCCAACAACATCTGCATTAATGAATGATGTTGAAGAGG |
| MjFH_R84L_RP | CCTCTTCAACATCATTCATTAATGCAGATGTTGTTGGGC |
| MjFH_K104L_FP | GCAATTGTTGGACTGGGAGGAATGAAAAAAGAG |
| MjFH_K104L_RP | CTCTTTTTTCATTCCTCCCAGTCCAACAATTGC |
| MjFH_R102L_FP | GGAAGAGGTTCCTTTATTACCTAATGTAGTTCATCC |
| MjFH_R102L_RP | GGATGAACTACATTAGGTAATAAAGGAACCTCTTCC |
| MjFH_Q61L_FP | GAAACGCAAGTTCCTCTATGTTTAGATACTGGTGTCCCAATAG |
| MjFH_Q61L_RP | CTATTGGGACACCAGTATCTAAACATAGAGGAACTTGCGTTTC |
| MjFH_R32L_FP | GGCAAAATATACACTGCGTTAGATGAAGCACATTTAAAAATTATTG |
| MjFH_R32L_RP | CAATAATTTTTAAATGTGCTTCATCTAACGCAGTGTATATTTTGCC |
| MjFH_K144L_FP | GATAATTGCATTTCCATTAGGGGCAGGAAG |
| MjFH_K144L_RP | CTTCCTGCCCCTAATGGAAATGCAATTATC |
| MjFH_S148A_FP | GCATTTCCAAAAGGGGCAGGAGCCGAAAACATGAGTGC |
| MjFH_S148A_RP | GCACTCATGTTTTCGGCTCCTGCCCCTTTTGGAAATGC |
| MjFH_R32K_FP | GGCAAAATATACACTGCGAAAGATGAAGCACATTTAAAAATTATTG |
| MjFH_R32K_RP | CAATAATTTTTAAATGTGCTTCATCTTTCGCAGTGTATATTTTGCC |
| MjFH_R84K_FP | GCCCAACAACATCTGCAAAAATGAATGATGTTGAAGAGG |
| MjFH_R84K_RP | CCTCTTCAACATCATTCATTTTTGCAGATGTTGTTGGGC |
| MjFH_R102K_FP | GGAAGAGGTTCCTTTAAAACCTAATGTAGTTCATCC |
| MjFH_R102K_RP | GGATGAACTACATTAGGTTTTAAAGGAACCTCTTCC |

**Table S2.** Refinement statistics of MjFH β and MjFHαβ protein structures.

| **Protein** | **MjFHβ** | **MjFHαβ** |
|---|---|---|
| PDB ID | 5DNI | 7XKY |
| Space group | $P6_1$ | C121 |
| a, b. c (Å) | 85.61 85.61 121.83 | 116.95 61.70 83.88 |
| α, β, γ ($^0$) | 90.00 90.00 120.00 | 90.00 123.90 90.00 |
| Molecules per asymmetric unit | 2 | 2 |
| Crystallization conditions | 0.1 M Bis-Tris, pH 5.5, and 25 % polyethylene glycol-3350 | 0.2 M magnesium chloride hexahydrate, 0.1 M BIS-TRIS, pH 5.5, 25% w/v polyethylene glycol 3,350 |
| | **Data collection**[a] | |
| Wavelength (Å) | 1.54 | 0.9537, 1.54 |
| Resolution (Å) | 47.07-2.23 (2.30-2.23) | 52.07-2.46 (2.54-2.46) |
| Total reflections | 109381 (15166) | 142561 (19194) |
| Unique reflections | 22550 (3282) | 17961 (2578) |
| Multiplicity | 4.9 (4.6) | 7.9 (7.4) |
| Completeness (%) | 100.0 (100.0) | 98.10 (97.50) |
| Mean I/σ (I) | 7.0 (1.6) | 11.2 (3.9) |
| Wilson B factor (Å$^2$) | 39.6 | 30.3 |
| $R_{merge}$ | 0.127 (0.865) | 0.124 (0.525) |
| $R_{measure}$ | 0.143 (0.978) | 0.145 (0.598) |
| $CC_{1/2}$ | 0.994 (0.709) | 0.976 (0.871) |
| | **Refinement** | |
| $R_{work}$ | 0.185 | 0.205 |
| $R_{free}$ | 0.233 | 0.245 |
| Number of non-hydrogen atoms | 2927 | 3567 |
| $RMS_{bonds}$ | 0.008 | 0.015 |
| $RMS_{angles}$ | 0.176 | 2.05 |
| Ramachandran | | |
| Favoured (%) | 94.07 | 96.18 |
| Outliers (%) | 0.56 | 3.14 |
| Clashscore | 3.25 | 6 |
| Average B-factor | 45.26 | 34.34 |

[a]Values of highest resolution shell are shown in parenthesis

**Table S3.** Comparison of subunit/domain interfaces between MjFH and LmFH.

| Organism | Chains | Interface residues | Interface area (Å$^2$) | Salt bridges | Hydrogen bonds | Van der Waals interactions |
|---|---|---|---|---|---|---|
| *M. jannaschii* | A : B | 21 : 19 | 1275 : 1295 | - | 5 | 74 |
| | A : C | 48 : 48 | 2521 : 2521 | 2 | 34 | 241 |
| *L. major* | A : B | 31 : 34 | 1503 : 1520 | - | 19 | 193 |
| | A : C | 77 : 76 | 3712 : 3718 | 10 | 42 | 494 |
| | A : D | 2 : 3 | 310 : 271 | - | - | 11 |

*Chains A and C refer to α-subunit of MjFH and NTD of LmFH, and chains B and D refer to β-subunit of MjFH and CTD of LmFH**.** LmFH reveals a larger area of interface interactions compared to MjFH.

**Table S4.** Covalent catalytic contacts in holo-LmFH crystal structure (CS) and holo-MjFH docked structure (DS). Refer to **Figure S8 A** for residue and atom names.

| Contact ID | holo-LmFH (chain A) | | holo-MjFH (chain A) | |
|---|---|---|---|---|
| | Contact | Distance in CS (Å) | Contact | Distance in DS (Å) |
| **1** | Cys133 - Fe1 | 2.33 | Cys60 - Fe1 | 2.55 |
| **2** | Cys252 - Fe3 | 2.29 | Cys182 - Fe3 | 2.08 |
| **3** | Cys346 - Fe2 | 2.37 | Cys269 - Fe2 | 1.79 |

**Table S5.** Non-covalent catalytic contacts in holo-LmFH crystal structure (CS) and holo-MjFH docked structure (DS). M stands for malate, for example MO1 means O1 atom of malate. Refer to **Figure S8 A** for residue and atom names.

| Contact ID | holo-LmFH (chain A) | | holo-MjFH (chain A) | |
|---|---|---|---|---|
| | Contact | Distance in CS (Å) | Contact | Distance in DS (Å) |
| **1** | Lys491 NZ - MO1 | 2.68 | Lys104 NZ - MO1 | 2.56 |
| **2** | His334 NE2 -Asp135 OD1 | 2.81 | His257 NE2 - Asp62 OD1 | 2.93 |
| **3** | Asp135 OD2 - MO | 2.64 | Asp62 OD2 - MO | 4.27 |
| **4** | Gly216 N - MO | 3.10 | Gly147 N - MO | 5.09 |
| **5** | Arg173 NH2 - MO3 | 2.86 | Arg102 NH2 - MO3 | 9.41 |
| **6** | Arg173 NH1 - MO4 | 2.81 | Arg102 NH1 - MO4 | 12.22 |
| **7** | Arg421 NH2 - MO3 | 2.78 | Arg32 NH2 - MO3 | 5.15 |
| **8** | Arg421 NH1 - MO4 | 2.92 | Arg32 NH1 - MO4 | 7.64 |
| **9** | Gln134 NE2 - MO4 | 2.96 | Gln61 NE2 - MO4 | 4.45 |
| **10** | Thr467 OG1 - MC2 | 3.34 | Thr80 OG1 - MC2 | 7.26 |
| **11** | Thr467 OG1 - MO2 | 2.75 | Thr80 OG1 - MO2 | 8.31 |
| **12** | Thr468 OG1 - MO2 | 2.89 | Thr81 OG1 - MO2 | 5.64 |
| **13** | Thr468 N - MO2 | 3.12 | Thr81 N - MO2 | 4.20 |
| **14** | Arg471 NH2 - MO2 | 2.92 | Arg84 NH2 - MO2 | 7.29 |

**Table S6.** Details of systems studied using MD simulations

| Enzyme | Form | Number of atoms | Cubic box length (nm) | Simulation Length (ns) |
|---|---|---|---|---|
| holo-LmFH | dimer | 5,34,194 | 17.40 | 200 |
| apo-LmFH | dimer | 5,33,769 | 17.36 | 800 |
| apo-MjFH | dimer of a dimer | 2,05,371 | 12.63 | 200 |
| holo-MjFH-1 | dimer of a dimer | 2,06,056 | 12.65 | 200 |
| holo-MjFH-2 | dimer of a dimer | 2,06,056 | 12.65 | 400 |

**Table S7.** Size comparison between LmFH and MjFH. The quantities from simulation are the mean and standard deviation obtained from the last 100 ns. The Rg for LmFH is calculated excluding the highly flexible 63-residues modeled fragment.

| Enzyme | Rg (Å) | Inter-CTD distance (Å) |
|---|---|---|
| holo-LmFH (Crystal structure) | 30.3 | 56.85 |
| apo-MjFH (Crystal structure) | 31.8 | 65.63 |
| holo-LmFH (Simulation) | 30.8 ± 0.1 | 57.4 ± 0.4 |
| apo-LmFH (Simulation) | 33.4 ± 0.2 | 73.6 ± 1.2 |
| apo-MjFH (Simulation) | 33.0 ± 0.3 | 68.7 ± 1.2 |
| holo-MjFH-1 (Simulation) | 33.7 ± 0.2 | 68.5 ± 1.1 |
| holo-MjFH-2 (Simulation) | 31.2 ± 0.1 | 61.0 ± 0.6 |

#### 7. FIGURES

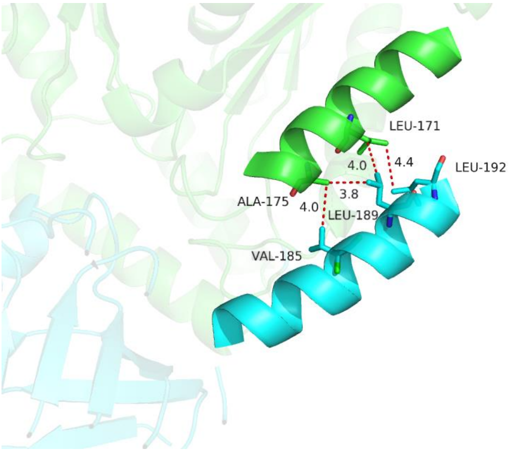


**Figure S1. Helix-helix interaction between α and β subunits stabilizes the C-terminal helix in β subunit of the protein complex**. Highlighted in blue is the C-terminal helix from β subunit, which initially disordered in the structure of β subunit, becomes ordered in the complex on interacting with helix (α4) from α subunit highlighted in green. The helix-helix interactions have been highlighted.

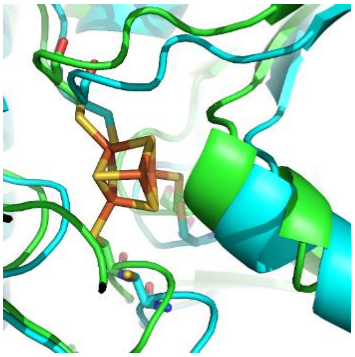

**Figure S2.** Overlay of 4Fe-4S cluster from the crystal structure of holo-LmFH onto that from apo-MjFH shows that the active site pocket with three cysteine residues are well poised to accommodate the cluster.

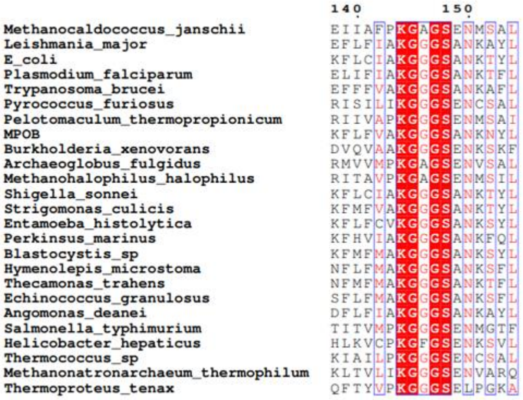


**Figure S3. Motif KGXGS is identical and conserved across all class-I FH.**

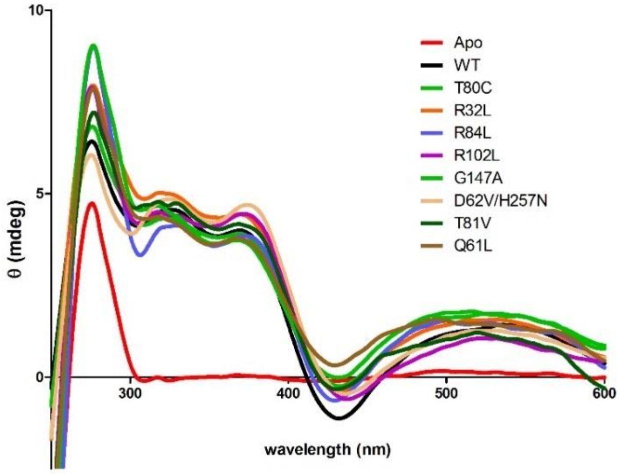


**Figure S4. Circular dichroism spectra of apo- and 4Fe-4S reconstituted wild type and mutants of MjFH.** The CD spectrum of apo wild type MjFH is also shown.

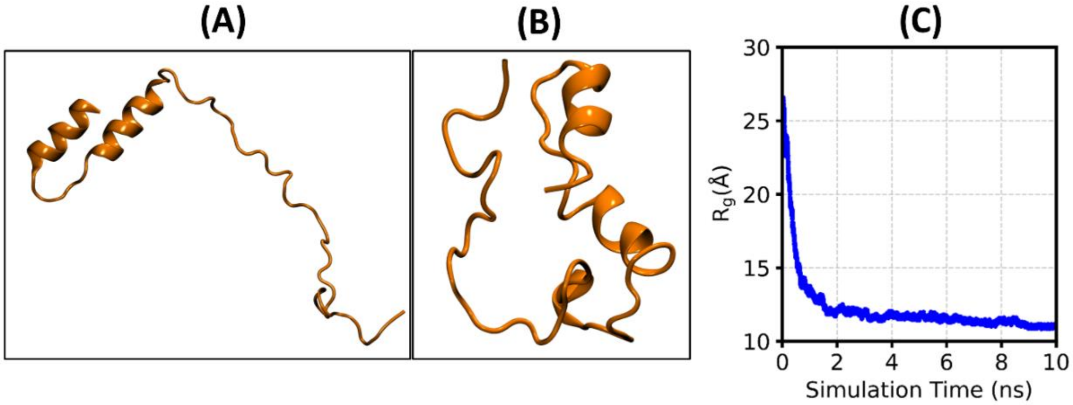


**Figure S5. Modelling and WTMetaD simulation of missing 63 residue peptide stretch from LmFH.** The structure of the 63 residues polypeptide **A**. after joining the two independently modelled fragments, **B.** with a compact globular architecture (Rg = 12 Å) from the WTMetaD simulation. **C.** The evolution of the Rg of the 63 residues modelled peptide in the WTMetaD simulation. Rg spans a wide range of values within the simulation time.

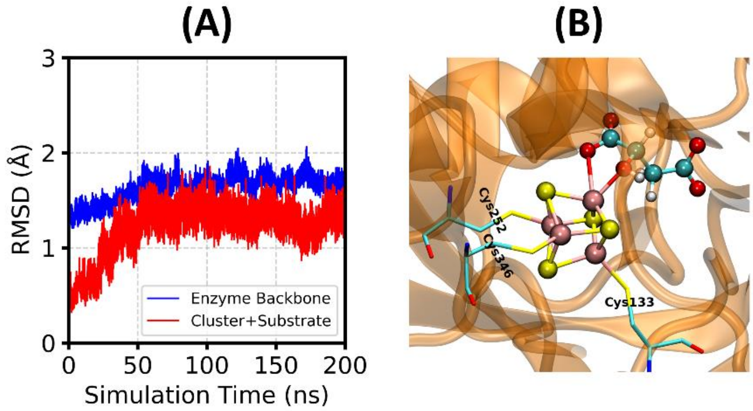


**Figure S6. Validation of force field parameters for 4Fe-4S cluster and malate. A**. RMSD of backbone atoms of holo-LmFH (blue, without the modelled 63-residues long fragment from the N-terminal chain) and the heavy atoms of the cluster and substrate (red) considering the energy minimized structure as a reference. The convergence of the RMSDs within 2 Å shows the stability of the enzyme and of the 4Fe-4S cluster (and substrate) during the simulation and hence validates the force field parameters employed. **B**. Snapshot of the cluster and malate bound to LmFH at 200 ns simulation time also showing that the structures of the cluster and substrate are intact and stable during the simulation.

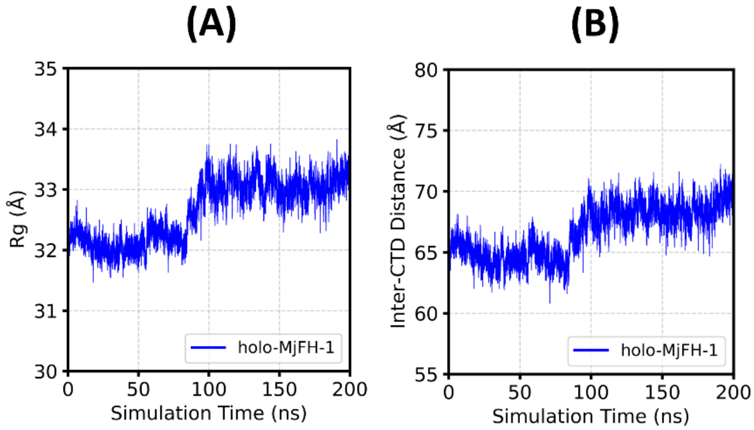


**Figure S7.** The **A**. Rg and **B**. inter-CTD distance values plotted against simulation time for holo-MjFH-1. Simulation of holo-MjFH just after docking of the cluster and substrate does not show a contraction.

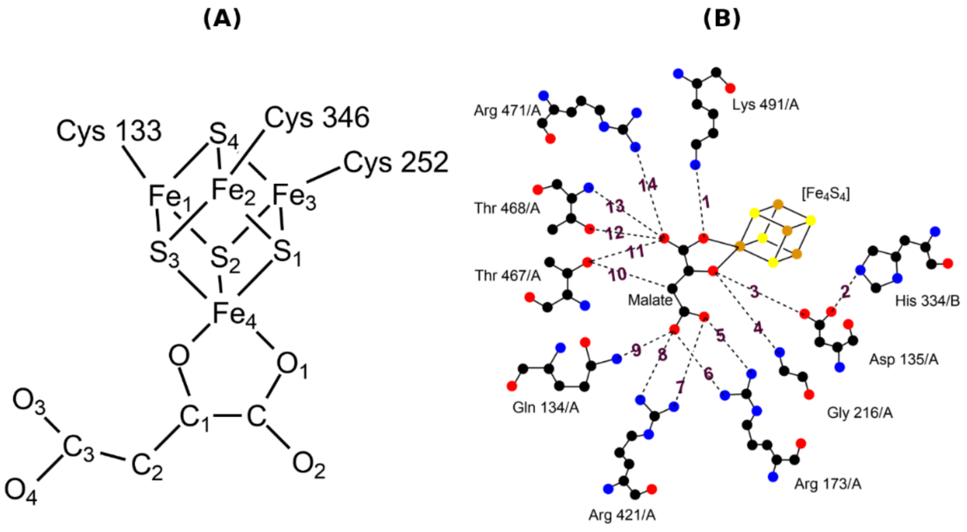


**Figure S8. Contacts of LmFH active site with the bound substrate, malate. A**. The 4Fe-4S cluster with malate and three cysteine residues (from chain A) bound to it, in holo-LmFH. **B**. The catalytic contacts of protein residues with malate present in the holo-LmFH, the corresponding distances in LmFH and MjFH are listed in **Table S5**. To show which chain a residue belongs, "/A" and "/B" are added at the end of the residue name. The contacts are numbered from 1 to 14 for better reference. Color code: Yellow - Sulfur, Orange - Iron, Black - Carbon, Blue - Nitrogen, and Red – Oxygen.

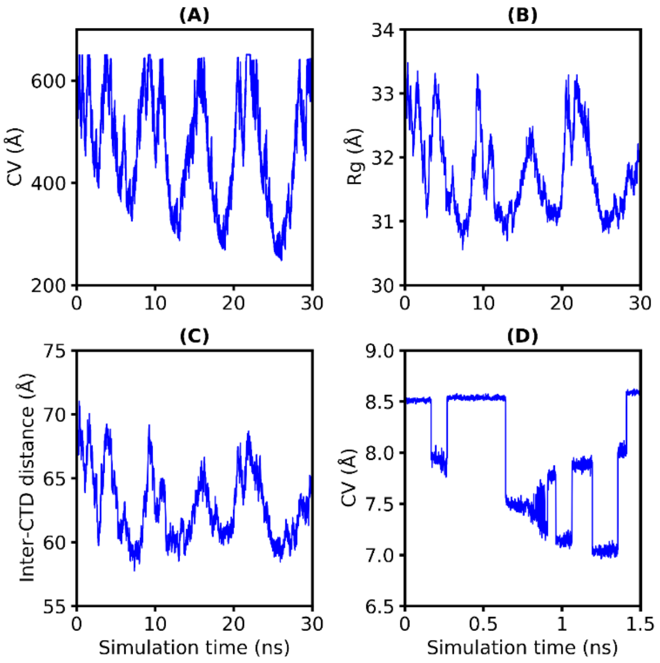


**Figure S9. CV, Rg and Inter-CTD distance evolution from metadynamics run of holo-MjFH. A**. CV (linear combination of all the 86 pair distances), **B**. Rg, and **C**. Inter-CTD distance from the first metadynamics run of holo-MjFH, **D**. The evolution of CV from the second metadynamics run, where the CV is linear combination of only the 28 catalytic contacts (14 catalytic contacts per chain as in **Figure S8. B**).

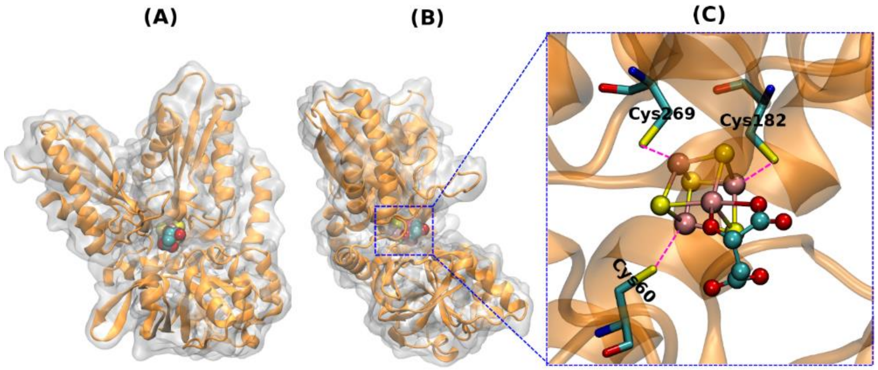


**Figure S10. Ligand docked in the cavity of MjFH.** Two views (**A** and **B**) of the ligand docked in the chain A of MjFH showing the ligand bound to the cavity. Enzyme is represented as orange cartoon as well as transparent white surface and the ligand is shown in ball representation. **C**. A zoomed-in view of the catalytic site showing close contact between three cysteine residues and three Fe from 4Fe-4S cluster with dashed magenta lines (refer to **Table S4** for the values).

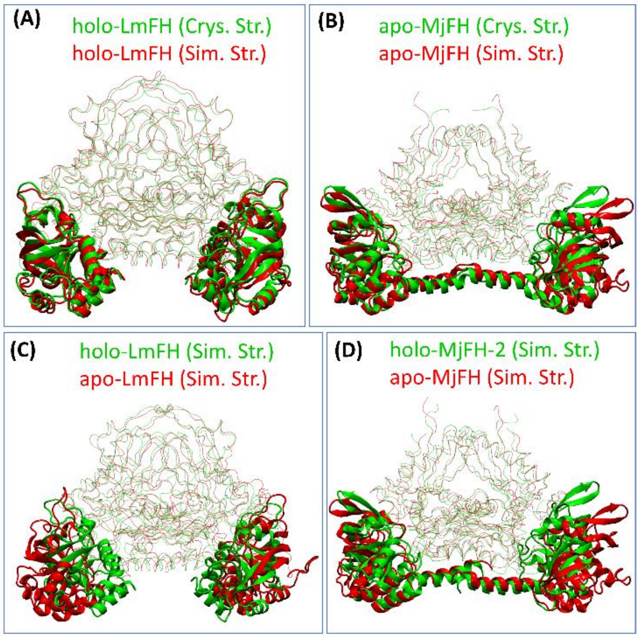


**Figure S11. Superposition of apo- and holo- Mj and LmFH structures to observe difference in structures post MD. A**. The alignment of holo-LmFH obtained from simulation on the crystal structure of the same showing that the holo-LmFH does not undergo any contraction/expansion during the simulation. CTDs/β-subunits are shown in cartoon representations, whereas the remaining parts are shown in transparent lines. **B**. The overlay of apo-MjFH from the simulation on the crystal structure of the same. The apo-MjFH expands marginally in the simulation as can be inferred from the CTD regions (refer to **Table S7** for the inter-CTD distance values). The alignment of **C**. apo-LmFH on holo-LmFH and **D**. apo-MjFH on holo-MjFH-2, all being simulation structures. Panels C and D demonstrate that the apo- structure of class-I FH is an expanded form of their holo- forms. The structures of all the enzyme forms are obtained as the structure representing the major cluster (obtained using cluster analysis) present in the last 100 ns of their respective simulation. The highly flexible 63-residues long modelled peptide fragment, present in the LmFH structures, is not shown.

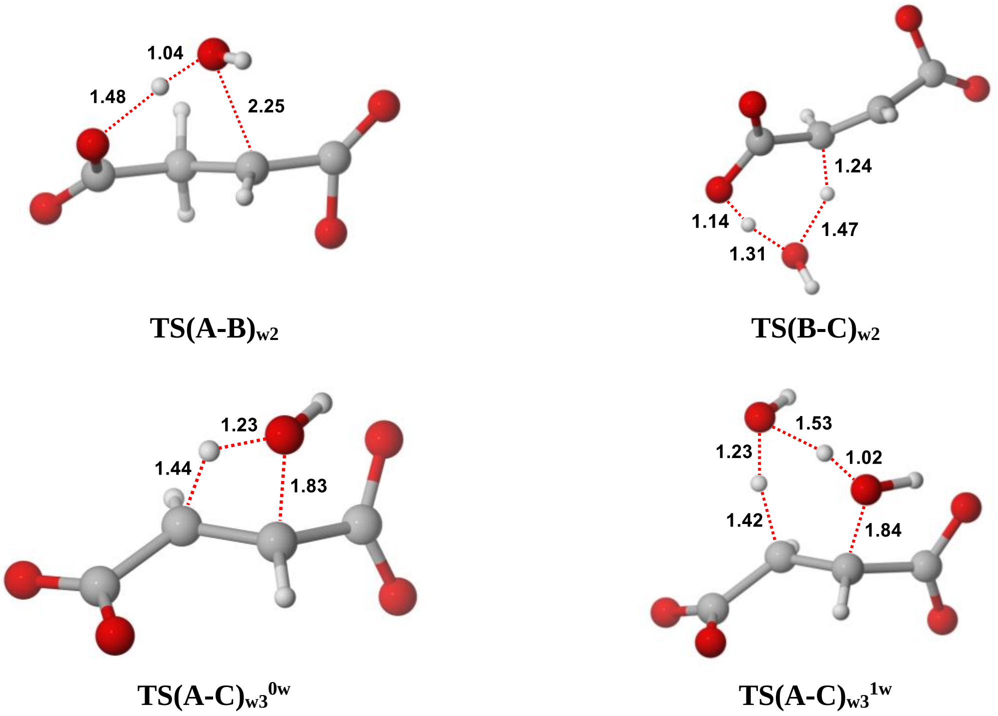


**Figure S12.** Optimized geometries of the transition states involved in carbocation pathway (w2) and concerted pathway (w3). Distances shown are in Å.

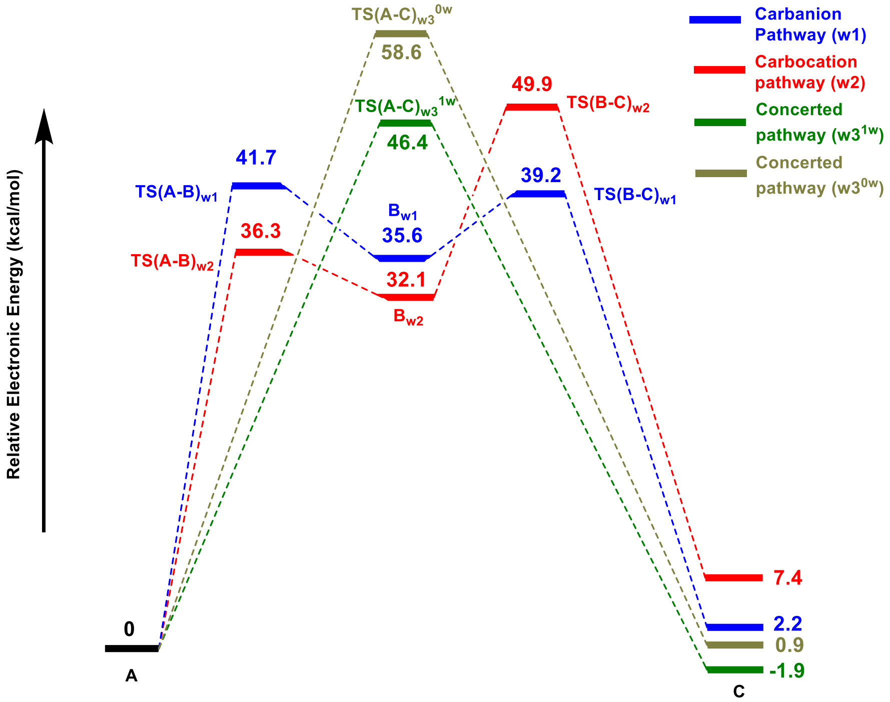


**Figure S13.** Electronic energy profiles for different plausible pathways in aqueous medium.

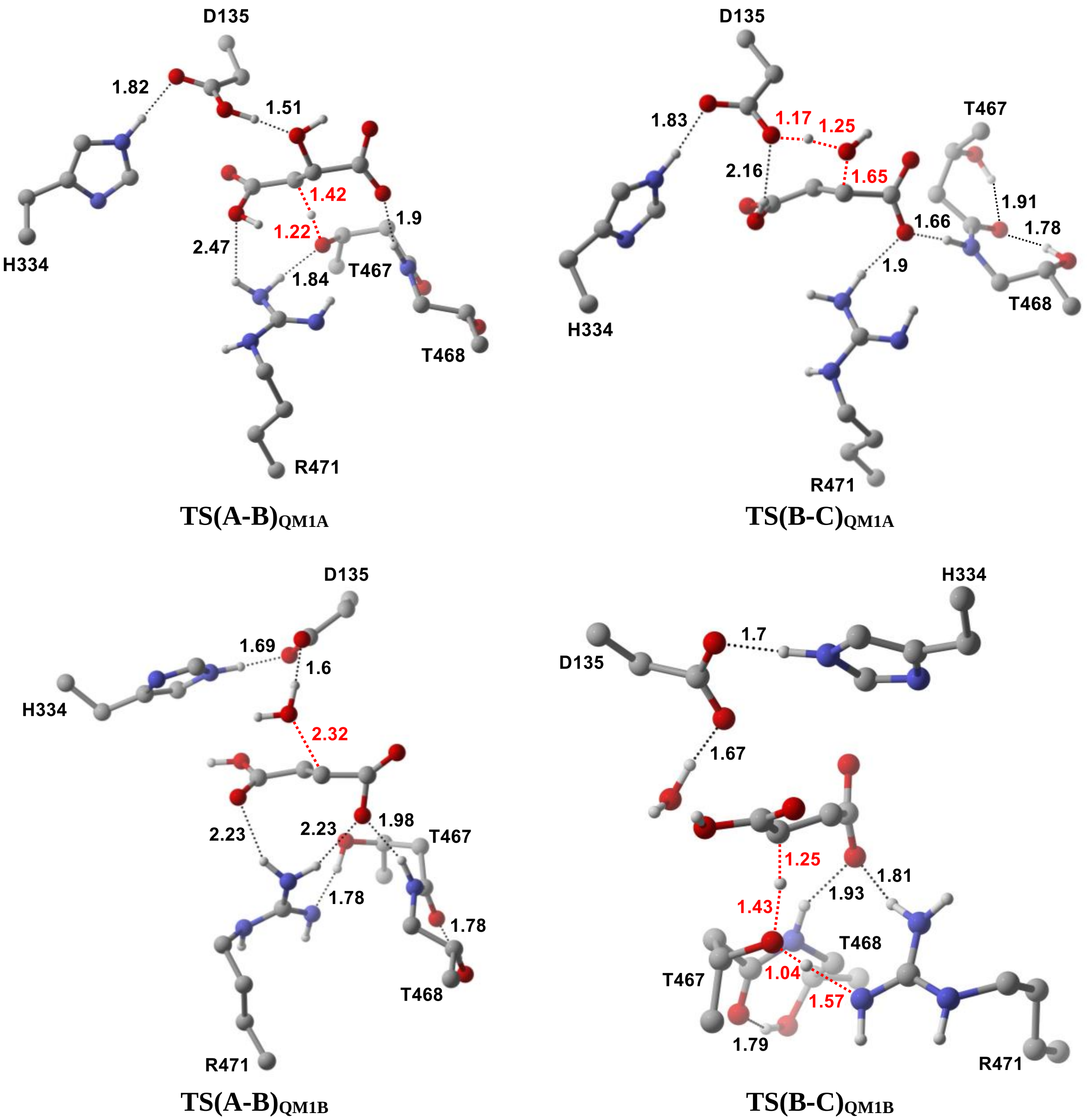


**Figure S14.** Optimized geometries of the transition states in carbanion pathway (QM1A) and carbocation pathway (QM1B) for QM1 cluster model. Distances are in Å.

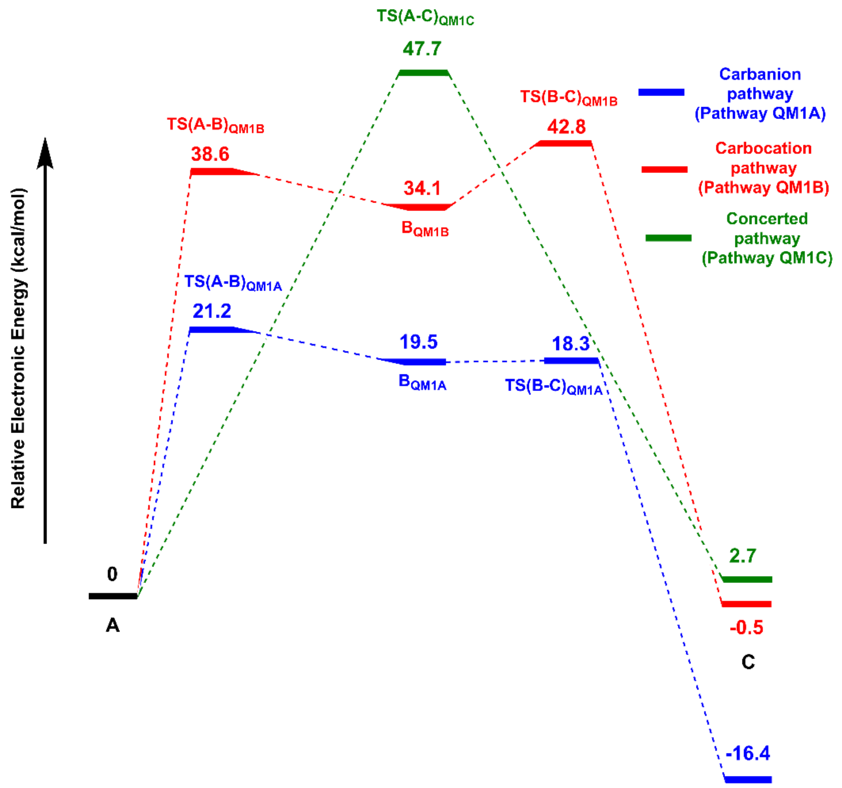


**Figure S15.** Electronic energy profiles of various plausible pathways obtained from QM1 model.

#### 9. CARTESIAN COORDINATES

---

**$A_{w1}$**

---

Number of imaginary frequencies: 0 Electronic energy: HF=-607.9706985
Zero-point correction= 0.111536 (Hartree/Particle)
Thermal correction to Energy= 0.122146
Thermal correction to Enthalpy= 0.123090
Thermal correction to Gibbs Free Energy= 0.073954
Sum of electronic and zero-point Energies= -607.8591625
Sum of electronic and thermal Energies= -607.8485525
Sum of electronic and thermal Enthalpies= -607.8476085
Sum of electronic and thermal Free Energies= -607.8967445

...........................................................

Cartesian Coordinates

...........................................................

| | | | |
|---|---|---|---|
| C | 1.901816 | 0.058561 | 0.174727 |
| O | 1.928796 | 1.247219 | 0.590987 |
| O | 2.870293 | -0.764256 | 0.139943 |
| C | 0.547578 | -0.497782 | -0.374026 |
| O | 0.794304 | -1.800267 | -0.914115 |
| C | -2.024116 | -0.398087 | 0.162427 |
| O | -2.242078 | 0.575608 | -0.628566 |
| O | -2.863562 | -1.234234 | 0.580546 |
| H | 1.741841 | -1.885860 | -0.646991 |
| H | 0.214444 | 0.181230 | -1.172618 |
| H | -1.077183 | 2.047343 | -0.441627 |
| C | -0.561813 | -0.508864 | 0.698167 |
| H | -0.398507 | 0.367753 | 1.339448 |
| H | -0.452821 | -1.411765 | 1.311346 |
| O | -0.434305 | 2.783905 | -0.314013 |
| H | 0.363857 | 2.314536 | 0.004415 |

---

**$B_{w1}$**

---

Number of imaginary frequencies: 0 Electronic energy: HF=-607.9123865
Zero-point correction= 0.109840 (Hartree/Particle)
Thermal correction to Energy= 0.120635
Thermal correction to Enthalpy= 0.121579
Thermal correction to Gibbs Free Energy= 0.072152
Sum of electronic and zero-point Energies= -607.8025465
Sum of electronic and thermal Energies= -607.7917515
Sum of electronic and thermal Enthalpies= -607.7908075
Sum of electronic and thermal Free Energies= -607.8402345

...........................................................

Cartesian Coordinates

...........................................................

| | | | |
|---|---|---|---|
| C | -2.201793 | -0.039433 | -0.260162 |
| O | -2.656156 | -0.154684 | -1.418936 |
| O | -2.803833 | -0.275896 | 0.843798 |
| C | -0.724870 | 0.464428 | -0.069657 |
| O | -0.592722 | 0.763966 | 1.351289 |
| C | 1.503403 | -0.909075 | 0.005962 |
| O | 1.907302 | -0.095019 | 1.134572 |
| O | 2.314679 | -1.806936 | -0.313444 |

| | | | |
|---|---|---|---|
| H | -1.424294 | 0.297333 | 1.650232 |
| H | -0.650364 | 1.444873 | -0.580949 |
| H | 1.057880 | 0.289859 | 1.455298 |
| C | 0.287901 | -0.549298 | -0.582191 |
| H | 1.676291 | 1.253986 | -1.274968 |
| H | -0.067960 | -1.196200 | -1.379374 |
| O | 2.306029 | 1.901919 | -0.896559 |
| H | 2.418210 | 1.443614 | -0.039697 |

---

**$C_{w1}$**

---

Number of imaginary frequencies: 0 Electronic energy: HF=-607.9654498
Zero-point correction= 0.108251 (Hartree/Particle)
Thermal correction to Energy= 0.120336
Thermal correction to Enthalpy= 0.121280
Thermal correction to Gibbs Free Energy= 0.068659
Sum of electronic and zero-point Energies= -607.8571988
Sum of electronic and thermal Energies= -607.8451138
Sum of electronic and thermal Enthalpies= -607.8441698
Sum of electronic and thermal Free Energies= -607.8967908

..........................................................

Cartesian Coordinates

..........................................................

| | | | |
|---|---|---|---|
| C | -1.540017 | -1.331837 | 0.029447 |
| O | -2.600304 | -0.623882 | -0.060050 |
| O | -1.471126 | -2.541144 | 0.359121 |
| C | -0.249317 | -0.573374 | -0.273622 |
| O | -2.409200 | 2.132226 | -0.596889 |
| C | 2.268227 | -0.171429 | -0.159544 |
| O | 2.183344 | 1.093894 | -0.324327 |
| O | 3.321995 | -0.855185 | -0.201808 |
| H | -2.497210 | 1.160025 | -0.426910 |
| H | -0.372555 | 0.381342 | -0.781159 |
| H | 0.946951 | 1.940914 | 0.413038 |
| H | -1.546825 | 2.341401 | -0.196649 |
| C | 0.982791 | -0.948607 | 0.102572 |
| H | 1.097815 | -1.921212 | 0.582064 |
| O | 0.205507 | 2.376567 | 0.940801 |
| H | -0.240000 | 1.589203 | 1.281720 |

---

**TS(A-B)$_{w1}$**

---

Number of imaginary frequencies: 1 Electronic energy: HF=-607.8976774
Zero-point correction= 0.105328 (Hartree/Particle)
Thermal correction to Energy= 0.115539
Thermal correction to Enthalpy= 0.116483
Thermal correction to Gibbs Free Energy= 0.068314
Sum of electronic and zero-point Energies= -607.7923494
Sum of electronic and thermal Energies= -607.7821384
Sum of electronic and thermal Enthalpies= -607.7811944
Sum of electronic and thermal Free Energies= -607.8293634

..........................................................

Cartesian Coordinates

..........................................................

| | | | |
|---|---|---|---|
| C | 2.087755 | 0.248359 | 0.119200 |
| O | 2.272387 | 1.449511 | 0.405941 |

| | | | |
|---|---|---|---|
| O | 2.935581 | -0.707218 | 0.203644 |
| C | 0.677512 | -0.209011 | -0.433397 |
| O | 0.830800 | -1.578681 | -0.867661 |
| C | -1.745300 | -0.588591 | 0.303818 |
| O | -2.336985 | -0.126160 | -0.874213 |
| O | -2.426559 | -1.341568 | 1.009223 |
| H | 1.745964 | -1.725940 | -0.517063 |
| H | 0.467433 | 0.403682 | -1.327455 |
| H | -1.940221 | 0.771379 | -1.003266 |
| C | -0.442140 | -0.000884 | 0.597145 |
| H | -0.818875 | 1.435590 | 0.304654 |
| H | -0.120112 | -0.324068 | 1.594019 |
| O | -1.356024 | 2.347790 | -0.237185 |
| H | -2.154752 | 2.390723 | 0.310524 |

---

**TS(B-C)$_{w1}$**

---

Number of imaginary frequencies: 1 Electronic energy: HF=-607.902868
Zero-point correction= 0.106215 (Hartree/Particle)
Thermal correction to Energy= 0.116716
Thermal correction to Enthalpy= 0.117661
Thermal correction to Gibbs Free Energy= 0.069390
Sum of electronic and zero-point Energies= -607.796653
Sum of electronic and thermal Energies= -607.786152
Sum of electronic and thermal Enthalpies= -607.785207
Sum of electronic and thermal Free Energies= -607.833478

..........................................................
Cartesian Coordinates
..........................................................

| | | | |
|---|---|---|---|
| C | 2.120706 | -0.416390 | 0.050988 |
| O | 2.752499 | -1.035291 | 0.946344 |
| O | 2.542700 | -0.025040 | -1.077903 |
| C | 0.633544 | -0.128027 | 0.355511 |
| O | 0.792585 | 1.827625 | 0.132518 |
| C | -1.705800 | -0.989325 | -0.078897 |
| O | -2.250958 | -0.163129 | 0.901746 |
| O | -2.446875 | -1.865576 | -0.555200 |
| H | 1.280728 | 1.649131 | -0.687791 |
| H | 0.402464 | 0.014733 | 1.407327 |
| H | -1.897389 | 0.748785 | 0.747432 |
| H | -0.526417 | 2.168953 | -0.062810 |
| C | -0.338396 | -0.754489 | -0.441710 |
| H | -0.010164 | -1.228738 | -1.363931 |
| O | -1.604825 | 2.335415 | -0.151370 |
| H | -1.790549 | 1.784481 | -0.924671 |

---

**A$_{w2}$**

---

Number of imaginary frequencies: 0 Electronic energy: HF=-531.9443357
Zero-point correction= 0.098166 (Hartree/Particle)
Thermal correction to Energy= 0.106285
Thermal correction to Enthalpy= 0.107229
Thermal correction to Gibbs Free Energy= 0.064331
Sum of electronic and zero-point Energies= -531.8461697
Sum of electronic and thermal Energies= -531.8380507

Sum of electronic and thermal Enthalpies= -531.8371067
Sum of electronic and thermal Free Energies= -531.8800047

............................................................

Cartesian Coordinates

............................................................

| | | | |
|---|---|---|---|
| C | -1.950236 | -0.287091 | -0.022677 |
| O | -2.339208 | -1.454338 | -0.180769 |
| O | -2.524032 | 0.711381 | 0.521745 |
| C | -0.505155 | 0.080930 | -0.514558 |
| O | -0.350981 | 1.488248 | -0.302293 |
| C | 0.548293 | -0.715400 | 0.278735 |
| C | 1.991839 | -0.256731 | 0.091124 |
| O | 2.158919 | 1.077474 | 0.062208 |
| O | 2.941572 | -1.013447 | 0.021010 |
| H | 0.473652 | -1.776662 | 0.039538 |
| H | 0.318600 | -0.603476 | 1.348432 |
| H | 1.246055 | 1.488206 | 0.035655 |
| H | -0.407476 | -0.143397 | -1.585859 |
| H | -1.229437 | 1.630536 | 0.191275 |

--------------------------------------------------------------------------------------------------------------

**$B_{w2}$**

--------------------------------------------------------------------------------------------------------------

Number of imaginary frequencies: 0 Electronic energy: HF=-531.8907448
Zero-point correction= 0.093815 (Hartree/Particle)
Thermal correction to Energy= 0.103927
Thermal correction to Enthalpy= 0.104871
Thermal correction to Gibbs Free Energy= 0.056869
Sum of electronic and zero-point Energies= -531.7969298
Sum of electronic and thermal Energies= -531.7868178
Sum of electronic and thermal Enthalpies= -531.7858738
Sum of electronic and thermal Free Energies= -531.8338758

............................................................

Cartesian Coordinates

............................................................

| | | | |
|---|---|---|---|
| C | 2.180369 | -0.036712 | 0.006874 |
| O | 1.534449 | -0.181817 | 1.154165 |
| O | 3.320099 | -0.121516 | -0.376764 |
| C | 0.761369 | 0.183008 | -0.192896 |
| O | -1.216543 | 2.747516 | -0.209398 |
| C | -0.178601 | -0.859730 | -0.705983 |
| C | -1.604963 | -0.859695 | -0.028306 |
| O | -2.093421 | 0.280940 | 0.233796 |
| O | -2.109280 | -1.980815 | 0.138036 |
| H | 0.261080 | -1.859135 | -0.646699 |
| H | -0.336676 | -0.630665 | -1.771144 |
| H | -1.594706 | 1.821382 | -0.164610 |
| H | 0.371878 | 1.200510 | -0.145118 |
| H | -1.133048 | 2.952222 | 0.730753 |

--------------------------------------------------------------------------------------------------------------

**$C_{w2}$**

--------------------------------------------------------------------------------------------------------------

Number of imaginary frequencies: 0 Electronic energy: HF=-531.9311027
Zero-point correction= 0.095204 (Hartree/Particle)
Thermal correction to Energy= 0.104841
Thermal correction to Enthalpy= 0.105785
Thermal correction to Gibbs Free Energy= 0.059839

Sum of electronic and zero-point Energies= -531.8358987
Sum of electronic and thermal Energies= -531.8262617
Sum of electronic and thermal Enthalpies= -531.8253177
Sum of electronic and thermal Free Energies= -531.8712637

............................................................

Cartesian Coordinates

............................................................

| | | | |
|---|---|---|---|
| O | 0.131951 | 2.359311 | 0.203928 |
| C | -1.993712 | -0.374308 | -0.002415 |
| O | -2.023736 | 0.866359 | -0.315426 |
| O | -2.948954 | -1.149762 | 0.178969 |
| C | -0.587250 | -0.942858 | 0.198246 |
| C | 0.443424 | -0.456204 | -0.500516 |
| C | 1.871727 | -0.550896 | -0.110801 |
| O | 2.527931 | -1.547974 | 0.117318 |
| H | -0.682449 | 1.799404 | -0.040957 |
| H | 0.216661 | 0.304137 | -1.242428 |
| H | -0.438378 | -1.688459 | 0.978626 |
| H | 0.059094 | 2.416750 | 1.165107 |
| O | 2.409488 | 0.692069 | 0.019776 |
| H | 1.666491 | 1.353742 | -0.003951 |

--------------------------------------------------------------------------------------------------------------

**TS(A-B)$_{w2}$**

--------------------------------------------------------------------------------------------------------------

Number of imaginary frequencies: 1 Electronic energy: HF=-531.8819296
Zero-point correction= 0.092783 (Hartree/Particle)
Thermal correction to Energy= 0.101782
Thermal correction to Enthalpy= 0.102726
Thermal correction to Gibbs Free Energy= 0.058035
Sum of electronic and zero-point Energies= -531.7891466
Sum of electronic and thermal Energies= -531.7801476
Sum of electronic and thermal Enthalpies= -531.7792036
Sum of electronic and thermal Free Energies= -531.8238946

............................................................

Cartesian Coordinates

............................................................

| | | | |
|---|---|---|---|
| C | 2.035174 | -0.265652 | 0.028305 |
| O | 1.912995 | -1.410760 | 0.613923 |
| O | 2.947226 | 0.336253 | -0.505309 |
| C | 0.605218 | -0.094622 | 0.320629 |
| O | 0.025861 | 2.022679 | -0.176971 |
| C | -0.502400 | -0.648647 | -0.531058 |
| C | -1.950374 | -0.285347 | -0.047544 |
| O | -2.080159 | 0.861719 | 0.511350 |
| O | -2.831740 | -1.125251 | -0.265200 |
| H | -0.413875 | -1.737690 | -0.544911 |
| H | -0.365925 | -0.298902 | -1.561882 |
| H | -0.906854 | 1.682752 | 0.136423 |
| H | 0.353798 | 0.171298 | 1.341184 |
| H | 0.413687 | 2.471029 | 0.584842 |

--------------------------------------------------------------------------------------------------------------

**TS(B-C)$_{w2}$**

--------------------------------------------------------------------------------------------------------------

Number of imaginary frequencies: 1 Electronic energy: HF=-531.854837
Zero-point correction= 0.087295 (Hartree/Particle)
Thermal correction to Energy= 0.096451

Thermal correction to Enthalpy= 0.097396
Thermal correction to Gibbs Free Energy= 0.051837
Sum of electronic and zero-point Energies= -531.767542
Sum of electronic and thermal Energies= -531.758386
Sum of electronic and thermal Enthalpies= -531.757441
Sum of electronic and thermal Free Energies= -531.80300

............................................................

Cartesian Coordinates

............................................................

| | | | |
|---|---|---|---|
| O | -1.932364 | 2.099886 | -0.033067 |
| C | 2.294371 | 0.153080 | -0.040821 |
| O | 3.007424 | 1.144306 | -0.042728 |
| O | 2.440323 | -1.062815 | 0.339064 |
| C | 0.909282 | -0.151898 | -0.459325 |
| C | -0.274681 | 0.115280 | 0.277579 |
| C | -1.492249 | -0.804244 | 0.058580 |
| O | -1.371549 | -1.971946 | -0.292880 |
| H | -0.760449 | 1.213863 | -0.051021 |
| H | -0.102540 | 0.354927 | 1.332117 |
| H | 0.801238 | -0.675517 | -1.407477 |
| H | -2.111772 | 2.252825 | -0.972510 |
| O | -2.643074 | -0.201718 | 0.265878 |
| H | -2.452897 | 0.918882 | 0.192681 |

---

**$A_{w3}^{1w}$**

---

Number of imaginary frequencies: 0 Electronic energy: HF=-607.9617119
Zero-point correction= 0.110772 (Hartree/Particle)
Thermal correction to Energy= 0.121735
Thermal correction to Enthalpy= 0.122680
Thermal correction to Gibbs Free Energy= 0.072360
Sum of electronic and zero-point Energies= -607.8509399
Sum of electronic and thermal Energies= -607.8399769
Sum of electronic and thermal Enthalpies= -607.8390319
Sum of electronic and thermal Free Energies= -607.8893519

............................................................

Cartesian Coordinates

............................................................

| | | | |
|---|---|---|---|
| C | -0.922696 | -0.048523 | -0.600406 |
| C | 0.111312 | -0.406634 | 0.466192 |
| H | -0.929623 | -0.881616 | -1.317530 |
| C | 1.406047 | -1.086992 | -0.094512 |
| O | 1.293370 | -2.113199 | -0.792071 |
| O | 2.505288 | -0.519700 | 0.269970 |
| C | -2.405670 | 0.141888 | -0.107168 |
| O | -3.102492 | 0.941100 | -0.792309 |
| O | -2.767467 | -0.563476 | 0.873808 |
| O | 0.514678 | 0.750587 | 1.229369 |
| H | -0.385436 | -1.125485 | 1.137265 |
| H | -0.601116 | 0.846127 | -1.149490 |
| H | 1.547888 | 2.149732 | -0.088154 |
| O | 2.441208 | 2.202929 | -0.469442 |
| H | 2.673627 | 1.251477 | -0.385041 |
| H | 1.484011 | 0.575398 | 1.263726 |

---

**$C_{w3}^{1w}$**

--------------------------------------------------------------------------------------------------------------
Number of imaginary frequencies:  0 Electronic energy:          HF=-607.9633216
Zero-point correction=                           0.108001 (Hartree/Particle)
Thermal correction to Energy=                    0.120330
Thermal correction to Enthalpy=                  0.121274
Thermal correction to Gibbs Free Energy=         0.067534
Sum of electronic and zero-point Energies=           -607.8553206
Sum of electronic and thermal Energies=              -607.8429916
Sum of electronic and thermal Enthalpies=            -607.8420476
Sum of electronic and thermal Free Energies=         -607.8957876
............................................................
Cartesian Coordinates
............................................................

| | | | |
|---|---|---|---|
| C | -1.020403 | -1.046272 | 0.064677 |
| C | 0.224492 | -0.584347 | -0.113717 |
| H | -1.153940 | -2.111254 | 0.254580 |
| C | 1.505611 | -1.406813 | -0.032851 |
| O | 1.423690 | -2.655196 | 0.053426 |
| O | 2.589662 | -0.722650 | -0.058159 |
| C | -2.303217 | -0.220793 | -0.002372 |
| O | -3.373901 | -0.881203 | 0.052176 |
| O | -2.211544 | 1.046990 | -0.112596 |
| O | 2.614290 | 1.972053 | 0.086601 |
| H | 0.348078 | 0.476722 | -0.309858 |
| H | 0.426556 | 2.512611 | 0.613534 |
| H | 1.886471 | 2.236996 | -0.492396 |
| O | -0.241854 | 2.900750 | 0.032023 |
| H | -0.934253 | 2.180546 | -0.032501 |
| H | 2.585446 | 0.967773 | 0.044458 |

--------------------------------------------------------------------------------------------------------------

**$TS(A\text{-}C)_{w3}^{1w}$**

--------------------------------------------------------------------------------------------------------------
Number of imaginary frequencies:  1 Electronic energy:          HF=-607.8807641
Zero-point correction=                           0.104584 (Hartree/Particle)
Thermal correction to Energy=                    0.114799
Thermal correction to Enthalpy=                  0.115743
Thermal correction to Gibbs Free Energy=         0.067709
Sum of electronic and zero-point Energies=           -607.7761801
Sum of electronic and thermal Energies=              -607.7659651
Sum of electronic and thermal Enthalpies=            -607.7650211
Sum of electronic and thermal Free Energies=         -607.8130551
............................................................
Cartesian Coordinates
............................................................

| | | | |
|---|---|---|---|
| C | -0.672116 | -0.063777 | -0.456538 |
| C | 0.433652 | -0.285324 | 0.421342 |
| H | -0.483027 | -0.350013 | -1.495893 |
| C | 1.765169 | -0.856454 | -0.125411 |
| O | 1.780127 | -2.057148 | -0.415430 |
| O | 2.733882 | -0.023743 | -0.226931 |
| C | -2.070824 | -0.470433 | 0.053421 |
| O | -2.942023 | -0.648018 | -0.836329 |
| O | -2.211473 | -0.567261 | 1.301205 |
| O | 1.148599 | 1.296865 | 1.020315 |
| H | 0.135543 | -0.631380 | 1.410681 |
| H | -0.677263 | 1.353978 | -0.566945 |

| | | | |
|---|---|---|---|
| H | 0.616991 | 1.945117 | 0.437807 |
| O | -0.327649 | 2.530566 | -0.621692 |
| H | 0.219513 | 2.465299 | -1.415548 |
| H | 2.001249 | 1.022846 | 0.503918 |

---

**$A_{w3}^{0w}$**

---

Number of imaginary frequencies: 0 Electronic energy: HF=-531.483962
Zero-point correction= 0.084910 (Hartree/Particle)
Thermal correction to Energy= 0.093282
Thermal correction to Enthalpy= 0.094226
Thermal correction to Gibbs Free Energy= 0.050235
Sum of electronic and zero-point Energies= -531.399052
Sum of electronic and thermal Energies= -531.390680
Sum of electronic and thermal Enthalpies= -531.389736
Sum of electronic and thermal Free Energies= -531.433727

Cartesian Coordinates

| | | | |
|---|---|---|---|
| C | -0.633228 | -0.342854 | -0.525582 |
| C | 0.490521 | 0.260968 | 0.325828 |
| H | -0.451458 | -1.427612 | -0.556665 |
| C | 1.903588 | -0.356109 | 0.013810 |
| O | 2.060224 | -1.593764 | 0.133638 |
| O | 2.783582 | 0.506058 | -0.329034 |
| C | -2.108995 | -0.140171 | -0.021950 |
| O | -2.984675 | -0.040385 | -0.929313 |
| O | -2.289970 | -0.162720 | 1.226809 |
| O | 0.594633 | 1.681775 | 0.161954 |
| H | 0.240644 | 0.027668 | 1.373879 |
| H | -0.552398 | 0.028265 | -1.556617 |
| H | 1.541541 | 1.712956 | -0.125657 |

---

**$C_{w3}^{0w}$**

---

Number of imaginary frequencies: 0 Electronic energy: HF=-531.4808339
Zero-point correction= 0.081542 (Hartree/Particle)
Thermal correction to Energy= 0.091628
Thermal correction to Enthalpy= 0.092572
Thermal correction to Gibbs Free Energy= 0.043852
Sum of electronic and zero-point Energies= -531.3992919
Sum of electronic and thermal Energies= -531.3892059
Sum of electronic and thermal Enthalpies= -531.3882619
Sum of electronic and thermal Free Energies= -531.4369819

Cartesian Coordinates

| | | | |
|---|---|---|---|
| C | -0.682635 | 0.016411 | -0.267770 |
| C | 0.250428 | -0.855599 | 0.145754 |
| H | -0.365085 | 0.919158 | -0.792363 |
| C | 1.757375 | -0.779133 | -0.084171 |
| O | 2.375177 | -1.871734 | 0.021532 |
| O | 2.282251 | 0.347784 | -0.378868 |
| C | -2.200771 | -0.126153 | -0.054028 |
| O | -2.882815 | 0.866594 | -0.447421 |
| O | -2.618158 | -1.188015 | 0.486949 |

| | | | |
|---|---|---|---|
| O | 1.297483 | 2.723123 | 0.439927 |
| H | -0.102971 | -1.761566 | 0.639336 |
| H | 0.393015 | 2.428326 | 0.608852 |
| H | 1.697164 | 1.858901 | 0.128506 |

---

**TS(A-C)$_{w3}$$^{0w}$**

---

Number of imaginary frequencies: 1 Electronic energy: HF=-531.3848812
Zero-point correction= 0.079221 (Hartree/Particle)
Thermal correction to Energy= 0.087628
Thermal correction to Enthalpy= 0.088572
Thermal correction to Gibbs Free Energy= 0.044911
Sum of electronic and zero-point Energies= -531.3056602
Sum of electronic and thermal Energies= -531.2972532
Sum of electronic and thermal Enthalpies= -531.2963092
Sum of electronic and thermal Free Energies= -531.3399702

........................................................

Cartesian Coordinates

........................................................

| | | | |
|---|---|---|---|
| C | -0.642185 | 0.027796 | -0.543909 |
| C | 0.443272 | 0.039513 | 0.392375 |
| H | -0.413724 | -0.315101 | -1.553758 |
| C | 1.850396 | -0.460837 | 0.002993 |
| O | 2.150206 | -1.588471 | 0.439357 |
| O | 2.537451 | 0.305351 | -0.733770 |
| C | -2.078345 | -0.177022 | -0.062466 |
| O | -2.926668 | -0.436161 | -0.959326 |
| O | -2.290996 | -0.041593 | 1.173585 |
| O | 0.719055 | 1.849686 | 0.329685 |
| H | 0.149265 | -0.057077 | 1.435406 |
| H | -0.184084 | 1.379123 | -0.355098 |
| H | 1.497333 | 1.705858 | -0.256760 |

---

**A$_{QM1A}$**

---

Number of imaginary frequencies: 2 Electronic energy: HF=-2024.133643
Zero-point correction= 0.725938 (Hartree/Particle)
Thermal correction to Energy= 0.773279
Thermal correction to Enthalpy= 0.774223
Thermal correction to Gibbs Free Energy= 0.635688
Sum of electronic and zero-point Energies= -2023.407705
Sum of electronic and thermal Energies= -2023.360364
Sum of electronic and thermal Enthalpies= -2023.35942
Sum of electronic and thermal Free Energies= -2023.497955

........................................................

Cartesian Coordinates

........................................................

| | | | |
|---|---|---|---|
| C | -5.265552 | -4.552785 | 0.699076 |
| C | -4.390729 | -3.475452 | 0.047650 |
| C | -5.181045 | -2.523871 | -0.831307 |
| O | -6.392843 | -2.361033 | -0.689123 |
| O | -4.522168 | -1.832326 | -1.744119 |
| C | 2.909720 | -2.709402 | 1.240852 |
| C | 4.151497 | -2.038799 | 0.640828 |
| O | 5.238759 | -2.005183 | 1.258606 |
| C | 2.537199 | -2.331920 | 2.693441 |

| | | | |
|---|---|---|---|
| C | 3.643691 | -2.599621 | 3.727226 |
| N | 4.003968 | -1.564273 | -0.609987 |
| C | 4.973044 | -0.675199 | -1.247363 |
| C | 6.307773 | -1.323953 | -1.660358 |
| O | 7.172661 | -1.524247 | -0.557143 |
| C | 7.053252 | -0.442594 | -2.663806 |
| C | 6.611878 | 5.065979 | -0.449721 |
| C | 5.211642 | 4.918497 | 0.174562 |
| C | 4.172878 | 4.259763 | -0.749834 |
| C | 2.740582 | 4.170903 | -0.170160 |
| N | 2.581523 | 3.282086 | 0.982211 |
| C | 2.466795 | 1.885640 | 0.791393 |
| N | 3.252361 | 1.029634 | 1.373419 |
| N | 1.475033 | 1.499149 | -0.063876 |
| C | -7.063601 | 4.982176 | -0.295249 |
| C | -8.011272 | 3.896650 | 0.239405 |
| C | -7.332709 | 2.559421 | 0.266292 |
| N | -6.140902 | 2.403042 | 0.941921 |
| C | -7.669340 | 1.389326 | -0.374608 |
| C | -5.753992 | 1.157298 | 0.706477 |
| N | -6.655551 | 0.503452 | -0.081346 |
| C | 0.438174 | -1.793266 | -1.737421 |
| O | 1.461984 | -1.135429 | -1.368384 |
| O | 0.406570 | -2.833121 | -2.438948 |
| C | -0.924131 | -1.240878 | -1.268265 |
| O | -1.929291 | -2.056507 | -1.901018 |
| C | -2.146347 | -0.173890 | 0.683293 |
| O | -1.817702 | 1.025794 | 0.495623 |
| O | -3.242450 | -0.623011 | 1.138074 |
| H | -1.358049 | -2.729753 | -2.350596 |
| H | 3.255298 | -2.372007 | 4.726238 |
| H | 3.966803 | -3.648824 | 3.708646 |
| H | 4.518552 | -1.979333 | 3.527340 |
| H | 8.036600 | -0.874258 | -2.876398 |
| H | 6.500650 | -0.347544 | -3.605285 |
| H | 7.210185 | 0.558899 | -2.246079 |
| H | 6.586693 | -1.728123 | 0.212474 |
| H | 7.023568 | 4.089470 | -0.727057 |
| H | 6.576248 | 5.679753 | -1.357611 |
| H | 7.311397 | 5.538773 | 0.248504 |
| H | 4.845143 | 5.910930 | 0.475594 |
| H | 5.300387 | 4.334241 | 1.100500 |
| H | 4.506764 | 3.249939 | -1.021372 |
| H | 4.121298 | 4.832683 | -1.687552 |
| H | 2.061797 | 3.804089 | -0.942634 |
| H | 2.400377 | 5.176894 | 0.114022 |
| H | -4.678084 | -5.156032 | 1.399732 |
| H | -6.093516 | -4.090442 | 1.241711 |
| H | -5.698565 | -5.226001 | -0.050040 |
| H | -6.141897 | 4.977076 | 0.291270 |
| H | -6.791395 | 4.779020 | -1.336638 |
| H | -7.514745 | 5.982665 | -0.245592 |
| H | -8.333659 | 4.179897 | 1.252903 |
| H | -8.920873 | 3.844386 | -0.374355 |
| H | -1.018726 | -0.208301 | -1.628341 |
| H | 5.194638 | 0.169784 | -0.580002 |

| | | | |
|---|---|---|---|
| H | 3.041200 | -1.538090 | -1.004398 |
| H | 4.469488 | -0.280369 | -2.135334 |
| H | 2.045803 | -2.515192 | 0.602886 |
| H | 3.107967 | -3.790448 | 1.213695 |
| H | -6.581633 | -0.461902 | -0.407244 |
| H | -4.826022 | 0.681873 | 1.035515 |
| H | -8.506783 | 1.117924 | -1.001089 |
| H | 3.223988 | 3.490905 | 1.736948 |
| H | 4.049538 | 1.498074 | 1.804292 |
| H | 1.491963 | 0.513084 | -0.349104 |
| H | 0.521911 | 1.816829 | 0.118336 |
| H | -3.522657 | -1.947309 | -1.710914 |
| H | 1.693427 | -2.993451 | 2.944919 |
| H | 6.067124 | -2.285934 | -2.150873 |
| H | -3.915671 | -2.810103 | 0.788387 |
| H | -3.568539 | -3.910992 | -0.530071 |
| C | -1.083096 | -1.222799 | 0.257940 |
| H | -0.130824 | -0.937982 | 0.715018 |
| H | -1.342397 | -2.232087 | 0.596704 |
| O | 2.033632 | -1.021691 | 2.816936 |
| H | 2.549107 | -0.387653 | 2.260467 |

---

**$B_{QM1A}$**

---

Number of imaginary frequencies: 1 Electronic energy: HF=-2024.102967
Zero-point correction= 0.724912 (Hartree/Particle)
Thermal correction to Energy= 0.773759
Thermal correction to Enthalpy= 0.774703
Thermal correction to Gibbs Free Energy= 0.629924
Sum of electronic and zero-point Energies= -2023.378055
Sum of electronic and thermal Energies= -2023.329208
Sum of electronic and thermal Enthalpies= -2023.328264
Sum of electronic and thermal Free Energies= -2023.473043

............................................................

Cartesian Coordinates

............................................................

| | | | |
|---|---|---|---|
| C | -4.104159 | -5.275833 | 0.621776 |
| C | -3.322479 | -4.064568 | 0.097088 |
| C | -4.126175 | -3.243348 | -0.899362 |
| O | -5.353111 | -3.168485 | -0.815807 |
| O | -3.483955 | -2.584830 | -1.844359 |
| C | 3.581907 | -2.070365 | 1.689545 |
| C | 4.729367 | -1.297851 | 1.059643 |
| O | 5.897666 | -1.426685 | 1.520448 |
| C | 3.893762 | -3.581159 | 1.762638 |
| C | 4.100161 | -4.195178 | 0.373742 |
| N | 4.426549 | -0.512276 | 0.024619 |
| C | 5.372698 | 0.412241 | -0.595343 |
| C | 6.516863 | -0.243090 | -1.394606 |
| O | 7.545125 | -0.730799 | -0.547336 |
| C | 7.159944 | 0.760683 | -2.350837 |
| C | 3.370707 | 7.218635 | 1.451935 |
| C | 2.058835 | 6.510341 | 1.087780 |
| C | 2.053524 | 5.934249 | -0.333410 |
| C | 0.715198 | 5.303956 | -0.767542 |
| N | 0.306869 | 4.101429 | -0.067319 |

| | | | |
|---|---|---|---|
| C | 0.926045 | 2.876795 | -0.348918 |
| N | 0.493297 | 1.852849 | 0.475247 |
| N | 1.819419 | 2.809151 | -1.279727 |
| C | -7.502873 | 3.839343 | -0.081573 |
| C | -8.258799 | 2.545756 | 0.266668 |
| C | -7.338520 | 1.362939 | 0.345486 |
| N | -6.292738 | 1.343076 | 1.246186 |
| C | -7.316239 | 0.224294 | -0.427137 |
| C | -5.643977 | 0.211305 | 1.013233 |
| N | -6.232286 | -0.502696 | 0.013238 |
| C | 1.128175 | -1.484850 | -1.104547 |
| O | 1.823460 | -0.484513 | -0.713372 |
| O | 1.559588 | -2.610004 | -1.450248 |
| C | -0.406729 | -1.298221 | -1.100032 |
| O | -0.921649 | -2.291952 | -2.090433 |
| C | -2.072698 | -0.776444 | 0.791288 |
| O | -2.461859 | 0.450307 | 0.185383 |
| O | -2.755788 | -1.087218 | 1.793767 |
| H | -0.182270 | -2.938771 | -2.083578 |
| H | 4.210308 | -5.281944 | 0.457801 |
| H | 3.262044 | -3.955635 | -0.289886 |
| H | 5.017534 | -3.802023 | -0.079759 |
| H | 8.023153 | 0.300659 | -2.842082 |
| H | 6.452931 | 1.091482 | -3.119018 |
| H | 7.515583 | 1.638477 | -1.798957 |
| H | 7.099833 | -1.057667 | 0.266505 |
| H | 4.220661 | 6.530851 | 1.378993 |
| H | 3.568356 | 8.057384 | 0.772983 |
| H | 3.348501 | 7.614403 | 2.474099 |
| H | 1.224015 | 7.219161 | 1.198603 |
| H | 1.871821 | 5.703377 | 1.807935 |
| H | 2.829949 | 5.169402 | -0.442093 |
| H | 2.294875 | 6.740280 | -1.044322 |
| H | 0.791817 | 5.035560 | -1.824352 |
| H | -0.090975 | 6.045436 | -0.664156 |
| H | -3.545023 | -5.781313 | 1.416336 |
| H | -5.069583 | -4.954511 | 1.019398 |
| H | -4.298694 | -6.005513 | -0.173337 |
| H | -6.695346 | 3.995006 | 0.638367 |
| H | -7.048354 | 3.767084 | -1.075410 |
| H | -8.164683 | 4.715613 | -0.067558 |
| H | -8.769385 | 2.687069 | 1.230847 |
| H | -9.044028 | 2.354784 | -0.477196 |
| H | -0.615540 | -0.315909 | -1.542825 |
| H | 5.830243 | 1.048131 | 0.175013 |
| H | 3.436190 | -0.534909 | -0.330035 |
| H | 4.778685 | 1.056701 | -1.249996 |
| H | 3.467828 | -1.697171 | 2.714513 |
| H | 2.650238 | -1.887031 | 1.152359 |
| H | -5.914319 | -1.401031 | -0.357177 |
| H | -4.728985 | -0.127659 | 1.494431 |
| H | -7.949611 | -0.120736 | -1.231287 |
| H | 0.026278 | 4.207298 | 0.897411 |
| H | -0.503324 | 1.814921 | 0.669489 |
| H | 2.035830 | 1.822048 | -1.434822 |
| H | -1.987003 | 0.535172 | -0.652713 |

| | | | |
|---|---|---|---|
| H | -2.468229 | -2.601605 | -1.800206 |
| H | 3.030245 | -4.067186 | 2.235341 |
| H | 6.075673 | -1.063925 | -1.987961 |
| H | -3.086929 | -3.363930 | 0.912845 |
| H | -2.364061 | -4.356395 | -0.342005 |
| C | -0.990488 | -1.467838 | 0.246797 |
| H | 5.645465 | -3.127055 | 2.399982 |
| H | -0.695776 | -2.367985 | 0.779733 |
| O | 5.007257 | -3.831549 | 2.623094 |
| H | 0.868120 | 0.933926 | 0.217083 |

----------------------------------------------------------------------------------------------------------

**C<sub>QM1A</sub>**

----------------------------------------------------------------------------------------------------------

Number of imaginary frequencies: 2 Electronic energy: HF=-2024.159269
Zero-point correction= 0.723627 (Hartree/Particle)
Thermal correction to Energy= 0.772711
Thermal correction to Enthalpy= 0.773656
Thermal correction to Gibbs Free Energy= 0.626487
Sum of electronic and zero-point Energies= -2023.435642
Sum of electronic and thermal Energies= -2023.386558
Sum of electronic and thermal Enthalpies= -2023.385613
Sum of electronic and thermal Free Energies= -2023.532782

............................................................

Cartesian Coordinates

............................................................

| | | | |
|---|---|---|---|
| C | -4.445612 | -5.484139 | 0.745016 |
| C | -3.363242 | -4.526599 | 0.250402 |
| C | -3.905231 | -3.161435 | -0.205334 |
| O | -4.979334 | -2.755314 | 0.293986 |
| O | -3.172284 | -2.536463 | -1.044660 |
| C | 3.415043 | -2.722740 | 1.782827 |
| C | 4.559518 | -1.931809 | 1.168473 |
| O | 5.752809 | -2.208245 | 1.473863 |
| C | 3.686229 | -4.241783 | 1.756230 |
| C | 3.799607 | -4.773955 | 0.323312 |
| N | 4.227104 | -0.957723 | 0.320701 |
| C | 5.188350 | -0.022677 | -0.255280 |
| C | 6.169297 | -0.624504 | -1.281845 |
| O | 7.236907 | -1.316671 | -0.651921 |
| C | 6.791214 | 0.468363 | -2.149455 |
| C | 3.586883 | 6.516556 | 2.778179 |
| C | 2.547697 | 6.368463 | 1.661056 |
| C | 3.175246 | 6.150937 | 0.279495 |
| C | 2.157434 | 5.976867 | -0.863918 |
| N | 1.277951 | 4.824020 | -0.756482 |
| C | 1.800295 | 3.534960 | -0.925849 |
| N | 0.914151 | 2.549287 | -0.568809 |
| N | 3.013465 | 3.392291 | -1.354359 |
| C | -7.485556 | 3.757292 | 1.438825 |
| C | -7.694426 | 3.007428 | 0.112309 |
| C | -7.412877 | 1.538589 | 0.230345 |
| N | -8.300709 | 0.680273 | 0.854918 |
| C | -6.282594 | 0.853315 | -0.161932 |
| C | -7.694829 | -0.497401 | 0.828443 |
| N | -6.479240 | -0.449286 | 0.226662 |
| C | 0.894997 | -1.250401 | -0.751557 |

| | | | |
|---|---|---|---|
| O | 1.682811 | -0.316439 | -0.371509 |
| O | 1.163737 | -2.474872 | -0.791843 |
| C | -0.498494 | -0.839937 | -1.160445 |
| O | -0.948677 | -3.951495 | -2.220897 |
| C | -2.461901 | 0.727456 | -0.943273 |
| O | -3.409913 | -0.053161 | -1.434811 |
| O | -2.614146 | 1.946212 | -0.854607 |
| H | -0.215766 | -3.587805 | -1.687190 |
| H | 3.893563 | -5.864928 | 0.333421 |
| H | 2.924004 | -4.485736 | -0.266984 |
| H | 4.691530 | -4.364663 | -0.164148 |
| H | 7.544631 | 0.028840 | -2.810572 |
| H | 6.036434 | 0.972491 | -2.761734 |
| H | 7.288883 | 1.216230 | -1.521679 |
| H | 6.857081 | -1.738981 | 0.149177 |
| H | 4.217069 | 5.622976 | 2.845328 |
| H | 4.248529 | 7.372279 | 2.594536 |
| H | 3.113196 | 6.665000 | 3.755359 |
| H | 1.903893 | 7.260984 | 1.639738 |
| H | 1.890816 | 5.521894 | 1.897379 |
| H | 3.811249 | 5.259618 | 0.289349 |
| H | 3.823071 | 7.007681 | 0.037184 |
| H | 2.705601 | 5.873147 | -1.803892 |
| H | 1.522590 | 6.871188 | -0.940193 |
| H | -4.011102 | -6.401548 | 1.161051 |
| H | -5.055164 | -4.999128 | 1.511766 |
| H | -5.120126 | -5.772477 | -0.069556 |
| H | -8.117664 | 3.323094 | 2.220192 |
| H | -6.444588 | 3.673525 | 1.767271 |
| H | -7.731594 | 4.823784 | 1.348008 |
| H | -8.730333 | 3.155487 | -0.222733 |
| H | -7.044809 | 3.441021 | -0.657659 |
| H | -0.956462 | -1.420328 | -1.955306 |
| H | 5.788135 | 0.438399 | 0.541612 |
| H | 3.216639 | -0.805889 | 0.073429 |
| H | 4.595275 | 0.767530 | -0.723237 |
| H | 3.323871 | -2.405973 | 2.829656 |
| H | 2.479193 | -2.505194 | 1.261434 |
| H | -5.818877 | -1.246942 | 0.127358 |
| H | -8.096709 | -1.419344 | 1.228146 |
| H | -5.374994 | 1.161863 | -0.659888 |
| H | 0.585710 | 4.874588 | -0.021243 |
| H | -0.075972 | 2.700127 | -0.725415 |
| H | 3.172118 | 2.403930 | -1.553907 |
| H | -3.290371 | -1.073357 | -1.262019 |
| H | -1.731309 | -3.495868 | -1.856968 |
| H | 2.831079 | -4.731656 | 2.240647 |
| H | 5.590059 | -1.305773 | -1.930196 |
| H | -2.654071 | -4.315204 | 1.063461 |
| H | -2.761125 | -4.955701 | -0.555997 |
| C | -1.155298 | 0.143195 | -0.533762 |
| H | 5.489592 | -3.902701 | 2.295259 |
| H | -0.657468 | 0.690019 | 0.260865 |
| O | 4.829236 | -4.579402 | 2.541846 |
| H | 1.215969 | 1.572022 | -0.628655 |

---------------------------------------------------------------------------------------------------------------------

**TS(A-B)$_{QM1A}$**

-----------------------------------------------------------------------------------------------------------------------------

Number of imaginary frequencies: 1 Electronic energy: HF=-2024.095582
Zero-point correction= 0.720175 (Hartree/Particle)
Thermal correction to Energy= 0.768951
Thermal correction to Enthalpy= 0.769895
Thermal correction to Gibbs Free Energy= 0.624547
Sum of electronic and zero-point Energies= -2023.375407
Sum of electronic and thermal Energies= -2023.326631
Sum of electronic and thermal Enthalpies= -2023.325687
Sum of electronic and thermal Free Energies= -2023.471035

..............................................................

Cartesian Coordinates

..............................................................

| | | | |
|---|---|---|---|
| C | -4.928493 | -4.604886 | 1.119287 |
| C | -3.915910 | -3.776851 | 0.323643 |
| C | -4.578555 | -2.764086 | -0.595530 |
| O | -5.754333 | -2.428922 | -0.429262 |
| O | -3.858024 | -2.221311 | -1.550866 |
| C | 3.180857 | -2.518714 | 1.763532 |
| C | 4.386546 | -1.801801 | 1.183323 |
| O | 5.382106 | -1.521709 | 1.889083 |
| C | 2.099847 | -1.532177 | 2.314345 |
| C | 2.628851 | -0.691158 | 3.484399 |
| N | 4.303684 | -1.471628 | -0.122350 |
| C | 5.262108 | -0.584279 | -0.787798 |
| C | 6.666739 | -1.178702 | -1.005910 |
| O | 7.440989 | -1.176671 | 0.180682 |
| C | 7.440640 | -0.374995 | -2.052430 |
| C | 4.363985 | 6.620599 | 0.557066 |
| C | 3.058100 | 6.029088 | 0.016877 |
| C | 3.284912 | 4.910674 | -1.005313 |
| C | 1.991916 | 4.277873 | -1.552284 |
| N | 1.151538 | 3.607250 | -0.575328 |
| C | 1.587202 | 2.403616 | 0.002039 |
| N | 0.763653 | 1.982862 | 1.043012 |
| N | 2.656201 | 1.828873 | -0.425696 |
| C | -6.931563 | 4.803050 | -0.498090 |
| C | -7.638872 | 3.661373 | 0.249252 |
| C | -6.832045 | 2.396281 | 0.254783 |
| N | -5.593396 | 2.343594 | 0.861680 |
| C | -7.123576 | 1.181740 | -0.324335 |
| C | -5.149806 | 1.113368 | 0.646865 |
| N | -6.041307 | 0.373068 | -0.065934 |
| C | 0.898970 | -3.229617 | -1.509151 |
| O | 2.082457 | -2.900776 | -1.272713 |
| O | 0.474954 | -4.339548 | -1.952883 |
| C | -0.252053 | -2.186150 | -1.273136 |
| O | -1.411880 | -2.787389 | -1.923673 |
| C | -1.558367 | -0.909265 | 0.484624 |
| O | -1.554976 | 0.263722 | -0.254347 |
| O | -2.419259 | -0.990963 | 1.360746 |
| H | -1.006985 | -3.694695 | -2.105156 |
| H | 1.828162 | -0.044943 | 3.859459 |
| H | 2.986708 | -1.322023 | 4.308794 |
| H | 3.461104 | -0.059297 | 3.160237 |

| | | | |
|---|---|---|---|
| H | 8.464771 | -0.755239 | -2.126782 |
| H | 6.969451 | -0.440464 | -3.039697 |
| H | 7.493530 | 0.679440 | -1.758132 |
| H | 6.800082 | -1.297379 | 0.923613 |
| H | 4.968729 | 5.848039 | 1.044506 |
| H | 4.969801 | 7.049057 | -0.251172 |
| H | 4.179745 | 7.414005 | 1.291373 |
| H | 2.453090 | 6.830071 | -0.435551 |
| H | 2.469863 | 5.636842 | 0.856235 |
| H | 3.876710 | 4.104067 | -0.560364 |
| H | 3.862915 | 5.308210 | -1.854152 |
| H | 2.265437 | 3.529658 | -2.300994 |
| H | 1.379472 | 5.044708 | -2.048903 |
| H | -4.414960 | -5.266254 | 1.825494 |
| H | -5.599806 | -3.947896 | 1.676675 |
| H | -5.547652 | -5.223615 | 0.460191 |
| H | -5.940350 | 4.966326 | -0.066642 |
| H | -6.793883 | 4.549172 | -1.554744 |
| H | -7.499036 | 5.741149 | -0.440658 |
| H | -7.828136 | 3.982992 | 1.283501 |
| H | -8.621020 | 3.468040 | -0.202418 |
| H | 0.015259 | -1.284869 | -1.847046 |
| H | 5.367277 | 0.346531 | -0.215769 |
| H | 3.446218 | -1.752594 | -0.612619 |
| H | 4.816295 | -0.323862 | -1.751534 |
| H | 2.711063 | -3.125067 | 0.984701 |
| H | 3.542516 | -3.164880 | 2.571588 |
| H | -5.933851 | -0.611398 | -0.338254 |
| H | -4.199880 | 0.702215 | 0.971820 |
| H | -7.979301 | 0.832004 | -0.883093 |
| H | 0.675634 | 4.209875 | 0.082628 |
| H | -0.227552 | 2.046440 | 0.830395 |
| H | 2.721119 | 0.906942 | 0.015193 |
| H | -0.788794 | 0.241664 | -0.846938 |
| H | -2.869030 | -2.504713 | -1.597058 |
| H | 1.297964 | -2.187189 | 2.714323 |
| H | 6.532122 | -2.209491 | -1.383967 |
| H | -3.286494 | -3.176202 | 0.995429 |
| H | -3.229413 | -4.405590 | -0.253476 |
| C | -0.496074 | -1.865706 | 0.190688 |
| H | 0.665630 | -1.272461 | 0.753864 |
| H | -0.677562 | -2.784152 | 0.756842 |
| O | 1.600533 | -0.713331 | 1.306928 |
| H | 1.007747 | 1.028210 | 1.350505 |

---

**TS(B-C)$_{QM1A}$**

---

Number of imaginary frequencies: 4 Electronic energy: HF=-2024.098575
Zero-point correction= 0.720706 (Hartree/Particle)
Thermal correction to Energy= 0.767453
Thermal correction to Enthalpy= 0.768398
Thermal correction to Gibbs Free Energy= 0.630733
Sum of electronic and zero-point Energies= -2023.377869
Sum of electronic and thermal Energies= -2023.331122
Sum of electronic and thermal Enthalpies= -2023.330177
Sum of electronic and thermal Free Energies= -2023.467842

.............................................................
Cartesian Coordinates
.............................................................

| | | | |
|---|---|---|---|
| C | -4.102575 | -5.412629 | 0.265535 |
| C | -3.229250 | -4.412150 | -0.490879 |
| C | -4.002471 | -3.192846 | -0.996143 |
| O | -5.184780 | -3.032338 | -0.667946 |
| O | -3.353878 | -2.362687 | -1.759345 |
| C | 3.523217 | -2.178283 | 1.635646 |
| C | 4.675516 | -1.372408 | 1.057951 |
| O | 5.834422 | -1.495543 | 1.543493 |
| C | 3.879613 | -3.675993 | 1.752436 |
| C | 4.152607 | -4.304092 | 0.381208 |
| N | 4.389892 | -0.555841 | 0.041781 |
| C | 5.342891 | 0.403102 | -0.512812 |
| C | 6.504283 | -0.208310 | -1.321334 |
| O | 7.516084 | -0.738920 | -0.479663 |
| C | 7.164104 | 0.843856 | -2.211903 |
| C | 3.224263 | 7.110831 | 1.733189 |
| C | 1.963829 | 6.496503 | 1.112566 |
| C | 2.167300 | 6.017750 | -0.329826 |
| C | 0.900628 | 5.443774 | -0.993677 |
| N | 0.371190 | 4.226855 | -0.407092 |
| C | 1.016055 | 3.003375 | -0.632690 |
| N | 0.504924 | 1.976948 | 0.135607 |
| N | 2.002856 | 2.941266 | -1.465610 |
| C | -7.575073 | 3.690586 | -0.185724 |
| C | -8.324682 | 2.441311 | 0.307253 |
| C | -7.379554 | 1.293702 | 0.496388 |
| N | -6.371623 | 1.365937 | 1.436978 |
| C | -7.252571 | 0.139948 | -0.242303 |
| C | -5.642970 | 0.274087 | 1.255595 |
| N | -6.143010 | -0.503195 | 0.257065 |
| C | 1.068541 | -1.485498 | -0.978897 |
| O | 1.810356 | -0.459466 | -0.798900 |
| O | 1.448985 | -2.671665 | -1.119517 |
| C | -0.443048 | -1.230730 | -0.935684 |
| O | -0.968769 | -2.347592 | -2.034335 |
| C | -2.116311 | -0.620306 | 0.858951 |
| O | -2.832484 | 0.128508 | -0.103713 |
| O | -2.519913 | -0.534683 | 2.027147 |
| H | -0.343078 | -3.062071 | -1.796839 |
| H | 4.298471 | -5.384956 | 0.484813 |
| H | 3.322129 | -4.102387 | -0.303417 |
| H | 5.067685 | -3.884029 | -0.052412 |
| H | 8.038083 | 0.410619 | -2.708463 |
| H | 6.471873 | 1.212838 | -2.976110 |
| H | 7.506446 | 1.692699 | -1.608844 |
| H | 7.058173 | -1.089862 | 0.316505 |
| H | 4.046367 | 6.386532 | 1.747388 |
| H | 3.564381 | 7.981907 | 1.159308 |
| H | 3.050420 | 7.438094 | 2.764883 |
| H | 1.148376 | 7.235164 | 1.142265 |
| H | 1.634478 | 5.652410 | 1.731753 |
| H | 2.941678 | 5.244011 | -0.370735 |
| H | 2.523575 | 6.862063 | -0.940797 |

| | | | |
|---|---|---|---|
| H | 1.130949 | 5.212161 | -2.036540 |
| H | 0.102111 | 6.200268 | -0.981763 |
| H | -3.503156 | -6.250048 | 0.641368 |
| H | -4.593004 | -4.925826 | 1.111676 |
| H | -4.892078 | -5.813869 | -0.378173 |
| H | -6.767512 | 3.929098 | 0.511158 |
| H | -7.121249 | 3.505836 | -1.165031 |
| H | -8.238129 | 4.562542 | -0.271142 |
| H | -8.826957 | 2.679323 | 1.256079 |
| H | -9.111987 | 2.165673 | -0.407289 |
| H | -0.720932 | -0.320799 | -1.466230 |
| H | 5.782803 | 0.999969 | 0.297841 |
| H | 3.409100 | -0.564703 | -0.336314 |
| H | 4.759726 | 1.078846 | -1.145320 |
| H | 3.331870 | -1.789716 | 2.643399 |
| H | 2.621434 | -2.052009 | 1.036753 |
| H | -5.738042 | -1.372106 | -0.111126 |
| H | -4.718649 | 0.024500 | 1.767658 |
| H | -7.827368 | -0.264527 | -1.062655 |
| H | -0.016221 | 4.311804 | 0.522641 |
| H | -0.500749 | 1.959716 | 0.262233 |
| H | 2.232430 | 1.955405 | -1.605663 |
| H | -2.915625 | -0.420971 | -0.899081 |
| H | -2.197773 | -2.492436 | -1.854641 |
| H | 3.012009 | -4.179717 | 2.198402 |
| H | 6.077139 | -0.998394 | -1.964818 |
| H | -2.426550 | -4.019073 | 0.146948 |
| H | -2.726147 | -4.885066 | -1.344363 |
| C | -1.004235 | -1.354527 | 0.384690 |
| H | 5.592875 | -3.165903 | 2.442954 |
| H | -0.489039 | -1.971814 | 1.114501 |
| O | 4.964761 | -3.882259 | 2.658039 |
| H | 0.891154 | 1.051850 | -0.083153 |

----------------------------------------------------------------------------------------------------------------

**$A_{QM1B}$**

----------------------------------------------------------------------------------------------------------------

Number of imaginary frequencies: 2 Electronic energy: HF=-2024.645386
Zero-point correction= 0.738594 (Hartree/Particle)
Thermal correction to Energy= 0.786992
Thermal correction to Enthalpy= 0.787936
Thermal correction to Gibbs Free Energy= 0.645007
Sum of electronic and zero-point Energies= -2023.906792
Sum of electronic and thermal Energies= -2023.858394
Sum of electronic and thermal Enthalpies= -2023.85745
Sum of electronic and thermal Free Energies= -2024.000379

............................................................

Cartesian Coordinates

............................................................

| | | | |
|---|---|---|---|
| C | -5.533553 | -5.112255 | 0.898851 |
| C | -5.019072 | -4.497843 | -0.414112 |
| C | -3.984704 | -3.385685 | -0.204482 |
| O | -4.257864 | -2.378097 | 0.453004 |
| O | -2.846886 | -3.651547 | -0.788049 |
| C | 2.586091 | -3.026440 | 1.129291 |
| C | 3.816827 | -2.528035 | 0.396427 |
| O | 4.960810 | -2.733899 | 0.850481 |

| | | | |
|---|---|---|---|
| C | 2.207333 | -2.265919 | 2.428266 |
| O | 1.655907 | -0.987552 | 2.177295 |
| C | 3.345564 | -2.192254 | 3.453755 |
| N | 3.581920 | -1.897475 | -0.768437 |
| C | 4.628345 | -1.201096 | -1.509202 |
| C | 5.700773 | -2.100406 | -2.157400 |
| O | 6.685423 | -2.516558 | -1.229588 |
| C | 6.427606 | -1.355478 | -3.277077 |
| C | 6.014209 | 4.909812 | -0.952206 |
| C | 4.920516 | 4.801560 | 0.111841 |
| C | 3.513136 | 4.674329 | -0.482617 |
| C | 2.409913 | 4.484418 | 0.571186 |
| N | 2.574431 | 3.301992 | 1.420911 |
| C | 2.579875 | 2.006163 | 0.890773 |
| N | 3.339115 | 1.047007 | 1.326506 |
| N | 1.701616 | 1.801136 | -0.137873 |
| C | -7.629655 | 4.333841 | -0.526748 |
| C | -6.198090 | 3.768934 | -0.506609 |
| C | -6.112525 | 2.266643 | -0.572818 |
| N | -6.786827 | 1.544388 | -1.538644 |
| C | -5.354167 | 1.414044 | 0.202693 |
| C | -6.433010 | 0.287408 | -1.343493 |
| N | -5.566478 | 0.146491 | -0.304096 |
| C | -0.233247 | -1.246773 | -0.958393 |
| O | 0.959981 | -0.886073 | -1.154147 |
| O | -0.657866 | -2.429193 | -0.874319 |
| C | -1.231093 | -0.093691 | -0.716655 |
| O | -2.473862 | -0.417766 | -1.326900 |
| C | -1.365055 | 0.107246 | 0.813068 |
| C | -1.659674 | 1.528673 | 1.204901 |
| O | -1.214149 | 2.532850 | 0.675381 |
| H | -0.415870 | -0.155023 | 1.308847 |
| H | -2.130559 | -0.570721 | 1.198796 |
| H | -3.098654 | 0.299270 | -1.152143 |
| H | 2.988750 | -1.680971 | 4.354002 |
| H | 3.697935 | -3.191937 | 3.731002 |
| H | 4.202505 | -1.648159 | 3.048736 |
| H | 7.242945 | -1.975311 | -3.661725 |
| H | 5.751592 | -1.112189 | -4.104145 |
| H | 6.864732 | -0.426227 | -2.894192 |
| H | 6.212236 | -2.662395 | -0.378951 |
| H | 6.019274 | 4.025524 | -1.598388 |
| H | 5.861782 | 5.786317 | -1.593192 |
| H | 7.008103 | 4.996935 | -0.500288 |
| H | 4.961839 | 5.677281 | 0.776393 |
| H | 5.126590 | 3.925696 | 0.739364 |
| H | 3.478562 | 3.821737 | -1.171116 |
| H | 3.279762 | 5.569520 | -1.076593 |
| H | 1.433635 | 4.435177 | 0.081638 |
| H | 2.373994 | 5.351256 | 1.242966 |
| H | -6.327251 | -5.839927 | 0.702058 |
| H | -4.728532 | -5.625070 | 1.435152 |
| H | -5.930358 | -4.330900 | 1.553143 |
| H | -8.163718 | 3.972209 | -1.409454 |
| H | -8.193007 | 4.013901 | 0.356511 |
| H | -7.618234 | 5.429061 | -0.548088 |

| | | | |
|---|---|---|---|
| H | -5.649050 | 4.192322 | -1.359726 |
| H | -5.671541 | 4.114309 | 0.391720 |
| H | -0.808408 | 0.812774 | -1.164524 |
| H | 5.135260 | -0.481995 | -0.851598 |
| H | 2.603010 | -1.674436 | -0.996454 |
| H | 4.112964 | -0.632315 | -2.289128 |
| H | 1.710363 | -3.018674 | 0.474062 |
| H | 2.799474 | -4.067076 | 1.403103 |
| H | -5.130966 | -0.725407 | 0.013572 |
| H | -6.767542 | -0.557729 | -1.929690 |
| H | -4.704691 | 1.581065 | 1.048472 |
| H | 3.278804 | 3.414140 | 2.137282 |
| H | 4.023111 | 1.391018 | 1.999352 |
| H | 1.601387 | 0.831146 | -0.457896 |
| H | 0.809352 | 2.278002 | -0.054718 |
| H | -2.076094 | -2.948752 | -0.740488 |
| H | 1.386882 | -2.846673 | 2.872158 |
| H | 2.350729 | -0.386240 | 1.808586 |
| H | 5.177942 | -2.971072 | -2.595309 |
| H | -4.585894 | -5.265965 | -1.059258 |
| H | -5.861957 | -4.047349 | -0.953629 |
| O | -2.483225 | 1.627438 | 2.282384 |
| H | -2.541843 | 2.579087 | 2.475425 |

---

**$B_{QM1B}$**

---

Number of imaginary frequencies: 1 Electronic energy: HF=-2024.590207
Zero-point correction= 0.735535 (Hartree/Particle)
Thermal correction to Energy= 0.786136
Thermal correction to Enthalpy= 0.787081
Thermal correction to Gibbs Free Energy= 0.638095
Sum of electronic and zero-point Energies= -2023.854672
Sum of electronic and thermal Energies= -2023.804071
Sum of electronic and thermal Enthalpies= -2023.803126
Sum of electronic and thermal Free Energies= -2023.952112

............................................................

Cartesian Coordinates

............................................................

| | | | |
|---|---|---|---|
| C | -5.521934 | -4.691455 | 1.330680 |
| C | -5.265559 | -4.257818 | -0.115202 |
| C | -5.095034 | -2.736459 | -0.340193 |
| O | -4.711954 | -2.039308 | 0.653187 |
| O | -5.324617 | -2.317347 | -1.503646 |
| C | 2.755239 | -3.210786 | 1.462837 |
| C | 3.967959 | -2.508618 | 0.873843 |
| O | 4.943278 | -2.208121 | 1.584767 |
| C | 1.836956 | -2.274749 | 2.291853 |
| O | 1.267642 | -1.269162 | 1.476978 |
| C | 2.511775 | -1.696956 | 3.539479 |
| N | 3.912058 | -2.238681 | -0.449972 |
| C | 4.879126 | -1.382089 | -1.142372 |
| C | 6.310159 | -1.947101 | -1.255281 |
| O | 7.057860 | -1.750293 | -0.070942 |
| C | 7.073597 | -1.257246 | -2.385007 |
| C | 6.711577 | 4.550711 | -0.297544 |
| C | 5.321064 | 4.579686 | 0.343949 |

| | | | |
|---|---|---|---|
| C | 4.181742 | 4.412915 | -0.669858 |
| C | 2.775945 | 4.441323 | -0.046422 |
| N | 2.507894 | 3.361615 | 0.909682 |
| C | 2.309708 | 2.041905 | 0.503392 |
| N | 2.805916 | 1.004288 | 1.100218 |
| N | 1.519468 | 1.893610 | -0.610462 |
| C | -6.926251 | 4.967969 | 0.411595 |
| C | -5.667818 | 4.202970 | 0.849119 |
| C | -5.644141 | 2.770292 | 0.390830 |
| N | -5.845973 | 2.435232 | -0.930265 |
| C | -5.402562 | 1.626861 | 1.131337 |
| C | -5.721676 | 1.115600 | -0.971687 |
| N | -5.459177 | 0.573982 | 0.242683 |
| C | 0.314296 | -2.010809 | -1.435174 |
| O | 0.375424 | -0.714314 | -1.686209 |
| O | 1.084453 | -2.938101 | -1.449623 |
| C | -0.994187 | -1.407564 | -1.272812 |
| O | -3.508524 | -0.890600 | -2.991035 |
| C | -1.533736 | -0.987452 | 0.061944 |
| C | -2.099678 | 0.414815 | 0.012811 |
| O | -2.224981 | 1.096031 | -0.987898 |
| H | -0.778499 | -1.057732 | 0.850785 |
| H | -2.382555 | -1.640022 | 0.327465 |
| H | -3.385878 | -0.048641 | -2.530240 |
| H | 1.806814 | -1.044346 | 4.063494 |
| H | 2.822235 | -2.495644 | 4.221993 |
| H | 3.405646 | -1.126125 | 3.276317 |
| H | 8.110586 | -1.604333 | -2.390559 |
| H | 6.627009 | -1.470297 | -3.362124 |
| H | 7.083763 | -0.172414 | -2.231017 |
| H | 6.438723 | -1.909658 | 0.674308 |
| H | 6.887508 | 3.600114 | -0.812358 |
| H | 6.824424 | 5.353193 | -1.035633 |
| H | 7.500047 | 4.673104 | 0.452061 |
| H | 5.186718 | 5.525477 | 0.888462 |
| H | 5.263614 | 3.780954 | 1.095021 |
| H | 4.304645 | 3.469041 | -1.216050 |
| H | 4.240614 | 5.215824 | -1.417999 |
| H | 2.024890 | 4.394809 | -0.837584 |
| H | 2.618393 | 5.389242 | 0.482772 |
| H | -5.543950 | -5.784020 | 1.425143 |
| H | -4.743208 | -4.294329 | 1.986852 |
| H | -6.478860 | -4.301801 | 1.695139 |
| H | -7.039587 | 4.904834 | -0.673704 |
| H | -7.825568 | 4.536128 | 0.864149 |
| H | -6.870954 | 6.024302 | 0.699684 |
| H | -4.789193 | 4.726396 | 0.444718 |
| H | -5.567341 | 4.234631 | 1.941734 |
| H | -1.701882 | -1.395836 | -2.109880 |
| H | 4.930940 | -0.406974 | -0.640769 |
| H | 3.041492 | -2.453025 | -0.929145 |
| H | 4.470914 | -1.221598 | -2.144494 |
| H | 2.149253 | -3.660078 | 0.672890 |
| H | 3.131930 | -4.008800 | 2.112415 |
| H | -5.208784 | -0.433783 | 0.420849 |
| H | -5.792227 | 0.497636 | -1.855990 |

| | | | |
|---|---|---|---|
| H | -5.249928 | 1.471521 | 2.189855 |
| H | 3.070755 | 3.415758 | 1.748147 |
| H | 3.501530 | 1.263357 | 1.797190 |
| H | 1.307237 | 0.932776 | -0.860285 |
| H | 0.716436 | 2.504277 | -0.681925 |
| H | -4.175159 | -1.376475 | -2.446421 |
| H | 0.995008 | -2.899589 | 2.619578 |
| H | 1.899110 | -0.513437 | 1.346335 |
| H | 6.221141 | -3.023123 | -1.493953 |
| H | -4.345763 | -4.732572 | -0.486697 |
| H | -6.063469 | -4.598691 | -0.783562 |
| O | -2.474122 | 0.831910 | 1.233872 |
| H | -3.054931 | 1.608421 | 1.101533 |

------------------------------------------------------------------------------------------------------------

**$C_{QM1B}$**

------------------------------------------------------------------------------------------------------------

Number of imaginary frequencies: 3 Electronic energy: HF=-2024.641969
Zero-point correction= 0.734019 (Hartree/Particle)
Thermal correction to Energy= 0.782848
Thermal correction to Enthalpy= 0.783792
Thermal correction to Gibbs Free Energy= 0.637506
Sum of electronic and zero-point Energies= -2023.90795
Sum of electronic and thermal Energies= -2023.859121
Sum of electronic and thermal Enthalpies= -2023.858177
Sum of electronic and thermal Free Energies= -2024.004463

............................................................

Cartesian Coordinates

............................................................

| | | | |
|---|---|---|---|
| C | -5.674508 | -4.528254 | -0.011994 |
| C | -4.258479 | -3.943709 | -0.086547 |
| C | -4.244469 | -2.412378 | -0.286677 |
| O | -5.213192 | -1.926442 | -0.927478 |
| O | -3.269845 | -1.750925 | 0.188913 |
| C | 2.514340 | -3.038393 | 0.947277 |
| C | 3.664830 | -2.066506 | 0.705500 |
| O | 4.492006 | -1.763830 | 1.594217 |
| C | 1.718628 | -2.984662 | 2.267056 |
| O | 0.910023 | -1.809148 | 2.345969 |
| C | 2.557018 | -3.102634 | 3.539735 |
| N | 3.757095 | -1.660144 | -0.574313 |
| C | 4.793073 | -0.782463 | -1.111000 |
| C | 6.236433 | -1.309392 | -1.001655 |
| O | 6.793823 | -1.076066 | 0.281593 |
| C | 7.144105 | -0.613244 | -2.013089 |
| C | 6.461866 | 4.923761 | 0.907935 |
| C | 4.972653 | 4.606755 | 1.071638 |
| C | 4.377772 | 3.894772 | -0.151556 |
| C | 2.889419 | 3.540972 | -0.020326 |
| N | 2.650852 | 2.591223 | 1.062140 |
| C | 1.984118 | 1.410556 | 0.973407 |
| N | 2.266417 | 0.469161 | 1.879099 |
| N | 1.042838 | 1.198839 | 0.062642 |
| C | -6.769829 | 4.675509 | 1.658407 |
| C | -6.223150 | 3.345835 | 2.195872 |
| C | -6.347165 | 2.220770 | 1.208979 |
| N | -7.558896 | 1.924659 | 0.613715 |

| | | | |
|---|---|---|---|
| C | -5.381974 | 1.338620 | 0.770684 |
| C | -7.305631 | 0.878261 | -0.157793 |
| N | -6.006933 | 0.488964 | -0.109570 |
| C | 1.122229 | -0.682086 | -2.558919 |
| O | 1.114379 | -1.025047 | -1.312062 |
| O | 2.058755 | -0.868326 | -3.353173 |
| C | -0.106527 | 0.036160 | -3.066507 |
| O | -1.495532 | -2.702500 | 1.779494 |
| C | -1.264394 | 0.096914 | -2.392915 |
| C | -2.427077 | 0.846309 | -2.928369 |
| O | -2.515404 | 1.353362 | -4.033644 |
| H | -0.009735 | -2.041249 | 2.017440 |
| H | -1.398770 | -0.403665 | -1.438193 |
| H | -1.975179 | -2.533102 | 2.599989 |
| H | 1.894921 | -3.126573 | 4.410788 |
| H | 3.152709 | -4.021975 | 3.527186 |
| H | 3.250508 | -2.265856 | 3.635019 |
| H | 8.178017 | -0.937047 | -1.863367 |
| H | 6.846813 | -0.842970 | -3.041116 |
| H | 7.109451 | 0.473262 | -1.872246 |
| H | 6.096257 | -1.321020 | 0.925943 |
| H | 7.044731 | 4.008831 | 0.758360 |
| H | 6.634687 | 5.572548 | 0.041972 |
| H | 6.859353 | 5.433265 | 1.791282 |
| H | 4.412508 | 5.532413 | 1.261793 |
| H | 4.851135 | 3.985691 | 1.970224 |
| H | 4.941626 | 2.976757 | -0.359063 |
| H | 4.491368 | 4.540090 | -1.031999 |
| H | 2.541299 | 3.081098 | -0.948862 |
| H | 2.297914 | 4.453815 | 0.137449 |
| H | -5.656197 | -5.623628 | 0.043512 |
| H | -6.208216 | -4.154929 | 0.869063 |
| H | -6.246604 | -4.222919 | -0.890530 |
| H | -6.192499 | 5.013896 | 0.790808 |
| H | -7.805739 | 4.545651 | 1.334065 |
| H | -6.736093 | 5.464218 | 2.420076 |
| H | -5.170992 | 3.457920 | 2.486699 |
| H | -6.770049 | 3.091837 | 3.116220 |
| H | -0.024177 | 0.508881 | -4.042406 |
| H | 4.773348 | 0.201305 | -0.613643 |
| H | 2.877784 | -1.674198 | -1.111592 |
| H | 4.513906 | -0.633754 | -2.157716 |
| H | 1.798587 | -2.955665 | 0.124722 |
| H | 2.969231 | -4.037290 | 0.874550 |
| H | -5.593063 | -0.389857 | -0.522213 |
| H | -8.034543 | 0.364661 | -0.771209 |
| H | -4.333110 | 1.224996 | 0.998693 |
| H | 3.147256 | 2.739436 | 1.927919 |
| H | 3.224123 | 0.355192 | 2.184669 |
| H | 0.934426 | 0.227904 | -0.388024 |
| H | 0.689555 | 1.981555 | -0.466699 |
| H | -2.148053 | -2.359448 | 1.066009 |
| H | 1.034855 | -3.843317 | 2.225970 |
| H | 1.661959 | -0.379271 | 1.972215 |
| H | 6.212973 | -2.389710 | -1.231545 |
| H | -3.667379 | -4.189499 | 0.802324 |

| | | | |
|---|---|---|---|
| H | -3.720508 | -4.385147 | -0.937545 |
| O | -3.406302 | 0.932173 | -2.002744 |
| H | -4.159377 | 1.393091 | -2.406532 |

---

**TS(A-B)$_{QM1B}$**

---

Number of imaginary frequencies: 2 Electronic energy: HF=-2024.580996
Zero-point correction= 0.733883 (Hartree/Particle)
Thermal correction to Energy= 0.784129
Thermal correction to Enthalpy= 0.785073
Thermal correction to Gibbs Free Energy= 0.634990
Sum of electronic and zero-point Energies= -2023.847113
Sum of electronic and thermal Energies= -2023.796867
Sum of electronic and thermal Enthalpies= -2023.795923
Sum of electronic and thermal Free Energies= -2023.946006

............................................................

Cartesian Coordinates

............................................................

| | | | |
|---|---|---|---|
| C | -5.503603 | -4.834328 | 1.172776 |
| C | -5.156769 | -4.332122 | -0.229883 |
| C | -4.903281 | -2.815551 | -0.366465 |
| O | -4.766694 | -2.132208 | 0.689499 |
| O | -4.837433 | -2.383404 | -1.555211 |
| C | 2.694122 | -3.031465 | 1.444109 |
| C | 3.927799 | -2.408893 | 0.819043 |
| O | 5.009789 | -2.371126 | 1.430135 |
| C | 2.016911 | -2.202434 | 2.568318 |
| O | 1.372668 | -1.048065 | 2.065030 |
| C | 2.951801 | -1.867993 | 3.734974 |
| N | 3.771418 | -1.937758 | -0.436538 |
| C | 4.791537 | -1.163013 | -1.137134 |
| C | 6.073325 | -1.934841 | -1.515193 |
| O | 6.976486 | -2.033740 | -0.430573 |
| C | 6.813972 | -1.228082 | -2.649474 |
| C | 6.380120 | 4.858927 | -0.340702 |
| C | 4.960798 | 4.835456 | 0.233583 |
| C | 3.901509 | 4.391316 | -0.783287 |
| C | 2.474629 | 4.321459 | -0.213346 |
| N | 2.307964 | 3.364909 | 0.885898 |
| C | 2.352805 | 1.987997 | 0.679344 |
| N | 2.946144 | 1.141983 | 1.462041 |
| N | 1.693682 | 1.551363 | -0.447207 |
| C | -7.274342 | 4.738317 | 0.116233 |
| C | -5.954930 | 4.108004 | 0.594549 |
| C | -5.792519 | 2.661876 | 0.210005 |
| N | -5.903413 | 2.247707 | -1.099741 |
| C | -5.494928 | 1.578253 | 1.017494 |
| C | -5.675908 | 0.940690 | -1.065644 |
| N | -5.421829 | 0.483676 | 0.186120 |
| C | -0.057038 | -2.113797 | -1.746161 |
| O | 0.985378 | -1.425028 | -1.346173 |
| O | -0.227143 | -3.217346 | -2.194009 |
| C | -0.757696 | -0.901094 | -1.316636 |
| O | -2.893048 | -0.770812 | -2.231999 |
| C | -1.232968 | -0.702951 | 0.087279 |
| C | -1.714041 | 0.713451 | 0.307241 |

| | | | |
|---|---|---|---|
| O | -1.429818 | 1.672685 | -0.391805 |
| H | -0.408331 | -0.907720 | 0.791568 |
| H | -2.050494 | -1.392100 | 0.324661 |
| H | -3.270827 | 0.096760 | -2.034103 |
| H | 2.393788 | -1.315789 | 4.497548 |
| H | 3.360510 | -2.777703 | 4.187618 |
| H | 3.797641 | -1.262543 | 3.400042 |
| H | 7.762814 | -1.737377 | -2.839901 |
| H | 6.225941 | -1.224651 | -3.573458 |
| H | 7.039249 | -0.192594 | -2.370473 |
| H | 6.429017 | -2.190590 | 0.368969 |
| H | 6.678515 | 3.865085 | -0.691285 |
| H | 6.452567 | 5.545325 | -1.192244 |
| H | 7.110195 | 5.180298 | 0.409410 |
| H | 4.701510 | 5.832710 | 0.617409 |
| H | 4.943601 | 4.158863 | 1.097994 |
| H | 4.164023 | 3.404805 | -1.185873 |
| H | 3.899531 | 5.085457 | -1.635289 |
| H | 1.773545 | 4.058493 | -1.007693 |
| H | 2.169237 | 5.304482 | 0.165400 |
| H | -5.616007 | -5.925167 | 1.188514 |
| H | -4.723960 | -4.552154 | 1.885479 |
| H | -6.438208 | -4.389797 | 1.530522 |
| H | -7.371875 | 4.620860 | -0.966251 |
| H | -8.135633 | 4.247634 | 0.582334 |
| H | -7.314766 | 5.806426 | 0.359091 |
| H | -5.122752 | 4.688869 | 0.171359 |
| H | -5.869712 | 4.200831 | 1.684965 |
| H | -0.744915 | -0.059952 | -1.991944 |
| H | 5.082551 | -0.296301 | -0.528818 |
| H | 2.829765 | -1.924242 | -0.820455 |
| H | 4.310002 | -0.788250 | -2.045074 |
| H | 1.937377 | -3.236218 | 0.682025 |
| H | 3.014940 | -3.989451 | 1.869465 |
| H | -5.153047 | -0.507918 | 0.425353 |
| H | -5.694331 | 0.266276 | -1.911205 |
| H | -5.369817 | 1.489893 | 2.087274 |
| H | 2.785637 | 3.635086 | 1.735199 |
| H | 3.519812 | 1.610342 | 2.161399 |
| H | 1.688771 | 0.541552 | -0.544709 |
| H | 0.773405 | 1.951514 | -0.606533 |
| H | -3.616683 | -1.418033 | -1.953250 |
| H | 1.208828 | -2.840777 | 2.950672 |
| H | 2.039779 | -0.341906 | 1.861038 |
| H | 5.762805 | -2.936455 | -1.865275 |
| H | -4.251127 | -4.831115 | -0.600537 |
| H | -5.942132 | -4.590575 | -0.949938 |
| O | -2.506706 | 0.807404 | 1.381178 |
| H | -2.914152 | 1.692678 | 1.370271 |

------------------------------------------------------------------------------------------------------------

**TS(B-C)$_{QM1B}$**

------------------------------------------------------------------------------------------------------------

Number of imaginary frequencies: 3 Electronic energy: HF=-2024.568006
Zero-point correction= 0.728537 (Hartree/Particle)
Thermal correction to Energy= 0.777914
Thermal correction to Enthalpy= 0.778858

Thermal correction to Gibbs Free Energy= 0.631944
Sum of electronic and zero-point Energies= -2023.839469
Sum of electronic and thermal Energies= -2023.790092
Sum of electronic and thermal Enthalpies= -2023.789148
Sum of electronic and thermal Free Energies= -2023.936062

............................................................

Cartesian Coordinates

............................................................

| | | | |
|---|---|---|---|
| C | 6.164037 | -4.524205 | -0.098130 |
| C | 4.917090 | -4.014821 | 0.621506 |
| C | 4.680700 | -2.495406 | 0.466303 |
| O | 5.700613 | -1.765809 | 0.346599 |
| O | 3.478747 | -2.093435 | 0.511404 |
| C | -2.065126 | -3.049574 | -0.893902 |
| C | -3.367936 | -2.640034 | -0.232212 |
| O | -4.457492 | -3.029470 | -0.691548 |
| C | -1.698348 | -2.289075 | -2.187593 |
| O | -1.306847 | -0.930722 | -1.943867 |
| C | -2.780032 | -2.330688 | -3.266710 |
| N | -3.255712 | -1.876659 | 0.869785 |
| C | -4.403309 | -1.302654 | 1.563261 |
| C | -5.343572 | -2.314918 | 2.250647 |
| O | -6.274008 | -2.880999 | 1.345181 |
| C | -6.152992 | -1.635879 | 3.354085 |
| C | -6.182161 | 4.660745 | 0.729205 |
| C | -5.227697 | 4.609430 | -0.464478 |
| C | -3.764445 | 4.788280 | -0.048999 |
| C | -2.761450 | 4.607191 | -1.198634 |
| N | -2.808919 | 3.305356 | -1.865515 |
| C | -2.613042 | 2.089458 | -1.235169 |
| N | -3.073696 | 0.955002 | -1.694199 |
| N | -1.893178 | 2.113323 | -0.087389 |
| C | 7.060185 | 4.847043 | 0.174596 |
| C | 5.616800 | 4.572007 | 0.615355 |
| C | 5.265238 | 3.116176 | 0.767398 |
| N | 4.118839 | 2.737653 | 1.437115 |
| C | 5.929036 | 1.994095 | 0.311672 |
| C | 4.109635 | 1.411357 | 1.377326 |
| N | 5.177815 | 0.913371 | 0.707850 |
| C | 0.109407 | 0.001296 | 1.612191 |
| O | -1.151896 | -0.035869 | 1.587632 |
| O | 0.919758 | -0.124776 | 2.542264 |
| C | 0.893471 | 0.208844 | 0.359939 |
| O | 1.493852 | -3.748171 | -0.068069 |
| C | 1.010955 | -0.778588 | -0.624599 |
| C | 2.045940 | -0.566997 | -1.685695 |
| O | 2.582471 | 0.482169 | -1.961330 |
| H | -0.110005 | -0.747225 | -1.184415 |
| H | 0.926946 | -1.823539 | -0.282257 |
| H | 1.732931 | -3.730374 | -1.005151 |
| H | -2.438214 | -1.786969 | -4.152571 |
| H | -2.996274 | -3.365371 | -3.549681 |
| H | -3.713684 | -1.891582 | -2.907136 |
| H | -6.875970 | -2.345016 | 3.767078 |
| H | -5.507477 | -1.280064 | 4.163866 |
| H | -6.711829 | -0.785059 | 2.948152 |

| | | | |
|---|---|---|---|
| H | -5.780280 | -3.041974 | 0.513301 |
| H | -5.952165 | 3.866927 | 1.447687 |
| H | -6.107679 | 5.617715 | 1.258766 |
| H | -7.223916 | 4.534311 | 0.416558 |
| H | -5.502190 | 5.382256 | -1.197038 |
| H | -5.343072 | 3.644330 | -0.972770 |
| H | -3.523733 | 4.070502 | 0.742963 |
| H | -3.616447 | 5.789334 | 0.378658 |
| H | -1.742548 | 4.796256 | -0.837964 |
| H | -2.939755 | 5.355503 | -1.978668 |
| H | 6.361085 | -5.579823 | 0.126311 |
| H | 6.060282 | -4.428469 | -1.185455 |
| H | 7.031425 | -3.928398 | 0.196334 |
| H | 7.776455 | 4.422777 | 0.886485 |
| H | 7.267570 | 4.401808 | -0.805099 |
| H | 7.252588 | 5.922977 | 0.097286 |
| H | 5.423676 | 5.072262 | 1.573379 |
| H | 4.928655 | 5.045716 | -0.101415 |
| H | 1.425730 | 1.152185 | 0.226977 |
| H | -5.005618 | -0.704157 | 0.866067 |
| H | -2.350480 | -1.469363 | 1.107475 |
| H | -3.979949 | -0.625249 | 2.309700 |
| H | -1.221253 | -2.992741 | -0.200385 |
| H | -2.182301 | -4.105167 | -1.161865 |
| H | 5.364710 | -0.103387 | 0.527047 |
| H | 3.362459 | 0.764769 | 1.818418 |
| H | 6.846130 | 1.871869 | -0.244857 |
| H | -3.392252 | 3.250745 | -2.685958 |
| H | -3.687149 | 1.086619 | -2.494618 |
| H | -1.732506 | 1.244957 | 0.441483 |
| H | -1.280018 | 2.888323 | 0.104052 |
| H | 2.246051 | -3.207519 | 0.320383 |
| H | -0.792725 | -2.770105 | -2.576524 |
| H | -2.087162 | -0.270468 | -1.761883 |
| H | -4.709482 | -3.096684 | 2.706799 |
| H | 4.018502 | -4.549671 | 0.297124 |
| H | 5.009804 | -4.206145 | 1.700191 |
| O | 2.263287 | -1.707019 | -2.394869 |
| H | 2.972912 | -1.489627 | -3.021707 |

---

**A$_{QM1C}$**

---

Number of imaginary frequencies: 2 Electronic energy: HF=-2024.144225
Zero-point correction= 0.723368 (Hartree/Particle)
Thermal correction to Energy= 0.771497
Thermal correction to Enthalpy= 0.772441
Thermal correction to Gibbs Free Energy= 0.626061
Sum of electronic and zero-point Energies= -2023.420857
Sum of electronic and thermal Energies= -2023.372728
Sum of electronic and thermal Enthalpies= -2023.371784
Sum of electronic and thermal Free Energies= -2023.518164

.............................................................

Cartesian Coordinates

.............................................................

| | | | |
|---|---|---|---|
| C | -6.152166 | -4.271768 | 1.428934 |
| C | -5.469673 | -2.930592 | 1.697658 |

| | | | |
|---|---|---|---|
| C | -6.108060 | -1.728384 | 0.975972 |
| O | -7.217219 | -1.903084 | 0.395241 |
| O | -5.477365 | -0.631645 | 1.024898 |
| C | 2.158413 | -3.272068 | 0.909228 |
| C | 3.031675 | -3.551518 | -0.308781 |
| O | 3.477484 | -4.687629 | -0.551536 |
| C | 2.614044 | -2.057804 | 1.757491 |
| O | 2.495348 | -0.897112 | 0.946842 |
| C | 4.023341 | -2.241342 | 2.330942 |
| N | 3.256625 | -2.469745 | -1.089696 |
| C | 4.244522 | -2.406239 | -2.157287 |
| C | 3.977279 | -3.326134 | -3.367521 |
| O | 4.366997 | -4.662201 | -3.113725 |
| C | 4.765144 | -2.856298 | -4.589662 |
| C | 6.388274 | 3.412042 | -3.148317 |
| C | 6.794146 | 3.127950 | -1.697148 |
| C | 6.126810 | 4.072936 | -0.690368 |
| C | 6.563200 | 3.858787 | 0.770552 |
| N | 6.278342 | 2.540938 | 1.344949 |
| C | 4.949693 | 2.045045 | 1.380960 |
| N | 4.664466 | 0.803136 | 1.110010 |
| N | 4.004451 | 2.945457 | 1.715335 |
| C | -7.176776 | 4.759135 | -2.230659 |
| C | -6.538515 | 3.446185 | -2.714395 |
| C | -7.198612 | 2.215612 | -2.161676 |
| N | -8.482100 | 1.850810 | -2.531227 |
| C | -6.690519 | 1.303494 | -1.259903 |
| C | -8.714286 | 0.733607 | -1.854611 |
| N | -7.669878 | 0.362057 | -1.073214 |
| C | 0.700233 | 1.913163 | 2.472553 |
| O | 1.448893 | 2.427173 | 1.559259 |
| O | 0.961378 | 1.769119 | 3.676024 |
| C | -0.672347 | 1.423797 | 1.920003 |
| O | -0.781695 | 1.734699 | 0.540928 |
| C | -0.805821 | -0.087664 | 2.162255 |
| C | -2.110604 | -0.722521 | 1.705018 |
| O | -3.180855 | 0.000171 | 2.005721 |
| O | -2.144774 | -1.821028 | 1.159564 |
| H | -0.695299 | -0.261350 | 3.240529 |
| H | 0.009325 | -0.603549 | 1.649218 |
| H | 0.117553 | 2.081314 | 0.360858 |
| H | 4.317037 | -1.356778 | 2.902120 |
| H | 4.056684 | -3.115438 | 2.991379 |
| H | 4.760996 | -2.388831 | 1.533760 |
| H | 4.631744 | -3.569800 | -5.408059 |
| H | 4.432494 | -1.868999 | -4.927728 |
| H | 5.835663 | -2.808163 | -4.357973 |
| H | 4.067491 | -4.861790 | -2.196418 |
| H | 5.305199 | 3.314151 | -3.277104 |
| H | 6.665914 | 4.430515 | -3.445591 |
| H | 6.870897 | 2.717197 | -3.844217 |
| H | 7.888362 | 3.202049 | -1.602415 |
| H | 6.529863 | 2.093001 | -1.447341 |
| H | 5.037780 | 3.961581 | -0.742660 |
| H | 6.352077 | 5.114776 | -0.963361 |
| H | 6.094159 | 4.626037 | 1.395263 |

| | | | |
|---|---|---|---|
| H | 7.646625 | 4.013922 | 0.863084 |
| H | -5.682410 | -5.082345 | 2.001541 |
| H | -7.214244 | -4.226531 | 1.688441 |
| H | -6.096880 | -4.530459 | 0.366517 |
| H | -8.242895 | 4.773352 | -2.479663 |
| H | -7.088098 | 4.854688 | -1.143117 |
| H | -6.700962 | 5.635243 | -2.690531 |
| H | -6.580883 | 3.420869 | -3.812961 |
| H | -5.476436 | 3.433384 | -2.442393 |
| H | -1.469645 | 1.936726 | 2.476593 |
| H | 5.248870 | -2.654491 | -1.780143 |
| H | 2.945483 | -1.593217 | -0.654037 |
| H | 4.261240 | -1.363954 | -2.490635 |
| H | 1.130689 | -3.078764 | 0.577533 |
| H | 2.149294 | -4.179870 | 1.517410 |
| H | -7.558433 | -0.498522 | -0.472594 |
| H | -9.628589 | 0.155214 | -1.905221 |
| H | -5.751655 | 1.220730 | -0.733484 |
| H | 6.918595 | 1.828983 | 1.016918 |
| H | 5.472476 | 0.317073 | 0.722515 |
| H | 2.964899 | 2.668373 | 1.750356 |
| H | 4.294964 | 3.712384 | 2.302292 |
| H | -4.043593 | -0.390678 | 1.617003 |
| H | 1.909735 | -1.978235 | 2.598923 |
| H | 3.165120 | -0.196674 | 1.180306 |
| H | 2.898114 | -3.262062 | -3.594812 |
| H | -5.485149 | -2.703746 | 2.773511 |
| H | -4.409415 | -2.950116 | 1.420122 |

--------------------------------------------------------------------------------------------------------------------

**$C_{QM1C}$**

--------------------------------------------------------------------------------------------------------------------

Number of imaginary frequencies: 2 Electronic energy: HF=-2024.139564
Zero-point correction= 0.722150 (Hartree/Particle)
Thermal correction to Energy= 0.771176
Thermal correction to Enthalpy= 0.772120
Thermal correction to Gibbs Free Energy= 0.626014
Sum of electronic and zero-point Energies= -2023.417414
Sum of electronic and thermal Energies= -2023.368388
Sum of electronic and thermal Enthalpies= -2023.367444
Sum of electronic and thermal Free Energies= -2023.513550

............................................................

Cartesian Coordinates

............................................................

| | | | |
|---|---|---|---|
| C | -5.138004 | -4.575609 | 1.145553 |
| C | -5.611677 | -3.813437 | -0.097743 |
| C | -5.907010 | -2.316471 | 0.164028 |
| O | -6.626474 | -2.057039 | 1.182647 |
| O | -5.454631 | -1.475435 | -0.653175 |
| C | 3.071300 | -3.025486 | 2.107423 |
| C | 4.389880 | -2.682043 | 1.422744 |
| O | 5.453422 | -3.250139 | 1.731286 |
| C | 2.179184 | -1.815642 | 2.494726 |
| O | 1.764354 | -1.093994 | 1.347648 |
| C | 2.863905 | -0.852017 | 3.464193 |

| | | | |
|---|---|---|---|
| N | 4.292135 | -1.749239 | 0.450138 |
| C | 5.415914 | -1.124474 | -0.237928 |
| C | 6.253282 | -2.067809 | -1.124328 |
| O | 7.159256 | -2.846588 | -0.362215 |
| C | 7.083115 | -1.271856 | -2.130162 |
| C | 7.086623 | 4.815956 | 0.915126 |
| C | 5.673457 | 4.375025 | 1.312520 |
| C | 4.575835 | 4.961000 | 0.415189 |
| C | 3.148139 | 4.564220 | 0.840676 |
| N | 2.834007 | 3.152411 | 0.738560 |
| C | 2.561162 | 2.516133 | -0.469829 |
| N | 2.577177 | 1.140805 | -0.373008 |
| N | 2.320350 | 3.083474 | -1.611112 |
| C | -6.561450 | 5.073267 | 0.339544 |
| C | -7.185600 | 3.905316 | -0.444301 |
| C | -7.319844 | 2.639309 | 0.355039 |
| N | -8.100499 | 2.584730 | 1.496659 |
| C | -6.754516 | 1.402914 | 0.112278 |
| C | -7.988022 | 1.327724 | 1.909998 |
| N | -7.190622 | 0.574871 | 1.113713 |
| C | -0.232038 | 0.664869 | -2.951557 |
| O | 1.123257 | 0.844209 | -2.862397 |
| O | -0.955149 | 1.524957 | -3.418100 |
| C | -0.702515 | -0.627573 | -2.402656 |
| O | -3.628760 | -2.374885 | -2.651166 |
| C | 0.081189 | -1.398780 | -1.632263 |
| C | -0.346311 | -2.622847 | -0.844246 |
| O | -1.253750 | -3.355754 | -1.262605 |
| O | 0.327397 | -2.790095 | 0.240057 |
| H | 1.106197 | -1.734824 | 0.817192 |
| H | 1.117658 | -1.111208 | -1.479401 |
| H | -2.873405 | -2.752077 | -2.164801 |
| H | 2.178621 | -0.037981 | 3.720449 |
| H | 3.162532 | -1.361203 | 4.387477 |
| H | 3.761222 | -0.415740 | 3.010594 |
| H | 7.733815 | -1.951308 | -2.688676 |
| H | 6.445383 | -0.733674 | -2.839065 |
| H | 7.721396 | -0.548206 | -1.609796 |
| H | 6.672185 | -3.118657 | 0.448174 |
| H | 7.320780 | 4.509327 | -0.110389 |
| H | 7.191460 | 5.906391 | 0.965938 |
| H | 7.842823 | 4.376191 | 1.573986 |
| H | 5.481116 | 4.666568 | 2.355318 |
| H | 5.617580 | 3.279759 | 1.283910 |
| H | 4.737034 | 4.641715 | -0.622195 |
| H | 4.645986 | 6.058666 | 0.423480 |
| H | 2.415103 | 5.118752 | 0.244265 |
| H | 2.979214 | 4.865742 | 1.882195 |
| H | -5.050942 | -5.653131 | 0.953983 |
| H | -4.154324 | -4.217871 | 1.468668 |
| H | -5.842383 | -4.415962 | 1.967741 |
| H | -7.139572 | 5.262671 | 1.249347 |
| H | -5.536316 | 4.833441 | 0.641713 |
| H | -6.536179 | 5.994684 | -0.257157 |
| H | -8.176700 | 4.222408 | -0.803915 |
| H | -6.584476 | 3.698917 | -1.338245 |

| | | | |
|---|---|---|---|
| H | -1.748256 | -0.876821 | -2.580064 |
| H | 6.105704 | -0.666735 | 0.487489 |
| H | 3.373237 | -1.304374 | 0.394732 |
| H | 4.982990 | -0.321778 | -0.840405 |
| H | 2.477769 | -3.646858 | 1.425464 |
| H | 3.315639 | -3.627160 | 2.987775 |
| H | -6.950458 | -0.465292 | 1.183080 |
| H | -8.473710 | 0.918755 | 2.787615 |
| H | -6.098576 | 1.016691 | -0.653783 |
| H | 3.255321 | 2.552468 | 1.433361 |
| H | 2.214655 | 0.710872 | 0.478661 |
| H | 1.338562 | 1.807382 | -2.839298 |
| H | 2.413586 | 4.094785 | -1.563790 |
| H | -4.219780 | -2.014817 | -1.953312 |
| H | 1.293232 | -2.237625 | 2.993617 |
| H | 2.162761 | 0.717367 | -1.200106 |
| H | 5.547024 | -2.711716 | -1.678394 |
| H | -4.888513 | -3.889242 | -0.913985 |
| H | -6.552199 | -4.260696 | -0.454085 |

-----------------------------------------------------------------------------------------------------------------

**TS(A-C)$_{QM1C}$**

-----------------------------------------------------------------------------------------------------------------

Number of imaginary frequencies: 1 Electronic energy: HF=-2024.065353
Zero-point correction= 0.719703 (Hartree/Particle)
Thermal correction to Energy= 0.768678
Thermal correction to Enthalpy= 0.769622
Thermal correction to Gibbs Free Energy= 0.624301
Sum of electronic and zero-point Energies= -2023.345650
Sum of electronic and thermal Energies= -2023.296675
Sum of electronic and thermal Enthalpies= -2023.295731
Sum of electronic and thermal Free Energies= -2023.441052

............................................................

Cartesian Coordinates

............................................................

| | | | |
|---|---|---|---|
| C | -5.154600 | -4.697328 | 0.178679 |
| C | -4.387573 | -3.489093 | -0.361271 |
| C | -5.195570 | -2.211745 | -0.393758 |
| O | -6.358987 | -2.153759 | 0.034318 |
| O | -4.624137 | -1.138553 | -0.894661 |
| C | 3.022586 | -3.145940 | 1.294461 |
| C | 4.263708 | -2.747543 | 0.524274 |
| O | 5.301097 | -3.444316 | 0.537059 |
| C | 2.325447 | -2.015619 | 2.085146 |
| O | 1.859146 | -0.994458 | 1.209445 |
| C | 3.214581 | -1.374148 | 3.151469 |
| N | 4.173237 | -1.599005 | -0.189589 |
| C | 5.315178 | -0.923914 | -0.794172 |
| C | 5.958832 | -1.650867 | -1.992032 |
| O | 6.800412 | -2.711985 | -1.580354 |
| C | 6.817874 | -0.689141 | -2.813940 |
| C | 6.908667 | 4.847303 | 1.096355 |
| C | 5.388163 | 5.031341 | 1.042412 |
| C | 4.790711 | 4.728026 | -0.337026 |
| C | 3.255085 | 4.867002 | -0.397433 |
| N | 2.518325 | 3.956027 | 0.458043 |
| C | 2.448233 | 2.581444 | 0.216127 |

| | | | |
|---|---|---|---|
| N | 1.989083 | 1.888401 | 1.322579 |
| N | 2.767278 | 1.985285 | -0.890047 |
| C | -6.744909 | 4.955548 | 0.611010 |
| C | -6.922720 | 4.095438 | -0.651925 |
| C | -7.284231 | 2.669742 | -0.350686 |
| N | -8.525699 | 2.329516 | 0.157665 |
| C | -6.502568 | 1.541440 | -0.479546 |
| C | -8.469459 | 1.016532 | 0.323777 |
| N | -7.273548 | 0.492247 | -0.045983 |
| C | -0.512746 | 0.374605 | -0.648806 |
| O | 0.377411 | 0.329320 | -1.668966 |
| O | -0.644450 | 1.378430 | 0.045086 |
| C | -1.209066 | -0.933056 | -0.378293 |
| O | -2.256691 | -1.138335 | -1.646356 |
| C | -0.277431 | -2.007476 | -0.180286 |
| C | -0.661778 | -3.235093 | 0.604998 |
| O | -1.790926 | -3.244207 | 1.197041 |
| O | 0.204513 | -4.165277 | 0.655750 |
| H | 0.977510 | -1.346977 | 0.731992 |
| H | 0.460248 | -2.150564 | -0.972568 |
| H | -1.930290 | -1.995998 | -1.957615 |
| H | 2.656634 | -0.602791 | 3.692395 |
| H | 3.567882 | -2.122336 | 3.871362 |
| H | 4.090144 | -0.897038 | 2.696321 |
| H | 7.336613 | -1.241088 | -3.604129 |
| H | 6.210913 | 0.097596 | -3.275659 |
| H | 7.578861 | -0.220264 | -2.178463 |
| H | 6.352561 | -3.119195 | -0.798130 |
| H | 7.186908 | 3.816917 | 0.850420 |
| H | 7.415766 | 5.506297 | 0.381103 |
| H | 7.304413 | 5.069725 | 2.093364 |
| H | 5.130970 | 6.060629 | 1.333699 |
| H | 4.923528 | 4.373272 | 1.786789 |
| H | 5.062985 | 3.706940 | -0.630543 |
| H | 5.233212 | 5.401377 | -1.085831 |
| H | 2.912218 | 4.723465 | -1.428928 |
| H | 2.965889 | 5.888749 | -0.119668 |
| H | -4.494906 | -5.569980 | 0.212855 |
| H | -5.510374 | -4.501291 | 1.194062 |
| H | -6.031536 | -4.939434 | -0.433545 |
| H | -7.656913 | 4.928479 | 1.215704 |
| H | -5.924938 | 4.572315 | 1.227006 |
| H | -6.525449 | 6.001502 | 0.360613 |
| H | -7.705369 | 4.545471 | -1.278975 |
| H | -5.997736 | 4.114407 | -1.240841 |
| H | -1.906141 | -0.828257 | 0.450723 |
| H | 6.109865 | -0.765654 | -0.048525 |
| H | 3.304961 | -1.076531 | -0.044586 |
| H | 4.943356 | 0.055785 | -1.109413 |
| H | 2.240059 | -3.573500 | 0.639664 |
| H | 3.324863 | -3.948678 | 1.975288 |
| H | -6.964294 | -0.501253 | -0.013172 |
| H | -9.273189 | 0.400258 | 0.705875 |
| H | -5.493148 | 1.377307 | -0.823763 |
| H | 2.549975 | 4.162528 | 1.446649 |
| H | 1.041764 | 2.168456 | 1.565259 |

| | | | |
|---|---|---|---|
| H | 1.043265 | 1.036287 | -1.514524 |
| H | 3.197307 | 2.632795 | -1.549919 |
| H | -3.602701 | -1.230457 | -1.199095 |
| H | 1.462013 | -2.488750 | 2.570428 |
| H | 1.991919 | 0.867160 | 1.193175 |
| H | 5.135869 | -2.021406 | -2.629728 |
| H | -3.463060 | -3.313882 | 0.233808 |
| H | -4.030741 | -3.679315 | -1.383718 |

---

**$A_{wt}$**

---

Number of imaginary frequencies: 5 Electronic energy: HF=-6984.290353
Zero-point correction= 1.770971 (Hartree/Particle)
Thermal correction to Energy= 1.896650
Thermal correction to Enthalpy= 1.897594
Thermal correction to Gibbs Free Energy= 1.573316
Sum of electronic and zero-point Energies= -6982.519382
Sum of electronic and thermal Energies= -6982.393703
Sum of electronic and thermal Enthalpies= -6982.392759
Sum of electronic and thermal Free Energies= -6982.717037

..........................................................

Cartesian Coordinates

..........................................................

| | | | |
|---|---|---|---|
| C | -3.113025 | -6.205941 | -3.261031 |
| C | -3.057823 | -5.501526 | -1.901156 |
| S | -2.662891 | -3.697639 | -2.080445 |
| C | 0.654044 | -5.786553 | -3.810158 |
| C | 1.856398 | -5.417902 | -4.690439 |
| C | 2.482632 | -4.067059 | -4.318668 |
| C | 1.702910 | -2.884176 | -4.893015 |
| O | 1.083134 | -2.943775 | -5.940113 |
| N | 1.851696 | -1.685909 | -4.197040 |
| C | 0.845951 | -5.019360 | -0.103169 |
| C | -0.231981 | -4.657159 | 0.922961 |
| C | 0.283425 | -4.061647 | 2.220424 |
| O | 1.423516 | -4.235695 | 2.646125 |
| O | -0.580175 | -3.360532 | 2.949977 |
| C | 5.585756 | -9.796092 | -0.945215 |
| C | 5.252930 | -8.302038 | -0.927434 |
| C | 6.458256 | -7.439065 | -0.533298 |
| C | 6.226738 | -5.932564 | -0.665703 |
| N | 5.268959 | -5.425899 | 0.319807 |
| C | 4.623068 | -4.241578 | 0.203378 |
| N | 4.766781 | -3.486087 | -0.888779 |
| N | 3.836283 | -3.801714 | 1.182744 |
| N | -0.349257 | 2.220370 | 3.940452 |
| C | -1.633959 | 1.831442 | 4.552714 |
| C | -8.103175 | 2.905972 | -0.302285 |
| C | -7.791565 | 1.867731 | -1.378488 |
| S | -6.003252 | 1.750543 | -1.866445 |
| C | -9.142483 | -0.944308 | 3.675745 |
| C | -7.642737 | -0.644228 | 3.748180 |
| S | -6.559724 | -2.115993 | 3.400507 |
| C | 8.972467 | 5.142103 | -2.394548 |
| C | 7.817730 | 4.133185 | -2.386098 |
| C | 8.293477 | 2.689035 | -2.606026 |

| | | | |
|---|---|---|---|
| C | 7.183645 | 1.632241 | -2.616563 |
| N | 6.544877 | 1.529779 | -1.305451 |
| C | 5.571734 | 0.659791 | -0.979103 |
| N | 5.144011 | -0.245898 | -1.858996 |
| N | 5.119215 | 0.670245 | 0.292116 |
| C | 1.223092 | 3.059540 | -2.364940 |
| C | 0.968846 | 4.528541 | -2.027760 |
| O | 1.213587 | 5.431530 | -2.831469 |
| C | 2.706630 | 2.638903 | -2.500530 |
| O | 3.236924 | 2.180704 | -1.266886 |
| C | 3.607436 | 3.685774 | -3.159170 |
| N | 0.423571 | 4.771527 | -0.805442 |
| C | 0.148262 | 6.127631 | -0.367258 |
| C | -0.955780 | 6.167902 | 0.697679 |
| O | -2.233686 | 5.780729 | 0.159686 |
| C | -1.055468 | 7.538388 | 1.365059 |
| C | 4.334709 | 9.745335 | 2.600278 |
| C | 4.439706 | 8.276954 | 3.040148 |
| C | 3.095860 | 7.533857 | 2.991980 |
| C | 3.125396 | 6.089138 | 3.525744 |
| N | 3.932931 | 5.169044 | 2.726730 |
| C | 3.479997 | 4.167251 | 1.897888 |
| N | 4.193747 | 3.677898 | 0.906586 |
| N | 2.225900 | 3.694720 | 2.137131 |
| C | -3.038843 | 7.359738 | -4.254631 |
| C | -3.238780 | 5.980540 | -4.901807 |
| C | -2.660304 | 4.797142 | -4.102696 |
| C | -3.410848 | 4.491308 | -2.796359 |
| C | -2.763798 | 3.345239 | -2.016905 |
| N | -3.386322 | 3.170880 | -0.661998 |
| C | 6.207834 | -2.760628 | 7.773262 |
| C | 5.927618 | -1.632033 | 6.775196 |
| C | 4.843189 | -1.914038 | 5.770450 |
| N | 4.622912 | -1.035776 | 4.724953 |
| C | 3.948418 | -2.960486 | 5.696913 |
| C | 3.615719 | -1.544532 | 4.042482 |
| N | 3.166453 | -2.710664 | 4.585210 |
| C | -0.596793 | 1.283827 | 0.715277 |
| O | 0.140096 | 2.295767 | 0.701823 |
| O | -1.867267 | 1.311884 | 0.843484 |
| C | 0.088640 | -0.093899 | 0.635461 |
| O | -0.886086 | -1.156013 | 0.624363 |
| C | 1.057077 | -0.185391 | -0.536174 |
| C | 2.223073 | -1.142990 | -0.350159 |
| O | 2.744750 | -1.250901 | 0.789949 |
| O | 2.698907 | -1.702902 | -1.402373 |
| Fe | -4.045601 | -2.651969 | -0.742634 |
| Fe | -5.548287 | -1.756816 | 1.554255 |
| Fe | -5.485297 | -0.107532 | -0.747684 |
| Fe | -2.813245 | -0.435735 | 0.969356 |
| S | -4.969675 | 0.365802 | 1.390492 |
| S | -3.337673 | -0.570713 | -1.257757 |
| S | -3.532680 | -2.646172 | 1.472944 |
| S | -6.318451 | -2.264738 | -0.365181 |
| H | 1.547738 | 0.788826 | -0.647275 |
| H | 0.523258 | -0.398717 | -1.468754 |

| | | | |
|---|---|---|---|
| H | -1.059575 | -1.453110 | -0.288305 |
| H | 4.603606 | 3.261169 | -3.317873 |
| H | 3.200180 | 4.017438 | -4.117373 |
| H | 3.703059 | 4.575121 | -2.530025 |
| H | -1.870843 | 7.544393 | 2.092739 |
| H | -0.126357 | 7.796312 | 1.882625 |
| H | -1.249882 | 8.322593 | 0.622138 |
| H | -2.524204 | 6.484190 | -0.437383 |
| H | -3.423135 | 8.154753 | -4.901631 |
| H | -2.764596 | 5.988274 | -5.890807 |
| H | -4.308616 | 5.809018 | -5.080191 |
| H | -1.602194 | 4.997834 | -3.882574 |
| H | -2.681950 | 3.901159 | -4.736427 |
| H | -4.455852 | 4.238636 | -3.009438 |
| H | -3.418559 | 5.386024 | -2.162036 |
| H | -1.705261 | 3.545177 | -1.845085 |
| H | -2.872424 | 2.389065 | -2.534328 |
| H | -1.975720 | 7.558520 | -4.075475 |
| H | -3.563916 | 7.443337 | -3.295938 |
| H | 3.965459 | 9.825733 | 1.571904 |
| H | 3.644744 | 10.300393 | 3.245200 |
| H | 5.308136 | 10.243219 | 2.645740 |
| H | 4.839455 | 8.230738 | 4.062521 |
| H | 5.171729 | 7.764782 | 2.399840 |
| H | 2.712957 | 7.526228 | 1.962619 |
| H | 2.364536 | 8.089077 | 3.595389 |
| H | 2.105897 | 5.702035 | 3.553985 |
| H | 3.494832 | 6.086257 | 4.559931 |
| H | 8.606374 | 6.161826 | -2.243001 |
| H | 9.697135 | 4.925524 | -1.601560 |
| H | 9.509837 | 5.120237 | -3.348935 |
| H | 7.090596 | 4.394010 | -3.165222 |
| H | 7.274789 | 4.211399 | -1.434722 |
| H | 9.035946 | 2.417236 | -1.842172 |
| H | 8.815347 | 2.623527 | -3.568010 |
| H | 7.612470 | 0.659586 | -2.893562 |
| H | 6.426574 | 1.886444 | -3.368963 |
| H | 4.713135 | -10.391790 | -1.228732 |
| H | 6.385758 | -10.014953 | -1.661157 |
| H | 5.918063 | -10.143312 | 0.039627 |
| H | 0.366783 | -5.491158 | -0.960555 |
| H | 1.372510 | -4.130842 | -0.460785 |
| H | 1.578696 | -5.715979 | 0.314703 |
| H | 5.321113 | -2.990839 | 8.374257 |
| H | 6.508327 | -3.680620 | 7.259509 |
| H | 7.011355 | -2.482057 | 8.461946 |
| H | 0.160507 | -6.689146 | -4.184028 |
| H | 0.974256 | -5.985809 | -2.784432 |
| H | -0.088992 | -4.983679 | -3.784860 |
| H | -3.314385 | -7.276731 | -3.130296 |
| H | -3.907134 | -5.781642 | -3.882847 |
| H | -2.171420 | -6.097820 | -3.803953 |
| H | -9.423168 | -1.745673 | 4.366679 |
| H | -9.716906 | -0.048082 | 3.940541 |
| H | -9.431001 | -1.253136 | 2.666332 |
| H | -7.568830 | 2.675736 | 0.624603 |

| | | | |
|---|---|---|---|
| H | -7.809715 | 3.910738 | -0.625702 |
| H | -9.178485 | 2.920572 | -0.084723 |
| H | 4.905112 | -7.993613 | -1.922441 |
| H | 4.418222 | -8.111625 | -0.242037 |
| H | 7.311007 | -7.688585 | -1.176926 |
| H | 6.780022 | -7.675503 | 0.491079 |
| H | 5.822245 | -5.728106 | -1.663053 |
| H | 7.182911 | -5.396938 | -0.575083 |
| H | 5.669265 | -0.716495 | 7.325326 |
| H | 6.846156 | -1.387480 | 6.225818 |
| H | 0.657403 | -0.219355 | 1.562647 |
| H | -0.128561 | 6.725348 | -1.242814 |
| H | 0.260335 | 3.993882 | -0.166209 |
| H | 1.055334 | 6.589763 | 0.050902 |
| H | 0.737010 | 2.400522 | -1.648191 |
| H | 0.744459 | 2.913193 | -3.340969 |
| H | 2.620414 | -6.201873 | -4.616320 |
| H | 1.551958 | -5.365380 | -5.740312 |
| H | 2.588358 | -3.959086 | -3.232560 |
| H | 3.495670 | -3.978593 | -4.736643 |
| H | -8.110927 | 0.872624 | -1.046150 |
| H | -8.343529 | 2.085694 | -2.298635 |
| H | -7.366770 | 0.150002 | 3.052335 |
| H | -7.369748 | -0.311099 | 4.755554 |
| H | -4.015820 | -5.604466 | -1.380945 |
| H | -2.291193 | -5.960442 | -1.266112 |
| H | 2.414892 | -3.280656 | 4.201630 |
| H | 3.180372 | -1.129944 | 3.143332 |
| H | 3.792938 | -3.828067 | 6.319409 |
| H | 6.956010 | 2.037087 | -0.536333 |
| H | 4.377274 | 0.008280 | 0.519787 |
| H | 4.240748 | -0.694471 | -1.685263 |
| H | 5.412307 | -0.254902 | -2.840611 |
| H | 4.898444 | 5.426585 | 2.589211 |
| H | 4.991792 | 4.286231 | 0.725518 |
| H | 1.845479 | 3.000511 | 1.501468 |
| H | 1.848424 | 3.658197 | 3.074917 |
| H | 5.344927 | -5.789104 | 1.260375 |
| H | 4.035913 | -2.777204 | -1.096259 |
| H | 5.484238 | -3.674373 | -1.567693 |
| H | 3.441229 | -2.850777 | 1.088433 |
| H | 1.195294 | -0.976930 | -4.506662 |
| H | 1.995673 | -1.727398 | -3.182467 |
| H | -2.884435 | 2.444337 | -0.106267 |
| H | -3.308360 | 4.054205 | -0.142386 |
| H | -2.309861 | 2.692308 | 4.548502 |
| H | -2.152760 | 1.000744 | 4.051944 |
| H | 0.273584 | 1.414767 | 3.946445 |
| H | -0.505448 | 2.440068 | 2.959741 |
| H | -1.467978 | 1.550058 | 5.597209 |
| H | -1.446536 | -3.229390 | 2.499505 |
| H | 2.707272 | 1.743411 | -3.137322 |
| H | -4.381265 | 2.861700 | -0.770601 |
| H | 3.437736 | 2.903485 | -0.623993 |
| H | -0.741610 | 5.407516 | 1.457123 |
| H | -0.971867 | -3.985628 | 0.475857 |

| | | | |
|---|---|---|---|
| H | -0.796734 | -5.555599 | 1.210529 |
| H | 4.960896 | 1.599231 | 0.697093 |
| H | 3.313464 | -4.423474 | 1.791215 |
| O | 4.682707 | -1.114042 | -4.619343 |
| H | 4.814106 | -0.815122 | -5.526811 |
| H | 3.709833 | -1.215713 | -4.530504 |

--------------------------------------------------------------------------------------------------------

**$B_{wt}$**

--------------------------------------------------------------------------------------------------------

Number of imaginary frequencies: 5 Electronic energy: HF=-6984.263417
Zero-point correction= 1.767775 (Hartree/Particle)
Thermal correction to Energy= 1.892553
Thermal correction to Enthalpy= 1.893498
Thermal correction to Gibbs Free Energy= 1.572352
Sum of electronic and zero-point Energies= -6982.495642
Sum of electronic and thermal Energies= -6982.370864
Sum of electronic and thermal Enthalpies= -6982.369919
Sum of electronic and thermal Free Energies= -6982.691065

............................................................

Cartesian Coordinates

............................................................

| | | | |
|---|---|---|---|
| C | -3.207777 | -6.319516 | -3.140594 |
| C | -3.161848 | -5.587289 | -1.794321 |
| S | -2.738036 | -3.793882 | -2.010185 |
| C | 0.537157 | -5.902224 | -3.825757 |
| C | 1.766593 | -5.472115 | -4.639634 |
| C | 2.370812 | -4.133627 | -4.188976 |
| C | 1.584956 | -2.930345 | -4.719552 |
| O | 0.934244 | -2.979391 | -5.751387 |
| N | 1.740341 | -1.752706 | -4.007858 |
| C | 0.853673 | -5.068549 | -0.141726 |
| C | -0.183969 | -4.699189 | 0.924260 |
| C | 0.370248 | -4.095087 | 2.203204 |
| O | 1.525449 | -4.256661 | 2.593652 |
| O | -0.476472 | -3.408138 | 2.963652 |
| C | 5.574697 | -9.849721 | -1.060515 |
| C | 5.111723 | -8.388658 | -1.079469 |
| C | 6.264710 | -7.409666 | -0.820466 |
| C | 5.919661 | -5.932416 | -1.022294 |
| N | 5.009694 | -5.418261 | 0.003325 |
| C | 4.494025 | -4.159193 | -0.043047 |
| N | 4.678225 | -3.386482 | -1.113581 |
| N | 3.806335 | -3.672932 | 0.987257 |
| N | -0.328149 | 2.366007 | 3.705683 |
| C | -1.483878 | 1.859166 | 4.473556 |
| C | -8.118277 | 2.833416 | -0.175194 |
| C | -7.851063 | 1.743650 | -1.212884 |
| S | -6.095029 | 1.630035 | -1.809959 |
| C | -9.010628 | -0.947359 | 3.903897 |
| C | -7.521456 | -0.588161 | 3.881830 |
| S | -6.399853 | -2.043089 | 3.586734 |
| C | 8.869793 | 5.067211 | -2.890292 |
| C | 7.914047 | 3.921202 | -2.515808 |
| C | 8.641879 | 2.669338 | -1.997993 |
| C | 7.774286 | 1.411388 | -1.845449 |
| N | 6.846746 | 1.480354 | -0.706108 |

| | | | |
|---|---|---|---|
| C | 5.792297 | 0.620241 | -0.551941 |
| N | 5.268683 | -0.001383 | -1.598615 |
| N | 5.275161 | 0.400985 | 0.659327 |
| C | 1.131490 | 2.969458 | -2.558228 |
| C | 0.979575 | 4.445404 | -2.193732 |
| O | 1.418356 | 5.343593 | -2.915238 |
| C | 2.552089 | 2.393428 | -2.357927 |
| O | 2.820318 | 2.152605 | -0.971747 |
| C | 3.670554 | 3.260991 | -2.924281 |
| N | 0.316349 | 4.707443 | -1.033162 |
| C | 0.117459 | 6.070450 | -0.579921 |
| C | -1.013124 | 6.163890 | 0.453040 |
| O | -2.276748 | 5.766133 | -0.100042 |
| C | -1.114366 | 7.561258 | 1.063555 |
| C | 4.393219 | 9.749318 | 2.177789 |
| C | 4.351527 | 8.344389 | 2.797335 |
| C | 2.929613 | 7.769758 | 2.910562 |
| C | 2.847773 | 6.355416 | 3.515105 |
| N | 3.542514 | 5.353848 | 2.698982 |
| C | 2.965401 | 4.416800 | 1.903988 |
| N | 3.630542 | 4.012638 | 0.815213 |
| N | 1.783962 | 3.883503 | 2.197294 |
| C | -3.204009 | 7.226367 | -4.377177 |
| C | -3.482744 | 5.858398 | -5.019230 |
| C | -2.897937 | 4.654638 | -4.256354 |
| C | -3.601243 | 4.348047 | -2.924431 |
| C | -2.896287 | 3.234893 | -2.147476 |
| N | -3.456433 | 3.079100 | -0.764673 |
| C | 6.475670 | -2.658404 | 7.505109 |
| C | 6.303872 | -1.624361 | 6.384220 |
| C | 5.146258 | -1.872569 | 5.454074 |
| N | 4.883657 | -1.006457 | 4.404254 |
| C | 4.216439 | -2.889649 | 5.461615 |
| C | 3.814136 | -1.497608 | 3.802505 |
| N | 3.369354 | -2.636008 | 4.400927 |
| C | -0.528428 | 1.333539 | 0.631691 |
| O | 0.152662 | 2.396245 | 0.589917 |
| O | -1.799724 | 1.339711 | 0.732950 |
| C | 0.217694 | -0.008903 | 0.620134 |
| O | -0.802844 | -1.076350 | 0.743157 |
| C | 1.120863 | -0.222784 | -0.533320 |
| C | 2.330075 | -0.930109 | -0.393190 |
| O | 2.944090 | -1.069270 | 0.749199 |
| O | 2.936581 | -1.396483 | -1.475886 |
| Fe | -4.027204 | -2.700544 | -0.625489 |
| Fe | -5.437098 | -1.738780 | 1.698939 |
| Fe | -5.469781 | -0.182700 | -0.655643 |
| Fe | -2.715116 | -0.415549 | 0.966051 |
| S | -4.884514 | 0.383088 | 1.440223 |
| S | -3.339724 | -0.638149 | -1.226524 |
| S | -3.414643 | -2.609870 | 1.559724 |
| S | -6.275315 | -2.329513 | -0.168022 |
| H | 2.206373 | 1.413924 | -0.682139 |
| H | 0.658286 | -0.214259 | -1.518796 |
| H | -0.805828 | -1.562185 | -0.098779 |
| H | 4.631505 | 2.759839 | -2.778496 |

| | | | |
|---|---|---|---|
| H | 3.519477 | 3.434015 | -3.993181 |
| H | 3.699441 | 4.243884 | -2.446203 |
| H | -1.944559 | 7.602033 | 1.773025 |
| H | -0.194266 | 7.835483 | 1.589982 |
| H | -1.288643 | 8.315827 | 0.286374 |
| H | -2.511355 | 6.403043 | -0.789530 |
| H | -3.603331 | 8.037656 | -4.994111 |
| H | -3.064064 | 5.858363 | -6.032925 |
| H | -4.565466 | 5.721156 | -5.138712 |
| H | -1.828219 | 4.834483 | -4.075333 |
| H | -2.956910 | 3.765477 | -4.896484 |
| H | -4.643972 | 4.062049 | -3.101763 |
| H | -3.621074 | 5.252451 | -2.303396 |
| H | -1.835957 | 3.463368 | -2.029262 |
| H | -3.003129 | 2.263758 | -2.636640 |
| H | -2.126545 | 7.394071 | -4.262145 |
| H | -3.668075 | 7.317914 | -3.387949 |
| H | 3.976288 | 9.748153 | 1.164938 |
| H | 3.813859 | 10.461119 | 2.775434 |
| H | 5.419944 | 10.121874 | 2.117503 |
| H | 4.812201 | 8.367262 | 3.794058 |
| H | 4.977765 | 7.679026 | 2.185124 |
| H | 2.455081 | 7.759662 | 1.920197 |
| H | 2.323690 | 8.433347 | 3.540810 |
| H | 1.803805 | 6.049542 | 3.607195 |
| H | 3.276869 | 6.344996 | 4.523721 |
| H | 8.315144 | 5.937056 | -3.254342 |
| H | 9.468175 | 5.387889 | -2.029853 |
| H | 9.563628 | 4.758532 | -3.679676 |
| H | 7.322241 | 3.645991 | -3.398696 |
| H | 7.190836 | 4.266066 | -1.766277 |
| H | 9.149027 | 2.889323 | -1.046993 |
| H | 9.442165 | 2.408022 | -2.702044 |
| H | 8.420987 | 0.527749 | -1.749255 |
| H | 7.189757 | 1.280269 | -2.759527 |
| H | 4.739275 | -10.531206 | -1.246076 |
| H | 6.332924 | -10.033706 | -1.829812 |
| H | 6.012571 | -10.117702 | -0.092256 |
| H | 0.338741 | -5.528372 | -0.984493 |
| H | 1.374601 | -4.180436 | -0.509079 |
| H | 1.595348 | -5.773911 | 0.244918 |
| H | 5.588670 | -2.699113 | 8.146398 |
| H | 6.638340 | -3.662649 | 7.098372 |
| H | 7.334205 | -2.409838 | 8.136384 |
| H | 0.060643 | -6.776869 | -4.280158 |
| H | 0.819806 | -6.174303 | -2.805539 |
| H | -0.208039 | -5.103009 | -3.773250 |
| H | -3.433762 | -7.383035 | -2.991361 |
| H | -3.981594 | -5.892277 | -3.785660 |
| H | -2.254340 | -6.242078 | -3.667709 |
| H | -9.227117 | -1.704289 | 4.664622 |
| H | -9.608555 | -0.055550 | 4.129203 |
| H | -9.331348 | -1.342193 | 2.935026 |
| H | -7.524212 | 2.661506 | 0.727764 |
| H | -7.865594 | 3.824400 | -0.568416 |
| H | -9.178606 | 2.840829 | 0.107026 |

| | | | |
|---|---|---|---|
| H | 4.663791 | -8.160366 | -2.055614 |
| H | 4.316906 | -8.237192 | -0.339104 |
| H | 7.089549 | -7.637414 | -1.507255 |
| H | 6.671778 | -7.557719 | 0.190254 |
| H | 5.435311 | -5.813480 | -1.999135 |
| H | 6.848115 | -5.341234 | -1.037214 |
| H | 6.189455 | -0.624728 | 6.825331 |
| H | 7.227176 | -1.572752 | 5.790275 |
| H | 0.808054 | -0.061977 | 1.541818 |
| H | -0.097686 | 6.701577 | -1.450630 |
| H | 0.068034 | 3.929616 | -0.418061 |
| H | 1.038255 | 6.478098 | -0.132279 |
| H | 0.429003 | 2.341078 | -2.008398 |
| H | 0.887227 | 2.901764 | -3.624440 |
| H | 2.536694 | -6.251975 | -4.579860 |
| H | 1.494841 | -5.369246 | -5.694627 |
| H | 2.460217 | -4.075399 | -3.097674 |
| H | 3.390260 | -4.028968 | -4.588785 |
| H | -8.127414 | 0.762914 | -0.808158 |
| H | -8.464674 | 1.902201 | -2.105789 |
| H | -7.310225 | 0.161197 | 3.116751 |
| H | -7.217787 | -0.168889 | 4.847463 |
| H | -4.127970 | -5.666624 | -1.285241 |
| H | -2.409537 | -6.043538 | -1.140384 |
| H | 2.591812 | -3.204225 | 4.067841 |
| H | 3.327091 | -1.093050 | 2.924570 |
| H | 4.077372 | -3.739331 | 6.111785 |
| H | 7.259648 | 1.803014 | 0.158913 |
| H | 4.401715 | -0.176259 | 0.747550 |
| H | 4.376987 | -0.537770 | -1.490443 |
| H | 5.406788 | 0.291936 | -2.558175 |
| H | 4.504960 | 5.563031 | 2.476899 |
| H | 4.312810 | 4.640008 | 0.416813 |
| H | 1.328192 | 3.236837 | 1.532073 |
| H | 1.386134 | 3.865730 | 3.133990 |
| H | 5.099941 | -5.812140 | 0.929965 |
| H | 4.013683 | -2.571690 | -1.258153 |
| H | 5.284305 | -3.682202 | -1.859767 |
| H | 3.469461 | -2.683053 | 0.921967 |
| H | 1.064109 | -1.046003 | -4.274838 |
| H | 1.992991 | -1.762429 | -3.007665 |
| H | -2.905154 | 2.387752 | -0.210894 |
| H | -3.394983 | 3.978102 | -0.273388 |
| H | -2.222755 | 2.658562 | 4.582807 |
| H | -1.988047 | 0.999362 | 4.012403 |
| H | 0.337454 | 1.601357 | 3.598895 |
| H | -0.650522 | 2.561833 | 2.760449 |
| H | -1.158150 | 1.570189 | 5.477175 |
| H | -1.352512 | -3.263220 | 2.534868 |
| H | 2.583765 | 1.424482 | -2.872464 |
| H | -4.446917 | 2.733050 | -0.824700 |
| H | 3.257574 | 3.272886 | 0.172927 |
| H | -0.826881 | 5.431374 | 1.246767 |
| H | -0.941597 | -4.032228 | 0.499583 |
| H | -0.739443 | -5.595139 | 1.237150 |
| H | 5.764835 | 0.657228 | 1.503014 |

| | | | |
|---|---|---|---|
| H | 3.264016 | -4.275400 | 1.601096 |
| O | 4.329017 | -0.309910 | -4.159114 |
| H | 4.276766 | 0.030877 | -5.060008 |
| H | 3.509404 | -0.837387 | -4.060687 |

---

**$C_{wt}$**

---

Number of imaginary frequencies: 6 Electronic energy: HF=-6984.273194
Zero-point correction= 1.768011 (Hartree/Particle)
Thermal correction to Energy= 1.893014
Thermal correction to Enthalpy= 1.893958
Thermal correction to Gibbs Free Energy= 1.573208
Sum of electronic and zero-point Energies= -6982.505183
Sum of electronic and thermal Energies= -6982.380180
Sum of electronic and thermal Enthalpies= -6982.379236
Sum of electronic and thermal Free Energies= -6982.699986

............................................................

Cartesian Coordinates

............................................................

| | | | |
|---|---|---|---|
| C | -2.762313 | -6.817271 | -2.558596 |
| C | -2.982750 | -5.917646 | -1.337598 |
| S | -2.645080 | -4.140910 | -1.744503 |
| C | 0.946300 | -6.195441 | -3.284701 |
| C | 2.131722 | -5.753025 | -4.153277 |
| C | 2.494899 | -4.268846 | -3.985120 |
| C | 1.469127 | -3.347310 | -4.656892 |
| O | 0.933951 | -3.643576 | -5.714025 |
| N | 1.251763 | -2.139852 | -4.032830 |
| C | 1.201797 | -5.089046 | 0.334989 |
| C | 0.745459 | -4.877905 | 1.789631 |
| C | 1.446829 | -3.661695 | 2.397949 |
| O | 2.587248 | -3.880615 | 2.952707 |
| O | 0.920477 | -2.531962 | 2.263045 |
| C | 6.237640 | -9.591257 | -0.272282 |
| C | 5.562020 | -8.230305 | -0.464274 |
| C | 6.586789 | -7.113122 | -0.699082 |
| C | 5.996527 | -5.758680 | -1.092033 |
| N | 5.275175 | -5.118268 | 0.010095 |
| C | 4.783453 | -3.861762 | -0.059207 |
| N | 4.868110 | -3.186499 | -1.227984 |
| N | 4.274071 | -3.268519 | 1.007992 |
| N | -0.295909 | 2.635782 | 4.696389 |
| C | -1.605577 | 1.966609 | 4.466080 |
| C | -8.286110 | 2.164634 | -0.214385 |
| C | -7.942036 | 1.260395 | -1.395411 |
| S | -6.148441 | 1.237016 | -1.875421 |
| C | -8.923616 | -1.376682 | 4.119461 |
| C | -7.500067 | -0.844557 | 3.937514 |
| S | -6.228192 | -2.180817 | 3.712625 |
| C | 8.281200 | 5.170527 | -3.682712 |
| C | 7.612219 | 3.832669 | -3.352178 |
| C | 8.624162 | 2.697524 | -3.154468 |
| C | 7.998972 | 1.333283 | -2.850197 |
| N | 7.324805 | 1.313564 | -1.544890 |
| C | 6.209814 | 0.609808 | -1.237650 |
| N | 5.365385 | 0.208858 | -2.194235 |

| | | | |
|---|---|---|---|
| N | 5.941773 | 0.301249 | 0.030130 |
| C | 0.935009 | 2.763020 | -2.635261 |
| C | 0.439780 | 4.218840 | -2.524730 |
| O | 0.667163 | 5.051067 | -3.401400 |
| C | 2.316491 | 2.677960 | -1.919702 |
| O | 2.225416 | 3.051646 | -0.536099 |
| C | 3.384290 | 3.562187 | -2.552649 |
| N | -0.162155 | 4.543723 | -1.338301 |
| C | -0.288845 | 5.917061 | -0.872433 |
| C | -1.373043 | 6.038162 | 0.210556 |
| O | -2.642151 | 5.536876 | -0.230533 |
| C | -1.512564 | 7.473221 | 0.716358 |
| C | 3.724344 | 10.059500 | 1.608650 |
| C | 3.562970 | 8.901321 | 2.613178 |
| C | 2.139261 | 8.315070 | 2.670428 |
| C | 1.938525 | 7.146092 | 3.660981 |
| N | 2.733732 | 5.948691 | 3.324329 |
| C | 2.309959 | 4.993486 | 2.436739 |
| N | 3.126751 | 4.682012 | 1.423756 |
| N | 1.141323 | 4.380320 | 2.596017 |
| C | -3.677497 | 6.579565 | -4.728041 |
| C | -3.934689 | 5.165099 | -5.270565 |
| C | -3.238175 | 4.031798 | -4.493882 |
| C | -3.755031 | 3.849911 | -3.059009 |
| C | -3.049736 | 2.708390 | -2.326602 |
| N | -3.449881 | 2.656044 | -0.877933 |
| C | 6.639589 | -1.780053 | 7.772685 |
| C | 6.576624 | -0.726371 | 6.659107 |
| C | 5.576347 | -1.026365 | 5.575809 |
| N | 5.370131 | -0.151834 | 4.523278 |
| C | 4.747001 | -2.120387 | 5.439536 |
| C | 4.429733 | -0.722577 | 3.784115 |
| N | 4.016872 | -1.911726 | 4.292439 |
| C | -0.413133 | 1.360144 | 0.756490 |
| O | -0.121447 | 2.614627 | 0.792015 |
| O | -1.584078 | 0.930931 | 0.641765 |
| C | 0.817757 | 0.506884 | 0.760240 |
| O | -0.918752 | -1.986225 | 0.373589 |
| C | 1.189874 | -0.205027 | -0.310376 |
| C | 2.587544 | -0.671922 | -0.471582 |
| O | 3.405359 | -0.580168 | 0.491266 |
| O | 2.935405 | -1.107485 | -1.635382 |
| Fe | -4.155015 | -2.963238 | -0.618236 |
| Fe | -5.466392 | -1.788953 | 1.759080 |
| Fe | -5.427708 | -0.446988 | -0.616715 |
| Fe | -2.571728 | -0.874806 | 0.853566 |
| S | -4.658651 | 0.222325 | 1.384907 |
| S | -3.364406 | -0.961982 | -1.306466 |
| S | -3.572290 | -2.890205 | 1.578355 |
| S | -6.407437 | -2.429461 | -0.050470 |
| H | 1.411776 | 2.688283 | -0.125021 |
| H | 0.504860 | -0.389626 | -1.129426 |
| H | -1.301077 | -2.782596 | -0.046358 |
| H | 4.320735 | 3.464888 | -1.990874 |
| H | 3.561378 | 3.249146 | -3.582367 |
| H | 3.079881 | 4.611772 | -2.556284 |

| | | | |
|---|---|---|---|
| H | -2.285476 | 7.527584 | 1.487119 |
| H | -0.572296 | 7.841611 | 1.139454 |
| H | -1.793475 | 8.149525 | -0.100516 |
| H | -2.970435 | 6.114862 | -0.933623 |
| H | -4.137693 | 7.335289 | -5.372723 |
| H | -3.595096 | 5.123260 | -6.312771 |
| H | -5.015512 | 4.973580 | -5.295794 |
| H | -2.156002 | 4.223217 | -4.472477 |
| H | -3.380171 | 3.092143 | -5.042804 |
| H | -4.833264 | 3.655087 | -3.065550 |
| H | -3.589624 | 4.776207 | -2.497238 |
| H | -1.969089 | 2.846540 | -2.348710 |
| H | -3.294779 | 1.731187 | -2.747880 |
| H | -2.602981 | 6.791834 | -4.676494 |
| H | -4.096563 | 6.715153 | -3.724215 |
| H | 3.485795 | 9.737590 | 0.589204 |
| H | 3.059490 | 10.892810 | 1.858841 |
| H | 4.750866 | 10.436851 | 1.611721 |
| H | 3.849156 | 9.245945 | 3.615800 |
| H | 4.282452 | 8.111984 | 2.348566 |
| H | 1.835727 | 7.986373 | 1.666841 |
| H | 1.437779 | 9.109326 | 2.957217 |
| H | 0.887580 | 6.850263 | 3.686120 |
| H | 2.213041 | 7.451880 | 4.675903 |
| H | 7.536729 | 5.959584 | -3.824114 |
| H | 8.955888 | 5.488070 | -2.879509 |
| H | 8.871755 | 5.102547 | -4.602861 |
| H | 6.920116 | 3.559667 | -4.159931 |
| H | 6.998430 | 3.937559 | -2.448916 |
| H | 9.331189 | 2.959648 | -2.353681 |
| H | 9.232476 | 2.581775 | -4.060199 |
| H | 8.770005 | 0.551523 | -2.870730 |
| H | 7.268163 | 1.077418 | -3.619559 |
| H | 5.498235 | -10.379448 | -0.103382 |
| H | 6.824807 | -9.867893 | -1.155000 |
| H | 6.915595 | -9.582482 | 0.588637 |
| H | 0.653157 | -5.918452 | -0.119517 |
| H | 1.009843 | -4.197420 | -0.269686 |
| H | 2.270467 | -5.325675 | 0.284795 |
| H | 5.670530 | -1.885787 | 8.271845 |
| H | 6.916132 | -2.762339 | 7.374123 |
| H | 7.380020 | -1.506303 | 8.530885 |
| H | 0.685321 | -7.239181 | -3.486820 |
| H | 1.177979 | -6.106064 | -2.220270 |
| H | 0.062984 | -5.583401 | -3.486423 |
| H | -2.965044 | -7.866308 | -2.306177 |
| H | -3.427190 | -6.527092 | -3.378042 |
| H | -1.735185 | -6.747737 | -2.922423 |
| H | -8.987056 | -2.055395 | 4.976088 |
| H | -9.620305 | -0.546185 | 4.288913 |
| H | -9.250373 | -1.925061 | 3.230723 |
| H | -7.745486 | 1.850444 | 0.683653 |
| H | -8.019591 | 3.207206 | -0.422114 |
| H | -9.361949 | 2.124534 | -0.001589 |
| H | 4.879227 | -8.275899 | -1.322563 |
| H | 4.936999 | -7.999640 | 0.407691 |

| | | | |
|---|---|---|---|
| H | 7.257126 | -7.411665 | -1.514427 |
| H | 7.227595 | -6.988511 | 0.185023 |
| H | 5.311662 | -5.898848 | -1.939807 |
| H | 6.809918 | -5.097926 | -1.423625 |
| H | 6.343255 | 0.254291 | 7.096231 |
| H | 7.572762 | -0.611005 | 6.208290 |
| H | 1.530298 | 0.670712 | 1.563821 |
| H | -0.521356 | 6.553157 | -1.733707 |
| H | -0.255019 | 3.823004 | -0.624820 |
| H | 0.665627 | 6.281614 | -0.458743 |
| H | 0.242969 | 2.051749 | -2.173136 |
| H | 1.063152 | 2.506518 | -3.686394 |
| H | 3.013760 | -6.364906 | -3.918915 |
| H | 1.901043 | -5.916265 | -5.210757 |
| H | 2.610040 | -4.009683 | -2.925755 |
| H | 3.458923 | -4.067200 | -4.475280 |
| H | -8.232208 | 0.227533 | -1.179953 |
| H | -8.487250 | 1.572278 | -2.292882 |
| H | -7.445962 | -0.171322 | 3.077354 |
| H | -7.189070 | -0.271802 | 4.818295 |
| H | -4.013981 | -6.010235 | -0.981512 |
| H | -2.326430 | -6.220589 | -0.514614 |
| H | 3.346190 | -2.573134 | 3.849336 |
| H | 4.005129 | -0.322000 | 2.872667 |
| H | 4.608534 | -3.002988 | 6.044743 |
| H | 7.883790 | 1.612813 | -0.757548 |
| H | 4.996747 | -0.069950 | 0.272780 |
| H | 4.526065 | -0.324475 | -1.918552 |
| H | 5.231357 | 0.735015 | -3.049366 |
| H | 3.725675 | 6.135513 | 3.249536 |
| H | 3.810293 | 5.371301 | 1.149014 |
| H | 0.797018 | 3.710013 | 1.891492 |
| H | 0.704403 | 4.224942 | 3.513381 |
| H | 5.349314 | -5.520502 | 0.933693 |
| H | 4.189264 | -2.427218 | -1.384939 |
| H | 5.152702 | -3.690820 | -2.052087 |
| H | 3.939671 | -2.307431 | 0.890288 |
| H | 0.414037 | -1.657028 | -4.333375 |
| H | 1.515129 | -2.020206 | -3.060834 |
| H | -2.882095 | 1.970828 | -0.341999 |
| H | -3.321671 | 3.581646 | -0.446327 |
| H | -1.676991 | 1.645065 | 3.425129 |
| H | -1.772223 | 1.092684 | 5.108141 |
| H | -0.242938 | 2.900624 | 5.679212 |
| H | 0.436710 | 1.935545 | 4.582990 |
| H | -2.411021 | 2.684178 | 4.643450 |
| H | -0.327491 | -2.271976 | 1.128193 |
| H | 2.639630 | 1.639286 | -1.967585 |
| H | -4.445829 | 2.342949 | -0.818041 |
| H | 2.880660 | 3.976198 | 0.702720 |
| H | -1.108915 | 5.380335 | 1.045395 |
| H | -0.337005 | -4.720634 | 1.815222 |
| H | 0.980324 | -5.764838 | 2.385834 |
| H | 6.635649 | 0.392682 | 0.756434 |
| H | 3.796883 | -3.761064 | 1.802434 |
| O | 3.164791 | 0.781106 | -3.980673 |

| | | | |
|---|---|---|---|
| H | 2.827057 | 0.071835 | -3.403302 |
| H | 2.936408 | 0.473821 | -4.868370 |

---

**TS(A-B)$_{wt}$**

---

Number of imaginary frequencies: 6 Electronic energy: HF=-6984.255081
Zero-point correction= 1.763427 (Hartree/Particle)
Thermal correction to Energy= 1.887803
Thermal correction to Enthalpy= 1.888747
Thermal correction to Gibbs Free Energy= 1.569249
Sum of electronic and zero-point Energies= -6982.491654
Sum of electronic and thermal Energies= -6982.367278
Sum of electronic and thermal Enthalpies= -6982.366334
Sum of electronic and thermal Free Energies= -6982.685832

..........................................................

Cartesian Coordinates

..........................................................

| | | | |
|---|---|---|---|
| C | -3.293417 | -6.227470 | -3.202953 |
| C | -3.186091 | -5.513551 | -1.851613 |
| S | -2.749892 | -3.723331 | -2.062168 |
| C | 0.455212 | -5.859898 | -3.896383 |
| C | 1.696331 | -5.423076 | -4.689222 |
| C | 2.311049 | -4.101131 | -4.203608 |
| C | 1.568817 | -2.874805 | -4.742597 |
| O | 0.941111 | -2.894929 | -5.787701 |
| N | 1.743314 | -1.701353 | -4.017362 |
| C | 0.796146 | -5.069081 | -0.205056 |
| C | -0.247604 | -4.691106 | 0.851835 |
| C | 0.299676 | -4.142630 | 2.158469 |
| O | 1.442459 | -4.347824 | 2.566522 |
| O | -0.540683 | -3.452315 | 2.921327 |
| C | 5.441091 | -9.911505 | -1.188682 |
| C | 5.010071 | -8.441443 | -1.143844 |
| C | 6.182009 | -7.500900 | -0.832589 |
| C | 5.873924 | -6.009872 | -0.986267 |
| N | 4.968691 | -5.505781 | 0.049584 |
| C | 4.443025 | -4.253702 | 0.015156 |
| N | 4.618184 | -3.468814 | -1.049041 |
| N | 3.752584 | -3.779357 | 1.050198 |
| N | -0.180050 | 2.317473 | 3.844751 |
| C | -1.421246 | 1.845313 | 4.489057 |
| C | -8.054880 | 2.967757 | -0.127371 |
| C | -7.816160 | 1.876643 | -1.170677 |
| S | -6.066872 | 1.737992 | -1.781173 |
| C | -8.991434 | -0.840887 | 3.915727 |
| C | -7.488667 | -0.544843 | 3.883696 |
| S | -6.436834 | -2.038443 | 3.531176 |
| C | 8.956346 | 4.971875 | -2.876474 |
| C | 7.900606 | 3.925108 | -2.486339 |
| C | 8.499015 | 2.528405 | -2.250698 |
| C | 7.484873 | 1.384723 | -2.125722 |
| N | 6.730710 | 1.418766 | -0.863018 |
| C | 5.714727 | 0.546075 | -0.606050 |
| N | 5.163477 | -0.147878 | -1.591435 |
| N | 5.298769 | 0.344243 | 0.649041 |
| C | 1.188235 | 2.988269 | -2.540033 |

| | | | |
|---|---|---|---|
| C | 1.055514 | 4.462794 | -2.168246 |
| O | 1.507455 | 5.360949 | -2.882505 |
| C | 2.575157 | 2.359754 | -2.239852 |
| O | 2.738917 | 2.078339 | -0.866342 |
| C | 3.738999 | 3.205382 | -2.764770 |
| N | 0.395748 | 4.725982 | -1.005304 |
| C | 0.227714 | 6.083681 | -0.526567 |
| C | -0.907790 | 6.182735 | 0.500628 |
| O | -2.179734 | 5.844153 | -0.074602 |
| C | -0.969062 | 7.563469 | 1.152835 |
| C | 4.567610 | 9.668769 | 2.254360 |
| C | 4.577368 | 8.227529 | 2.784024 |
| C | 3.169472 | 7.633353 | 2.948654 |
| C | 3.131970 | 6.196689 | 3.500180 |
| N | 3.772859 | 5.225249 | 2.609056 |
| C | 3.147305 | 4.289537 | 1.844848 |
| N | 3.741723 | 3.848869 | 0.735017 |
| N | 1.967340 | 3.787342 | 2.206453 |
| C | -3.087997 | 7.328845 | -4.300567 |
| C | -3.362274 | 5.970095 | -4.963197 |
| C | -2.805169 | 4.756093 | -4.196553 |
| C | -3.537970 | 4.452017 | -2.880233 |
| C | -2.853707 | 3.337471 | -2.087555 |
| N | -3.440327 | 3.192860 | -0.714802 |
| C | 6.478551 | -2.823284 | 7.447143 |
| C | 6.186365 | -1.710130 | 6.433833 |
| C | 5.025092 | -1.969656 | 5.512131 |
| N | 4.713935 | -1.070323 | 4.505972 |
| C | 4.135130 | -3.021827 | 5.488353 |
| C | 3.655800 | -1.575057 | 3.897585 |
| N | 3.262909 | -2.755223 | 4.450805 |
| C | -0.583538 | 1.427928 | 0.698811 |
| O | 0.066706 | 2.506120 | 0.724617 |
| O | -1.856511 | 1.391511 | 0.755312 |
| C | 0.192650 | 0.102660 | 0.667707 |
| O | -0.783025 | -0.996997 | 0.687782 |
| C | 1.174909 | -0.016303 | -0.461425 |
| C | 2.266288 | -0.964915 | -0.298054 |
| O | 2.847163 | -1.130290 | 0.833487 |
| O | 2.762509 | -1.531670 | -1.364766 |
| Fe | -4.035100 | -2.644486 | -0.667975 |
| Fe | -5.446095 | -1.736206 | 1.657966 |
| Fe | -5.473138 | -0.115910 | -0.674783 |
| Fe | -2.730422 | -0.377440 | 0.956343 |
| S | -4.899776 | 0.393011 | 1.438623 |
| S | -3.340615 | -0.577767 | -1.242154 |
| S | -3.414889 | -2.584854 | 1.511318 |
| S | -6.278740 | -2.281761 | -0.225624 |
| H | 1.983506 | 1.171063 | -0.600528 |
| H | 0.677300 | -0.108488 | -1.432094 |
| H | -0.823849 | -1.381802 | -0.204024 |
| H | 4.684486 | 2.700377 | -2.547339 |
| H | 3.658571 | 3.360789 | -3.845571 |
| H | 3.753497 | 4.198765 | -2.307078 |
| H | -1.805181 | 7.610727 | 1.854960 |
| H | -0.046206 | 7.789371 | 1.697117 |

| | | | |
|---|---|---|---|
| H | -1.109320 | 8.347144 | 0.397796 |
| H | -2.375067 | 6.495888 | -0.762404 |
| H | -3.463040 | 8.150149 | -4.919581 |
| H | -2.919037 | 5.976094 | -5.966397 |
| H | -4.442949 | 5.843000 | -5.109881 |
| H | -1.737565 | 4.923809 | -3.993623 |
| H | -2.860676 | 3.870694 | -4.842241 |
| H | -4.577959 | 4.170554 | -3.080106 |
| H | -3.567687 | 5.356110 | -2.259272 |
| H | -1.794620 | 3.560984 | -1.950660 |
| H | -2.958307 | 2.365085 | -2.574850 |
| H | -2.012326 | 7.484889 | -4.155938 |
| H | -3.577137 | 7.415454 | -3.323140 |
| H | 4.085981 | 9.726544 | 1.272288 |
| H | 4.021807 | 10.333722 | 2.932274 |
| H | 5.584857 | 10.058097 | 2.151987 |
| H | 5.098594 | 8.194242 | 3.750354 |
| H | 5.168011 | 7.607317 | 2.094191 |
| H | 2.642115 | 7.657984 | 1.985747 |
| H | 2.593315 | 8.264748 | 3.637457 |
| H | 2.095596 | 5.887041 | 3.646744 |
| H | 3.621459 | 6.153317 | 4.480563 |
| H | 8.497346 | 5.952047 | -3.035693 |
| H | 9.719255 | 5.082536 | -2.097422 |
| H | 9.467266 | 4.687962 | -3.802833 |
| H | 7.147716 | 3.858378 | -3.282024 |
| H | 7.360644 | 4.255683 | -1.589838 |
| H | 9.160292 | 2.538291 | -1.371661 |
| H | 9.145589 | 2.273088 | -3.099525 |
| H | 7.999900 | 0.419234 | -2.225287 |
| H | 6.772628 | 1.459506 | -2.953384 |
| H | 4.592340 | -10.565108 | -1.410295 |
| H | 6.200519 | -10.076847 | -1.960958 |
| H | 5.865991 | -10.233011 | -0.231035 |
| H | 0.281513 | -5.492680 | -1.066583 |
| H | 1.350234 | -4.189477 | -0.543913 |
| H | 1.508733 | -5.806548 | 0.176775 |
| H | 5.621162 | -2.994932 | 8.106982 |
| H | 6.705152 | -3.769751 | 6.943839 |
| H | 7.335812 | -2.563721 | 8.075769 |
| H | -0.030126 | -6.713526 | -4.380293 |
| H | 0.726661 | -6.167232 | -2.883155 |
| H | -0.279729 | -5.052660 | -3.824101 |
| H | -3.523735 | -7.290752 | -3.058668 |
| H | -4.088331 | -5.784620 | -3.810703 |
| H | -2.361359 | -6.152446 | -3.767759 |
| H | -9.231072 | -1.610762 | 4.656255 |
| H | -9.547004 | 0.068500 | 4.176253 |
| H | -9.341620 | -1.191338 | 2.940007 |
| H | -7.456489 | 2.782173 | 0.769990 |
| H | -7.787720 | 3.955115 | -0.520014 |
| H | -9.112404 | 2.993733 | 0.164144 |
| H | 4.575437 | -8.159639 | -2.112051 |
| H | 4.212719 | -8.306101 | -0.403084 |
| H | 7.010893 | -7.722175 | -1.516579 |
| H | 6.571725 | -7.695425 | 0.176977 |

| | | | |
|---|---|---|---|
| H | 5.398122 | -5.848466 | -1.960903 |
| H | 6.814950 | -5.439387 | -0.975785 |
| H | 6.005875 | -0.768245 | 6.970154 |
| H | 7.080375 | -1.526204 | 5.822341 |
| H | 0.734158 | 0.022624 | 1.616815 |
| H | 0.034473 | 6.737196 | -1.385934 |
| H | 0.147163 | 3.950968 | -0.388593 |
| H | 1.154526 | 6.459821 | -0.063699 |
| H | 0.427873 | 2.381222 | -2.044491 |
| H | 1.011632 | 2.940564 | -3.620632 |
| H | 2.457864 | -6.211814 | -4.639995 |
| H | 1.436495 | -5.294389 | -5.744368 |
| H | 2.367314 | -4.057561 | -3.109391 |
| H | 3.344167 | -4.010376 | -4.570465 |
| H | -8.106318 | 0.899717 | -0.765948 |
| H | -8.435300 | 2.048286 | -2.057263 |
| H | -7.254113 | 0.216842 | 3.138273 |
| H | -7.154927 | -0.169179 | 4.857246 |
| H | -4.132632 | -5.587426 | -1.306111 |
| H | -2.414656 | -5.986364 | -1.232598 |
| H | 2.498952 | -3.334115 | 4.106720 |
| H | 3.147501 | -1.152897 | 3.041040 |
| H | 4.038196 | -3.903682 | 6.102382 |
| H | 7.252790 | 1.727662 | -0.054233 |
| H | 4.426692 | -0.189835 | 0.810245 |
| H | 4.275733 | -0.647405 | -1.419345 |
| H | 5.259711 | 0.080263 | -2.578996 |
| H | 4.719399 | 5.438978 | 2.331472 |
| H | 4.426971 | 4.456722 | 0.309137 |
| H | 1.446124 | 3.177136 | 1.558468 |
| H | 1.617606 | 3.805521 | 3.158826 |
| H | 5.056877 | -5.912533 | 0.970971 |
| H | 3.935659 | -2.679902 | -1.184693 |
| H | 5.235407 | -3.739282 | -1.795447 |
| H | 3.421359 | -2.795027 | 0.994741 |
| H | 1.081698 | -0.983196 | -4.293148 |
| H | 1.928133 | -1.744812 | -3.005306 |
| H | -2.908850 | 2.492525 | -0.151010 |
| H | -3.371595 | 4.092431 | -0.225686 |
| H | -2.144074 | 2.665969 | 4.522998 |
| H | -1.903885 | 0.998550 | 3.981553 |
| H | 0.474430 | 1.537687 | 3.800075 |
| H | -0.398759 | 2.540340 | 2.876709 |
| H | -1.209466 | 1.549851 | 5.521129 |
| H | -1.410932 | -3.283431 | 2.486654 |
| H | 2.597097 | 1.405070 | -2.791298 |
| H | -4.433847 | 2.859856 | -0.792454 |
| H | 3.282710 | 3.102155 | 0.080914 |
| H | -0.752588 | 5.419461 | 1.271330 |
| H | -0.970561 | -3.983925 | 0.432034 |
| H | -0.842767 | -5.573599 | 1.127458 |
| H | 5.660866 | 0.888692 | 1.415381 |
| H | 3.187094 | -4.383512 | 1.642746 |
| O | 4.330452 | -0.322503 | -4.185128 |
| H | 4.197474 | 0.269773 | -4.934595 |
| H | 3.495321 | -0.839069 | -4.137047 |

-----------------------------------------------------------------------------------------------------------------

**TS(B-C)$_{wt}$**

-----------------------------------------------------------------------------------------------------------------

Number of imaginary frequencies: 9 Electronic energy: HF=-6984.254185
Zero-point correction= 1.765480 (Hartree/Particle)
Thermal correction to Energy= 1.887029
Thermal correction to Enthalpy= 1.887973
Thermal correction to Gibbs Free Energy= 1.579474
Sum of electronic and zero-point Energies= -6982.488705
Sum of electronic and thermal Energies= -6982.367156
Sum of electronic and thermal Enthalpies= -6982.366212
Sum of electronic and thermal Free Energies= -6982.674711

............................................................

Cartesian Coordinates

............................................................

| | | | |
|---|---|---|---|
| C | -3.006553 | -6.613173 | -2.694234 |
| C | -3.092828 | -5.789222 | -1.404003 |
| S | -2.652778 | -4.011299 | -1.697745 |
| C | 0.738387 | -6.151680 | -3.350158 |
| C | 1.813664 | -5.833706 | -4.397934 |
| C | 2.333101 | -4.388747 | -4.324233 |
| C | 1.326353 | -3.388640 | -4.906770 |
| O | 0.671277 | -3.650031 | -5.904804 |
| N | 1.268907 | -2.166331 | -4.284554 |
| C | 0.979763 | -5.094672 | 0.281539 |
| C | 0.230480 | -4.572612 | 1.517311 |
| C | 1.076137 | -3.637089 | 2.373258 |
| O | 2.163034 | -4.049554 | 2.841229 |
| O | 0.650630 | -2.439164 | 2.605482 |
| C | 5.818446 | -9.812742 | -0.285182 |
| C | 5.188891 | -8.447881 | -0.574718 |
| C | 6.247550 | -7.388213 | -0.905680 |
| C | 5.690889 | -6.024542 | -1.315634 |
| N | 5.016901 | -5.353832 | -0.202407 |
| C | 4.536129 | -4.093835 | -0.264813 |
| N | 4.628081 | -3.404256 | -1.415569 |
| N | 4.019486 | -3.509134 | 0.809346 |
| N | -0.455663 | 2.590255 | 3.659917 |
| C | -1.580201 | 2.038015 | 4.444574 |
| C | -8.162104 | 2.584189 | -0.346650 |
| C | -7.832781 | 1.506250 | -1.378721 |
| S | -6.050345 | 1.418390 | -1.895588 |
| C | -9.033693 | -0.968580 | 3.936774 |
| C | -7.553111 | -0.584231 | 3.952542 |
| S | -6.412141 | -2.023288 | 3.679125 |
| C | 8.581616 | 4.875829 | -3.498904 |
| C | 7.848695 | 3.596906 | -3.071345 |
| C | 8.789077 | 2.548428 | -2.462617 |
| C | 8.139647 | 1.200228 | -2.131694 |
| N | 7.199681 | 1.266062 | -0.999218 |
| C | 6.064687 | 0.525834 | -0.892682 |
| N | 5.404789 | 0.135458 | -1.988851 |
| N | 5.594880 | 0.180874 | 0.304114 |
| C | 1.117584 | 2.791145 | -2.603233 |
| C | 0.674102 | 4.256253 | -2.478132 |
| O | 0.860346 | 5.080931 | -3.372413 |

| | | | |
|---|---|---|---|
| C | 2.509152 | 2.630650 | -1.922291 |
| O | 2.460921 | 2.863407 | -0.506754 |
| C | 3.577847 | 3.545802 | -2.506605 |
| N | 0.155483 | 4.599456 | -1.257571 |
| C | 0.005727 | 5.979060 | -0.827214 |
| C | -1.137919 | 6.142450 | 0.186135 |
| O | -2.390353 | 5.676924 | -0.331004 |
| C | -1.260147 | 7.585951 | 0.673910 |
| C | 4.156617 | 9.911180 | 1.767035 |
| C | 3.969757 | 8.717267 | 2.717762 |
| C | 2.525771 | 8.187462 | 2.770158 |
| C | 2.316317 | 6.952365 | 3.669814 |
| N | 3.092170 | 5.782569 | 3.221135 |
| C | 2.657937 | 4.887765 | 2.290977 |
| N | 3.504002 | 4.492146 | 1.336547 |
| N | 1.418375 | 4.395335 | 2.342925 |
| C | -3.281669 | 6.832612 | -4.732588 |
| C | -3.544456 | 5.418618 | -5.275974 |
| C | -2.930612 | 4.274046 | -4.445756 |
| C | -3.612949 | 4.041431 | -3.087552 |
| C | -2.920344 | 2.944486 | -2.275091 |
| N | -3.462590 | 2.864064 | -0.875810 |
| C | 6.429321 | -2.109551 | 7.850265 |
| C | 6.531894 | -1.211914 | 6.606882 |
| C | 5.497300 | -1.481651 | 5.546073 |
| N | 5.352263 | -0.648120 | 4.447973 |
| C | 4.575149 | -2.504506 | 5.475812 |
| C | 4.355724 | -1.170085 | 3.749783 |
| N | 3.851756 | -2.292180 | 4.323863 |
| C | -0.464175 | 1.242770 | 0.638975 |
| O | 0.141337 | 2.375151 | 0.614126 |
| O | -1.699252 | 1.146074 | 0.838889 |
| C | 0.501273 | 0.089311 | 0.484754 |
| O | -0.870211 | -1.415091 | 1.033246 |
| C | 1.105310 | -0.245116 | -0.700378 |
| C | 2.425255 | -0.830245 | -0.714097 |
| O | 3.111427 | -0.950113 | 0.364777 |
| O | 2.948275 | -1.184243 | -1.859090 |
| Fe | -4.067063 | -2.832901 | -0.497324 |
| Fe | -5.529170 | -1.733603 | 1.751068 |
| Fe | -5.395684 | -0.322198 | -0.635917 |
| Fe | -2.693341 | -0.603147 | 1.120553 |
| S | -4.862794 | 0.347997 | 1.442172 |
| S | -3.248184 | -0.817259 | -1.110349 |
| S | -3.574194 | -2.709793 | 1.710063 |
| S | -6.332400 | -2.389814 | -0.117111 |
| H | 1.672421 | 2.410533 | -0.119291 |
| H | 0.610174 | -0.090259 | -1.652369 |
| H | -0.921310 | -1.949438 | 0.223347 |
| H | 4.528727 | 3.378860 | -1.986459 |
| H | 3.718629 | 3.320802 | -3.565131 |
| H | 3.304383 | 4.600535 | -2.412754 |
| H | -2.081988 | 7.673553 | 1.388851 |
| H | -0.338672 | 7.922996 | 1.160416 |
| H | -1.459520 | 8.265383 | -0.163899 |
| H | -2.629225 | 6.233348 | -1.085674 |

| | | | |
|---|---|---|---|
| H | -3.695481 | 7.592816 | -5.402824 |
| H | -3.136838 | 5.357634 | -6.292412 |
| H | -4.626079 | 5.256854 | -5.373421 |
| H | -1.861315 | 4.477395 | -4.294261 |
| H | -2.991165 | 3.346384 | -5.028999 |
| H | -4.664463 | 3.769869 | -3.230980 |
| H | -3.599331 | 4.972135 | -2.507436 |
| H | -1.853399 | 3.148957 | -2.184638 |
| H | -3.064023 | 1.955713 | -2.716620 |
| H | -2.206029 | 7.020547 | -4.634383 |
| H | -3.743070 | 6.986224 | -3.749567 |
| H | 3.889175 | 9.645102 | 0.738684 |
| H | 3.527223 | 10.755983 | 2.066011 |
| H | 5.195816 | 10.252475 | 1.766924 |
| H | 4.286830 | 9.000103 | 3.730391 |
| H | 4.651162 | 7.914563 | 2.399217 |
| H | 2.184149 | 7.946409 | 1.754133 |
| H | 1.863277 | 8.981555 | 3.137804 |
| H | 1.261787 | 6.671062 | 3.689849 |
| H | 2.611746 | 7.168961 | 4.701454 |
| H | 7.886474 | 5.602444 | -3.929535 |
| H | 9.079808 | 5.353770 | -2.647696 |
| H | 9.347328 | 4.659837 | -4.251869 |
| H | 7.343407 | 3.160816 | -3.943708 |
| H | 7.056773 | 3.843139 | -2.353256 |
| H | 9.271648 | 2.952922 | -1.561150 |
| H | 9.605535 | 2.340722 | -3.166258 |
| H | 8.919783 | 0.462465 | -1.900061 |
| H | 7.596670 | 0.821169 | -2.999380 |
| H | 5.053874 | -10.559407 | -0.052033 |
| H | 6.387210 | -10.176347 | -1.148132 |
| H | 6.504639 | -9.762603 | 0.567726 |
| H | 0.336275 | -5.779904 | -0.273952 |
| H | 1.247495 | -4.278489 | -0.396698 |
| H | 1.890163 | -5.629427 | 0.569276 |
| H | 5.453807 | -2.001694 | 8.335715 |
| H | 6.557039 | -3.166129 | 7.590274 |
| H | 7.201558 | -1.851390 | 8.581333 |
| H | 0.344080 | -7.162879 | -3.491614 |
| H | 1.146233 | -6.090955 | -2.336591 |
| H | -0.099325 | -5.451450 | -3.413908 |
| H | -3.258216 | -7.663470 | -2.497429 |
| H | -3.704291 | -6.228818 | -3.444620 |
| H | -2.003618 | -6.577554 | -3.123920 |
| H | -9.253840 | -1.733192 | 4.688822 |
| H | -9.656899 | -0.090621 | 4.149643 |
| H | -9.321698 | -1.366900 | 2.959054 |
| H | -7.603800 | 2.415087 | 0.579288 |
| H | -7.908967 | 3.582262 | -0.721746 |
| H | -9.233592 | 2.572202 | -0.110065 |
| H | 4.490511 | -8.532380 | -1.417207 |
| H | 4.585954 | -8.131146 | 0.286114 |
| H | 6.864163 | -7.745680 | -1.739108 |
| H | 6.934313 | -7.257677 | -0.057564 |
| H | 4.981087 | -6.153714 | -2.143975 |
| H | 6.516777 | -5.394206 | -1.674455 |

| | | | |
|---|---|---|---|
| H | 6.452445 | -0.158986 | 6.907919 |
| H | 7.535761 | -1.312347 | 6.169166 |
| H | 1.084540 | -0.102698 | 1.382425 |
| H | -0.170672 | 6.590883 | -1.718719 |
| H | 0.046386 | 3.866814 | -0.560028 |
| H | 0.935405 | 6.351926 | -0.367080 |
| H | 0.405708 | 2.107020 | -2.137184 |
| H | 1.219271 | 2.542028 | -3.659417 |
| H | 2.659163 | -6.525837 | -4.283799 |
| H | 1.411896 | -5.983451 | -5.404954 |
| H | 2.597565 | -4.117202 | -3.294720 |
| H | 3.246063 | -4.293709 | -4.929441 |
| H | -8.110019 | 0.518298 | -0.995164 |
| H | -8.405519 | 1.665400 | -2.298494 |
| H | -7.335284 | 0.171524 | 3.194829 |
| H | -7.279450 | -0.163330 | 4.926715 |
| H | -4.105082 | -5.840549 | -0.989814 |
| H | -2.411624 | -6.194390 | -0.646933 |
| H | 3.111476 | -2.886016 | 3.923060 |
| H | 3.938890 | -0.773957 | 2.832518 |
| H | 4.366619 | -3.335658 | 6.131409 |
| H | 7.606244 | 1.550804 | -0.117885 |
| H | 4.641674 | -0.248053 | 0.375581 |
| H | 4.535856 | -0.417576 | -1.874721 |
| H | 5.401893 | 0.704301 | -2.826218 |
| H | 4.091501 | 5.935034 | 3.196766 |
| H | 4.282480 | 5.094704 | 1.116034 |
| H | 1.115017 | 3.733350 | 1.626303 |
| H | 0.904064 | 4.256831 | 3.219959 |
| H | 5.082139 | -5.768370 | 0.715840 |
| H | 4.018085 | -2.578168 | -1.563987 |
| H | 4.970324 | -3.870035 | -2.239586 |
| H | 3.657653 | -2.544863 | 0.696444 |
| H | 0.474283 | -1.596292 | -4.544600 |
| H | 1.631755 | -2.040202 | -3.345146 |
| H | -2.953281 | 2.166061 | -0.305418 |
| H | -3.365202 | 3.778873 | -0.419116 |
| H | -2.412259 | 2.747559 | 4.430454 |
| H | -1.953636 | 1.077817 | 4.067482 |
| H | 0.294827 | 1.899684 | 3.663861 |
| H | -0.759037 | 2.627714 | 2.688231 |
| H | -1.271805 | 1.906439 | 5.485876 |
| H | -0.161409 | -2.043340 | 1.907361 |
| H | 2.810727 | 1.600599 | -2.072166 |
| H | -4.469265 | 2.549382 | -0.919649 |
| H | 3.196993 | 3.814316 | 0.599721 |
| H | -0.945120 | 5.487060 | 1.043019 |
| H | -0.704818 | -4.092784 | 1.220886 |
| H | -0.036849 | -5.418381 | 2.161825 |
| H | 6.165268 | 0.234093 | 1.136092 |
| H | 3.598041 | -4.035726 | 1.579247 |
| O | 3.279437 | 0.724567 | -3.931914 |
| H | 2.967013 | 0.006527 | -3.343740 |
| H | 3.076798 | 0.398661 | -4.819014 |

-----------------------------------------------------------------------------------------------------------------------

**$A_{T467A}$**

------------------------------------------------------------------------------------------------------------

Number of imaginary frequencies: 9 Electronic energy: HF=-6869.693626
Zero-point correction= 1.736224 (Hartree/Particle)
Thermal correction to Energy= 1.856069
Thermal correction to Enthalpy= 1.857014
Thermal correction to Gibbs Free Energy= 1.547975
Sum of electronic and zero-point Energies= -6867.957402
Sum of electronic and thermal Energies= -6867.837557
Sum of electronic and thermal Enthalpies= -6867.836612
Sum of electronic and thermal Free Energies= -6868.145651

............................................................

Cartesian Coordinates

............................................................

| | | | |
|---|---|---|---|
| C | -3.448712 | -5.987844 | -3.354885 |
| C | -3.472721 | -5.311024 | -1.975263 |
| S | -2.801099 | -3.577260 | -2.038079 |
| C | 0.290712 | -5.873850 | -4.026987 |
| C | 1.389750 | -5.340102 | -4.957210 |
| C | 1.892599 | -3.942160 | -4.566987 |
| C | 0.925689 | -2.836274 | -5.024039 |
| N | 0.947588 | -1.689870 | -4.285133 |
| O | 0.239489 | -2.968734 | -6.028627 |
| C | 0.708058 | -4.995198 | -0.343909 |
| C | -0.327980 | -4.695955 | 0.741940 |
| C | 0.279992 | -4.038778 | 1.960363 |
| O | 1.437076 | -4.225902 | 2.322911 |
| C | 4.988075 | -10.097645 | -1.282042 |
| C | 4.462484 | -8.658018 | -1.306581 |
| C | 5.583474 | -7.639634 | -1.555098 |
| C | 5.120745 | -6.197945 | -1.774659 |
| N | 4.578974 | -5.590195 | -0.553838 |
| C | 4.145605 | -4.312568 | -0.493982 |
| N | 4.116782 | -3.561289 | -1.601857 |
| N | 3.781073 | -3.771599 | 0.671009 |
| C | -1.040816 | 2.083565 | 4.169749 |
| C | -7.598571 | 3.556377 | -0.317107 |
| C | -7.400995 | 2.312677 | -1.185456 |
| S | -5.658555 | 2.043352 | -1.774252 |
| C | -8.736721 | -0.144587 | 3.702416 |
| C | -7.236674 | -0.095112 | 3.999597 |
| S | -6.336458 | -1.663560 | 3.577714 |
| C | 9.299920 | 4.144512 | -3.610256 |
| C | 8.852846 | 3.187991 | -2.492046 |
| C | 8.066378 | 1.974448 | -3.015146 |
| C | 7.868578 | 0.838867 | -1.997035 |
| N | 6.887732 | 1.079887 | -0.924896 |
| C | 5.716460 | 0.415220 | -0.757711 |
| N | 5.199605 | -0.333520 | -1.737708 |
| N | 5.090199 | 0.495227 | 0.422299 |
| C | 1.718740 | 3.182235 | -2.947125 |
| C | 1.392415 | 4.638797 | -2.626986 |
| O | 1.270745 | 5.500681 | -3.496275 |
| C | 1.310068 | 2.795593 | -4.371091 |
| N | 1.271594 | 4.901332 | -1.297599 |
| C | 0.856646 | 6.210814 | -0.829178 |
| C | 0.065898 | 6.131169 | 0.490957 |

| | | | |
|---|---|---|---|
| C | -0.506748 | 7.494191 | 0.864747 |
| O | -1.010681 | 5.201407 | 0.412017 |
| C | 5.232649 | 9.153492 | 2.018706 |
| C | 5.268845 | 7.775743 | 2.691350 |
| C | 3.871306 | 7.195201 | 2.948379 |
| C | 3.866910 | 5.844030 | 3.683473 |
| N | 4.493335 | 4.747806 | 2.940103 |
| C | 3.865635 | 3.863973 | 2.092232 |
| N | 2.573312 | 3.565457 | 2.367993 |
| C | -2.517758 | 7.335024 | -4.746846 |
| C | -2.905269 | 5.916102 | -5.182928 |
| C | -2.357800 | 4.799176 | -4.277030 |
| C | -2.989709 | 4.752852 | -2.876851 |
| C | -2.435412 | 3.599155 | -2.043424 |
| N | -3.090407 | 3.534937 | -0.696306 |
| S | -4.739538 | 0.675647 | 1.482087 |
| S | -3.031883 | -0.386962 | -1.052373 |
| S | -3.273111 | -2.293662 | 1.612136 |
| S | -6.104336 | -1.975527 | -0.199667 |
| Fe | -3.897651 | -2.405500 | -0.555095 |
| Fe | -5.313637 | -1.455413 | 1.714869 |
| Fe | -5.187003 | 0.161447 | -0.650251 |
| Fe | -2.550153 | -0.071400 | 1.183471 |
| C | -0.390542 | 1.734089 | 0.800531 |
| O | 0.282291 | 2.792058 | 0.718154 |
| O | -1.663393 | 1.703198 | 0.863151 |
| C | 0.344098 | 0.385334 | 0.869665 |
| C | 0.699882 | -0.215958 | -0.502536 |
| C | 2.033554 | -0.958501 | -0.513156 |
| O | 2.616424 | -1.226001 | 0.575145 |
| O | 2.516581 | -1.287479 | -1.658873 |
| O | 2.697513 | 1.812102 | -0.587847 |
| C | 6.516356 | -2.955798 | 7.226566 |
| C | 5.114018 | -2.333920 | 7.335949 |
| C | 4.350072 | -2.367327 | 6.044329 |
| C | 3.279161 | -3.163021 | 5.703734 |
| N | 4.688262 | -1.553531 | 4.978217 |
| C | 3.830032 | -1.856935 | 4.022800 |
| N | 2.953866 | -2.826623 | 4.407997 |
| H | 1.007212 | -5.267649 | -5.979621 |
| H | 2.233524 | -6.043225 | -4.976932 |
| H | 2.852478 | -3.737946 | -5.064287 |
| H | 2.065695 | -3.869440 | -3.485952 |
| H | 0.236822 | -1.007772 | -4.510967 |
| H | 1.384176 | -1.629417 | -3.370412 |
| H | -0.164165 | -6.780404 | -4.438725 |
| H | -0.802446 | -5.625279 | 1.087005 |
| H | -1.135558 | -4.077096 | 0.335216 |
| H | 1.161548 | -4.072876 | -0.715688 |
| H | 3.947937 | -8.434563 | -0.363258 |
| H | 3.706385 | -8.556403 | -2.095827 |
| H | 6.136563 | -7.930855 | -2.456243 |
| H | 6.315303 | -7.666899 | -0.735271 |
| H | 4.343909 | -6.182735 | -2.550037 |
| H | 5.969138 | -5.598583 | -2.134796 |
| H | 4.799643 | -6.030677 | 0.328289 |

| | | | |
|---|---|---|---|
| H | 3.352232 | -4.353031 | 1.384299 |
| H | 3.413442 | -2.803033 | 0.651975 |
| H | 4.356639 | -3.976193 | -2.487427 |
| H | 3.513269 | -2.712164 | -1.615970 |
| H | 5.459772 | -10.361135 | -2.234976 |
| H | -1.357898 | 2.585392 | 3.244604 |
| H | 0.053490 | 2.007635 | 4.162601 |
| H | 8.246825 | 3.734530 | -1.757497 |
| H | 9.741568 | 2.829346 | -1.953794 |
| H | 8.618013 | 1.538236 | -3.858845 |
| H | 7.093461 | 2.290121 | -3.411891 |
| H | 8.825529 | 0.616272 | -1.510421 |
| H | 7.590263 | -0.081949 | -2.517588 |
| H | 7.255220 | 1.469594 | -0.068495 |
| H | 5.672378 | -0.417125 | -2.621800 |
| H | 4.221740 | -0.668686 | -1.682984 |
| H | 5.071517 | 1.414336 | 0.894284 |
| H | 4.195561 | 0.001497 | 0.494883 |
| H | 8.441560 | 4.539600 | -4.164089 |
| H | 1.735232 | 3.493880 | -5.095329 |
| H | 1.646188 | 1.781611 | -4.612396 |
| H | 0.223199 | 2.824923 | -4.490188 |
| H | 2.807745 | 3.079619 | -2.844263 |
| H | 0.757535 | 5.803610 | 1.284862 |
| H | -1.017705 | 7.437804 | 1.829450 |
| H | 0.279832 | 8.253314 | 0.928319 |
| H | -1.233959 | 7.815704 | 0.111954 |
| H | -0.635805 | 4.306007 | 0.576071 |
| H | 0.242584 | 6.665777 | -1.613014 |
| H | 1.275016 | 4.117165 | -0.654396 |
| H | 5.810609 | 7.848191 | 3.644759 |
| H | 5.846263 | 7.085647 | 2.060178 |
| H | 3.331161 | 7.087048 | 1.998639 |
| H | 3.293972 | 7.904711 | 3.556153 |
| H | 2.838430 | 5.563318 | 3.916778 |
| H | 4.388373 | 5.938561 | 4.643631 |
| H | 5.483036 | 4.848783 | 2.770259 |
| H | 2.207475 | 3.680538 | 3.298505 |
| H | 1.976243 | 3.090444 | 1.691468 |
| H | 4.697918 | 9.880778 | 2.639184 |
| H | -2.529055 | 5.745309 | -6.199219 |
| H | -3.998634 | 5.831456 | -5.246049 |
| H | -1.270216 | 4.918042 | -4.186757 |
| H | -2.533997 | 3.832432 | -4.767472 |
| H | -4.076898 | 4.635592 | -2.965975 |
| H | -2.800637 | 5.693871 | -2.343796 |
| H | -1.365457 | 3.719797 | -1.865065 |
| H | -2.617277 | 2.631663 | -2.518034 |
| H | -2.644791 | 2.804631 | -0.098755 |
| H | -2.964433 | 4.427910 | -0.209271 |
| H | -4.099766 | 3.266841 | -0.818706 |
| H | -2.875292 | 8.078811 | -5.466712 |
| H | 5.209866 | -1.291851 | 7.665615 |
| H | 4.537599 | -2.855671 | 8.109145 |
| H | 2.727146 | -3.907306 | 6.257455 |
| H | 2.229147 | -3.246289 | 3.827657 |

| | | | |
|---|---|---|---|
| H | 3.781184 | -1.403171 | 3.041924 |
| H | 6.454506 | -4.015398 | 6.955733 |
| H | -4.044726 | -5.419109 | -4.074890 |
| H | -2.872990 | -5.888782 | -1.262071 |
| H | -4.495869 | -5.275652 | -1.589262 |
| H | -7.300851 | 4.463705 | -0.853746 |
| H | -8.024061 | 2.368563 | -2.083706 |
| H | -7.713941 | 1.417386 | -0.635522 |
| H | -9.213847 | 0.790708 | 4.020514 |
| H | -7.066091 | 0.057281 | 5.071097 |
| H | -6.757079 | 0.723475 | 3.460377 |
| H | 1.253849 | 0.514511 | 1.457566 |
| H | 0.748137 | 0.550009 | -1.275105 |
| H | -0.102066 | -0.900429 | -0.806960 |
| H | 3.055727 | 1.744799 | -1.478541 |
| H | 3.322767 | 2.404398 | -0.103643 |
| H | 7.061448 | -2.877852 | 8.173871 |
| H | 7.098130 | -2.446316 | 6.452237 |
| H | 5.733886 | -10.237415 | -0.491525 |
| H | 4.177576 | -10.809904 | -1.103147 |
| H | 9.852432 | 4.995090 | -3.200787 |
| H | 9.952598 | 3.633107 | -4.325826 |
| H | 4.724212 | 9.107825 | 1.049399 |
| H | 6.242469 | 9.539359 | 1.848321 |
| H | 1.256485 | 2.533577 | -2.205844 |
| H | -1.429140 | 7.435144 | -4.669779 |
| H | -2.945240 | 7.594849 | -3.772055 |
| H | -7.004300 | 3.487635 | 0.599532 |
| H | -8.653253 | 3.661174 | -0.033575 |
| H | -9.221283 | -0.972029 | 4.230577 |
| H | -8.917173 | -0.276815 | 2.631301 |
| H | -2.433701 | -6.064665 | -3.747039 |
| H | -3.868026 | -6.999380 | -3.285728 |
| H | -0.498918 | -5.130634 | -3.888263 |
| H | 0.698036 | -6.126859 | -3.043201 |
| H | 0.209561 | -5.475645 | -1.185315 |
| H | 1.496553 | -5.656631 | 0.024538 |
| H | -1.306469 | 2.731254 | 5.012941 |
| O | -0.560232 | -0.480813 | 1.582075 |
| H | -0.184561 | -1.358797 | 1.754315 |
| H | 1.725533 | 6.870409 | -0.683016 |
| N | -1.605208 | 0.750245 | 4.387835 |
| H | -2.620311 | 0.769023 | 4.327194 |
| H | -1.279124 | 0.104035 | 3.675397 |
| N | 4.473730 | 3.297088 | 1.068921 |
| H | 5.310795 | 3.827070 | 0.829994 |
| O | -0.483783 | -3.198053 | 2.665614 |
| H | -1.433652 | -3.144916 | 2.356330 |

---

**$A_{M1}$**

---

Number of imaginary frequencies: 7 Electronic energy: HF=-6869.676843
Zero-point correction= 1.736859 (Hartree/Particle)
Thermal correction to Energy= 1.858505
Thermal correction to Enthalpy= 1.859449
Thermal correction to Gibbs Free Energy= 1.544689

Sum of electronic and zero-point Energies= -6867.939984
Sum of electronic and thermal Energies= -6867.818338
Sum of electronic and thermal Enthalpies= -6867.817394
Sum of electronic and thermal Free Energies= -6868.132154

........................................................

Cartesian Coordinates

........................................................

| | | | |
|---|---|---|---|
| C | -3.467544 | -6.050940 | -3.151826 |
| C | -3.294945 | -5.398375 | -1.776448 |
| S | -2.846984 | -3.606507 | -1.937215 |
| C | 0.285222 | -5.872566 | -3.895506 |
| C | 1.496395 | -5.566892 | -4.787632 |
| C | 2.173781 | -4.226083 | -4.467226 |
| C | 1.403543 | -3.036941 | -5.061570 |
| O | 0.787566 | -3.136506 | -6.115265 |
| N | 1.512513 | -1.862373 | -4.378169 |
| C | 0.709739 | -5.042312 | -0.221558 |
| C | -0.303719 | -4.666284 | 0.864224 |
| C | 0.290049 | -4.069415 | 2.126454 |
| O | 1.453123 | -4.254593 | 2.484995 |
| O | -0.519721 | -3.361744 | 2.905710 |
| C | 5.107587 | -10.114427 | -1.183071 |
| C | 4.695241 | -8.639921 | -1.191064 |
| C | 5.897673 | -7.705150 | -1.007532 |
| C | 5.600966 | -6.223203 | -1.236205 |
| N | 4.764315 | -5.642233 | -0.180366 |
| C | 4.321874 | -4.363615 | -0.226262 |
| N | 4.502096 | -3.631150 | -1.332600 |
| N | 3.732526 | -3.807659 | 0.830304 |
| N | 0.053873 | 2.425012 | 3.580997 |
| C | -1.118604 | 2.040832 | 4.387639 |
| C | -7.749424 | 3.406223 | -0.166707 |
| C | -7.616696 | 2.329454 | -1.240185 |
| S | -5.874089 | 1.976089 | -1.768152 |
| C | -8.814047 | -0.291155 | 3.947357 |
| C | -7.301143 | -0.228234 | 4.164323 |
| S | -6.395783 | -1.780207 | 3.692524 |
| C | 9.301546 | 4.557472 | -3.149305 |
| C | 8.215468 | 3.564526 | -2.713330 |
| C | 8.802651 | 2.254951 | -2.165579 |
| C | 7.797354 | 1.113064 | -1.981876 |
| N | 6.895501 | 1.305353 | -0.839987 |
| C | 5.810335 | 0.521138 | -0.613099 |
| N | 5.426748 | -0.364135 | -1.539546 |
| N | 5.144543 | 0.625838 | 0.537543 |
| C | 1.453137 | 2.950484 | -2.688235 |
| C | 1.551300 | 4.411585 | -2.247361 |
| O | 2.247452 | 5.243242 | -2.828012 |
| C | 2.588364 | 2.547545 | -3.634156 |
| N | 0.733386 | 4.749238 | -1.203171 |
| C | 0.665639 | 6.118595 | -0.713975 |
| C | -0.398420 | 6.257071 | 0.380824 |
| O | -1.714956 | 5.935528 | -0.104904 |
| C | -0.393263 | 7.648277 | 1.011981 |
| C | 5.205086 | 9.535721 | 1.957857 |
| C | 4.999752 | 8.162945 | 2.620020 |

| | | | |
|---|---|---|---|
| C | 3.579617 | 7.603577 | 2.435016 |
| C | 3.294863 | 6.273068 | 3.159948 |
| N | 4.072526 | 5.140845 | 2.657103 |
| C | 3.612675 | 4.096485 | 1.888656 |
| N | 4.386903 | 3.435484 | 1.052126 |
| N | 2.299305 | 3.764545 | 2.014482 |
| C | -2.634791 | 7.461860 | -4.467556 |
| C | -3.062573 | 6.097017 | -5.031158 |
| C | -2.544729 | 4.872933 | -4.250046 |
| C | -3.228409 | 4.643050 | -2.891937 |
| C | -2.595671 | 3.479895 | -2.124262 |
| N | -3.164453 | 3.351716 | -0.740669 |
| C | 6.588190 | -2.951855 | 7.325839 |
| C | 6.331915 | -1.860355 | 6.280476 |
| C | 5.152734 | -2.095987 | 5.374946 |
| N | 4.875067 | -1.206631 | 4.352003 |
| C | 4.216971 | -3.108477 | 5.379350 |
| C | 3.794047 | -1.676297 | 3.760402 |
| N | 3.351261 | -2.826655 | 4.339864 |
| C | -0.537456 | 1.300686 | 0.493833 |
| O | 0.220030 | 2.283836 | 0.362263 |
| O | -1.788832 | 1.389832 | 0.757056 |
| C | 0.082475 | -0.113154 | 0.428472 |
| O | -0.924592 | -1.125915 | 0.641297 |
| C | 0.887063 | -0.386411 | -0.842039 |
| C | 2.233699 | -1.069625 | -0.635121 |
| O | 2.740084 | -1.173316 | 0.512460 |
| O | 2.831376 | -1.473299 | -1.701469 |
| Fe | -4.172459 | -2.513454 | -0.575568 |
| Fe | -5.509956 | -1.504571 | 1.767187 |
| Fe | -5.501412 | 0.119232 | -0.586635 |
| Fe | -2.784885 | -0.300180 | 1.020021 |
| S | -4.880208 | 0.597013 | 1.523608 |
| S | -3.412681 | -0.478167 | -1.171601 |
| S | -3.547025 | -2.483395 | 1.606460 |
| S | -6.404785 | -1.996688 | -0.104114 |
| H | 0.298269 | -0.960174 | -1.567460 |
| H | -1.181800 | -1.530857 | -0.207983 |
| H | -1.175093 | 7.719819 | 1.772374 |
| H | 0.570471 | 7.869326 | 1.481086 |
| H | -0.577799 | 8.421311 | 0.255218 |
| H | -1.951745 | 6.604392 | -0.762472 |
| H | -3.007609 | 8.277997 | -5.094454 |
| H | -2.700615 | 6.020547 | -6.063677 |
| H | -4.157920 | 6.050180 | -5.091655 |
| H | -1.460531 | 4.976822 | -4.100790 |
| H | -2.686200 | 3.976013 | -4.866125 |
| H | -4.296324 | 4.443026 | -3.036826 |
| H | -3.151898 | 5.551955 | -2.282249 |
| H | -1.521814 | 3.633804 | -2.008825 |
| H | -2.774449 | 2.523941 | -2.622395 |
| H | -1.542534 | 7.545102 | -4.425041 |
| H | -3.024208 | 7.628787 | -3.456210 |
| H | 5.023649 | 9.486122 | 0.878632 |
| H | 4.519119 | 10.280323 | 2.376200 |
| H | 6.226065 | 9.898912 | 2.110015 |

| | | | |
|---|---|---|---|
| H | 5.218916 | 8.240150 | 3.693861 |
| H | 5.734612 | 7.457504 | 2.206574 |
| H | 3.367276 | 7.479824 | 1.364786 |
| H | 2.859737 | 8.341761 | 2.814625 |
| H | 2.239202 | 6.019668 | 3.049245 |
| H | 3.485000 | 6.390426 | 4.235066 |
| H | 8.857733 | 5.480703 | -3.532626 |
| H | 9.958468 | 4.825517 | -2.314086 |
| H | 9.927826 | 4.132986 | -3.941527 |
| H | 7.573293 | 3.333279 | -3.573394 |
| H | 7.561065 | 4.026916 | -1.964316 |
| H | 9.337358 | 2.435152 | -1.222115 |
| H | 9.561325 | 1.885377 | -2.866721 |
| H | 8.339920 | 0.162466 | -1.872011 |
| H | 7.186551 | 1.046126 | -2.889820 |
| H | 4.239993 | -10.768022 | -1.311847 |
| H | 5.811893 | -10.331353 | -1.993723 |
| H | 5.592297 | -10.389224 | -0.239450 |
| H | 0.176720 | -5.503225 | -1.051960 |
| H | 1.224473 | -4.158534 | -0.607597 |
| H | 1.456546 | -5.749292 | 0.151403 |
| H | 5.731144 | -3.067966 | 7.998227 |
| H | 6.772279 | -3.922473 | 6.851735 |
| H | 7.461078 | -2.707182 | 7.938612 |
| H | -0.252959 | -6.753852 | -4.258807 |
| H | 0.600372 | -6.079306 | -2.869682 |
| H | -0.417986 | -5.034848 | -3.874743 |
| H | -3.696670 | -7.119014 | -3.045190 |
| H | -4.286711 | -5.577887 | -3.701576 |
| H | -2.562194 | -5.952919 | -3.755106 |
| H | -9.261169 | -1.126674 | 4.495516 |
| H | -9.283309 | 0.637284 | 4.296080 |
| H | -9.050706 | -0.419698 | 2.886707 |
| H | -7.219702 | 3.111169 | 0.744304 |
| H | -7.333746 | 4.359878 | -0.510844 |
| H | -8.804459 | 3.569744 | 0.087093 |
| H | 4.202143 | -8.403594 | -2.143185 |
| H | 3.949052 | -8.454044 | -0.408818 |
| H | 6.679608 | -7.981695 | -1.725475 |
| H | 6.345890 | -7.843890 | -0.013343 |
| H | 5.072336 | -6.118643 | -2.191889 |
| H | 6.545563 | -5.664881 | -1.310510 |
| H | 6.194572 | -0.895468 | 6.787311 |
| H | 7.226235 | -1.731739 | 5.655851 |
| H | 0.763517 | -0.185210 | 1.282170 |
| H | 0.444264 | 6.792526 | -1.551995 |
| H | 0.389557 | 3.999498 | -0.604944 |
| H | 1.636993 | 6.440859 | -0.311394 |
| H | 1.311032 | 2.318050 | -1.830831 |
| H | 0.519079 | 2.903370 | -3.274098 |
| H | 2.231732 | -6.377502 | -4.698990 |
| H | 1.186222 | -5.526209 | -5.836158 |
| H | 2.312085 | -4.094045 | -3.386466 |
| H | 3.174256 | -4.199511 | -4.923522 |
| H | -8.056213 | 1.388718 | -0.888320 |
| H | -8.161641 | 2.614514 | -2.146163 |

| | | | |
|---|---|---|---|
| H | -6.858741 | 0.600180 | 3.608475 |
| H | -7.076063 | -0.078873 | 5.226314 |
| H | -4.219029 | -5.483703 | -1.195297 |
| H | -2.505502 | -5.906660 | -1.210101 |
| H | 2.550511 | -3.367351 | 4.019654 |
| H | 3.290773 | -1.238474 | 2.908727 |
| H | 4.084381 | -3.971593 | 6.013145 |
| H | 7.296966 | 1.724455 | -0.011850 |
| H | 4.312953 | 0.034587 | 0.627141 |
| H | 4.466699 | -0.747281 | -1.546528 |
| H | 5.999679 | -0.528118 | -2.349833 |
| H | 5.073230 | 5.268134 | 2.630501 |
| H | 5.227357 | 3.978737 | 0.857801 |
| H | 1.922825 | 3.081227 | 1.363702 |
| H | 1.820093 | 3.780906 | 2.911202 |
| H | 4.845267 | -6.045092 | 0.743409 |
| H | 3.885902 | -2.804130 | -1.479908 |
| H | 4.941459 | -4.046797 | -2.136668 |
| H | 3.434088 | -2.822524 | 0.753302 |
| H | 0.968932 | -1.089293 | -4.734254 |
| H | 1.920266 | -1.783027 | -3.450214 |
| H | -2.681220 | 2.604663 | -0.188852 |
| H | -3.026603 | 4.236231 | -0.238290 |
| H | -1.811424 | 2.886041 | 4.441515 |
| H | -1.677328 | 1.178201 | 3.997913 |
| H | 0.688188 | 1.629629 | 3.529057 |
| H | -0.251699 | 2.587095 | 2.623677 |
| H | -0.799945 | 1.810359 | 5.408840 |
| H | -1.405481 | -3.201721 | 2.504927 |
| H | 2.608129 | 1.463011 | -3.789606 |
| H | -4.177368 | 3.085153 | -0.807192 |
| H | 3.403502 | 2.473893 | -0.219198 |
| H | -0.204877 | 5.503161 | 1.151997 |
| H | -1.065369 | -3.994387 | 0.454897 |
| H | -0.855976 | -5.559697 | 1.189289 |
| H | 5.029161 | 1.565632 | 0.951741 |
| H | 3.185238 | -4.355349 | 1.492313 |
| O | 2.906284 | 1.903181 | -0.852360 |
| H | 3.440895 | 1.931822 | -1.652482 |
| H | 1.138952 | 0.558052 | -1.320536 |
| H | 3.567031 | 2.878304 | -3.266541 |
| H | 2.468394 | 3.027748 | -4.608050 |

---

**$B_{M1}$**

---

Number of imaginary frequencies: 7 Electronic energy: HF=-6869.668158
Zero-point correction= 1.732896 (Hartree/Particle)
Thermal correction to Energy= 1.853764
Thermal correction to Enthalpy= 1.854709
Thermal correction to Gibbs Free Energy= 1.544625
Sum of electronic and zero-point Energies= -6867.935262
Sum of electronic and thermal Energies= -6867.814394
Sum of electronic and thermal Enthalpies= -6867.813449
Sum of electronic and thermal Free Energies= -6868.123533

..........................................................

Cartesian Coordinates

..........................................................

| | | | |
|---|---|---|---|
| C | -2.688804 | -6.593277 | -2.963810 |
| C | -2.696702 | -5.815763 | -1.642641 |
| S | -2.448734 | -4.000242 | -1.929537 |
| C | 1.005677 | -5.889440 | -3.687365 |
| C | 2.316130 | -5.551823 | -4.411551 |
| C | 2.839062 | -4.138689 | -4.117489 |
| C | 2.073181 | -3.058539 | -4.896785 |
| O | 1.534173 | -3.302469 | -5.971087 |
| N | 2.098092 | -1.812730 | -4.348508 |
| C | 1.273315 | -4.939935 | -0.027544 |
| C | 0.253139 | -4.589658 | 1.059497 |
| C | 0.808284 | -3.811796 | 2.237981 |
| O | 1.989763 | -3.844101 | 2.579223 |
| O | -0.057492 | -3.114244 | 2.967827 |
| C | 6.364084 | -9.341884 | -0.863486 |
| C | 5.810452 | -7.926121 | -1.035128 |
| C | 6.903186 | -6.859439 | -0.896296 |
| C | 6.437565 | -5.431372 | -1.177946 |
| N | 5.493813 | -4.955468 | -0.165593 |
| C | 4.912924 | -3.728866 | -0.217624 |
| N | 5.082981 | -2.950025 | -1.293100 |
| N | 4.198107 | -3.275151 | 0.805287 |
| N | -0.550400 | 2.397916 | 3.522345 |
| C | -1.596981 | 1.888462 | 4.432557 |
| C | -8.314349 | 2.200711 | -0.188613 |
| C | -7.936447 | 1.115047 | -1.194935 |
| S | -6.180676 | 1.165904 | -1.802143 |
| C | -8.872298 | -1.535996 | 3.989334 |
| C | -7.443023 | -0.998117 | 3.849045 |
| S | -6.146498 | -2.325780 | 3.687274 |
| C | 8.419171 | 5.743079 | -3.086365 |
| C | 7.677323 | 4.439007 | -2.738923 |
| C | 8.603792 | 3.344491 | -2.184221 |
| C | 7.974628 | 1.951826 | -2.015701 |
| N | 7.039460 | 1.850770 | -0.881671 |
| C | 5.959888 | 1.009279 | -0.826415 |
| N | 5.296343 | 0.671498 | -1.938594 |
| N | 5.515065 | 0.558719 | 0.341114 |
| C | 0.880349 | 3.030127 | -2.644124 |
| C | 0.670215 | 4.503304 | -2.297949 |
| O | 1.058955 | 5.428792 | -3.009245 |
| C | 2.043001 | 2.827463 | -3.615104 |
| N | -0.046889 | 4.733483 | -1.152008 |
| C | -0.373021 | 6.087338 | -0.735360 |
| C | -1.462914 | 6.099589 | 0.343802 |
| O | -2.697035 | 5.535445 | -0.128007 |
| C | -1.701653 | 7.502969 | 0.899920 |
| C | 3.602989 | 10.171040 | 1.898078 |
| C | 3.592293 | 8.793350 | 2.580410 |
| C | 2.214234 | 8.110054 | 2.555254 |
| C | 2.155775 | 6.729045 | 3.236670 |
| N | 3.011707 | 5.738850 | 2.575909 |
| C | 2.608802 | 4.670100 | 1.844150 |
| N | 3.418652 | 4.236215 | 0.868443 |
| N | 1.461414 | 4.043838 | 2.083189 |

| | | | |
|---|---|---|---|
| C | -3.798691 | 6.872225 | -4.534931 |
| C | -4.020664 | 5.468360 | -5.119442 |
| C | -3.305232 | 4.329534 | -4.367746 |
| C | -3.898555 | 4.010250 | -2.985924 |
| C | -3.082319 | 2.947684 | -2.248217 |
| N | -3.555786 | 2.753999 | -0.836421 |
| C | 6.721630 | -1.886985 | 7.514664 |
| C | 6.662895 | -1.010679 | 6.253835 |
| C | 5.530504 | -1.316600 | 5.308427 |
| N | 5.303019 | -0.529309 | 4.189108 |
| C | 4.592146 | -2.324381 | 5.364614 |
| C | 4.245158 | -1.058515 | 3.597938 |
| N | 3.776080 | -2.146311 | 4.265897 |
| C | -0.414578 | 1.156813 | 0.462987 |
| O | 0.255097 | 2.222998 | 0.411551 |
| O | -1.680863 | 1.145817 | 0.635161 |
| C | 0.349272 | -0.181678 | 0.388635 |
| O | -0.644488 | -1.263599 | 0.507140 |
| C | 1.244079 | -0.340185 | -0.785211 |
| C | 2.573047 | -0.733604 | -0.676280 |
| O | 3.218493 | -0.792254 | 0.469166 |
| O | 3.292048 | -0.981627 | -1.780412 |
| Fe | -3.788202 | -2.968911 | -0.546478 |
| Fe | -5.200964 | -1.998471 | 1.794205 |
| Fe | -5.392684 | -0.537965 | -0.596855 |
| Fe | -2.555388 | -0.625508 | 0.894240 |
| S | -4.726903 | 0.126612 | 1.450262 |
| S | -3.274275 | -0.890917 | -1.256498 |
| S | -3.150733 | -2.811397 | 1.640771 |
| S | -6.053255 | -2.699343 | -0.024013 |
| H | 0.798643 | -0.300082 | -1.775297 |
| H | -0.707956 | -1.713492 | -0.353399 |
| H | -2.497203 | 7.480875 | 1.648891 |
| H | -0.797176 | 7.908851 | 1.364382 |
| H | -2.000014 | 8.193462 | 0.101178 |
| H | -3.051512 | 6.126407 | -0.807097 |
| H | -4.294242 | 7.633420 | -5.145882 |
| H | -3.669355 | 5.465351 | -6.158395 |
| H | -5.096462 | 5.253405 | -5.163303 |
| H | -2.241293 | 4.585819 | -4.260942 |
| H | -3.343424 | 3.421346 | -4.982176 |
| H | -4.934120 | 3.666804 | -3.086992 |
| H | -3.921132 | 4.923367 | -2.378039 |
| H | -2.034875 | 3.245317 | -2.183340 |
| H | -3.149600 | 1.971319 | -2.733787 |
| H | -2.730897 | 7.117811 | -4.496138 |
| H | -4.202310 | 6.962713 | -3.519455 |
| H | 3.312224 | 10.093826 | 0.844940 |
| H | 2.904303 | 10.858142 | 2.387130 |
| H | 4.598956 | 10.621422 | 1.939546 |
| H | 3.925173 | 8.896351 | 3.621940 |
| H | 4.336816 | 8.155270 | 2.082139 |
| H | 1.871398 | 8.007760 | 1.517111 |
| H | 1.486446 | 8.756076 | 3.063117 |
| H | 1.132233 | 6.349037 | 3.219591 |
| H | 2.453313 | 6.810107 | 4.288886 |

| | | | |
|---|---|---|---|
| H | 7.726762 | 6.493445 | -3.478603 |
| H | 8.912364 | 6.169326 | -2.205489 |
| H | 9.188644 | 5.567481 | -3.845721 |
| H | 7.181368 | 4.060013 | -3.642852 |
| H | 6.876478 | 4.644892 | -2.017633 |
| H | 9.040982 | 3.666842 | -1.228448 |
| H | 9.453088 | 3.217143 | -2.867952 |
| H | 8.772016 | 1.206176 | -1.891839 |
| H | 7.443719 | 1.685760 | -2.932693 |
| H | 5.573995 | -10.091508 | -0.966591 |
| H | 7.130726 | -9.560980 | -1.615088 |
| H | 6.820579 | -9.475221 | 0.123946 |
| H | 0.784001 | -5.559219 | -0.778299 |
| H | 1.645195 | -4.039221 | -0.522220 |
| H | 2.122509 | -5.495606 | 0.380852 |
| H | 5.805130 | -1.791260 | 8.106400 |
| H | 6.844208 | -2.945301 | 7.259126 |
| H | 7.564501 | -1.597472 | 8.149401 |
| H | 0.618115 | -6.857888 | -4.020570 |
| H | 1.157553 | -5.946363 | -2.608447 |
| H | 0.238214 | -5.135081 | -3.883007 |
| H | -2.806187 | -7.668465 | -2.777650 |
| H | -3.509717 | -6.265841 | -3.609046 |
| H | -1.754521 | -6.438545 | -3.508394 |
| H | -8.964031 | -2.203042 | 4.852495 |
| H | -9.573695 | -0.703517 | 4.124345 |
| H | -9.168193 | -2.095999 | 3.097037 |
| H | -7.702840 | 2.120661 | 0.715597 |
| H | -8.168689 | 3.200735 | -0.611782 |
| H | -9.368654 | 2.104193 | 0.099524 |
| H | 5.341076 | -7.830149 | -2.022951 |
| H | 5.013227 | -7.745806 | -0.303633 |
| H | 7.715392 | -7.078788 | -1.600118 |
| H | 7.355715 | -6.905384 | 0.104948 |
| H | 5.940652 | -5.407877 | -2.156182 |
| H | 7.309401 | -4.762067 | -1.228640 |
| H | 6.591806 | 0.045330 | 6.546744 |
| H | 7.614123 | -1.094798 | 5.709009 |
| H | 0.959558 | -0.241613 | 1.296998 |
| H | -0.692550 | 6.658557 | -1.615330 |
| H | -0.191051 | 3.954207 | -0.511330 |
| H | 0.515213 | 6.609673 | -0.344587 |
| H | 0.960082 | 2.436580 | -1.747389 |
| H | -0.044352 | 2.706938 | -3.147298 |
| H | 3.085704 | -6.286344 | -4.137776 |
| H | 2.169390 | -5.624680 | -5.493563 |
| H | 2.819567 | -3.916811 | -3.043671 |
| H | 3.889569 | -4.060076 | -4.436153 |
| H | -8.107458 | 0.122960 | -0.760459 |
| H | -8.565100 | 1.181365 | -2.088993 |
| H | -7.359608 | -0.338079 | 2.983016 |
| H | -7.167986 | -0.413976 | 4.733994 |
| H | -3.643969 | -5.968634 | -1.115229 |
| H | -1.895418 | -6.174577 | -0.986292 |
| H | 3.004387 | -2.734810 | 3.950966 |
| H | 3.771093 | -0.708171 | 2.689964 |

| | | | |
|---|---|---|---|
| H | 4.428640 | -3.122084 | 6.072380 |
| H | 7.459754 | 2.040273 | 0.019042 |
| H | 4.591287 | 0.014791 | 0.399828 |
| H | 4.472792 | -0.047363 | -1.901126 |
| H | 5.549945 | 1.102068 | -2.813343 |
| H | 3.975022 | 6.008406 | 2.439055 |
| H | 4.031238 | 4.911417 | 0.436076 |
| H | 1.133581 | 3.314588 | 1.430994 |
| H | 0.980869 | 4.031414 | 2.982238 |
| H | 5.565760 | -5.365726 | 0.754991 |
| H | 4.386136 | -2.176927 | -1.475247 |
| H | 5.604043 | -3.308454 | -2.075930 |
| H | 3.779277 | -2.314711 | 0.719806 |
| H | 1.553298 | -1.111300 | -4.829082 |
| H | 2.444873 | -1.600962 | -3.411203 |
| H | -2.936114 | 2.097033 | -0.319168 |
| H | -3.524277 | 3.654589 | -0.342301 |
| H | -2.406392 | 2.621562 | 4.497474 |
| H | -2.036637 | 0.932602 | 4.118849 |
| H | 0.183096 | 1.692056 | 3.464392 |
| H | -0.948741 | 2.424826 | 2.584668 |
| H | -1.179999 | 1.765064 | 5.436627 |
| H | -0.957707 | -3.086861 | 2.567058 |
| H | 2.192984 | 1.761024 | -3.813496 |
| H | -4.522429 | 2.349836 | -0.839509 |
| H | 3.194620 | 3.388764 | 0.299259 |
| H | -1.160059 | 5.432002 | 1.159161 |
| H | -0.593584 | -4.049217 | 0.625505 |
| H | -0.176242 | -5.507519 | 1.486542 |
| H | 6.044839 | 0.687813 | 1.191821 |
| H | 3.744831 | -3.896806 | 1.467708 |
| O | 2.871713 | 2.197536 | -0.833865 |
| H | 3.536480 | 1.790422 | -1.402963 |
| H | 2.227512 | 1.477791 | -0.633243 |
| H | 2.970340 | 3.239182 | -3.207568 |
| H | 1.850045 | 3.336219 | -4.562529 |

---

**$C_{M1}$**

---

Number of imaginary frequencies: 8 Electronic energy: HF=-6869.678115
Zero-point correction= 1.734249 (Hartree/Particle)
Thermal correction to Energy= 1.855751
Thermal correction to Enthalpy= 1.856695
Thermal correction to Gibbs Free Energy= 1.544917
Sum of electronic and zero-point Energies= -6867.943866
Sum of electronic and thermal Energies= -6867.822364
Sum of electronic and thermal Enthalpies= -6867.821420
Sum of electronic and thermal Free Energies= -6868.133198

..........................................................

Cartesian Coordinates

..........................................................

| | | | |
|---|---|---|---|
| C | -2.126064 | -6.854621 | -2.698745 |
| C | -2.395612 | -6.022202 | -1.442473 |
| S | -2.228495 | -4.215357 | -1.808984 |
| C | 1.546404 | -5.975962 | -3.338710 |
| C | 2.943332 | -5.618422 | -3.859080 |

| | | | |
|---|---|---|---|
| C | 3.272675 | -4.123143 | -3.736777 |
| C | 2.484901 | -3.265342 | -4.733568 |
| O | 2.294714 | -3.628833 | -5.886103 |
| N | 2.082857 | -2.041399 | -4.265415 |
| C | 1.658358 | -4.921580 | 0.303732 |
| C | 1.151160 | -4.726056 | 1.744393 |
| C | 1.780779 | -3.471258 | 2.347742 |
| O | 2.905472 | -3.632770 | 2.954316 |
| O | 1.228765 | -2.362684 | 2.150474 |
| C | 6.992732 | -9.069497 | -0.277274 |
| C | 6.241321 | -7.735348 | -0.383701 |
| C | 7.199364 | -6.535730 | -0.431828 |
| C | 6.572096 | -5.198194 | -0.835405 |
| N | 5.742980 | -4.597370 | 0.215611 |
| C | 5.171538 | -3.378836 | 0.087343 |
| N | 5.236213 | -2.736405 | -1.102178 |
| N | 4.594260 | -2.781707 | 1.119543 |
| N | -0.363203 | 2.488213 | 4.727888 |
| C | -1.689827 | 1.856762 | 4.495821 |
| C | -8.277844 | 1.698619 | -0.315489 |
| C | -7.907896 | 0.788507 | -1.483425 |
| S | -6.119228 | 0.843211 | -1.974520 |
| C | -8.761797 | -1.955608 | 3.943849 |
| C | -7.360963 | -1.345106 | 3.857778 |
| S | -6.009594 | -2.599479 | 3.625085 |
| C | 8.116805 | 5.854825 | -3.394767 |
| C | 7.147711 | 4.669346 | -3.367982 |
| C | 7.639635 | 3.477332 | -4.202676 |
| C | 6.736892 | 2.240728 | -4.123292 |
| N | 6.733411 | 1.704093 | -2.758711 |
| C | 5.748532 | 0.978851 | -2.190270 |
| N | 4.748173 | 0.495107 | -2.928165 |
| N | 5.779065 | 0.740350 | -0.875033 |
| C | 0.927918 | 2.948816 | -2.537811 |
| C | 0.607864 | 4.412685 | -2.246365 |
| O | 0.988260 | 5.320556 | -2.985257 |
| C | 0.305338 | 2.515611 | -3.882729 |
| N | -0.148153 | 4.642220 | -1.135687 |
| C | -0.536174 | 5.982298 | -0.747134 |
| C | -1.661663 | 5.967644 | 0.298021 |
| O | -2.856852 | 5.356621 | -0.207616 |
| C | -1.963901 | 7.370069 | 0.825043 |
| C | 3.144859 | 10.334771 | 1.884944 |
| C | 3.152856 | 9.037110 | 2.712852 |
| C | 1.762777 | 8.394747 | 2.877867 |
| C | 1.743023 | 7.066702 | 3.665003 |
| N | 2.518478 | 6.010120 | 2.998354 |
| C | 2.005564 | 5.013557 | 2.224192 |
| N | 2.653518 | 4.750509 | 1.074379 |
| N | 0.945002 | 4.306136 | 2.576721 |
| C | -3.886810 | 6.489925 | -4.659026 |
| C | -4.205442 | 5.065156 | -5.137065 |
| C | -3.469248 | 3.949363 | -4.371590 |
| C | -3.991259 | 3.707362 | -2.946157 |
| C | -3.134767 | 2.691128 | -2.188240 |
| N | -3.608086 | 2.488733 | -0.778794 |

| | | | |
|---|---|---|---|
| C | 6.720960 | -1.396978 | 7.905422 |
| C | 6.425537 | -0.241954 | 6.939842 |
| C | 5.496964 | -0.597006 | 5.810488 |
| N | 5.227185 | 0.301453 | 4.793734 |
| C | 4.801381 | -1.768022 | 5.592217 |
| C | 4.384671 | -0.331175 | 3.992119 |
| N | 4.090210 | -1.583027 | 4.428552 |
| C | -0.513033 | 1.437979 | 0.752077 |
| O | -0.416728 | 2.703929 | 0.899449 |
| O | -1.609554 | 0.851611 | 0.569211 |
| C | 0.817445 | 0.743503 | 0.713883 |
| O | -0.709003 | -1.983645 | 0.308452 |
| C | 1.213623 | -0.035632 | -0.305580 |
| C | 2.636360 | -0.438825 | -0.484045 |
| O | 3.518962 | -0.047250 | 0.349802 |
| O | 2.938440 | -1.100820 | -1.536799 |
| Fe | -3.857301 | -3.201288 | -0.694115 |
| Fe | -5.265062 | -2.153200 | 1.676204 |
| Fe | -5.322413 | -0.785793 | -0.693909 |
| Fe | -2.447454 | -1.012714 | 0.781276 |
| S | -4.615961 | -0.082774 | 1.320871 |
| S | -3.228357 | -1.145161 | -1.375421 |
| S | -3.290170 | -3.100601 | 1.504633 |
| S | -6.149378 | -2.844768 | -0.141591 |
| H | 0.509704 | -0.377393 | -1.054352 |
| H | -1.010956 | -2.805016 | -0.130547 |
| H | -2.785593 | 7.330969 | 1.544542 |
| H | -1.091293 | 7.810231 | 1.318765 |
| H | -2.254277 | 8.042082 | 0.007669 |
| H | -3.216282 | 5.928945 | -0.899765 |
| H | -4.407791 | 7.234962 | -5.268909 |
| H | -3.939007 | 4.987545 | -6.198208 |
| H | -5.287836 | 4.889640 | -5.083509 |
| H | -2.398299 | 4.193643 | -4.338911 |
| H | -3.556370 | 3.012706 | -4.936158 |
| H | -5.024330 | 3.345719 | -2.983392 |
| H | -3.995260 | 4.652167 | -2.390052 |
| H | -2.101844 | 3.033498 | -2.119526 |
| H | -3.158974 | 1.706504 | -2.661621 |
| H | -2.812037 | 6.695805 | -4.724220 |
| H | -4.196445 | 6.652476 | -3.619670 |
| H | 2.768948 | 10.156583 | 0.871699 |
| H | 2.506332 | 11.094340 | 2.348173 |
| H | 4.153108 | 10.750492 | 1.801930 |
| H | 3.576383 | 9.237507 | 3.705903 |
| H | 3.839691 | 8.327567 | 2.227880 |
| H | 1.314790 | 8.226312 | 1.889102 |
| H | 1.102513 | 9.098250 | 3.400993 |
| H | 0.717504 | 6.707306 | 3.775801 |
| H | 2.148046 | 7.212518 | 4.672558 |
| H | 7.733513 | 6.690242 | -2.801546 |
| H | 9.096565 | 5.578942 | -2.988858 |
| H | 8.272280 | 6.218143 | -4.416292 |
| H | 6.163799 | 4.982922 | -3.739383 |
| H | 6.989321 | 4.355466 | -2.327718 |
| H | 8.658207 | 3.195262 | -3.898058 |

| | | | |
|---|---|---|---|
| H | 7.713751 | 3.767766 | -5.256749 |
| H | 7.079288 | 1.476143 | -4.832633 |
| H | 5.710290 | 2.517863 | -4.390280 |
| H | 6.295685 | -9.911540 | -0.239578 |
| H | 7.652539 | -9.218928 | -1.139064 |
| H | 7.610838 | -9.109328 | 0.626709 |
| H | 1.178783 | -5.790635 | -0.154648 |
| H | 1.428854 | -4.047726 | -0.313455 |
| H | 2.741322 | -5.089186 | 0.287358 |
| H | 5.803899 | -1.759702 | 8.382183 |
| H | 7.179484 | -2.243000 | 7.381934 |
| H | 7.405688 | -1.080316 | 8.698481 |
| H | 1.362113 | -7.051330 | -3.427915 |
| H | 1.422711 | -5.696565 | -2.290630 |
| H | 0.771863 | -5.457964 | -3.911950 |
| H | -2.202691 | -7.927600 | -2.478779 |
| H | -2.848761 | -6.613098 | -3.484312 |
| H | -1.126780 | -6.658177 | -3.094913 |
| H | -8.828367 | -2.682750 | 4.759531 |
| H | -9.507209 | -1.170789 | 4.123729 |
| H | -9.019808 | -2.469672 | 3.013000 |
| H | -7.713513 | 1.424846 | 0.581278 |
| H | -8.059920 | 2.747773 | -0.546170 |
| H | -9.348249 | 1.616957 | -0.087292 |
| H | 5.626915 | -7.736920 | -1.293579 |
| H | 5.541366 | -7.631777 | 0.454588 |
| H | 7.980448 | -6.741076 | -1.174471 |
| H | 7.725111 | -6.421869 | 0.526522 |
| H | 5.954542 | -5.356684 | -1.729719 |
| H | 7.374133 | -4.495985 | -1.103594 |
| H | 6.004116 | 0.603385 | 7.502198 |
| H | 7.366773 | 0.134295 | 6.516334 |
| H | 1.540365 | 1.040539 | 1.469931 |
| H | -0.846916 | 6.538036 | -1.640817 |
| H | -0.355965 | 3.878649 | -0.490423 |
| H | 0.321220 | 6.537256 | -0.332543 |
| H | 2.000064 | 2.873709 | -2.656281 |
| H | 0.569793 | 2.288978 | -1.746220 |
| H | 3.701485 | -6.193211 | -3.309096 |
| H | 3.033377 | -5.896028 | -4.914336 |
| H | 3.107259 | -3.772393 | -2.712387 |
| H | 4.336037 | -3.963768 | -3.974658 |
| H | -8.149354 | -0.252148 | -1.245502 |
| H | -8.476282 | 1.058084 | -2.380271 |
| H | -7.301892 | -0.623811 | 3.038006 |
| H | -7.120080 | -0.807863 | 4.781973 |
| H | -3.404216 | -6.219137 | -1.064689 |
| H | -1.692176 | -6.288677 | -0.646060 |
| H | 3.511211 | -2.288961 | 3.933959 |
| H | 3.964900 | 0.063023 | 3.074980 |
| H | 4.744684 | -2.691085 | 6.148261 |
| H | 7.503321 | 1.971545 | -2.163565 |
| H | 4.966283 | 0.286462 | -0.385666 |
| H | 4.016179 | -0.093707 | -2.491149 |
| H | 4.777467 | 0.552228 | -3.933003 |
| H | 3.459400 | 6.276912 | 2.742872 |

| | | | |
|---|---|---|---|
| H | 3.155063 | 5.501851 | 0.625856 |
| H | 0.511372 | 3.639164 | 1.898733 |
| H | 0.632059 | 4.160640 | 3.540887 |
| H | 5.832056 | -4.951096 | 1.158222 |
| H | 4.454895 | -2.090196 | -1.295396 |
| H | 5.558664 | -3.261530 | -1.899727 |
| H | 4.209185 | -1.851200 | 0.955651 |
| H | 1.417731 | -1.554925 | -4.851728 |
| H | 2.051981 | -1.848991 | -3.269005 |
| H | -2.975792 | 1.845568 | -0.256652 |
| H | -3.612959 | 3.390147 | -0.286596 |
| H | -1.790666 | 1.592912 | 3.441014 |
| H | -1.860673 | 0.952485 | 5.093642 |
| H | -0.278474 | 2.695818 | 5.722096 |
| H | 0.353462 | 1.783698 | 4.555106 |
| H | -2.476584 | 2.578955 | 4.730818 |
| H | -0.081780 | -2.218537 | 1.049306 |
| H | 0.645728 | 1.510649 | -4.153100 |
| H | -4.556889 | 2.049657 | -0.795809 |
| H | 2.451092 | 3.947150 | 0.470950 |
| H | -1.366477 | 5.318647 | 1.128667 |
| H | 0.061809 | -4.623029 | 1.741280 |
| H | 1.416060 | -5.593976 | 2.355187 |
| H | 6.477328 | 1.191980 | -0.304813 |
| H | 4.105548 | -3.292328 | 1.903540 |
| O | 2.867985 | 2.716846 | -0.903811 |
| H | 3.544720 | 2.259724 | -0.375432 |
| H | 2.113853 | 2.114753 | -0.805957 |
| H | 0.613533 | 3.209201 | -4.669869 |
| H | -0.785913 | 2.493127 | -3.858684 |

---

**TS(A-B)$_{M1}$**

---

Number of imaginary frequencies: 9 Electronic energy: HF=-6869.644109
Zero-point correction= 1.729330 (Hartree/Particle)
Thermal correction to Energy= 1.848697
Thermal correction to Enthalpy= 1.849641
Thermal correction to Gibbs Free Energy= 1.542912
Sum of electronic and zero-point Energies= -6867.914779
Sum of electronic and thermal Energies= -6867.795412
Sum of electronic and thermal Enthalpies= -6867.794468
Sum of electronic and thermal Free Energies= -6868.101197

........................................................

Cartesian Coordinates

........................................................

| | | | |
|---|---|---|---|
| C | -3.142184 | -6.303080 | -3.052068 |
| C | -2.997753 | -5.588218 | -1.706283 |
| S | -2.649354 | -3.785124 | -1.947873 |
| C | 0.590767 | -5.887941 | -3.800980 |
| C | 1.811493 | -5.513096 | -4.653142 |
| C | 2.404674 | -4.138125 | -4.311306 |
| C | 1.587416 | -2.989885 | -4.923567 |
| O | 0.990632 | -3.125728 | -5.985216 |
| N | 1.634676 | -1.807178 | -4.248514 |
| C | 0.963332 | -5.008102 | -0.132952 |
| C | -0.045806 | -4.586599 | 0.938617 |

| | | | |
|---|---|---|---|
| C | 0.547359 | -3.937555 | 2.175134 |
| O | 1.715875 | -4.084057 | 2.532818 |
| O | -0.275396 | -3.220921 | 2.933945 |
| C | 5.678462 | -9.792242 | -1.066437 |
| C | 5.186745 | -8.345826 | -1.164345 |
| C | 6.340093 | -7.337955 | -1.082808 |
| C | 5.938066 | -5.881766 | -1.317934 |
| N | 5.100572 | -5.359231 | -0.235876 |
| C | 4.590281 | -4.104204 | -0.237956 |
| N | 4.744941 | -3.313568 | -1.306764 |
| N | 3.959978 | -3.626852 | 0.832098 |
| N | -0.141860 | 2.420762 | 3.659186 |
| C | -1.314863 | 1.972336 | 4.432336 |
| C | -8.023014 | 2.877562 | -0.123511 |
| C | -7.801311 | 1.765259 | -1.146291 |
| S | -6.056306 | 1.603632 | -1.758597 |
| C | -8.844220 | -0.853849 | 4.015539 |
| C | -7.338993 | -0.569880 | 4.019637 |
| S | -6.288834 | -2.065160 | 3.671069 |
| C | 8.916208 | 5.106419 | -3.132092 |
| C | 7.974856 | 3.946216 | -2.757563 |
| C | 8.712070 | 2.749790 | -2.131850 |
| C | 7.895623 | 1.457270 | -1.976345 |
| N | 6.922278 | 1.494019 | -0.866761 |
| C | 5.837698 | 0.671615 | -0.774277 |
| N | 5.226851 | 0.243797 | -1.886937 |
| N | 5.373523 | 0.290143 | 0.410738 |
| C | 1.188242 | 2.999733 | -2.652006 |
| C | 1.284541 | 4.430687 | -2.125360 |
| O | 2.089055 | 5.259758 | -2.550220 |
| C | 2.356881 | 2.621455 | -3.564264 |
| N | 0.320710 | 4.764918 | -1.205799 |
| C | 0.199226 | 6.123271 | -0.697495 |
| C | -0.893032 | 6.209147 | 0.374777 |
| O | -2.178016 | 5.813578 | -0.135633 |
| C | -0.977040 | 7.600729 | 1.000060 |
| C | 4.510667 | 9.843378 | 1.947233 |
| C | 4.458554 | 8.426568 | 2.534815 |
| C | 3.032125 | 7.860625 | 2.623256 |
| C | 2.937741 | 6.432769 | 3.190011 |
| N | 3.622715 | 5.452118 | 2.346554 |
| C | 3.050404 | 4.442790 | 1.634917 |
| N | 3.650197 | 3.994346 | 0.534910 |
| N | 1.911763 | 3.877558 | 2.041159 |
| C | -3.183465 | 7.226415 | -4.456074 |
| C | -3.544626 | 5.854703 | -5.047796 |
| C | -2.960309 | 4.645864 | -4.291869 |
| C | -3.607179 | 4.377903 | -2.923548 |
| C | -2.894848 | 3.256850 | -2.164612 |
| N | -3.422036 | 3.111648 | -0.767080 |
| C | 6.699801 | -2.493740 | 7.394354 |
| C | 6.631348 | -1.583009 | 6.159133 |
| C | 5.443387 | -1.807337 | 5.260691 |
| N | 5.193161 | -0.967601 | 4.185243 |
| C | 4.469050 | -2.779880 | 5.326067 |
| C | 4.086554 | -1.430790 | 3.629991 |

| | | | |
|---|---|---|---|
| N | 3.607434 | -2.526120 | 4.277974 |
| C | -0.554148 | 1.357517 | 0.557228 |
| O | 0.083106 | 2.438190 | 0.512812 |
| O | -1.823191 | 1.313342 | 0.696174 |
| C | 0.225442 | 0.025800 | 0.510376 |
| O | -0.735934 | -1.077247 | 0.563724 |
| C | 1.180368 | -0.070024 | -0.645251 |
| C | 2.367972 | -0.895336 | -0.533142 |
| O | 2.950629 | -1.074521 | 0.597915 |
| O | 2.963841 | -1.279167 | -1.632409 |
| Fe | -3.964541 | -2.738325 | -0.563320 |
| Fe | -5.337151 | -1.799764 | 1.771623 |
| Fe | -5.459401 | -0.215532 | -0.599785 |
| Fe | -2.672561 | -0.447394 | 0.947516 |
| S | -4.821424 | 0.328828 | 1.489764 |
| S | -3.346536 | -0.675057 | -1.221241 |
| S | -3.307975 | -2.646895 | 1.607250 |
| S | -6.209204 | -2.387101 | -0.080796 |
| H | 0.686975 | -0.142750 | -1.617353 |
| H | -0.843922 | -1.441524 | -0.332161 |
| H | -1.779854 | 7.632125 | 1.740978 |
| H | -0.038505 | 7.874782 | 1.492678 |
| H | -1.184526 | 8.360711 | 0.236383 |
| H | -2.424313 | 6.443035 | -0.827823 |
| H | -3.595059 | 8.036959 | -5.065840 |
| H | -3.185206 | 5.817834 | -6.083384 |
| H | -4.636506 | 5.753777 | -5.103379 |
| H | -1.878872 | 4.795388 | -4.161273 |
| H | -3.074088 | 3.749337 | -4.913886 |
| H | -4.664038 | 4.116439 | -3.048972 |
| H | -3.573024 | 5.290820 | -2.315749 |
| H | -1.828154 | 3.469869 | -2.079861 |
| H | -3.030719 | 2.287000 | -2.649079 |
| H | -2.096352 | 7.361861 | -4.411050 |
| H | -3.580717 | 7.353579 | -3.442061 |
| H | 4.101428 | 9.865423 | 0.931462 |
| H | 3.928802 | 10.544389 | 2.555398 |
| H | 5.538860 | 10.214676 | 1.902433 |
| H | 4.911629 | 8.423705 | 3.535306 |
| H | 5.085822 | 7.772286 | 1.912110 |
| H | 2.566717 | 7.878109 | 1.628931 |
| H | 2.427775 | 8.514409 | 3.265360 |
| H | 1.890217 | 6.133255 | 3.266594 |
| H | 3.358694 | 6.399197 | 4.202901 |
| H | 8.355658 | 5.935132 | -3.573705 |
| H | 9.446593 | 5.491095 | -2.253763 |
| H | 9.668018 | 4.783331 | -3.860131 |
| H | 7.452387 | 3.606520 | -3.661971 |
| H | 7.194659 | 4.302716 | -2.073724 |
| H | 9.144561 | 3.032620 | -1.161352 |
| H | 9.567977 | 2.490718 | -2.768165 |
| H | 8.579043 | 0.610534 | -1.824886 |
| H | 7.358369 | 1.264031 | -2.907705 |
| H | 4.844942 | -10.498084 | -1.126646 |
| H | 6.374710 | -10.028896 | -1.878635 |
| H | 6.199254 | -9.973971 | -0.119453 |

| | | | |
|---|---|---|---|
| H | 0.428498 | -5.510977 | -0.937794 |
| H | 1.470691 | -4.139767 | -0.561125 |
| H | 1.716308 | -5.693528 | 0.267816 |
| H | 5.815886 | -2.367628 | 8.028328 |
| H | 6.757189 | -3.549931 | 7.108618 |
| H | 7.583090 | -2.263633 | 7.997627 |
| H | 0.117630 | -6.799404 | -4.180601 |
| H | 0.881261 | -6.074808 | -2.764264 |
| H | -0.159819 | -5.092171 | -3.806824 |
| H | -3.313818 | -7.376898 | -2.902732 |
| H | -3.988022 | -5.896587 | -3.614736 |
| H | -2.245972 | -6.181322 | -3.664811 |
| H | -9.105750 | -1.631667 | 4.740212 |
| H | -9.398447 | 0.056425 | 4.276118 |
| H | -9.175577 | -1.188679 | 3.027832 |
| H | -7.419696 | 2.704177 | 0.773036 |
| H | -7.749363 | 3.854583 | -0.537117 |
| H | -9.077951 | 2.920101 | 0.175669 |
| H | 4.652867 | -8.201518 | -2.112672 |
| H | 4.454531 | -8.146520 | -0.372089 |
| H | 7.091977 | -7.587885 | -1.841274 |
| H | 6.854277 | -7.423172 | -0.114587 |
| H | 5.374723 | -5.815391 | -2.257410 |
| H | 6.842457 | -5.264563 | -1.424022 |
| H | 6.626540 | -0.532796 | 6.479394 |
| H | 7.552198 | -1.700587 | 5.569926 |
| H | 0.795630 | -0.045634 | 1.443316 |
| H | -0.029096 | 6.807957 | -1.526313 |
| H | -0.046762 | 4.007763 | -0.630917 |
| H | 1.151349 | 6.474255 | -0.271680 |
| H | 0.992615 | 2.308240 | -1.852528 |
| H | 0.280882 | 3.009025 | -3.282597 |
| H | 2.587225 | -6.282459 | -4.541253 |
| H | 1.532396 | -5.487695 | -5.710906 |
| H | 2.506907 | -4.001479 | -3.227579 |
| H | 3.414833 | -4.055769 | -4.739314 |
| H | -8.097433 | 0.798699 | -0.721282 |
| H | -8.423841 | 1.924996 | -2.032761 |
| H | -7.082416 | 0.199695 | 3.289711 |
| H | -7.023603 | -0.210752 | 5.005576 |
| H | -3.911377 | -5.700662 | -1.113232 |
| H | -2.176265 | -6.026640 | -1.127435 |
| H | 2.798944 | -3.070256 | 3.978523 |
| H | 3.581241 | -1.024447 | 2.764232 |
| H | 4.308034 | -3.597698 | 6.011009 |
| H | 7.309878 | 1.777717 | 0.023764 |
| H | 4.453259 | -0.198222 | 0.496020 |
| H | 4.361425 | -0.345405 | -1.818225 |
| H | 5.390862 | 0.734290 | -2.752042 |
| H | 4.558763 | 5.691832 | 2.055526 |
| H | 4.201379 | 4.664637 | 0.016539 |
| H | 1.418433 | 3.237139 | 1.406061 |
| H | 1.571617 | 3.888793 | 2.997895 |
| H | 5.188705 | -5.797626 | 0.670228 |
| H | 4.077793 | -2.511359 | -1.429073 |
| H | 5.216292 | -3.670310 | -2.121267 |

| | | | |
|---|---|---|---|
| H | 3.582779 | -2.658737 | 0.774599 |
| H | 1.044175 | -1.071328 | -4.609132 |
| H | 2.015977 | -1.703224 | -3.309577 |
| H | -2.869889 | 2.411948 | -0.219756 |
| H | -3.340496 | 4.011000 | -0.279272 |
| H | -2.032188 | 2.795174 | 4.508103 |
| H | -1.841748 | 1.110877 | 3.999517 |
| H | 0.508342 | 1.639855 | 3.581557 |
| H | -0.447253 | 2.604913 | 2.705872 |
| H | -1.005783 | 1.710595 | 5.448940 |
| H | -1.169370 | -3.111304 | 2.531660 |
| H | 2.390012 | 1.537914 | -3.723520 |
| H | -4.415427 | 2.772724 | -0.807145 |
| H | 3.238952 | 3.122085 | -0.025875 |
| H | -0.676593 | 5.468419 | 1.152695 |
| H | -0.803990 | -3.926602 | 0.505249 |
| H | -0.603715 | -5.462451 | 1.299742 |
| H | 5.894625 | 0.441773 | 1.262991 |
| H | 3.456618 | -4.232317 | 1.475292 |
| O | 2.700759 | 2.052314 | -0.813127 |
| H | 3.335982 | 1.766900 | -1.477752 |
| H | 1.921314 | 1.083639 | -0.680318 |
| H | 3.310158 | 2.956418 | -3.146704 |
| H | 2.260914 | 3.102159 | -4.541770 |

------------------------------------------------------------------------------------------------------------

**TS(B-C)$_{M1}$**

------------------------------------------------------------------------------------------------------------

Number of imaginary frequencies: 9 Electronic energy: HF=-6869.668492
Zero-point correction= 1.732550 (Hartree/Particle)
Thermal correction to Energy= 1.852268
Thermal correction to Enthalpy= 1.853212
Thermal correction to Gibbs Free Energy= 1.547081
Sum of electronic and zero-point Energies= -6867.935942
Sum of electronic and thermal Energies= -6867.816224
Sum of electronic and thermal Enthalpies= -6867.815280
Sum of electronic and thermal Free Energies= -6868.121411

............................................................

Cartesian Coordinates

............................................................

| | | | |
|---|---|---|---|
| C | -2.887583 | -6.561938 | -2.429081 |
| C | -2.915362 | -5.713640 | -1.150719 |
| S | -2.438013 | -3.950270 | -1.479385 |
| C | 0.888445 | -6.200051 | -2.951413 |
| C | 1.423531 | -5.958978 | -4.369036 |
| C | 1.689537 | -4.475309 | -4.673384 |
| C | 0.368806 | -3.708738 | -4.836605 |
| O | -0.524443 | -4.157570 | -5.544609 |
| N | 0.287711 | -2.516221 | -4.184979 |
| C | 1.015356 | -5.046936 | 0.656249 |
| C | 0.429708 | -4.383181 | 1.911158 |
| C | 1.398494 | -3.391485 | 2.538521 |
| O | 2.532188 | -3.778709 | 2.896624 |
| O | 1.030754 | -2.156304 | 2.689301 |
| C | 5.764056 | -9.882233 | 0.406528 |
| C | 4.975429 | -8.599442 | 0.104306 |
| C | 5.759384 | -7.639355 | -0.803601 |

| | | | |
|---|---|---|---|
| C | 4.984143 | -6.423271 | -1.324077 |
| N | 4.697744 | -5.440837 | -0.274081 |
| C | 4.292503 | -4.173614 | -0.501215 |
| N | 4.195770 | -3.713692 | -1.758633 |
| N | 4.051315 | -3.352996 | 0.522894 |
| N | -0.359832 | 2.633253 | 3.714677 |
| C | -1.539504 | 2.253594 | 4.521199 |
| C | -7.919841 | 2.806730 | -0.534604 |
| C | -7.542580 | 1.655464 | -1.467242 |
| S | -5.748535 | 1.570121 | -1.947748 |
| C | -9.034393 | -0.604613 | 3.807068 |
| C | -7.556543 | -0.206895 | 3.743303 |
| S | -6.400485 | -1.660435 | 3.652015 |
| C | 8.979519 | 4.647183 | -3.096875 |
| C | 7.759921 | 3.729452 | -3.221280 |
| C | 7.935988 | 2.646127 | -4.296156 |
| C | 6.777737 | 1.644956 | -4.373637 |
| N | 6.700792 | 0.868925 | -3.134545 |
| C | 5.628160 | 0.180156 | -2.689906 |
| N | 4.544538 | 0.031191 | -3.448429 |
| N | 5.660098 | -0.383251 | -1.474124 |
| C | 1.441474 | 2.750097 | -2.434568 |
| C | 1.248313 | 4.250332 | -2.240414 |
| O | 1.672540 | 5.082941 | -3.041983 |
| C | 0.976656 | 2.321708 | -3.840454 |
| N | 0.562124 | 4.616517 | -1.115687 |
| C | 0.335121 | 6.009597 | -0.790759 |
| C | -0.777571 | 6.176517 | 0.252862 |
| O | -2.040108 | 5.680905 | -0.219510 |
| C | -0.915864 | 7.627271 | 0.712588 |
| C | 4.471980 | 9.922829 | 1.854404 |
| C | 4.311537 | 8.650447 | 2.703717 |
| C | 2.880300 | 8.085664 | 2.699903 |
| C | 2.675443 | 6.810243 | 3.541969 |
| N | 3.463350 | 5.670086 | 3.054008 |
| C | 3.011044 | 4.665756 | 2.260953 |
| N | 3.852012 | 4.148136 | 1.357375 |
| N | 1.771630 | 4.191029 | 2.381246 |
| C | -2.780724 | 6.823972 | -4.842271 |
| C | -3.176340 | 5.413628 | -5.305407 |
| C | -2.606049 | 4.266634 | -4.449650 |
| C | -3.243297 | 4.136320 | -3.057052 |
| C | -2.586881 | 3.030721 | -2.228023 |
| N | -3.157305 | 2.962526 | -0.840455 |
| C | 6.238355 | -1.968375 | 8.346544 |
| C | 6.395129 | -0.937091 | 7.222379 |
| C | 5.557834 | -1.210126 | 6.002915 |
| N | 5.572947 | -0.349416 | 4.919694 |
| C | 4.698458 | -2.258675 | 5.752081 |
| C | 4.735301 | -0.879687 | 4.046837 |
| N | 4.178362 | -2.034842 | 4.495885 |
| C | -0.172414 | 1.274955 | 0.627528 |
| O | 0.538619 | 2.343959 | 0.631481 |
| O | -1.404344 | 1.279708 | 0.880632 |
| C | 0.668618 | 0.058965 | 0.329238 |
| O | -0.653898 | -1.291642 | 1.166072 |

| | | | |
|---|---|---|---|
| C | 1.007039 | -0.358277 | -0.938724 |
| C | 2.313066 | -0.907828 | -1.198354 |
| O | 3.269256 | -0.763993 | -0.321593 |
| O | 2.572740 | -1.477559 | -2.331481 |
| Fe | -3.854593 | -2.696222 | -0.362796 |
| Fe | -5.365537 | -1.471586 | 1.787214 |
| Fe | -5.134904 | -0.153822 | -0.640773 |
| Fe | -2.467351 | -0.428461 | 1.189454 |
| S | -4.627645 | 0.577841 | 1.427067 |
| S | -2.989559 | -0.718894 | -1.043137 |
| S | -3.424768 | -2.484602 | 1.854287 |
| S | -6.117611 | -2.192242 | -0.074686 |
| H | 0.305214 | -0.290656 | -1.762163 |
| H | -0.754423 | -1.914354 | 0.425916 |
| H | -1.728569 | 7.716364 | 1.437823 |
| H | 0.006776 | 7.989574 | 1.177250 |
| H | -1.137944 | 8.286180 | -0.136174 |
| H | -2.345965 | 6.270476 | -0.922987 |
| H | -3.177673 | 7.585420 | -5.521276 |
| H | -2.828261 | 5.276215 | -6.336415 |
| H | -4.270411 | 5.329626 | -5.341915 |
| H | -1.520703 | 4.405501 | -4.347913 |
| H | -2.749352 | 3.320483 | -4.986621 |
| H | -4.313921 | 3.923906 | -3.151394 |
| H | -3.145386 | 5.087464 | -2.519316 |
| H | -1.516678 | 3.211178 | -2.120238 |
| H | -2.743515 | 2.044189 | -2.670421 |
| H | -1.690671 | 6.936732 | -4.812438 |
| H | -3.168838 | 7.051172 | -3.842291 |
| H | 4.215074 | 9.735008 | 0.806312 |
| H | 3.820523 | 10.724298 | 2.218540 |
| H | 5.502402 | 10.288256 | 1.887814 |
| H | 4.613461 | 8.860149 | 3.738632 |
| H | 5.013491 | 7.891786 | 2.327948 |
| H | 2.565997 | 7.882469 | 1.667279 |
| H | 2.193255 | 8.847910 | 3.089820 |
| H | 1.623257 | 6.519901 | 3.531444 |
| H | 2.949058 | 6.991913 | 4.587056 |
| H | 8.820215 | 5.414154 | -2.333053 |
| H | 9.877019 | 4.082698 | -2.820096 |
| H | 9.189653 | 5.159809 | -4.041811 |
| H | 6.867355 | 4.322896 | -3.457007 |
| H | 7.559854 | 3.260937 | -2.248656 |
| H | 8.872232 | 2.094141 | -4.127581 |
| H | 8.038973 | 3.115266 | -5.281083 |
| H | 6.913730 | 0.977470 | -5.234960 |
| H | 5.833215 | 2.186260 | -4.509068 |
| H | 5.192534 | -10.551412 | 1.055961 |
| H | 5.995357 | -10.426774 | -0.515375 |
| H | 6.711765 | -9.658513 | 0.908808 |
| H | 0.310341 | -5.779379 | 0.256228 |
| H | 1.196842 | -4.312860 | -0.133925 |
| H | 1.952150 | -5.560866 | 0.890661 |
| H | 5.201862 | -2.017988 | 8.697140 |
| H | 6.523544 | -2.970793 | 8.008637 |
| H | 6.869794 | -1.712634 | 9.202992 |

| | | | |
|---|---|---|---|
| H | 0.611903 | -7.248989 | -2.803022 |
| H | 1.641348 | -5.943547 | -2.194392 |
| H | 0.007920 | -5.584459 | -2.749170 |
| H | -3.136027 | -7.606387 | -2.199918 |
| H | -3.615584 | -6.187166 | -3.154848 |
| H | -1.907046 | -6.539915 | -2.908112 |
| H | -9.233990 | -1.256930 | 4.663341 |
| H | -9.662478 | 0.289725 | 3.905094 |
| H | -9.333843 | -1.138894 | 2.900231 |
| H | -7.380192 | 2.728194 | 0.414358 |
| H | -7.680820 | 3.776544 | -0.985149 |
| H | -8.995958 | 2.787422 | -0.321077 |
| H | 4.025961 | -8.858124 | -0.381610 |
| H | 4.708552 | -8.106618 | 1.049238 |
| H | 6.094009 | -8.193148 | -1.689361 |
| H | 6.671682 | -7.295512 | -0.298059 |
| H | 4.049217 | -6.750510 | -1.799260 |
| H | 5.598461 | -5.935064 | -2.090626 |
| H | 6.146007 | 0.062423 | 7.603786 |
| H | 7.450757 | -0.877298 | 6.923154 |
| H | 1.434025 | -0.090616 | 1.087363 |
| H | 0.085798 | 6.550129 | -1.711975 |
| H | 0.308323 | 3.897172 | -0.441298 |
| H | 1.252382 | 6.482097 | -0.403387 |
| H | 2.504185 | 2.552732 | -2.358494 |
| H | 0.923356 | 2.167065 | -1.682182 |
| H | 2.343967 | -6.538144 | -4.527241 |
| H | 0.694091 | -6.310597 | -5.105302 |
| H | 2.313632 | -4.015525 | -3.896974 |
| H | 2.230277 | -4.386997 | -5.624388 |
| H | -7.805229 | 0.696106 | -1.007700 |
| H | -8.101348 | 1.722732 | -2.406557 |
| H | -7.362346 | 0.437403 | 2.882972 |
| H | -7.273380 | 0.352101 | 4.642064 |
| H | -3.915611 | -5.734163 | -0.705870 |
| H | -2.222250 | -6.123197 | -0.406545 |
| H | 3.508055 | -2.622031 | 3.985173 |
| H | 4.490781 | -0.470501 | 3.074701 |
| H | 4.411468 | -3.116867 | 6.339611 |
| H | 7.514666 | 0.877644 | -2.538428 |
| H | 4.741166 | -0.654712 | -1.027046 |
| H | 3.741626 | -0.545557 | -3.097186 |
| H | 4.532141 | 0.375204 | -4.394373 |
| H | 4.462289 | 5.816605 | 3.017594 |
| H | 4.620038 | 4.722999 | 1.045579 |
| H | 1.425127 | 3.488351 | 1.718516 |
| H | 1.237190 | 4.193856 | 3.252275 |
| H | 4.789665 | -5.724580 | 0.690443 |
| H | 3.642516 | -2.856304 | -1.954339 |
| H | 4.293087 | -4.361748 | -2.522984 |
| H | 3.735713 | -2.396201 | 0.304137 |
| H | -0.633443 | -2.101496 | -4.129126 |
| H | 0.974364 | -2.236972 | -3.490924 |
| H | -2.673616 | 2.250836 | -0.266243 |
| H | -3.041494 | 3.870865 | -0.374492 |
| H | -2.304699 | 3.029644 | 4.428135 |

| | | | |
|---|---|---|---|
| H | -1.993565 | 1.299642 | 4.223749 |
| H | 0.326512 | 1.883489 | 3.799139 |
| H | -0.647275 | 2.603921 | 2.737229 |
| H | -1.256989 | 2.189464 | 5.576263 |
| H | 0.164595 | -1.840226 | 2.074329 |
| H | 1.223848 | 1.269603 | -4.016845 |
| H | -4.168342 | 2.671504 | -0.901771 |
| H | 3.596558 | 3.343447 | 0.741075 |
| H | -0.549432 | 5.542279 | 1.117281 |
| H | -0.527897 | -3.908640 | 1.686196 |
| H | 0.238192 | -5.148350 | 2.672729 |
| H | 6.401510 | -0.110208 | -0.846242 |
| H | 3.750255 | -3.714616 | 1.432266 |
| O | 3.112416 | 2.232466 | -0.462499 |
| H | 2.165017 | 2.211478 | -0.194095 |
| H | 3.379312 | 1.306233 | -0.291380 |
| H | 1.467173 | 2.934361 | -4.601998 |
| H | -0.103799 | 2.428068 | -3.975177 |

---

**$A_{M2}$**

---

Number of imaginary frequencies: 7 Electronic energy: HF=-6796.0743519
Zero-point correction= 1.766819 (Hartree/Particle)
Thermal correction to Energy= 1.889129
Thermal correction to Enthalpy= 1.890073
Thermal correction to Gibbs Free Energy= 1.574570
Sum of electronic and zero-point Energies= -6794.307533
Sum of electronic and thermal Energies= -6794.185223
Sum of electronic and thermal Enthalpies= -6794.184279
Sum of electronic and thermal Free Energies= -6794.499782

..........................................................

Cartesian Coordinates

..........................................................

| | | | |
|---|---|---|---|
| C | -2.462846 | -6.259235 | -3.804472 |
| C | -2.564787 | -5.706473 | -2.375126 |
| S | -2.297351 | -3.868006 | -2.331939 |
| C | 1.236766 | -5.432165 | -4.350559 |
| C | 2.771730 | -5.426616 | -4.391332 |
| C | 3.425836 | -4.165303 | -3.804380 |
| C | 3.221822 | -2.920114 | -4.670037 |
| O | 3.175332 | -2.968164 | -5.889300 |
| N | 3.181836 | -1.727433 | -3.980321 |
| C | 1.390325 | -4.845536 | -0.610905 |
| C | 1.298171 | -5.828269 | 0.561390 |
| C | 6.578456 | -9.066873 | -1.732472 |
| C | 6.206925 | -7.604439 | -1.480378 |
| C | 7.335555 | -6.827683 | -0.789949 |
| C | 7.048643 | -5.337206 | -0.600002 |
| N | 5.948510 | -5.115716 | 0.347312 |
| C | 5.201785 | -4.003226 | 0.413895 |
| N | 5.471048 | -2.931948 | -0.327422 |
| N | 4.173931 | -3.934270 | 1.290670 |
| N | -0.610932 | 2.151724 | 3.719720 |
| C | -1.715842 | 1.464860 | 4.418795 |
| C | -8.313371 | 2.132458 | -0.335503 |

| | | | |
|---|---|---|---|
| C | -7.865444 | 1.127249 | -1.396043 |
| S | -6.070520 | 1.214966 | -1.873345 |
| C | -8.915495 | -2.006961 | 3.437058 |
| C | -7.477966 | -1.544557 | 3.681640 |
| S | -6.193231 | -2.829934 | 3.292421 |
| C | 8.425845 | 6.192892 | -2.401149 |
| C | 7.389139 | 5.069371 | -2.329022 |
| C | 8.026646 | 3.683893 | -2.503118 |
| C | 7.038108 | 2.516075 | -2.493873 |
| N | 6.391480 | 2.381855 | -1.186009 |
| C | 5.557289 | 1.396454 | -0.834840 |
| N | 5.390901 | 0.331513 | -1.634446 |
| N | 4.983241 | 1.445712 | 0.384539 |
| C | 0.926614 | 3.337788 | -2.439270 |
| C | 0.735291 | 4.758146 | -1.897238 |
| O | 1.197687 | 5.751199 | -2.458146 |
| C | 2.369412 | 2.886878 | -2.731053 |
| O | 3.053017 | 2.495628 | -1.537622 |
| C | 3.201933 | 3.864208 | -3.563755 |
| N | -0.067162 | 4.862092 | -0.798752 |
| C | -0.429921 | 6.171912 | -0.274695 |
| C | -1.502096 | 6.057050 | 0.815751 |
| O | -2.731499 | 5.514683 | 0.302621 |
| C | -1.758018 | 7.392692 | 1.510527 |
| C | 3.402771 | 10.034016 | 2.858278 |
| C | 3.542912 | 8.548483 | 3.217684 |
| C | 2.246833 | 7.754516 | 2.990377 |
| C | 2.318591 | 6.264484 | 3.364934 |
| N | 3.267006 | 5.525220 | 2.531891 |
| C | 3.048458 | 4.385195 | 1.805969 |
| N | 3.827096 | 4.029674 | 0.797209 |
| N | 2.026201 | 3.573157 | 2.176891 |
| C | -3.767527 | 7.277610 | -4.072761 |
| C | -4.001589 | 5.920649 | -4.755043 |
| C | -3.296056 | 4.722047 | -4.091120 |
| C | -3.886819 | 4.309241 | -2.733273 |
| C | -3.100471 | 3.167071 | -2.087679 |
| N | -3.619031 | 2.844606 | -0.714855 |
| C | 6.583789 | -2.472499 | 7.345560 |
| C | 5.035136 | -2.390448 | 7.443779 |
| C | 4.194944 | -2.505538 | 6.189796 |
| C | 3.046281 | -3.219052 | 5.950365 |
| C | 3.384576 | -1.951843 | 4.156448 |
| N | 2.569281 | -2.856848 | 4.698008 |
| C | -0.591891 | 1.162991 | 0.641201 |
| O | 0.060307 | 2.232590 | 0.555201 |
| O | -1.853161 | 1.087215 | 0.778330 |
| C | 0.213531 | -0.156102 | 0.686754 |
| O | -0.653754 | -1.307000 | 0.624421 |
| C | 1.338109 | -0.219367 | -0.336652 |
| C | 2.524211 | -1.102069 | 0.038067 |
| O | 2.734257 | -1.437442 | 1.238163 |
| O | 3.312379 | -1.413928 | -0.921993 |
| Fe | -3.717105 | -3.010631 | -0.897228 |
| Fe | -5.311896 | -2.282255 | 1.432040 |
| Fe | -5.345032 | -0.608336 | -0.813955 |

| | | | |
|---|---|---|---|
| Fe | -2.679397 | -0.744948 | 0.929714 |
| S | -4.887958 | -0.119913 | 1.335238 |
| S | -3.148920 | -0.861906 | -1.309745 |
| S | -3.239639 | -3.006427 | 1.338785 |
| S | -6.020363 | -2.818728 | -0.504011 |
| H | 1.740537 | 0.790197 | -0.497031 |
| H | 0.957827 | -0.543378 | -1.312436 |
| H | -0.820054 | -1.579519 | -0.300365 |
| H | 4.164935 | 3.403883 | -3.810789 |
| H | 2.693366 | 4.117928 | -4.498283 |
| H | 3.369780 | 4.800131 | -3.027620 |
| H | -2.541690 | 7.281608 | 2.264019 |
| H | -0.854542 | 7.766995 | 2.001510 |
| H | -2.079112 | 8.153316 | 0.787915 |
| H | -3.165985 | 6.207078 | -0.214923 |
| H | -4.260230 | 8.082254 | -4.626977 |
| H | -3.652713 | 5.988309 | -5.792395 |
| H | -5.078956 | 5.717866 | -4.812309 |
| H | -2.228610 | 4.957957 | -3.970120 |
| H | -3.348019 | 3.861678 | -4.770101 |
| H | -4.931814 | 4.002710 | -2.856133 |
| H | -3.877207 | 5.170033 | -2.053592 |
| H | -2.050020 | 3.439472 | -1.971887 |
| H | -3.171930 | 2.245127 | -2.669357 |
| H | -2.698204 | 7.514211 | -4.021282 |
| H | -4.166600 | 7.299192 | -3.051787 |
| H | 3.132214 | 10.163037 | 1.804777 |
| H | 2.625561 | 10.513658 | 3.462722 |
| H | 4.339118 | 10.572224 | 3.031653 |
| H | 3.848205 | 8.449025 | 4.267898 |
| H | 4.360712 | 8.121544 | 2.618686 |
| H | 1.939916 | 7.842019 | 1.939822 |
| H | 1.447338 | 8.206336 | 3.591660 |
| H | 1.330392 | 5.815478 | 3.235238 |
| H | 2.583754 | 6.160398 | 4.426486 |
| H | 7.956359 | 7.173067 | -2.278462 |
| H | 9.186069 | 6.085652 | -1.619479 |
| H | 8.941264 | 6.191163 | -3.367316 |
| H | 6.623461 | 5.215041 | -3.101090 |
| H | 6.860302 | 5.132435 | -1.366630 |
| H | 8.787827 | 3.516451 | -1.729209 |
| H | 8.555544 | 3.651609 | -3.462425 |
| H | 7.579707 | 1.588224 | -2.716709 |
| H | 6.274463 | 2.662523 | -3.268966 |
| H | 5.761655 | -9.600191 | -2.226444 |
| H | 7.462956 | -9.146019 | -2.373472 |
| H | 6.798719 | -9.590903 | -0.795768 |
| H | 1.910715 | -5.295983 | -1.459075 |
| H | 0.392265 | -4.552257 | -0.952647 |
| H | 1.925429 | -3.929999 | -0.339642 |
| H | 6.910871 | -3.429233 | 6.929757 |
| H | 7.002147 | -1.667688 | 6.731518 |
| H | 7.002504 | -2.371648 | 8.348769 |
| H | 0.851034 | -6.365085 | -4.772762 |
| H | 0.844732 | -5.334888 | -3.336309 |
| H | 0.827662 | -4.608924 | -4.944216 |

| | | | |
|---|---|---|---|
| H | -2.588985 | -7.348812 | -3.800952 |
| H | -3.238856 | -5.826225 | -4.442638 |
| H | -1.493317 | -6.025579 | -4.251010 |
| H | -9.148815 | -2.906268 | 4.015546 |
| H | -9.618518 | -1.218662 | 3.732319 |
| H | -9.077529 | -2.235515 | 2.379383 |
| H | -7.770019 | 1.977365 | 0.601832 |
| H | -8.139614 | 3.161986 | -0.666851 |
| H | -9.385313 | 2.018164 | -0.134115 |
| H | 5.967715 | -7.114906 | -2.433644 |
| H | 5.291585 | -7.554416 | -0.876676 |
| H | 8.252598 | -6.904685 | -1.385763 |
| H | 7.574750 | -7.281171 | 0.182863 |
| H | 6.758155 | -4.902890 | -1.563603 |
| H | 7.950194 | -4.817231 | -0.249234 |
| H | 4.679050 | -3.172961 | 8.120697 |
| H | 4.766691 | -1.440765 | 7.925666 |
| H | 0.659533 | -0.201853 | 1.686852 |
| H | -0.785286 | 6.801303 | -1.099423 |
| H | -0.274221 | 4.023655 | -0.257468 |
| H | 0.452047 | 6.682120 | 0.139301 |
| H | 0.465382 | 2.603582 | -1.777217 |
| H | 0.374152 | 3.321212 | -3.387970 |
| H | 3.158051 | -6.299381 | -3.848036 |
| H | 3.109108 | -5.523601 | -5.428302 |
| H | 3.079126 | -3.975864 | -2.782966 |
| H | 4.513995 | -4.318530 | -3.743982 |
| H | -8.065306 | 0.104264 | -1.056617 |
| H | -8.427623 | 1.268125 | -2.324567 |
| H | -7.246902 | -0.650906 | 3.098114 |
| H | -7.331035 | -1.298213 | 4.739138 |
| H | -3.547703 | -5.931971 | -1.949782 |
| H | -1.812871 | -6.175876 | -1.730829 |
| H | 1.715633 | -3.189688 | 4.234346 |
| H | 3.260228 | -1.509491 | 3.170100 |
| H | 2.526835 | -3.930932 | 6.571884 |
| H | 6.514770 | 3.124869 | -0.514125 |
| H | 4.339510 | 0.689399 | 0.578433 |
| H | 4.537466 | -0.224372 | -1.501954 |
| H | 5.737258 | 0.378495 | -2.580593 |
| H | 4.134117 | 5.998638 | 2.327549 |
| H | 4.417890 | 4.815448 | 0.529172 |
| H | 1.578593 | 2.987548 | 1.472755 |
| H | 1.470384 | 3.752264 | 3.000983 |
| H | 5.802006 | -5.813342 | 1.063694 |
| H | 4.675796 | -2.277573 | -0.534681 |
| H | 6.284393 | -2.922381 | -0.921628 |
| H | 3.618783 | -3.063123 | 1.261757 |
| H | 2.884882 | -0.934372 | -4.533482 |
| H | 2.955744 | -1.708841 | -2.991595 |
| H | -3.032800 | 2.133136 | -0.236213 |
| H | -3.598764 | 3.692378 | -0.134278 |
| H | -2.510189 | 2.186282 | 4.629722 |
| H | -2.165711 | 0.632630 | 3.858608 |
| H | 0.113129 | 1.467372 | 3.508710 |
| H | -0.950651 | 2.451574 | 2.808622 |

| | | | |
|---|---|---|---|
| H | -1.359081 | 1.083708 | 5.381219 |
| H | 2.278556 | 1.950992 | -3.298045 |
| H | -4.588533 | 2.445161 | -0.799802 |
| H | 3.108460 | 3.229210 | -0.889075 |
| H | -1.173167 | 5.317804 | 1.556648 |
| H | 0.707460 | -6.708263 | 0.288208 |
| H | 2.287326 | -6.214156 | 0.860054 |
| H | 4.605593 | 2.388389 | 0.659761 |
| H | 3.637944 | -4.780176 | 1.444176 |
| H | 0.821876 | -5.360230 | 1.427487 |
| O | -0.200280 | -3.590686 | 2.563102 |
| H | -0.238762 | -2.937730 | 1.843375 |
| H | -1.141143 | -3.811955 | 2.670444 |
| N | 4.369120 | -1.729740 | 5.042349 |
| H | 5.127429 | -1.078545 | 4.888852 |

---

**$B_{M2}$**

---

Number of imaginary frequencies: 5 Electronic energy: HF=-6796.0406602
Zero-point correction= 1.764570 (Hartree/Particle)
Thermal correction to Energy= 1.887596
Thermal correction to Enthalpy= 1.888540
Thermal correction to Gibbs Free Energy= 1.571910
Sum of electronic and zero-point Energies= -6794.276090
Sum of electronic and thermal Energies= -6794.153064
Sum of electronic and thermal Enthalpies= -6794.152120
Sum of electronic and thermal Free Energies= -6794.468750

.........................................................

Cartesian Coordinates

.........................................................

| | | | |
|---|---|---|---|
| C | -2.315738 | -6.446402 | -3.555607 |
| C | -2.452923 | -5.840779 | -2.152169 |
| S | -2.227230 | -3.998779 | -2.192157 |
| C | 1.375048 | -5.576262 | -4.094342 |
| C | 2.908668 | -5.530905 | -4.057972 |
| C | 3.492716 | -4.189264 | -3.591781 |
| C | 3.280189 | -3.052813 | -4.595395 |
| O | 3.237472 | -3.243891 | -5.802988 |
| N | 3.223420 | -1.797251 | -4.044291 |
| C | 1.478394 | -4.849682 | -0.377605 |
| C | 1.569256 | -5.711030 | 0.886289 |
| C | 6.748789 | -9.019237 | -1.289439 |
| C | 6.266649 | -7.571385 | -1.188224 |
| C | 7.377121 | -6.624951 | -0.715330 |
| C | 6.994561 | -5.145395 | -0.731233 |
| N | 5.963798 | -4.833910 | 0.264459 |
| C | 5.260390 | -3.686390 | 0.282316 |
| N | 5.458492 | -2.727796 | -0.610849 |
| N | 4.348834 | -3.467926 | 1.261812 |
| N | -0.760448 | 2.219723 | 3.526774 |
| C | -1.787930 | 1.587113 | 4.383211 |
| C | -8.344155 | 1.965125 | -0.459222 |
| C | -7.889641 | 0.895402 | -1.451310 |
| S | -6.109899 | 1.000969 | -1.977409 |
| C | -8.916945 | -2.042288 | 3.457675 |
| C | -7.496496 | -1.501456 | 3.641055 |

| | | | |
|---|---|---|---|
| S | -6.163949 | -2.768932 | 3.365730 |
| C | 8.345430 | 6.235109 | -2.503770 |
| C | 7.417717 | 5.027219 | -2.331461 |
| C | 8.184199 | 3.697314 | -2.344659 |
| C | 7.307424 | 2.443313 | -2.318214 |
| N | 6.600148 | 2.292444 | -1.040273 |
| C | 5.734869 | 1.287279 | -0.781858 |
| N | 5.408683 | 0.401890 | -1.706974 |
| N | 5.236086 | 1.169052 | 0.474308 |
| C | 0.896298 | 3.251477 | -2.512327 |
| C | 0.870402 | 4.621918 | -1.828887 |
| O | 1.726607 | 5.494876 | -2.007697 |
| C | 2.244970 | 2.513821 | -2.531117 |
| O | 2.708650 | 2.233621 | -1.200529 |
| C | 3.349250 | 3.237443 | -3.295311 |
| N | -0.233203 | 4.840481 | -1.058339 |
| C | -0.531114 | 6.139616 | -0.467975 |
| C | -1.599817 | 6.002466 | 0.624401 |
| O | -2.795205 | 5.389931 | 0.114296 |
| C | -1.921612 | 7.337619 | 1.290484 |
| C | 3.201834 | 10.180084 | 2.558604 |
| C | 3.266051 | 8.708421 | 2.986927 |
| C | 1.890412 | 8.020674 | 2.970295 |
| C | 1.912639 | 6.518314 | 3.297727 |
| N | 2.685397 | 5.779905 | 2.296539 |
| C | 2.355419 | 4.643020 | 1.653859 |
| N | 3.062351 | 4.345728 | 0.536733 |
| N | 1.427968 | 3.802735 | 2.094370 |
| C | -3.845778 | 7.046827 | -4.338061 |
| C | -4.208571 | 5.683504 | -4.948589 |
| C | -3.507472 | 4.466320 | -4.313261 |
| C | -4.001443 | 4.109498 | -2.901872 |
| C | -3.225788 | 2.936754 | -2.299341 |
| N | -3.671635 | 2.637552 | -0.896476 |
| C | 6.545022 | -2.095857 | 7.537954 |
| C | 5.007299 | -1.856115 | 7.445219 |
| C | 4.235110 | -2.087693 | 6.157479 |
| C | 3.093842 | -2.819832 | 5.940274 |
| C | 3.463271 | -1.677032 | 4.066583 |
| N | 2.641013 | -2.547773 | 4.656268 |
| C | -0.537032 | 1.130131 | 0.526414 |
| O | 0.051452 | 2.237084 | 0.370121 |
| O | -1.797984 | 1.036908 | 0.659694 |
| C | 0.330826 | -0.144134 | 0.648178 |
| O | -0.589308 | -1.305441 | 0.748530 |
| C | 1.371348 | -0.325859 | -0.389579 |
| C | 2.648106 | -0.824620 | -0.116374 |
| O | 3.198706 | -0.894451 | 1.078535 |
| O | 3.418808 | -1.183733 | -1.146719 |
| Fe | -3.657368 | -3.110239 | -0.800062 |
| Fe | -5.250451 | -2.296030 | 1.495853 |
| Fe | -5.307924 | -0.742176 | -0.823550 |
| Fe | -2.603843 | -0.779786 | 0.901419 |
| S | -4.832346 | -0.137703 | 1.293876 |
| S | -3.119304 | -0.981090 | -1.319964 |
| S | -3.175366 | -3.016104 | 1.431058 |

| | | | |
|---|---|---|---|
| S | -5.955732 | -2.943033 | -0.406593 |
| H | 2.169891 | 1.477633 | -0.837933 |
| H | 1.026281 | -0.409596 | -1.418329 |
| H | -0.568341 | -1.784418 | -0.100500 |
| H | 4.248505 | 2.615252 | -3.315727 |
| H | 3.037148 | 3.421204 | -4.327897 |
| H | 3.576293 | 4.203557 | -2.842153 |
| H | -2.693643 | 7.201944 | 2.051862 |
| H | -1.037216 | 7.772317 | 1.766815 |
| H | -2.287886 | 8.063425 | 0.554061 |
| H | -3.267327 | 6.048139 | -0.415256 |
| H | -4.353294 | 7.858318 | -4.868232 |
| H | -3.954837 | 5.702853 | -6.015131 |
| H | -5.295373 | 5.536970 | -4.901260 |
| H | -2.423182 | 4.649978 | -4.286062 |
| H | -3.656431 | 3.595442 | -4.963363 |
| H | -5.066893 | 3.854494 | -2.931802 |
| H | -3.889537 | 4.980285 | -2.243863 |
| H | -2.157620 | 3.158563 | -2.262910 |
| H | -3.373028 | 2.016323 | -2.868627 |
| H | -2.766815 | 7.232596 | -4.398631 |
| H | -4.139744 | 7.117099 | -3.283960 |
| H | 2.818053 | 10.278653 | 1.537688 |
| H | 2.544670 | 10.755386 | 3.219103 |
| H | 4.192529 | 10.641978 | 2.590385 |
| H | 3.698309 | 8.629459 | 3.992882 |
| H | 3.965174 | 8.189772 | 2.313775 |
| H | 1.422638 | 8.158605 | 1.986942 |
| H | 1.236384 | 8.510857 | 3.701364 |
| H | 0.893586 | 6.121389 | 3.302857 |
| H | 2.336483 | 6.350773 | 4.296409 |
| H | 7.780249 | 7.171536 | -2.493786 |
| H | 9.090215 | 6.286119 | -1.701591 |
| H | 8.885855 | 6.184046 | -3.454789 |
| H | 6.672446 | 5.017639 | -3.137088 |
| H | 6.849925 | 5.124240 | -1.396375 |
| H | 8.900180 | 3.660196 | -1.510747 |
| H | 8.790135 | 3.640093 | -3.256511 |
| H | 7.930872 | 1.556908 | -2.503352 |
| H | 6.563249 | 2.505617 | -3.120748 |
| H | 5.945331 | -9.679648 | -1.627430 |
| H | 7.576407 | -9.111634 | -2.000963 |
| H | 7.100129 | -9.393051 | -0.321059 |
| H | 1.986819 | -5.330112 | -1.216638 |
| H | 0.435684 | -4.691000 | -0.670069 |
| H | 1.927353 | -3.860464 | -0.236358 |
| H | 6.803268 | -3.134578 | 7.316466 |
| H | 7.106316 | -1.448317 | 6.856492 |
| H | 6.866072 | -1.868424 | 8.556326 |
| H | 1.030354 | -6.572023 | -4.390161 |
| H | 0.927238 | -5.337224 | -3.127245 |
| H | 0.985612 | -4.861127 | -4.825076 |
| H | -2.414414 | -7.538112 | -3.512091 |
| H | -3.092449 | -6.058669 | -4.221610 |
| H | -1.345696 | -6.205371 | -3.996931 |
| H | -9.110258 | -2.888000 | 4.124973 |

| | | | |
|---|---|---|---|
| H | -9.649302 | -1.256732 | 3.680281 |
| H | -9.075646 | -2.381346 | 2.429593 |
| H | -7.775845 | 1.896550 | 0.473733 |
| H | -8.208639 | 2.971125 | -0.871290 |
| H | -9.407515 | 1.836699 | -0.223328 |
| H | 5.903405 | -7.234595 | -2.167942 |
| H | 5.405833 | -7.512047 | -0.510112 |
| H | 8.252081 | -6.735825 | -1.366830 |
| H | 7.719014 | -6.906220 | 0.291364 |
| H | 6.598877 | -4.891495 | -1.721857 |
| H | 7.883459 | -4.524040 | -0.552032 |
| H | 4.514688 | -2.482569 | 8.195270 |
| H | 4.802324 | -0.821613 | 7.752107 |
| H | 0.820427 | -0.092385 | 1.629065 |
| H | -0.877736 | 6.832743 | -1.247021 |
| H | -0.689733 | 4.021100 | -0.669780 |
| H | 0.380815 | 6.580964 | -0.045720 |
| H | 0.151081 | 2.596812 | -2.058071 |
| H | 0.583864 | 3.426035 | -3.550893 |
| H | 3.286055 | -6.320503 | -3.393785 |
| H | 3.302543 | -5.740466 | -5.057823 |
| H | 3.096063 | -3.901938 | -2.612578 |
| H | 4.582496 | -4.291838 | -3.471780 |
| H | -8.048716 | -0.103861 | -1.030284 |
| H | -8.477011 | 0.948906 | -2.373351 |
| H | -7.305622 | -0.661862 | 2.969088 |
| H | -7.354809 | -1.143372 | 4.666903 |
| H | -3.436162 | -6.072171 | -1.730710 |
| H | -1.698622 | -6.263620 | -1.479165 |
| H | 1.790569 | -2.917661 | 4.212389 |
| H | 3.348608 | -1.277498 | 3.051593 |
| H | 2.560497 | -3.487043 | 6.598622 |
| H | 7.010998 | 2.744751 | -0.235790 |
| H | 4.553160 | 0.413961 | 0.665432 |
| H | 4.536556 | -0.201532 | -1.548880 |
| H | 5.826936 | 0.452381 | -2.622676 |
| H | 3.525009 | 6.229350 | 1.960354 |
| H | 3.230682 | 5.129297 | -0.086886 |
| H | 1.007530 | 3.116443 | 1.433712 |
| H | 0.968489 | 3.871181 | 2.997391 |
| H | 5.909336 | -5.421870 | 1.084067 |
| H | 4.631051 | -2.060131 | -0.803980 |
| H | 6.172520 | -2.850342 | -1.311386 |
| H | 3.847330 | -2.555746 | 1.188078 |
| H | 2.946552 | -1.066549 | -4.685489 |
| H | 3.041162 | -1.654075 | -3.051871 |
| H | -3.055365 | 1.945851 | -0.426744 |
| H | -3.653152 | 3.491538 | -0.327554 |
| H | -2.578680 | 2.312855 | 4.592006 |
| H | -2.258016 | 0.700545 | 3.937475 |
| H | -0.054137 | 1.513122 | 3.324103 |
| H | -1.193955 | 2.395082 | 2.621336 |
| H | -1.342218 | 1.303287 | 5.341876 |
| H | 2.059903 | 1.550269 | -3.026248 |
| H | -4.639136 | 2.215543 | -0.938290 |
| H | 2.814868 | 3.497052 | -0.010819 |

| | | | |
|---|---|---|---|
| H | -1.238041 | 5.290136 | 1.375370 |
| H | 1.082137 | -6.680183 | 0.737368 |
| H | 2.613244 | -5.938832 | 1.158423 |
| H | 5.039429 | 2.043124 | 0.943201 |
| H | 3.791512 | -4.264969 | 1.544853 |
| H | 1.078523 | -5.217393 | 1.730001 |
| O | -0.167320 | -3.506351 | 2.731373 |
| H | -0.256969 | -2.839222 | 2.026897 |
| H | -1.089866 | -3.784075 | 2.854766 |
| N | 4.429754 | -1.393113 | 4.958581 |
| H | 5.194060 | -0.759216 | 4.768934 |

----------------------------------------------------------------------------------------------------------------

**$C_{M2}$**

----------------------------------------------------------------------------------------------------------------

Number of imaginary frequencies: 6 Electronic energy: HF=-6796.0821845
Zero-point correction= 1.764406 (Hartree/Particle)
Thermal correction to Energy= 1.889008
Thermal correction to Enthalpy= 1.889952
Thermal correction to Gibbs Free Energy= 1.569579
Sum of electronic and zero-point Energies= -6794.317779
Sum of electronic and thermal Energies= -6794.193177
Sum of electronic and thermal Enthalpies= -6794.192233
Sum of electronic and thermal Free Energies= -6794.512606

............................................................

Cartesian Coordinates

............................................................

| | | | |
|---|---|---|---|
| C | -0.822268 | -6.931549 | -2.983965 |
| C | -1.308871 | -6.211894 | -1.722558 |
| S | -1.600780 | -4.409944 | -2.053662 |
| C | 2.596186 | -5.291875 | -3.525108 |
| C | 3.974819 | -4.873461 | -4.048289 |
| C | 4.201299 | -3.354947 | -4.005160 |
| C | 3.370786 | -2.608711 | -5.054655 |
| O | 3.276694 | -3.009189 | -6.205373 |
| N | 2.804856 | -1.434677 | -4.627169 |
| C | 2.474378 | -4.411777 | 0.174572 |
| C | 3.505225 | -5.523876 | 0.384282 |
| C | 8.551456 | -7.390582 | -0.549423 |
| C | 7.889358 | -6.024584 | -0.741793 |
| C | 8.831110 | -4.862340 | -0.403500 |
| C | 8.233361 | -3.477286 | -0.657848 |
| N | 7.136049 | -3.182292 | 0.270599 |
| C | 6.127157 | -2.307919 | 0.057377 |
| N | 5.845904 | -1.890401 | -1.197071 |
| N | 5.428816 | -1.847405 | 1.081035 |
| N | -1.247044 | 2.079717 | 3.835036 |
| C | -2.196781 | 1.292359 | 4.652725 |
| C | -8.596947 | 0.053192 | -0.254136 |
| C | -8.051775 | -0.565538 | -1.538099 |
| S | -6.276417 | -0.171411 | -1.922907 |
| C | -8.351046 | -3.845706 | 3.805435 |
| C | -7.148742 | -2.900264 | 3.783614 |
| S | -5.523661 | -3.762395 | 3.527297 |
| C | 6.615389 | 7.623816 | -2.880798 |
| C | 6.062959 | 6.193303 | -2.885284 |
| C | 7.021639 | 5.180478 | -3.530064 |

| | | | |
|---|---|---|---|
| C | 6.499986 | 3.738015 | -3.565587 |
| N | 6.361092 | 3.199488 | -2.207015 |
| C | 5.600753 | 2.147220 | -1.840420 |
| N | 4.965480 | 1.411415 | -2.753759 |
| N | 5.484150 | 1.844539 | -0.540702 |
| C | 0.154209 | 3.283947 | -2.295990 |
| C | -0.500400 | 4.624552 | -1.957740 |
| O | -0.308279 | 5.631921 | -2.636512 |
| C | 1.630424 | 3.231848 | -1.851560 |
| O | 1.736072 | 3.359252 | -0.422852 |
| C | 2.510124 | 4.301044 | -2.486695 |
| N | -1.279316 | 4.632719 | -0.841313 |
| C | -1.903842 | 5.856875 | -0.373702 |
| C | -2.971553 | 5.557874 | 0.688450 |
| O | -3.974354 | 4.651180 | 0.198917 |
| C | -3.615760 | 6.831950 | 1.228394 |
| C | 0.817572 | 10.702619 | 2.549144 |
| C | 1.075550 | 9.417860 | 3.351166 |
| C | -0.116168 | 8.443832 | 3.353767 |
| C | 0.142805 | 7.104283 | 4.070889 |
| N | 1.230972 | 6.339234 | 3.434748 |
| C | 1.069371 | 5.376066 | 2.492786 |
| N | 1.918667 | 5.360110 | 1.455685 |
| N | 0.118514 | 4.449011 | 2.603480 |
| C | -5.259147 | 5.860896 | -4.283225 |
| C | -5.226239 | 4.473894 | -4.941063 |
| C | -4.249092 | 3.474081 | -4.296529 |
| C | -4.616133 | 3.090175 | -2.855864 |
| C | -3.621734 | 2.103932 | -2.247412 |
| N | -3.952806 | 1.810816 | -0.809091 |
| C | 6.660411 | -0.374378 | 8.002076 |
| C | 6.628946 | -1.827322 | 7.486563 |
| C | 6.083482 | -1.980834 | 6.098365 |
| C | 6.699496 | -2.107452 | 4.877070 |
| C | 4.584616 | -2.130149 | 4.452101 |
| N | 5.766384 | -2.200022 | 3.863655 |
| C | -0.575003 | 1.072031 | 0.629094 |
| O | -0.544819 | 2.353342 | 0.761412 |
| O | -1.641325 | 0.421752 | 0.535604 |
| C | 0.805781 | 0.498608 | 0.547539 |
| O | -0.437688 | -2.470596 | 0.289497 |
| C | 1.323331 | -0.121546 | -0.521512 |
| C | 2.806498 | -0.234528 | -0.657361 |
| O | 3.543038 | 0.104645 | 0.320410 |
| O | 3.280493 | -0.601083 | -1.787050 |
| Fe | -3.397619 | -3.783522 | -0.871466 |
| Fe | -4.980483 | -2.996293 | 1.633475 |
| Fe | -5.207490 | -1.626045 | -0.682367 |
| Fe | -2.320191 | -1.513047 | 0.698956 |
| S | -4.533251 | -0.855807 | 1.304769 |
| S | -3.107683 | -1.623175 | -1.447587 |
| S | -2.889500 | -3.710934 | 1.370364 |
| S | -5.735538 | -3.750946 | -0.227608 |
| H | 0.990179 | 2.862764 | -0.015074 |
| H | 0.714955 | -0.455595 | -1.356011 |
| H | -0.564716 | -2.853656 | -0.616979 |

| | | | |
|---|---|---|---|
| H | 3.521412 | 4.239906 | -2.073124 |
| H | 2.564588 | 4.151955 | -3.569285 |
| H | 2.105491 | 5.300504 | -2.313338 |
| H | -4.369460 | 6.585742 | 1.980418 |
| H | -2.871626 | 7.491319 | 1.686274 |
| H | -4.102205 | 7.395800 | 0.422956 |
| H | -4.582009 | 5.146307 | -0.368364 |
| H | -5.914812 | 6.539341 | -4.837516 |
| H | -4.949379 | 4.590295 | -5.995560 |
| H | -6.235618 | 4.042570 | -4.939028 |
| H | -3.234996 | 3.899304 | -4.314729 |
| H | -4.217681 | 2.564317 | -4.908940 |
| H | -5.618308 | 2.647467 | -2.829616 |
| H | -4.633115 | 3.992817 | -2.234298 |
| H | -2.610933 | 2.514123 | -2.262020 |
| H | -3.622814 | 1.142180 | -2.764680 |
| H | -4.259541 | 6.311114 | -4.257595 |
| H | -5.636732 | 5.819642 | -3.254998 |
| H | 0.602082 | 10.478772 | 1.498783 |
| H | -0.036344 | 11.255860 | 2.953524 |
| H | 1.687760 | 11.364074 | 2.580494 |
| H | 1.334385 | 9.673603 | 4.386860 |
| H | 1.964725 | 8.926133 | 2.928259 |
| H | -0.426117 | 8.236985 | 2.320073 |
| H | -0.972562 | 8.923216 | 3.844196 |
| H | -0.756277 | 6.484798 | 4.063610 |
| H | 0.414460 | 7.270123 | 5.118367 |
| H | 5.904531 | 8.317716 | -2.422870 |
| H | 7.554599 | 7.688496 | -2.320329 |
| H | 6.812315 | 7.975640 | -3.898826 |
| H | 5.106426 | 6.169756 | -3.422832 |
| H | 5.837896 | 5.886538 | -1.854942 |
| H | 7.994648 | 5.198239 | -3.017812 |
| H | 7.227261 | 5.475819 | -4.565280 |
| H | 7.179876 | 3.109511 | -4.156126 |
| H | 5.513855 | 3.714793 | -4.042841 |
| H | 7.862118 | -8.201559 | -0.801079 |
| H | 9.436683 | -7.496603 | -1.186005 |
| H | 8.869814 | -7.536651 | 0.488825 |
| H | 2.817204 | -3.676465 | -0.561738 |
| H | 1.523229 | -4.812406 | -0.189177 |
| H | 2.281123 | -3.870642 | 1.105894 |
| H | 7.275638 | 0.255722 | 7.353016 |
| H | 5.656035 | 0.060772 | 8.036830 |
| H | 7.077291 | -0.336091 | 9.012694 |
| H | 2.487115 | -6.380323 | -3.549897 |
| H | 2.435530 | -4.960470 | -2.496137 |
| H | 1.796681 | -4.867088 | -4.140085 |
| H | -0.628099 | -7.990084 | -2.771011 |
| H | -1.573737 | -6.873579 | -3.777003 |
| H | 0.098629 | -6.483386 | -3.363860 |
| H | -8.245275 | -4.603571 | 4.587643 |
| H | -9.270044 | -3.280241 | 3.999499 |
| H | -8.459627 | -4.361714 | 2.846994 |
| H | -8.030652 | -0.292293 | 0.616042 |
| H | -8.539654 | 1.147023 | -0.282441 |

| | | | |
|---|---|---|---|
| H | -9.647539 | -0.228408 | -0.115361 |
| H | 7.558241 | -5.922079 | -1.784425 |
| H | 6.983713 | -5.959124 | -0.126694 |
| H | 9.743521 | -4.938234 | -1.007627 |
| H | 9.160559 | -4.932426 | 0.643354 |
| H | 7.853747 | -3.444251 | -1.682995 |
| H | 9.007678 | -2.702318 | -0.576034 |
| H | 7.645038 | -2.235005 | 7.492600 |
| H | 6.050937 | -2.451149 | 8.180411 |
| H | 1.494536 | 0.822016 | 1.324100 |
| H | -2.343791 | 6.379432 | -1.230951 |
| H | -1.314183 | 3.798474 | -0.258982 |
| H | -1.156985 | 6.543216 | 0.053771 |
| H | -0.381320 | 2.443467 | -1.849067 |
| H | 0.112427 | 3.168623 | -3.383347 |
| H | 4.759849 | -5.364973 | -3.458585 |
| H | 4.098848 | -5.202456 | -5.084905 |
| H | 3.989661 | -2.964208 | -3.003827 |
| H | 5.253961 | -3.134454 | -4.238905 |
| H | -8.133671 | -1.654875 | -1.503304 |
| H | -8.619674 | -0.219375 | -2.407933 |
| H | -7.265056 | -2.145415 | 2.997960 |
| H | -7.058009 | -2.363642 | 4.734254 |
| H | -2.243385 | -6.656564 | -1.366394 |
| H | -0.573892 | -6.310536 | -0.916356 |
| H | 1.069780 | -1.980501 | 1.700951 |
| H | 3.623026 | -2.164059 | 3.952173 |
| H | 7.761428 | -2.141013 | 4.677413 |
| H | 6.969856 | 3.586030 | -1.499926 |
| H | 4.806987 | 1.116293 | -0.221835 |
| H | 4.341894 | 0.637928 | -2.458599 |
| H | 5.197694 | 1.494961 | -3.730599 |
| H | 2.095894 | 6.855070 | 3.343178 |
| H | 2.370707 | 6.226765 | 1.205611 |
| H | -0.008869 | 3.746611 | 1.868683 |
| H | -0.294647 | 4.159589 | 3.491965 |
| H | 7.299133 | -3.418567 | 1.240747 |
| H | 4.950156 | -1.412237 | -1.356627 |
| H | 6.145483 | -2.472817 | -1.963941 |
| H | 4.652901 | -1.202030 | 0.896934 |
| H | 2.139105 | -1.021328 | -5.266064 |
| H | 2.697107 | -1.240736 | -3.636648 |
| H | -3.222352 | 1.235201 | -0.351664 |
| H | -4.048573 | 2.690598 | -0.280728 |
| H | -3.157241 | 1.813685 | 4.689079 |
| H | -2.379152 | 0.278669 | 4.273991 |
| H | -0.368558 | 1.564221 | 3.789606 |
| H | -1.587810 | 2.082603 | 2.874916 |
| H | -1.820898 | 1.219866 | 5.677543 |
| H | -0.674214 | -3.251842 | 0.826283 |
| H | 2.015907 | 2.241208 | -2.131785 |
| H | -4.836781 | 1.258021 | -0.771439 |
| H | 1.881953 | 4.629725 | 0.719911 |
| H | -2.500622 | 5.010763 | 1.513550 |
| H | 3.708550 | -6.063954 | -0.546279 |
| H | 4.456469 | -5.121938 | 0.752718 |

| | | | |
|---|---|---|---|
| H | 5.719285 | 2.556602 | 0.133941 |
| H | 5.680493 | -2.047152 | 2.071447 |
| H | 3.154367 | -6.257352 | 1.117533 |
| O | 1.399535 | -2.005826 | 2.615218 |
| H | 0.826619 | -2.656682 | 3.042093 |
| N | 4.729732 | -1.993351 | 5.798689 |
| H | 3.976190 | -1.944123 | 6.468838 |

---

**TS(A-B)$_{M2}$**

---

Number of imaginary frequencies: 6 Electronic energy: HF=-6796.031279
Zero-point correction= 1.760562 (Hartree/Particle)
Thermal correction to Energy= 1.883104
Thermal correction to Enthalpy= 1.884048
Thermal correction to Gibbs Free Energy= 1.569987
Sum of electronic and zero-point Energies= -6794.270717
Sum of electronic and thermal Energies= -6794.148175
Sum of electronic and thermal Enthalpies= -6794.147231
Sum of electronic and thermal Free Energies= -6794.461292

............................................................

Cartesian Coordinates

............................................................

| | | | |
|---|---|---|---|
| C | -2.464137 | -6.340650 | -3.625163 |
| C | -2.515462 | -5.757718 | -2.207481 |
| S | -2.258848 | -3.920361 | -2.230829 |
| C | 1.241664 | -5.545160 | -4.176167 |
| C | 2.776054 | -5.528086 | -4.131251 |
| C | 3.384947 | -4.211083 | -3.626022 |
| C | 3.217563 | -3.046002 | -4.604737 |
| O | 3.205494 | -3.204003 | -5.816715 |
| N | 3.170799 | -1.800751 | -4.022425 |
| C | 1.382956 | -4.866644 | -0.451593 |
| C | 1.429366 | -5.755358 | 0.795882 |
| C | 6.554826 | -9.139379 | -1.446512 |
| C | 6.081993 | -7.693112 | -1.283176 |
| C | 7.191753 | -6.779489 | -0.746630 |
| C | 6.833176 | -5.293571 | -0.715971 |
| N | 5.789824 | -4.996399 | 0.273195 |
| C | 5.112386 | -3.837084 | 0.323723 |
| N | 5.359625 | -2.843328 | -0.521325 |
| N | 4.179991 | -3.636194 | 1.286553 |
| N | -0.644614 | 2.233879 | 3.619659 |
| C | -1.712575 | 1.581470 | 4.407092 |
| C | -8.287448 | 2.162548 | -0.389761 |
| C | -7.872325 | 1.074610 | -1.379623 |
| S | -6.099105 | 1.141895 | -1.932728 |
| C | -8.925130 | -1.879037 | 3.481737 |
| C | -7.477458 | -1.408400 | 3.646378 |
| S | -6.211231 | -2.731688 | 3.322158 |
| C | 8.479716 | 6.090075 | -2.485674 |
| C | 7.490667 | 4.924375 | -2.380963 |
| C | 8.191612 | 3.559614 | -2.440708 |
| C | 7.253781 | 2.350014 | -2.423743 |
| N | 6.548122 | 2.236870 | -1.140721 |
| C | 5.673427 | 1.268249 | -0.829335 |
| N | 5.381115 | 0.297955 | -1.686145 |

| | | | |
|---|---|---|---|
| N | 5.147662 | 1.258772 | 0.424905 |
| C | 0.966799 | 3.270886 | -2.484279 |
| C | 0.970670 | 4.636563 | -1.794512 |
| O | 1.848343 | 5.489727 | -1.963880 |
| C | 2.270369 | 2.448968 | -2.413864 |
| O | 2.641928 | 2.138344 | -1.084346 |
| C | 3.424816 | 3.108907 | -3.177111 |
| N | -0.134605 | 4.876390 | -1.028098 |
| C | -0.384607 | 6.164394 | -0.396262 |
| C | -1.465818 | 6.034188 | 0.684076 |
| O | -2.689885 | 5.510574 | 0.142847 |
| C | -1.721524 | 7.352524 | 1.410265 |
| C | 3.454280 | 10.084598 | 2.655911 |
| C | 3.556593 | 8.583229 | 2.957626 |
| C | 2.187850 | 7.883131 | 2.959859 |
| C | 2.226101 | 6.372359 | 3.240756 |
| N | 2.951177 | 5.651443 | 2.192221 |
| C | 2.602547 | 4.498835 | 1.582139 |
| N | 3.265208 | 4.145153 | 0.459798 |
| N | 1.678162 | 3.680521 | 2.082185 |
| C | -3.701337 | 7.191490 | -4.234481 |
| C | -4.056329 | 5.845313 | -4.885389 |
| C | -3.395482 | 4.610929 | -4.241585 |
| C | -3.946994 | 4.244725 | -2.854390 |
| C | -3.188836 | 3.075092 | -2.225404 |
| N | -3.669490 | 2.788364 | -0.832096 |
| C | 6.555745 | -2.321843 | 7.465157 |
| C | 5.011374 | -2.121396 | 7.453984 |
| C | 4.206510 | -2.282815 | 6.177792 |
| C | 3.055612 | -2.995804 | 5.947527 |
| C | 3.410504 | -1.778994 | 4.120005 |
| N | 2.587749 | -2.665869 | 4.682746 |
| C | -0.620803 | 1.253253 | 0.569315 |
| O | -0.069798 | 2.383276 | 0.488136 |
| O | -1.879498 | 1.107491 | 0.670123 |
| C | 0.280006 | 0.001337 | 0.647380 |
| O | -0.577092 | -1.194284 | 0.676645 |
| C | 1.374055 | -0.073660 | -0.384359 |
| C | 2.557296 | -0.859046 | -0.060726 |
| O | 2.987901 | -1.025948 | 1.143230 |
| O | 3.285245 | -1.286250 | -1.059449 |
| Fe | -3.682542 | -3.039737 | -0.830248 |
| Fe | -5.264269 | -2.270105 | 1.465153 |
| Fe | -5.330607 | -0.651708 | -0.833203 |
| Fe | -2.631557 | -0.720841 | 0.887862 |
| S | -4.855302 | -0.106592 | 1.300146 |
| S | -3.141711 | -0.908754 | -1.333403 |
| S | -3.181959 | -2.970123 | 1.392507 |
| S | -5.974331 | -2.871757 | -0.449859 |
| H | 1.982079 | 1.160530 | -0.681116 |
| H | 0.991555 | -0.315931 | -1.380905 |
| H | -0.615866 | -1.584336 | -0.216030 |
| H | 4.294681 | 2.445961 | -3.171562 |
| H | 3.145918 | 3.291134 | -4.220616 |
| H | 3.693097 | 4.072142 | -2.738145 |
| H | -2.508948 | 7.224097 | 2.157004 |

| | | | |
|---|---|---|---|
| H | -0.820156 | 7.713923 | 1.915384 |
| H | -2.038831 | 8.131602 | 0.706229 |
| H | -3.094649 | 6.201117 | -0.401070 |
| H | -4.172859 | 8.020521 | -4.770882 |
| H | -3.758402 | 5.880172 | -5.940092 |
| H | -5.145955 | 5.712348 | -4.884995 |
| H | -2.311271 | 4.781392 | -4.169730 |
| H | -3.528972 | 3.749932 | -4.908080 |
| H | -5.009370 | 3.985918 | -2.928798 |
| H | -3.867808 | 5.112726 | -2.188102 |
| H | -2.122555 | 3.298400 | -2.163055 |
| H | -3.322643 | 2.151053 | -2.792312 |
| H | -2.618118 | 7.361388 | -4.244188 |
| H | -4.040850 | 7.247438 | -3.193363 |
| H | 3.008830 | 10.260975 | 1.671170 |
| H | 2.832899 | 10.594394 | 3.399637 |
| H | 4.441011 | 10.555989 | 2.666429 |
| H | 4.042821 | 8.431217 | 3.930290 |
| H | 4.220526 | 8.126444 | 2.208502 |
| H | 1.686611 | 8.050752 | 1.998005 |
| H | 1.554755 | 8.342867 | 3.728547 |
| H | 1.205344 | 5.981520 | 3.278357 |
| H | 2.687846 | 6.179916 | 4.218508 |
| H | 7.960878 | 7.051997 | -2.444700 |
| H | 9.210954 | 6.069340 | -1.669860 |
| H | 9.034400 | 6.052598 | -3.429053 |
| H | 6.756131 | 4.986216 | -3.193691 |
| H | 6.915432 | 5.014329 | -1.449023 |
| H | 8.916816 | 3.466799 | -1.619669 |
| H | 8.777805 | 3.495332 | -3.364553 |
| H | 7.835770 | 1.434970 | -2.599312 |
| H | 6.513964 | 2.444284 | -3.227301 |
| H | 5.751356 | -9.775126 | -1.828581 |
| H | 7.392832 | -9.204041 | -2.148763 |
| H | 6.887385 | -9.562402 | -0.491946 |
| H | 1.846051 | -5.367348 | -1.304619 |
| H | 0.349897 | -4.632366 | -0.727413 |
| H | 1.901603 | -3.913667 | -0.301931 |
| H | 6.831912 | -3.333448 | 7.156554 |
| H | 7.069885 | -1.609320 | 6.811643 |
| H | 6.916417 | -2.159963 | 8.482787 |
| H | 0.881467 | -6.526483 | -4.500068 |
| H | 0.791968 | -5.323383 | -3.205867 |
| H | 0.867923 | -4.804507 | -4.889653 |
| H | -2.578861 | -7.431303 | -3.595419 |
| H | -3.268697 | -5.928443 | -4.241457 |
| H | -1.515315 | -6.107590 | -4.114838 |
| H | -9.145448 | -2.727795 | 4.136717 |
| H | -9.614764 | -1.064417 | 3.734192 |
| H | -9.120093 | -2.189528 | 2.450856 |
| H | -7.705834 | 2.089130 | 0.534612 |
| H | -8.136408 | 3.161573 | -0.813079 |
| H | -9.349420 | 2.060067 | -0.135555 |
| H | 5.738591 | -7.306951 | -2.251788 |
| H | 5.209632 | -7.659930 | -0.617921 |
| H | 8.080592 | -6.877248 | -1.381167 |

| | | | |
|---|---|---|---|
| H | 7.506331 | -7.103256 | 0.256042 |
| H | 6.460397 | -4.997214 | -1.703824 |
| H | 7.728526 | -4.695307 | -0.496743 |
| H | 4.564405 | -2.814475 | 8.173302 |
| H | 4.791199 | -1.117670 | 7.841782 |
| H | 0.747462 | 0.022387 | 1.639827 |
| H | -0.697970 | 6.897797 | -1.152612 |
| H | -0.587135 | 4.061146 | -0.623831 |
| H | 0.540165 | 6.557474 | 0.046978 |
| H | 0.156983 | 2.659613 | -2.081372 |
| H | 0.727663 | 3.466805 | -3.538437 |
| H | 3.135203 | -6.342086 | -3.486771 |
| H | 3.172155 | -5.718315 | -5.134081 |
| H | 2.981008 | -3.937842 | -2.645984 |
| H | 4.470200 | -4.343263 | -3.493654 |
| H | -8.046686 | 0.082886 | -0.946280 |
| H | -8.474290 | 1.132247 | -2.291884 |
| H | -7.259719 | -0.566540 | 2.986135 |
| H | -7.299785 | -1.077731 | 4.675777 |
| H | -3.479381 | -5.978436 | -1.738201 |
| H | -1.735077 | -6.205525 | -1.581762 |
| H | 1.730024 | -3.008893 | 4.232155 |
| H | 3.289989 | -1.344210 | 3.125154 |
| H | 2.526122 | -3.686088 | 6.584833 |
| H | 6.873081 | 2.811414 | -0.376514 |
| H | 4.468045 | 0.522207 | 0.622121 |
| H | 4.500191 | -0.247323 | -1.538110 |
| H | 5.784626 | 0.305058 | -2.610403 |
| H | 3.771749 | 6.106401 | 1.819133 |
| H | 3.492694 | 4.926840 | -0.147963 |
| H | 1.167491 | 3.048890 | 1.431628 |
| H | 1.231726 | 3.826898 | 2.980147 |
| H | 5.687080 | -5.629072 | 1.054146 |
| H | 4.557132 | -2.165306 | -0.708286 |
| H | 6.081479 | -2.955217 | -1.215373 |
| H | 3.684138 | -2.726907 | 1.224360 |
| H | 2.904084 | -1.053662 | -4.649736 |
| H | 2.931612 | -1.689059 | -3.040261 |
| H | -3.067809 | 2.093285 | -0.344731 |
| H | -3.650677 | 3.647170 | -0.270538 |
| H | -2.476758 | 2.322025 | 4.659011 |
| H | -2.211518 | 0.750442 | 3.890327 |
| H | 0.037509 | 1.519698 | 3.369244 |
| H | -1.046889 | 2.514461 | 2.727213 |
| H | -1.297325 | 1.208267 | 5.348815 |
| H | 2.037909 | 1.502852 | -2.933722 |
| H | -4.639041 | 2.375229 | -0.892935 |
| H | 2.915987 | 3.299414 | -0.132701 |
| H | -1.147282 | 5.269385 | 1.402087 |
| H | 0.870691 | -6.683143 | 0.636224 |
| H | 2.457358 | -6.066477 | 1.046603 |
| H | 4.836269 | 2.171028 | 0.746183 |
| H | 3.605139 | -4.432862 | 1.534673 |
| H | 0.984209 | -5.246601 | 1.656010 |
| O | -0.214309 | -3.525793 | 2.683663 |
| H | -0.265543 | -2.863069 | 1.972736 |

| | | | |
|---|---|---|---|
| H | -1.150445 | -3.768204 | 2.785226 |
| N | 4.391447 | -1.539950 | 5.008001 |
| H | 5.156605 | -0.901414 | 4.837572 |

---

**TS(B-C)$_{M2}$**

---

Number of imaginary frequencies: 8 Electronic energy: HF=-6796.0295914
Zero-point correction= 1.758672 (Hartree/Particle)
Thermal correction to Energy= 1.879369
Thermal correction to Enthalpy= 1.880313
Thermal correction to Gibbs Free Energy= 1.573774
Sum of electronic and zero-point Energies= -6794.270919
Sum of electronic and thermal Energies= -6794.150222
Sum of electronic and thermal Enthalpies= -6794.149278
Sum of electronic and thermal Free Energies= -6794.455817

.............................................................
Cartesian Coordinates
.............................................................

| | | | |
|---|---|---|---|
| C | -2.393092 | -6.407744 | -3.677359 |
| C | -2.494927 | -5.898868 | -2.232772 |
| S | -2.197493 | -4.068984 | -2.148211 |
| C | 1.347833 | -5.648948 | -3.981564 |
| C | 2.878695 | -5.661430 | -3.897159 |
| C | 3.490562 | -4.304757 | -3.520750 |
| C | 3.312720 | -3.243484 | -4.609695 |
| O | 3.374278 | -3.514243 | -5.800617 |
| N | 3.157914 | -1.957823 | -4.151268 |
| C | 1.228549 | -4.879350 | -0.272335 |
| C | 1.402533 | -5.431836 | 1.146260 |
| C | 6.424829 | -9.206452 | -0.791810 |
| C | 5.989151 | -7.740232 | -0.778630 |
| C | 7.140297 | -6.796938 | -0.407327 |
| C | 6.805964 | -5.311493 | -0.542853 |
| N | 5.817822 | -4.874650 | 0.450329 |
| C | 5.129386 | -3.718160 | 0.370240 |
| N | 5.274316 | -2.896366 | -0.666632 |
| N | 4.304545 | -3.346828 | 1.364889 |
| N | -0.983105 | 2.330653 | 3.544169 |
| C | -2.156330 | 1.704429 | 4.194823 |
| C | -8.366978 | 2.207003 | -1.073036 |
| C | -7.660045 | 0.876151 | -1.336609 |
| S | -5.928683 | 1.030505 | -2.009333 |
| C | -9.310967 | -1.734371 | 2.839002 |
| C | -7.858766 | -1.294044 | 3.075035 |
| S | -6.640127 | -2.700454 | 3.009044 |
| C | 8.362203 | 5.845895 | -2.633180 |
| C | 7.489574 | 4.592097 | -2.522072 |
| C | 8.299866 | 3.295800 | -2.660362 |
| C | 7.461446 | 2.015235 | -2.621475 |
| N | 6.842913 | 1.822919 | -1.304769 |
| C | 5.867391 | 0.928928 | -1.033318 |
| N | 5.360084 | 0.153754 | -1.981995 |
| N | 5.435722 | 0.808013 | 0.238162 |
| C | 1.021062 | 3.208420 | -2.524993 |
| C | 0.966385 | 4.602412 | -1.891205 |
| O | 1.823095 | 5.472216 | -2.075943 |

| | | | |
|---|---|---|---|
| C | 2.380446 | 2.494187 | -2.484100 |
| O | 2.791748 | 2.243336 | -1.128193 |
| C | 3.504105 | 3.222783 | -3.212823 |
| N | -0.160437 | 4.843679 | -1.162043 |
| C | -0.452992 | 6.159677 | -0.610007 |
| C | -1.513898 | 6.062485 | 0.493900 |
| O | -2.722090 | 5.449901 | 0.010003 |
| C | -1.818138 | 7.417779 | 1.125887 |
| C | 3.188168 | 10.130787 | 2.614145 |
| C | 3.253160 | 8.683866 | 3.112961 |
| C | 1.880593 | 7.992267 | 3.100349 |
| C | 1.911977 | 6.494344 | 3.440350 |
| N | 2.694586 | 5.755561 | 2.446480 |
| C | 2.335699 | 4.660871 | 1.749504 |
| N | 3.033803 | 4.389770 | 0.626229 |
| N | 1.374601 | 3.836597 | 2.152423 |
| C | -3.479062 | 7.112329 | -4.699366 |
| C | -3.973271 | 5.732170 | -5.167662 |
| C | -3.313455 | 4.509961 | -4.492230 |
| C | -3.795951 | 4.216395 | -3.060075 |
| C | -3.085088 | 3.002874 | -2.451817 |
| N | -3.547775 | 2.726285 | -1.047740 |
| C | 5.841851 | -2.172210 | 7.930413 |
| C | 4.417672 | -1.586421 | 7.799514 |
| C | 3.766966 | -1.652054 | 6.443639 |
| N | 4.298947 | -1.081750 | 5.293937 |
| C | 2.546210 | -2.140143 | 6.041488 |
| C | 3.415352 | -1.236121 | 4.272898 |
| N | 2.338006 | -1.875604 | 4.699286 |
| C | -0.547255 | 1.216825 | 0.591044 |
| O | -0.028849 | 2.350279 | 0.368572 |
| O | -1.789454 | 1.047890 | 0.749159 |
| C | 0.464817 | 0.068854 | 0.726216 |
| O | -0.474178 | -1.243351 | 1.191427 |
| C | 1.293221 | -0.257409 | -0.397325 |
| C | 2.586984 | -0.818890 | -0.222354 |
| O | 3.201682 | -0.829924 | 0.927037 |
| O | 3.224924 | -1.274956 | -1.275905 |
| Fe | -3.429586 | -3.181625 | -0.612900 |
| Fe | -5.213252 | -2.400720 | 1.445638 |
| Fe | -5.097664 | -0.789892 | -0.924608 |
| Fe | -2.573241 | -0.763081 | 1.077434 |
| S | -4.827451 | -0.213719 | 1.242048 |
| S | -2.860230 | -1.069829 | -1.170239 |
| S | -3.096944 | -2.998496 | 1.653034 |
| S | -5.688060 | -3.070050 | -0.519831 |
| H | 2.206505 | 1.523100 | -0.779809 |
| H | 0.835080 | -0.339034 | -1.378745 |
| H | -0.458735 | -1.847308 | 0.421306 |
| H | 4.414784 | 2.619356 | -3.176016 |
| H | 3.234876 | 3.376692 | -4.262312 |
| H | 3.690760 | 4.203013 | -2.773081 |
| H | -2.584991 | 7.310577 | 1.897077 |
| H | -0.925982 | 7.856472 | 1.583101 |
| H | -2.182857 | 8.127068 | 0.372668 |
| H | -3.215970 | 6.113595 | -0.491880 |

| | | | |
|---|---|---|---|
| H | -3.985842 | 7.911826 | -5.247869 |
| H | -3.797019 | 5.654264 | -6.247021 |
| H | -5.061498 | 5.667682 | -5.038456 |
| H | -2.221834 | 4.643844 | -4.490559 |
| H | -3.513146 | 3.624010 | -5.107628 |
| H | -4.876675 | 4.032032 | -3.064210 |
| H | -3.611224 | 5.089677 | -2.422638 |
| H | -2.006367 | 3.164767 | -2.419985 |
| H | -3.285937 | 2.093699 | -3.022913 |
| H | -2.401334 | 7.223321 | -4.865635 |
| H | -3.671059 | 7.278184 | -3.632290 |
| H | 2.827272 | 10.177126 | 1.581136 |
| H | 2.511363 | 10.731728 | 3.230621 |
| H | 4.173723 | 10.603698 | 2.645963 |
| H | 3.666384 | 8.653137 | 4.129257 |
| H | 3.967709 | 8.138994 | 2.477978 |
| H | 1.417251 | 8.116075 | 2.112930 |
| H | 1.220907 | 8.488315 | 3.822201 |
| H | 0.896287 | 6.089927 | 3.445657 |
| H | 2.335202 | 6.334428 | 4.440029 |
| H | 7.760753 | 6.754577 | -2.537706 |
| H | 9.130081 | 5.870182 | -1.851752 |
| H | 8.873645 | 5.887682 | -3.600610 |
| H | 6.714136 | 4.612356 | -3.298489 |
| H | 6.955198 | 4.595847 | -1.563000 |
| H | 9.075877 | 3.246154 | -1.882473 |
| H | 8.838414 | 3.300078 | -3.615139 |
| H | 8.087838 | 1.148616 | -2.875202 |
| H | 6.663306 | 2.085213 | -3.368868 |
| H | 5.591562 | -9.862918 | -1.057732 |
| H | 7.225846 | -9.374330 | -1.519763 |
| H | 6.796055 | -9.522943 | 0.189434 |
| H | 0.191070 | -4.590686 | -0.467371 |
| H | 1.857320 | -3.998428 | -0.444741 |
| H | 1.501694 | -5.632350 | -1.016132 |
| H | 5.860916 | -3.233323 | 7.665887 |
| H | 6.560442 | -1.650831 | 7.287839 |
| H | 6.189767 | -2.069777 | 8.961273 |
| H | 0.965965 | -6.655767 | -4.177065 |
| H | 0.882389 | -5.282744 | -3.063235 |
| H | 1.012475 | -5.004801 | -4.800324 |
| H | -2.536504 | -7.494589 | -3.708089 |
| H | -3.159082 | -5.944676 | -4.306627 |
| H | -1.417235 | -6.175997 | -4.110838 |
| H | -9.621705 | -2.489386 | 3.567554 |
| H | -9.979671 | -0.870687 | 2.934815 |
| H | -9.434638 | -2.158718 | 1.838232 |
| H | -7.836028 | 2.792658 | -0.315186 |
| H | -8.432598 | 2.808216 | -1.985641 |
| H | -9.386108 | 2.026268 | -0.711721 |
| H | 5.606318 | -7.462435 | -1.769581 |
| H | 5.152395 | -7.604887 | -0.081826 |
| H | 7.996691 | -6.991148 | -1.064152 |
| H | 7.495499 | -7.005160 | 0.612356 |
| H | 6.385403 | -5.139498 | -1.540228 |
| H | 7.719007 | -4.706602 | -0.450945 |

| | | | |
|---|---|---|---|
| H | 3.750475 | -2.104480 | 8.495428 |
| H | 4.434506 | -0.538566 | 8.129826 |
| H | 1.051039 | 0.193154 | 1.639591 |
| H | -0.802712 | 6.830918 | -1.406637 |
| H | -0.622304 | 4.038312 | -0.750467 |
| H | 0.463662 | 6.609566 | -0.207981 |
| H | 0.273563 | 2.559366 | -2.065819 |
| H | 0.731886 | 3.342033 | -3.576324 |
| H | 3.204060 | -6.403783 | -3.155955 |
| H | 3.296420 | -5.967473 | -4.861781 |
| H | 3.087732 | -3.944937 | -2.568303 |
| H | 4.576485 | -4.425196 | -3.385310 |
| H | -7.608542 | 0.294553 | -0.409425 |
| H | -8.224306 | 0.286455 | -2.064205 |
| H | -7.548749 | -0.544892 | 2.344169 |
| H | -7.753918 | -0.849682 | 4.070523 |
| H | -3.480616 | -6.118806 | -1.812958 |
| H | -1.746385 | -6.390126 | -1.601506 |
| H | 1.162624 | -2.204902 | 3.784763 |
| H | 3.570674 | -0.887205 | 3.258883 |
| H | 1.807239 | -2.653877 | 6.638842 |
| H | 7.358041 | 2.175065 | -0.510134 |
| H | 4.651809 | 0.162755 | 0.465467 |
| H | 4.464623 | -0.366675 | -1.789280 |
| H | 5.744672 | 0.158178 | -2.913016 |
| H | 3.558121 | 6.185973 | 2.148217 |
| H | 3.263146 | 5.181897 | 0.037069 |
| H | 0.934647 | 3.195994 | 1.462522 |
| H | 0.919756 | 3.897888 | 3.057244 |
| H | 5.838408 | -5.332188 | 1.351109 |
| H | 4.476622 | -2.236111 | -0.890424 |
| H | 5.912402 | -3.144453 | -1.404931 |
| H | 3.812693 | -2.434028 | 1.248376 |
| H | 2.882620 | -1.289416 | -4.858351 |
| H | 2.875980 | -1.772466 | -3.190790 |
| H | -2.972921 | 2.011364 | -0.568110 |
| H | -3.502512 | 3.581360 | -0.480236 |
| H | -2.954471 | 2.446132 | 4.286589 |
| H | -2.566652 | 0.844043 | 3.648760 |
| H | -0.261839 | 1.614970 | 3.460671 |
| H | -1.254188 | 2.529002 | 2.583054 |
| H | -1.886816 | 1.385149 | 5.206179 |
| H | 0.007091 | -1.855401 | 2.182546 |
| H | 2.232294 | 1.518306 | -2.967487 |
| H | -4.533845 | 2.321637 | -1.098426 |
| H | 2.827296 | 3.541013 | 0.068857 |
| H | -1.153523 | 5.366940 | 1.260787 |
| H | 1.186216 | -4.671212 | 1.903949 |
| H | 0.728177 | -6.274411 | 1.329110 |
| H | 5.490014 | 1.637691 | 0.810592 |
| H | 3.859928 | -4.062262 | 1.921802 |
| H | 5.199181 | -0.630861 | 5.225468 |
| O | 0.405914 | -2.486192 | 3.094120 |
| H | -0.349009 | -2.932206 | 3.508012 |
| H | 2.423334 | -5.804708 | 1.312880 |
